# spAlignDE unifies cross-sample and cross-modal spatial alignment with mismatch-aware differential expression

**DOI:** 10.64898/2026.09.05.749632

**Authors:** Su Xu, Yu Wang, Lingzhen Meng, Aarya Dalal, Yuxin Yin, Dongyuan Song

## Abstract

Comparative analysis of spatial omics requires aligning data across samples and modalities to a common coordinate system. Existing methods can be computationally intensive for large datasets, and cross-modal alignment is difficult when datasets lack comparable molecular features. In addition, residual alignment errors can cause locations assigned to the same coordinates to represent different biological regions, producing false differential expression signals. Here we propose spAlignDE, a computational method that integrates structure-guided spatial alignment with mismatch-aware local differential expression analysis. spAlignDE represents structures from spatial transcriptomics, spatial ATAC-seq, histology, and anatomical atlases as continuous fields and aligns them by shooting-based diffeomorphic registration without requiring shared molecular features. In cross-sample benchmarks against 12 methods, spAlignDE achieved the highest agreement in gene expression patterns and anatomical annotations. It also scaled to 20 MERFISH brain sections containing 1.45 million cells. For cross-modal tasks, spAlignDE accurately aligned spatial transcriptomics with histology, the Allen Mouse Brain Common Coordinate Framework, and spatial ATAC-seq. After alignment, spAlignDE estimates local expression contrasts on a shared grid and inflates their variances according to mismatch risk estimated from putatively stable genes and local observation density. The analysis can also adjust for cell-type composition. Simulations showed improved false discovery control without systematic loss of power. In real-data applications, spAlignDE localized age-associated changes in gene expression and T-cell distribution in the mouse brain and spatially restricted expression differences between normal and injured kidney sections.

## 1 Introduction

Spatial transcriptomics (ST) technologies measure gene expression while preserving the locations of cells, spots or bins within tissue sections [1–3]. ST links molecular states to tissue architecture and enables studies of cellular organization, local tissue niches and anatomical changes during development and in disease [4, 5]. Emerging spatially resolved assays also measure other modalities, including chromatin accessibility, proteins and metabolites [6–8]. These molecular measurements can be analyzed together with histological images and anatomical atlases [2, 9, 10]. Comparisons across samples, conditions, time points or modalities require placing the corresponding datasets in a common coordinate system. However, tissue sections differ in orientation, scale, sectioning angle, local deformation, and tissue coverage. Therefore, spatial alignment is needed to identify corresponding anatomical locations for cross-sample comparison and multi-sample integration [11–14]. Like image registration, spatial alignment matches corresponding structures in a common coordinate system [15]. The resulting correspondence provides a basis for atlas-label transfer, cross-modal analysis and downstream differential expression analysis [16–18].

Existing alignment methods establish correspondence using optimal transport [11, 19, 20], probabilistic latent-coordinate models [13, 21], graph- and representation-learning approaches [12, 14, 22], or feature-based and continuous-field registration [16, 23, 24]. These methods have expanded spatial alignment beyond adjacent sections to include biological replicates, cross-platform datasets and datasets measured using different modalities. However, a recent benchmarking study found that performance varies substantially across datasets and evaluation criteria, and several limitations remain [25]. First, computational demands can restrict the use of existing methods for large, cell-resolved datasets or studies with many samples. Second, even close geometric overlap does not necessarily establish biologically meaningful correspondence. Overly flexible transformations may distort local cell neighborhoods, spatial domains or anatomical boundaries. They may also force regions present in only one sample into unsupported correspondence. Third, cross-modal alignment remains difficult when datasets lack directly comparable molecular features. Existing approaches often focus on specific modality pairs, such as ST with histology, anatomical atlases or spatial assay for transposase-accessible chromatin using sequencing (spatial ATAC-seq), rather than providing a unified framework for these modalities [17, 20, 24, 26]. Finally, recent methods can provide spatially resolved measures of correspondence confidence during alignment, but these measures do not necessarily indicate whether aligned neighborhoods remain comparable after registration. Residual differences in local anatomy, sampling density or cellular composition can therefore remain difficult to quantify and interpret. A broadly applicable framework should scale to large datasets, preserve biologically meaningful tissue organization without forcing unsupported matches, operate across diverse modalities and quantify the reliability of local correspondence.

Spatial alignment places tissue sections in a common coordinate system and provides the correspondence needed to test differential expression at matched locations, a task we refer to as location-resolved differential expression (“local DE”). Existing comparative methods, however, primarily provide section-wide or domain-level spatial summaries rather than directional hypothesis tests at each matched location. DESpace2 aggregates counts within annotated spatial domains for each sample and tests domain-by-condition interactions to identify genes whose domain-level expression patterns differ across conditions, followed by tests within individual domains [27]. CODA aligns samples, identifies their common tissue region and ranks genes by spatial cross-correlation across sections, distinguishing conserved from divergent spatial distributions [18]. STcompare rasterizes aligned tissues onto matched pixels and uses autocorrelation-preserving permutations to test gene-level spatial correlation. It summarizes spatial fold change with a similarity score, while fold-change maps show where and in which direction expression differs [28]. Together, these methods provide spatially informative results at the section or domain level and, for STcompare, descriptive pixel-level maps, but none of them provides a hypothesis test at each matched location. Such location-resolved inference is difficult because spatial alignment is imperfect: locations treated as matched may still differ in anatomy, sampling density or cell-type composition. This post-alignment mismatch increases the variability of local contrasts even when no true local DE exists. Treating estimated correspondences as exact can therefore underestimate uncertainty and inflate false discoveries. Reliable local DE requires quantifying local comparability and assigning greater uncertainty to poorly supported comparisons.

To address these challenges, we introduce spAlignDE, an integrated framework that combines spatial alignment across samples and modalities with mismatch-aware local differential expression analysis after alignment. spAlignDE converts modality-specific tissue structures into comparable, image-like continuous fields that guide shooting-based large-deformation diffeomorphic metric mapping [29]. This representation supports multi-sample ST alignment and cross-modal alignment with histology, anatomical atlases and spatial ATAC-seq without requiring directly comparable molecular measurements across modalities or feature fusion. For local differential expression analysis, spAlignDE estimates expression contrasts on a shared grid and inflates their variance according to mismatch risk quantified from discrepancies in putatively stable gene profiles and local observation density. Poorly supported local comparisons therefore receive lower inferential precision, while the estimated expression contrasts remain unchanged. In cross-sample benchmarks against 12 methods, spAlignDE consistently achieved the highest agreement in spatial gene-expression patterns and anatomical annotations. spAlignDE also scaled to 20 MERFISH brain sections containing 1.45 million cells in 2.43 minutes, excluding preprocessing. In cross-modal benchmarks, spAlignDE yielded high accuracy for ST-histology, ST-atlas and ST-spatial ATAC alignment and outperformed modality-specific competitors. Simulations showed improved false-discovery control without systematic loss of power. Applications to aging mouse brain and kidney injury resolved localized changes in expression and cell organization.

## 2 Results

### 2.1 Overview of the spAlignDE framework

spAlignDE has two complementary components: structure-guided spatial alignment and mismatch-aware local differential expression analysis (Fig. 1). For spatial alignment, spAlignDE represents modality-specific tissue organization as spatial structures and establishes correspondence using shared structure identities across samples or geometric compatibility across modalities. This design supports both cross-sample and cross-modal alignment. In particular, the cross-modal workflow does not require a shared molecular feature space or feature fusion. For local differential expression analysis, spAlignDE quantifies putative residual mismatch after alignment and propagates this risk into inferential uncertainty through location-specific variance inflation, rather than treating the estimated correspondence as fixed. The alignment and statistical models are described in Methods 5.1 and 5.2, with implementation details in Supplementary Methods S2 and S3.

**Fig 1:**
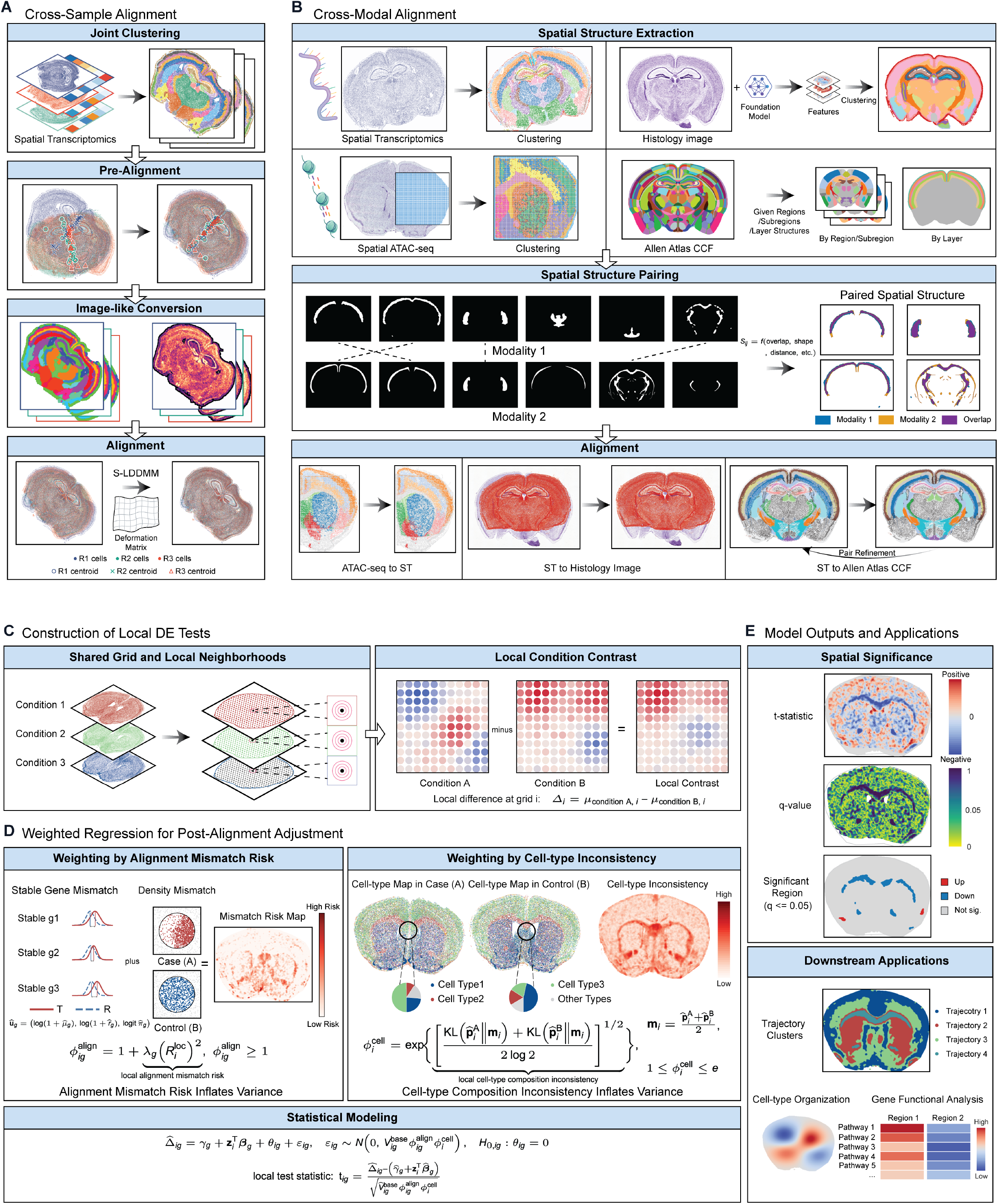
Overview of the spAlignDE framework. A, Cross-sample alignment of spatial transcriptomics data. Joint clustering identifies shared spatial domains. These domains guide global pre-alignment and the construction of image-like representations for registration by shooting-based large deformation diffeomorphic metric mapping (S-LDDMM). **B**, Cross-modal alignment. Spatial structures are extracted from spatial transcriptomics, spatial assay for transposase-accessible chromatin using sequencing (spatial ATAC-seq), histology and anatomical atlases. Structures from different modalities are paired according to spatial and geometric similarity and aligned using S-LDDMM. **C**, Construction of local differential expression tests. Each retained location on the shared grid defines a testing unit. Kernel-weighted neighborhoods of aligned observations provide the condition-specific local means and baseline variance for the local contrast. **D**, Mismatch-aware local regression. Profiles of putatively stable genes characterize the local transformed mean, negative-binomial size and excess-zero probability. Cross-sample discrepancies in these quantities, together with local sampling-density discordance, define a normalized local alignment-mismatch risk. Gene-specific calibration converts this risk into the spatially varying variance factor 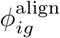.Kernel-smoothed cell-type composition vectors define the optional inconsistency factor 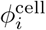. Weighted regression separates each estimated local contrast into a comparison-wide offset, an optional term for confounding variation such as effects of local library size and detection rate, and a local residual. The local statistic standardizes the residual 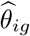 using the baseline variance scaled by 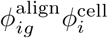. **E**, Model outputs and downstream applications. spAlignDE reports local test statistics, local *q*-values and connected significant regions. These outputs support analyses of spatial trajectories, cell-type organization and pathways.

For multi-sample spatial transcriptomics, spAlignDE identifies spatial structures jointly across sections using gene expression and spatial neighborhood information in a batch-adjusted representation [30, 31]. Structures assigned the same identity establish direct query–reference correspondence (Fig. 1A; Methods 5.1.1). For cross-modal alignment, spAlignDE identifies structures independently in spatial transcriptomics, spatial ATAC-seq and histological images, or obtains them from predefined anatomical atlas compartments (Fig. 1B; Methods 5.1.1). Because some metrics of geometric correspondence are unreliable when samples differ in position, orientation and scale, spAlignDE first performs a global pre-alignment to obtain approximate spatial correspondence (Methods 5.1.2). If automatic initialization is difficult, users can manually adjust the global alignment through an interactive interface (Supplementary Fig. S1; Methods 5.1.2). spAlignDE then represents candidate structures as masks and assigns each query–reference structure pair a composite score based on five complementary metrics of geometric correspondence (Methods 5.1.3). spAlignDE retains only high-confidence candidate pairs that satisfy modality-specific criteria and does not require every structure to have a corresponding match. For atlas alignment, the anatomical hierarchy provides a multiscale library of candidate regions. spAlignDE progressively refines the ST structure resolution while retaining all eligible atlas-hierarchy candidates at every stage (Methods 5.1.6). Users can also specify correspondences through an interactive interface when automatic pairing is ambiguous (Supplementary Fig. S2; Methods 5.1.6).

spAlignDE converts the shared cross-sample structures or accepted cross-modal structure pairs into continuous multichannel fields analogous to multichannel images, with each channel encoding spatial structure. For cross-sample alignment, spAlignDE uses smoothed fields of local structure composition and, when informative, an additional cell-density field for cell-resolved data. A spot-density field is not used for spot-based ST because the spots are evenly distributed and provide no informative spatial variation in density. For cross-modal alignment, spAlignDE uses paired signed-distance-transform fields derived from accepted structure pairs and, when informative, an additional field representing the overall tissue shape. spAlignDE then rasterizes these image-like fields for registration. spAlignDE jointly registers the rasterized fields using shooting-based large deformation diffeomorphic metric mapping (S-LDDMM), which estimates a smooth, invertible transformation with affine and diffeomorphic components [29, 32, 33]. Standard variational LDDMM optimizes a sequence of time-dependent velocity fields [32]. S-LDDMM instead estimates a single initial momentum field that determines the full geodesic trajectory [29]. By reducing the number of optimization variables and memory use, this parameterization improves scalability. The registration model must also accommodate locations without valid counterparts. Such locations can arise from missing tissue, weak overlap or modality-specific structures, and treating them as matched can distort the estimated transformation. spAlignDE therefore adapts the probabilistic missing-data formulation of Tward et al. [34] and fits an expectation–maximization mixture model with one matched component and two unmatched appearance components. The matched-component posterior probability *W*_*M*_ (**x**) weights each location during registration, so locations more consistent with the unmatched components contribute less to transformation estimation. Finally, spAlignDE applies the estimated transformation at the original spatial resolution to cells, spots or bins (Methods 5.1.4 and 5.1.5).

Spatial alignment estimates correspondence between samples, but nominally matched locations may still differ in local anatomy or cell-type organization. As a result, their expression differences may reflect residual mismatch rather than true between-sample changes. spAlignDE addresses this problem by allowing local mismatch risk to increase inferential uncertainty rather than treating the estimated correspondence as exact. spAlignDE constructs a shared testing grid over the aligned tissue sections (Fig. 1C). Each retained grid location is a testing unit, and nearby cells or spots form a kernel-weighted neighborhood for estimating the local expression mean *µ*. For post-alignment inference, sample A denotes the case or condition of interest, and sample B denotes the control or baseline condition, regardless of which sample served as the query or reference during alignment. For gene *g* at location *i*, the estimated directional contrast is

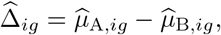

and the working model is

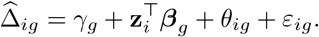

When included, *γ*_*g*_ represents the comparison-wide baseline of the contrast, 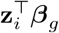 captures optional adjustment for confounding variation such as effects of library size and detection rate, and *θ* _*ig*_ is the location-specific change of interest. With the comparison-wide intercept included, spAlignDE tests whether the local contrast departs from the fitted baseline defined by the comparison-wide contrast and included confounders. The intercept *γ* _*g*_ can absorb approximately uniform between-section shifts. When the intercept is omitted, the test instead evaluates whether the covariate-adjusted local contrast differs from zero. Throughout the post-alignment analyses, *P* denotes a raw *P* value, and *q* denotes the corresponding Benjamini–Hochberg-adjusted *P* value within the testing family specified for each analysis.

Because the true residual mismatch is always unknown, spAlignDE infers it from the data. Similar to the use of stable control genes for batch-effect correction in scRNA-seq [35], spAlignDE quantifies potential mismatch severity from cross-sample discrepancies in the local distributions of putatively stable genes and from local sampling-density discordance (Fig. 1D). These quantities define a normalized local mismatch-risk score 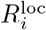. When cell-type annotations are available, an optional adjustment compares the kernel-smoothed local cell-type-composition vectors between the two samples to reflect discrepancies in local anatomical structures directly. The final adjusted local variance is

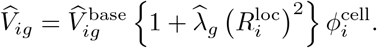

Here, 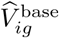 represents the baseline variance of the local comparison. The location-specific score 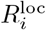 quantifies the risk that residual alignment mismatch makes the comparison at location *i* less reliable, and the gene-specific coefficient 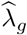 controls how strongly this risk inflates the variance for gene *g*. To estimate 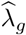 spAlignDE first computes local statistics without mismatch adjustment, groups locations by 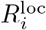, and estimates the excess dispersion of the statistics in each risk group relative to the corresponding Student-*t* null distribution. A non-negative weighted regression of excess dispersion on squared mismatch risk is then fitted; after a bounded reference-bin adjustment, its fitted coefficient gives a contrast-specific estimate of *λ* _*g*_. When multiple contrasts are available, these estimates are robustly combined to obtain 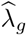.The optional factor 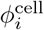 provides a further adjustment for local cell-type-composition inconsistency between samples. These adjustments increase inferential uncertainty where the aligned samples are less biologically comparable but leave the estimated effect and its direction unchanged. spAlignDE reports local *P* values, *q* values and connected significant regions for downstream gene-level and spatial analyses (Fig. 1E). Cells or spots contribute to estimating local spatial patterns but are not treated as independent biological replicates. Full details are provided in Methods 5.2.1–5.2.6.

### 2.2 spAlignDE enables accurate and scalable cross-sample alignment

Accurate cross-sample alignment of spatial transcriptomics data requires both reliable anatomical correspondence across sections and preservation of local tissue organization during deformation. The spAlignDE cross-sample workflow comprises two stages (Fig. 2A). First, spAlignDE identifies shared spatial domains from batch-corrected gene expression and spatial neighborhood information. These domains define a common anatomical representation that guides coarse alignment and subsequent diffeomorphic registration (Methods 5.1.1, 5.1.2, and 5.1.4). Second, spAlignDE uses shooting-based LDDMM to estimate smooth transformations that correct global and local deformations while preserving tissue continuity (Methods 5.1.5). We benchmarked spAlignDE against 12 existing spatial alignment methods [11–14, 16, 18, 19, 21–24, 36] using three datasets that differed in spatial resolution and alignment scale. These datasets included a paired MERFISH mouse brain dataset [37], a paired 10x Genomics Visium mouse kidney dataset comparing normal and ischemia–reperfusion conditions [28, 38, 39], and a multi-sample MERFISH mouse brain aging cohort [40, 41]. Comparator implementations, versions and method-specific settings are documented in Supplementary Methods S2.9. The benchmark covered cell- and spot-resolution platforms, healthy and disease conditions, and both single-pair and large-scale multi-sample alignment.

**Fig 2:**
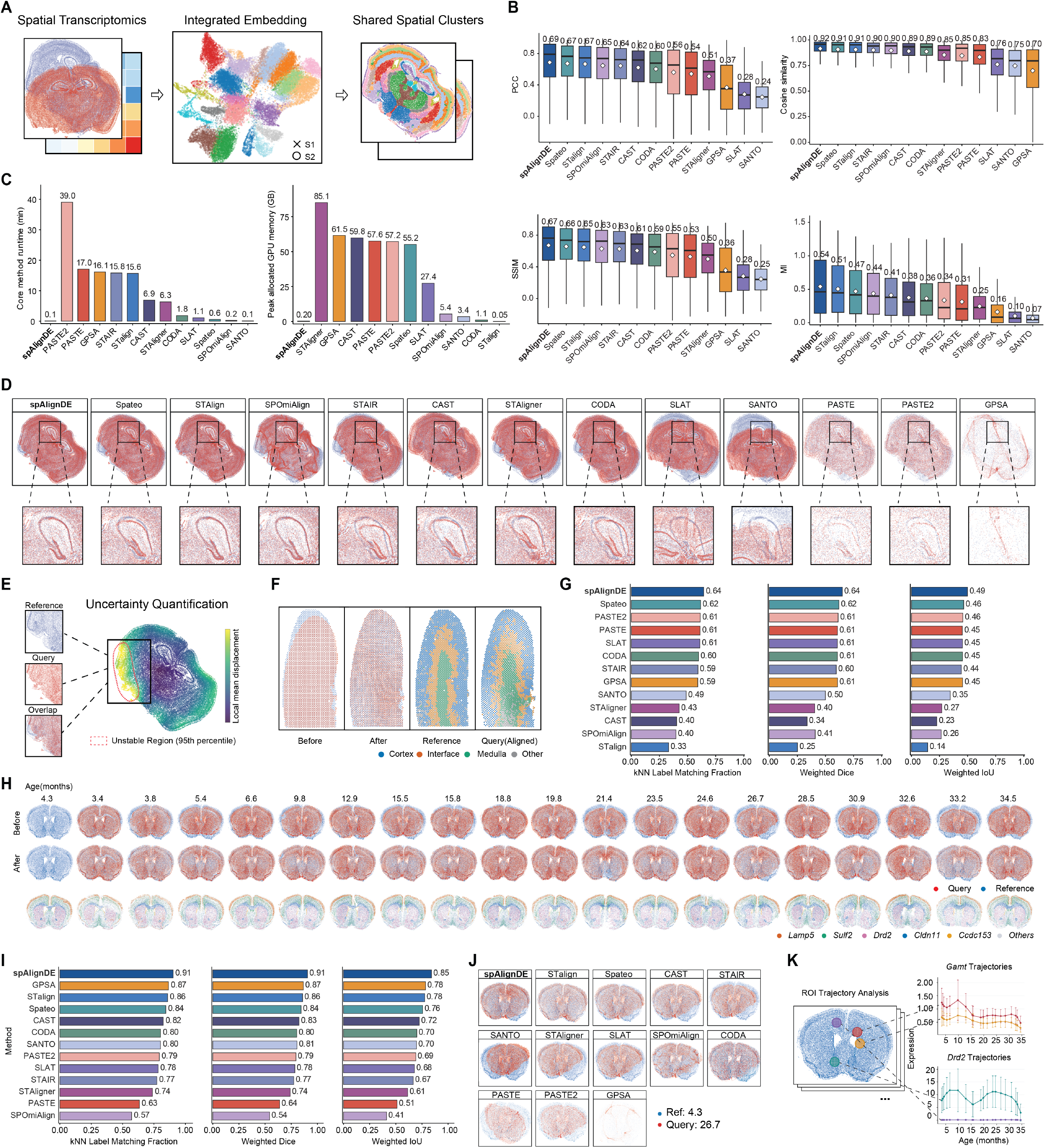
Cross-sample spatial alignment benchmarks and applications. A, The spatial transcriptomics sections were integrated with Harmony and jointly clustered to define shared structures for alignment. Crosses and circles distinguish the two sections in the uniform manifold approximation and projection (UMAP). **B–E**, Vizgen multiplexed error-robust fluorescence in situ hybridization (MERFISH) Mouse Brain Receptor Map benchmark. S2R3 was aligned as the query to the S2R2 reference. **B**, Agreement of spatial gene-expression patterns across 13 methods for 483 shared genes, assessed by the Pearson correlation coefficient (PCC), cosine similarity, structural similarity index measure (SSIM) and mutual information (MI). Boxes indicate interquartile ranges and medians, whiskers extend to 1.5×the interquartile range and diamonds indicate means. Outliers are not shown, and methods are ordered by the mean for each metric. **C**, Core alignment runtime and peak allocated graphics processing unit (GPU) memory. Values are shown above the bars, and methods without recorded memory measurements are omitted. **D**, Whole-section and hippocampal alignment results for all 13 methods (reference, blue; aligned query, red). For **B–D**, PASTE and PASTE2 used 30,000 cells per section, and GPSA used 10,000 cells per section; all other methods used the complete sections containing 85,958 query cells and 84,172 reference cells. **E**, Pointwise transformation stability, measured by mean Euclidean displacement from the repeat-mean aligned position across ten repeated alignments using independent 80% cell subsamples. Contours mark locations above the 95th percentile. **F–G**, 10x Genomics Visium kidney benchmark. The injured IL3 section was aligned as the query to the normal NL3 reference. **F**, Sections before and after alignment with spAlignDE, together with tissue-compartment annotations. **G**, Region agreement across 13 methods, assessed by the *k*-nearest-neighbor label-matching fraction, weighted Dice coefficient and weighted intersection over union (IoU). **H–K**, Aging-brain MERFISH series comprising 20 sections from 3.4 to 34.5 months, all aligned to the 4.3-month reference. **H**, Sections before and after alignment, together with representative aligned gene-expression patterns. **I**, Region-annotation agreement across 19 source-to-reference alignments, assessed using the metrics in **G**. Bars indicate means across alignments. **J**, Alignment results for the 26.7-month section across all 13 methods (reference, blue; query, red). The query lacks the lower-right tissue boundary. **K**, Region-of-interest expression trajectories for *Gamt* and *Drd2* across age.

The paired Vizgen MERFISH mouse brain benchmark included coronal sections from two biological replicates profiled at single-cell resolution, and we aligned S2R3 to S2R2. Following the benchmarking pipeline described in [25], we evaluated the preservation of spatial gene expression patterns across 483 shared genes using the Pearson correlation coefficient (PCC), cosine similarity, structural similarity index measure (SSIM) and mutual information (MI) (Fig. 2B; Methods 5.3.1). spAlignDE achieved the highest mean performance across all four metrics. The rankings of competing methods, however, varied by metric. Because these metrics are calculated after gene expression is aggregated on a spatial grid [25], the results may depend on the degree of spatial averaging. We therefore repeated the evaluation using finer grids. spAlignDE remained top-ranked at both resolutions (Supplementary Figs. S3 and S4). Alignment performance also remained stable across Leiden resolutions that produced 15–27 joint spatial clusters (Supplementary Figs. S5A, B and S6; Methods 5.3.1; Supplementary Methods S2.12). These results support the robustness of spAlignDE to evaluation-grid resolution and the granularity of the spatial structures used for alignment.

We next assessed computational efficiency by measuring core alignment runtime (excluding preprocessing) and peak GPU memory usage (Fig. 2C; Methods 5.5.2; Supplementary Methods S2.10.1). spAlignDE completed the core alignment in 0.1 min, tying for the shortest runtime among the evaluated methods. Its peak GPU memory usage was 0.20 GB, the second-lowest value. The accuracy and efficiency benchmarks therefore show that spAlignDE achieved the highest mean alignment performance while tying for the shortest runtime and ranking second lowest in peak GPU memory usage among the evaluated methods.

We further assessed anatomical preservation at global and local scales by visual inspection (Fig. 2D). spAlignDE matched the overall tissue geometry and boundaries and retained the laminar organization of the hippocampus with minimal local distortion. In contrast, competing methods showed excessive global deformation, residual local mismatch or forced alignment in regions with incomplete tissue support. These observations indicate that spAlignDE preserved anatomical structure at both spatial scales in this benchmark.

Beyond alignment accuracy, spAlignDE quantifies alignment uncertainty by repeating the alignment using independently subsampled cells from both sections (Fig. 2E; Methods 5.3.1; Supplementary Methods S2.11). Most locations showed small mean Euclidean displacement from their repeat-mean aligned positions across repeated alignments, consistent with locally stable transformation estimates. Larger mean displacement was concentrated in areas with incomplete tissue overlap at the bottom-left corner. We marked locations above the 95th percentile to identify regions with reduced transformation stability. These maps identify regions where downstream results should be interpreted with caution.

We next benchmarked spAlignDE on spot-resolution 10x Genomics Visium kidney sections by aligning the ischemia– reperfusion injury section IL3 to the normal section NL3 (Fig. 2F,G). Because injury-associated differences in gene expression make gene-expression concordance unreliable as a measure of alignment accuracy, we did not assess gene-expression concordance. We instead evaluated anatomical correspondence based on overall kidney morphology and annotations for three compartments and other tissue [39]. spAlignDE preserved overall kidney morphology and improved correspondence across the annotated compartments and other tissue. spAlignDE ranked first among all methods for each of the three compartment-label concordance metrics: k-nearest-neighbor (kNN) label matching, weighted Dice and weighted intersection over union (Methods 5.3.1). We subsequently used the aligned sections for local DE analysis (Results 2.5).

We next assessed the scalability and alignment accuracy of spAlignDE using the mouse brain aging MERFISH dataset from Sun et al. [40, 41], which comprised 20 coronal sections and 1,453,144 cells (Fig. 2H–K). We used the 4.3-month section as the reference because it contained the most complete set of anatomical structures and aligned the remaining 19 sections to this reference. We evaluated alignment accuracy for these 19 query-to-reference comparisons using anatomical annotations, spatial overlap and gene-expression concordance (Methods 5.3.1). spAlignDE mapped all sections to a common coordinate system while preserving anatomical organization and representative marker-gene patterns (Fig. 2H). spAlignDE achieved the highest region-level annotation agreement among the 13 methods (Fig. 2I) and remained top-ranked when evaluated using cell-type and subregion annotations (Supplementary Figs. S7 and S8x). spAlignDE also maintained consistent whole-section correspondence across the cohort (Supplementary Fig. S9). Using the default evaluation grid, spAlignDE achieved the highest gene-expression concordance across 300 genes. When we repeated the analysis using finer grids, spAlignDE remained topranked for all four metrics (Supplementary Figs. S10, S11 and S12). spAlignDE completed the 19 core query-to-reference alignments in 2.43 min, with peak allocated GPU memory below 0.1 GB (Methods 5.5.1 and 5.5.2). These results show that spAlignDE scaled well to a large cohort study.

We also evaluated spAlignDE under incomplete spatial coverage in the aging brain cohort. For the 26.7-month query, spAlignDE preserved the lower-right boundary created by incomplete tissue coverage while maintaining overall anatomical correspondence, rather than forcing unsupported overlap with the 4.3-month reference (Fig. 2J; Methods 5.1.5). Mean displacement from the repeat-mean aligned position was elevated in regions with discordant spatial structures or incomplete tissue support (Supplementary Fig. S13; Methods 5.3.1; Supplementary Methods S2.11). This larger mean displacement indicates that the estimated transformation was less stable where correspondence was poorly constrained. The resulting common coordinate system enabled comparison of spatial gene-expression trajectories across age (Fig. 2K). We further analyzed these trajectories in Results 2.5 (Methods 5.2.6 and 5.4.1).

The preceding benchmarks primarily evaluated whole-section alignment between tissues with broadly corresponding anatomy and well-defined spatial domains, even when tissue coverage was incomplete. To examine partial alignment between anatomically heterogeneous tissues, we aligned two consecutive sections from the 10x Genomics Xenium FFPE human breast cancer dataset [42], using manual pre-alignment for global initialization (Methods 5.1.2). Using the default evaluation grid, spAlignDE ranked first across all four gene-pattern metrics. At finer grid resolutions, spAlignDE remained close to the top-performing method (Supplementary Figs. S14–S16; Methods 5.3.1). Spatial overlays showed close correspondence within the shared tissue without forcing non-overlapping regions to match (Supplementary Fig. S17). Thus, spAlignDE can accommodate partial overlap between anatomically heterogeneous tissue sections when an appropriate global initialization is available.

### 2.3 spAlignDE enables structure-guided cross-modal alignment without requiring shared molecular features

Cross-modal alignment is more challenging than cross-sample alignment because the datasets may not share directly comparable molecular features. spAlignDE therefore establishes cross-modal correspondence through spatial structures rather than a shared molecular feature space. spAlignDE first identifies modality-specific structures from spatial transcriptomics, spatial ATAC-seq, histological images or anatomical atlases (Methods 5.1.1). After global pre-alignment, spAlignDE represents these structures as masks and pairs query and reference structures according to their geometric compatibility (Methods 5.1.2 and 5.1.3). spAlignDE then converts the accepted structure pairs into matched continuous fields and aligns these fields using SLDDMM (Fig. 3A; Methods 5.1.4 and 5.1.5). We evaluated this workflow by aligning spatial transcriptomics with histology, an anatomical atlas and spatial ATAC-seq.

**Fig 3:**
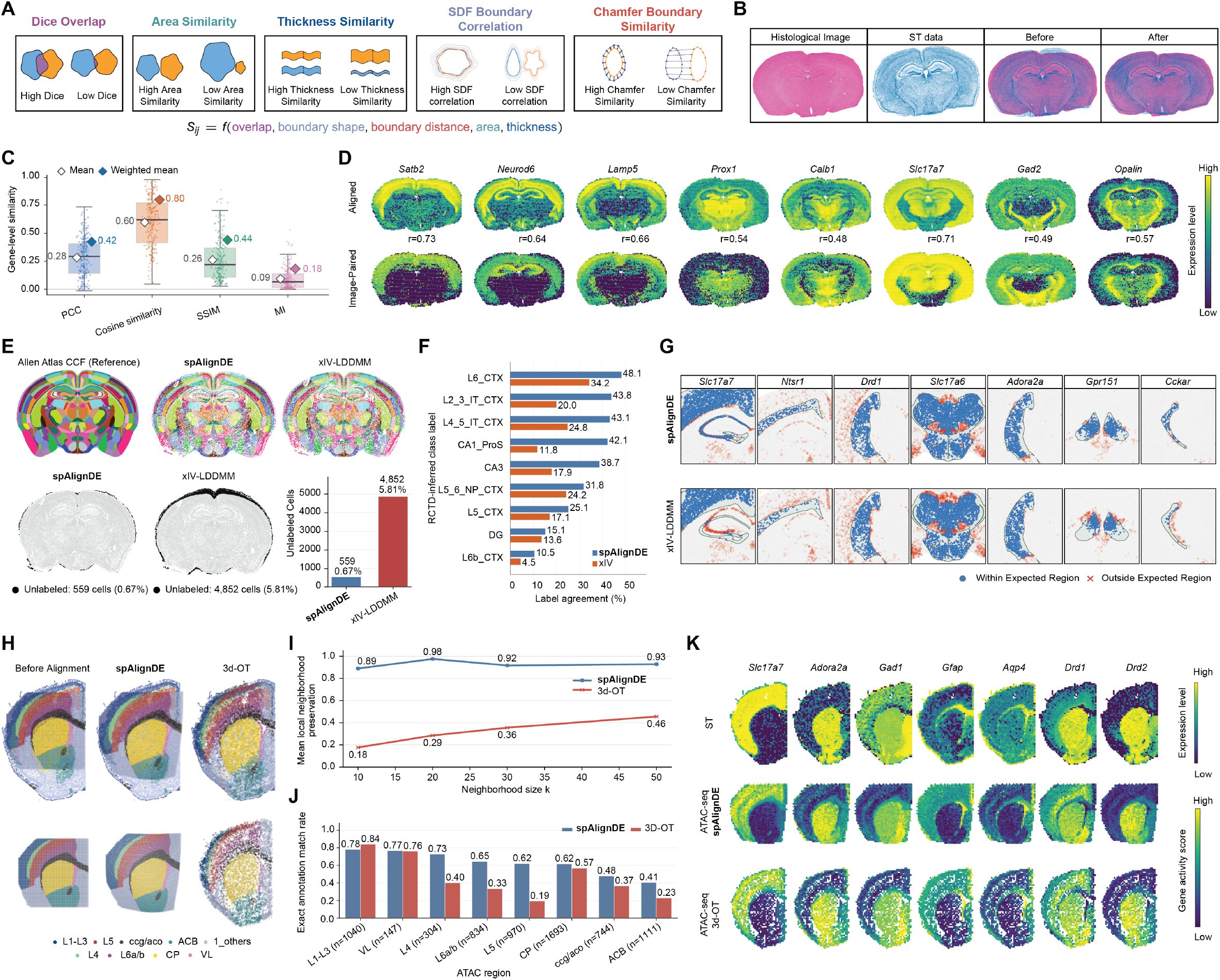
Cross-modal spatial alignment benchmarks. A, Candidate spatial structures are paired according to a modality-specific score, *s*_*ij*_, that combines overlap, boundary geometry, area and thickness. **B–D**, Alignment of Xenium spatial transcriptomics (ST) data to a hematoxylin and eosin (H&E)-stained image. **B**, H&E image, Xenium section and overlays before and after alignment with spAlignDE. **C**, Spatial concordance between aligned Xenium expression and Visium expression paired with the reference H&E image. Each dot represents one of *n* = 246 shared genes evaluated across 5,091 matched bins using the Pearson correlation coefficient (PCC), cosine similarity, structural similarity index measure (SSIM) and mutual information (MI). Boxes indicate interquartile ranges and medians; open and filled diamonds indicate unweighted and reliability-weighted means, respectively. **D**, Representative expression patterns in the aligned Xenium data and the H&E-paired Visium data. **E–G**, Alignment of ST data to the Allen Mouse Brain Common Coordinate Framework version 3 (CCFv3), compared with xIV-LDDMM. **E**, Atlas alignment and label transfer. The fixed-seed automatic workflow retained 18 matched spatial-structure pairs. Cells mapped outside the annotated atlas are shown in the bottom row. xIV-LDDMM left 4,852 cells unlabeled (5.81%), compared with 559 cells (0.67%) for spAlignDE. **F**, Agreement between transferred atlas labels and RCTD-inferred class labels. For each anatomically informative RCTD-inferred class, bars show the percentage of eligible ST cells assigned to one of its predefined compatible cortical, hippocampal or dentate-gyrus regions. **G**, Localization of regionally enriched marker genes. Blue circles and red crosses denote high-expression cells mapped inside and outside their expected atlas regions, respectively; outlines mark the expected regions. **H–K**, Alignment of a partial spatial assay for transposase-accessible chromatin using sequencing (spatial ATAC-seq) section to ST, compared with 3d-OT. **H**, Tissue overlays before and after alignment. **I**, Mean neighborhood preservation among 20-*µ*m ATAC-seq pixels. For each pixel, neighborhood preservation is the fraction of its original *K*-nearest neighbors retained after alignment. **J**, Agreement between the original ATAC-seq anatomical annotations and labels inferred from the ten nearest ST observations after alignment. **K**, Representative ST expression and ATAC-derived gene-activity patterns after alignment with spAlignDE or 3d-OT.

We first aligned a Xenium mouse brain section to an H&E image paired with Visium gene-expression measurements [43, 44] (Fig. 3B). We withheld the Visium measurements from transformation estimation and used them only for independent evaluation. Using spatial structures identified from the histology image, spAlignDE refined the initial global alignment with S-LDDMM and improved anatomical correspondence (Fig. 3B and Supplementary Fig. S18; Methods 5.1.1, 5.1.2, 5.1.4 and 5.1.5). We evaluated the resulting alignment by comparing spatial gene-expression patterns between the aligned Xenium data and the held-out Visium measurements. Across 246 shared genes, PCC, cosine similarity, SSIM and MI indicated molecular concordance between these data (Fig. 3C; Methods 5.3.1). Reliability weighting increased mean similarity for all four metrics; mean cosine similarity increased from 0.60 to 0.80 (Supplementary Methods S2.10.2). Representative genes had spatial PCC values of 0.48–0.73 (Fig. 3D). To evaluate alignment with a different histological stain, we aligned a MERFISH mouse brain section to a Nissl-stained section from the Allen Mouse Brain Atlas [9, 37]. S-LDDMM refinement improved correspondence of the tissue boundary and internal cytoarchitecture (Supplementary Fig. S19; Methods 5.1.1, 5.1.2, 5.1.4 and 5.1.5). These analyses show that spAlignDE can align Xenium and MERFISH data to H&E and Nissl images, respectively, without requiring shared molecular features.

Next, we evaluated alignment to the Allen Mouse Brain Common Coordinate Framework version 3 (CCFv3), a threedimensional reference atlas of the adult mouse brain with hierarchically organized anatomical regions [10]. Because the MERFISH data represent a single tissue section, we used a two-dimensional coronal section of CCFv3 at the corresponding anatomical depth. We aligned the MERFISH section to this atlas section and transferred region- and subregion-level atlas labels to the ST cells (Fig. 3E). Across three stages, spAlignDE progressively refined the ST structure partition while retaining all eligible atlas hierarchy candidates from depths 2–10 at each stage, and then applied iterative S-LDDMM refinement (Methods 5.1.6 and 5.1.5). The automatic workflow retained 18 matched ST–atlas structure pairs. We compared spAlignDE with xIV-LDDMM, a task-specific method for cross-modal atlas-to-spatial-omics registration [17, 26] (Supplementary Methods S2.9). spAlignDE more closely reproduced the whole-section geometry and local anatomical organization of the CCFv3 reference. In contrast, xIV-LDDMM showed larger discrepancies in tissue boundaries, section size and local deformation (Fig. 3E, top). After atlas labels were projected back to the native ST coordinates, spAlignDE left 559 cells unlabeled (0.67%), compared with 4,852 cells (5.81%) for xIV-LDDMM (Fig. 3E, bottom).

We then evaluated whether the transferred atlas labels were consistent with the MERFISH gene-expression profiles. Using RCTD with the Allen mouse cortex and hippocampus single-cell dataset as the reference, we independently assigned transcriptomic subclasses to the MERFISH cells [45–47]. We compared these expression-based subclasses with the corresponding broad CCFv3 regions assigned after alignment. spAlignDE showed higher agreement between the two label sets in all nine evaluated cortical, hippocampal and dentate gyrus categories (Fig. 3F; Methods 5.3.1). Across seven brain region marker genes, spAlignDE produced higher expected-region localization than xIV-LDDMM for all seven genes (Fig. 3G) [17, 48]. The marker-gene patterns were more concentrated within the expected atlas regions and showed less leakage across anatomical boundaries. These results show that spAlignDE improved both geometric atlas alignment and the biological consistency of atlas-label transfer in this comparison.

To assess whether atlas-alignment performance extended beyond the sample S2R1, we aligned two additional MERFISH sections, S1R1 and S3R1, to the corresponding coronal sections of Allen CCFv3 (Methods 5.1.5 and 5.1.6). For both sections, spAlignDE achieved close correspondence in overall tissue shape and local anatomical regions and transferred atlas labels to the MERFISH cells (Supplementary Figs. S20 and S21). The results for S1R1 and S3R1 indicate that the atlas-alignment workflow generalizes across independently profiled sections that differ in coronal depth and tissue geometry.

Matched structure pairs provide the anatomical constraints used to estimate the transformation, so alignment performance may depend on the number of pairs retained. We therefore assessed sensitivity to the number of matched pairs while holding the other components of the alignment procedure fixed. The automatic atlas-alignment workflow retained 18 pairs in S2R1. To isolate the effect of pair number, we used a single final-resolution S-LDDMM rerun from the same global pre-alignment with all 18 pairs as the full-pair baseline. We then independently repeated this step using the top-ranked 16, 14, 12, 10 or 8 pairs (Methods 5.3.1; Supplementary Methods S2.12). Across cortical, hippocampal and dentate gyrus categories, label agreement remained stable as the number of pairs decreased, and regionally enriched marker genes remained localized to their expected atlas regions (Supplementary Figs. S22 and S23). Because this sensitivity analysis varied the number of automatically selected pairs but not their identities, we separately evaluated an interactive interface that allows users to specify structure correspondences when automatic pairing is uncertain but anatomical knowledge is available (Methods 5.1.6; Supplementary Methods S2.5). The alignment based on UI-defined correspondences closely matched the Allen CCFv3 reference and achieved an overall label-agreement rate comparable to that of the automatic workflow, at 37.8% and 36.3%, respectively, with only modest category-specific differences (Supplementary Figs. S24 and S25). Marker-gene localization was also similar between the two workflows (Supplementary Fig. S26). In summary, alignment remained stable over the tested range of automatically selected structure pairs, and user-specified correspondences yielded results comparable to those obtained with automatic pairing.

Finally, we evaluated alignment across modalities and spatial resolutions by aligning a spatial ATAC-seq section covering part of the mouse brain, profiled at 20-*µ*m pixel resolution, to a cell-resolved MERFISH ST reference [6, 37]. Because the ATAC-seq section covered only a restricted brain region, we used the corresponding half-brain region of the MERFISH data as the reference (Methods 5.1.1–5.1.5). We compared spAlignDE with 3d-OT, a method for heterogeneous spatial multi-omics alignment [20] (Supplementary Methods S2.9). 3d-OT placed the query within the correct broad brain region but expanded the ATAC-seq pixels across most of the half-brain reference, altering the partial-tissue geometry and disrupting their internal spatial organization. In contrast, spAlignDE retained the compact query geometry and local structure (Fig. 3H). To quantify preservation of local spatial organization, we calculated the fraction of each pixel’s original *K*-nearest ATAC-seq neighbors retained after alignment. Across the evaluated values of *K*, mean neighborhood preservation ranged from 0.89 to 0.98 for spAlignDE and from 0.18 to 0.46 for 3d-OT (Fig. 3I; Methods 5.3.1). We next evaluated biological correspondence using the published anatomical annotations and gene-activity scores for the ATAC-seq pixels and the same coarse anatomical categories assigned to the ST reference from marker patterns [6, 49]. After alignment, each ATAC-seq pixel was assigned an ST-derived label by inverse-distance-weighted voting among its ten nearest ST cells. spAlignDE yielded a higher exact annotation match rate than 3d-OT (62.5% versus 49.6%) and higher match rates in seven of the eight anatomical categories (Fig. 3J). For representative genes, ATAC-derived gene activity and ST expression showed concordant spatial patterns after alignment with spAlignDE, while the ATAC-seq section retained its native geometry (Fig. 3K). In this cross-modal comparison, spAlignDE recovered anatomical and molecular correspondence while preserving the original spatial organization of the ATAC-seq section.

### 2.4 Mismatch-aware post-alignment testing improves false-discovery control in location-resolved comparisons

Even after spatial alignment, locations assigned to the same coordinates may remain biologically non-equivalent, which can produce spurious local expression differences. We therefore evaluated the mismatch-aware local differential expression component of spAlignDE. This component tests expression differences at each location on a shared grid and incorporates residual post-alignment mismatch through local variance inflation. To determine whether this adjustment improves post-alignment inference, we constructed semi-synthetic datasets with known local differential expression regions using a real mouse brain spatial template (Fig. 4A). Simulation sample A served as the baseline reference. Simulation sample B contained spatially localized up- and down-regulated regions and underwent global and local deformations before alignment. We defined the post-alignment contrast as sample B minus sample A. This design retained realistic tissue geometry and spatial sampling while providing known local differential expression regions [50] (Methods 5.3.2; Supplementary Methods S3.10). Because the local differential expression model can be applied to the output of any alignment algorithm, we compared its naive and mismatch-aware modes across alignments produced by spAlignDE and nine other alignment methods. The naive mode treats the aligned coordinates as exact. The mismatch-aware mode instead incorporates residual mismatch into the local variance (Methods 5.2.1–5.2.5).

**Fig 4:**
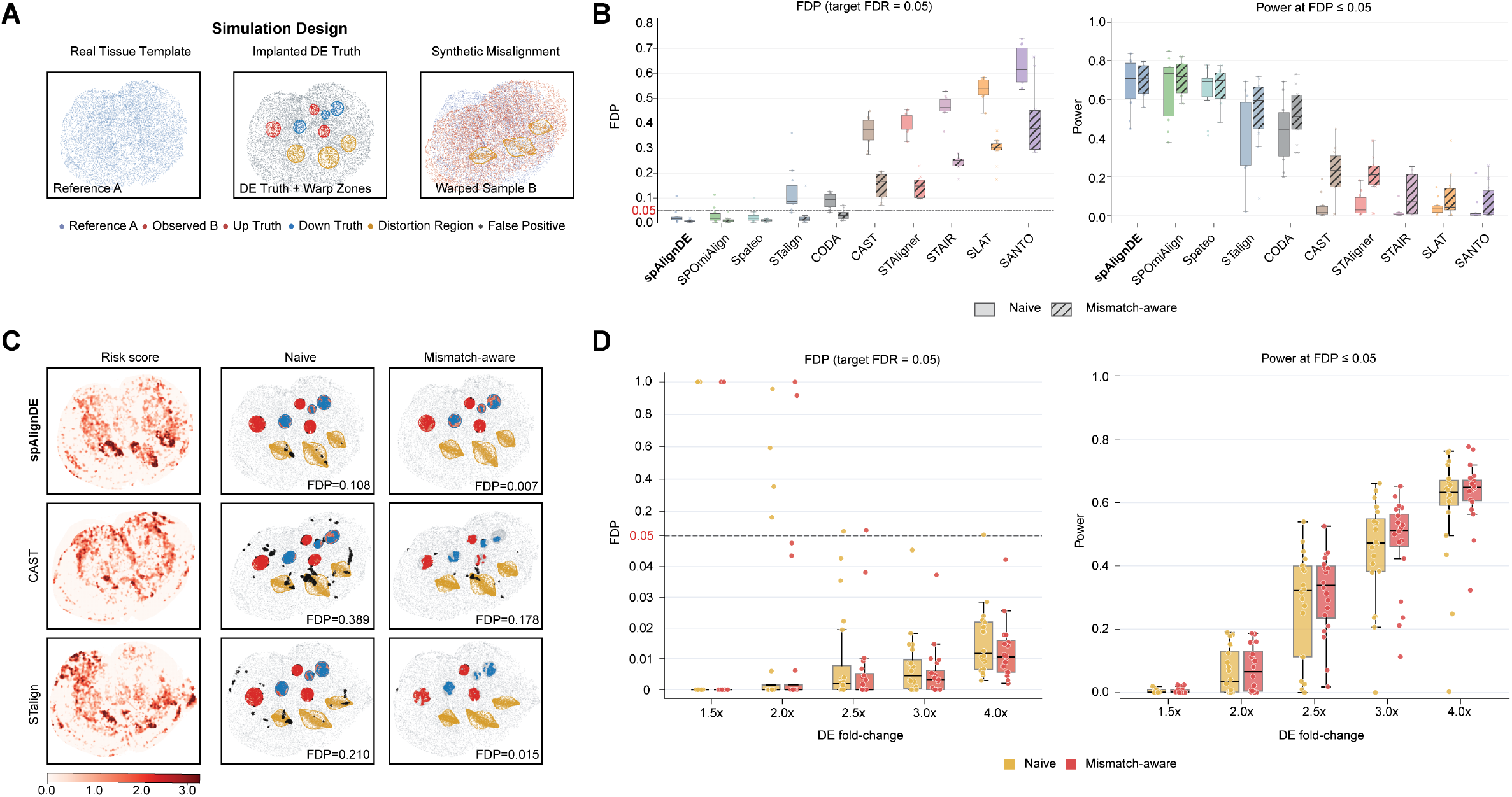
Simulation benchmark of mismatch-aware post-alignment spatial testing. A, Simulation design based on a real mouse brain spatial transcriptomics section. Spatially localized upregulated and downregulated regions were implanted together with independent distortion regions, and sample B was warped to introduce synthetic misalignment. The alignment methods received no information about the implanted truth. Post-alignment calls were evaluated against the known differential expression regions. **B**, Comparison of naive and mismatch-aware testing across ten alignment methods and ten matched simulation groups. Left, location-level FDP for calls made at a target FDR of 0.05; the dashed line indicates the target. Before evaluation against the implanted truth, significant grid calls and local statistics were projected onto cells in simulation sample B. Right, power at FDP ≤0.05, defined as the maximum fraction of correctly signed true DE cells recovered among q-value-ranked call sets whose location-level FDP did not exceed 0.05. Solid and hatched boxes indicate naive and mismatch-aware testing, respectively, and colors identify alignment methods. **C**, Representative results for alignments produced by spAlignDE, CAST and STalign, shown from top to bottom. Columns show the non-negative robust-standardized mismatchrisk score 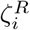, naive testing and mismatch-aware testing. The displayed value of 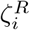is the score before 95th-percentile capping and rescaling and therefore has no fixed upper bound. Only the rescaled local score 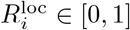 enters the variance-inflation model. Significant grid calls are projected onto sample-B cells. Red and blue points denote correctly signed true positives in upregulated and downregulated regions, respectively; orange points denote implanted distortion regions, black points denote false positives and gray points denote all cells. Displayed values are location-level FDP for calls made at a target FDR of 0.05. **D**, Sensitivity to differential expression signal strength. At each signal level, 20 matched simulation templates were evaluated. The implanted effect was scaled to 1.5×, 2.0×, 2.5×, 3.0× or 4.0× the baseline effect, with all other settings held fixed. Boxplots show location-level FDP at a target FDR of 0.05 and power at FDP ≤ 0.05, as defined in **B**. For **B** and **D**, boxes indicate interquartile ranges, center lines indicate medians and whiskers extend to 1.5× the interquartile range.

Across ten matched simulation groups, we first evaluated naive post-alignment inference for alignments produced by spAlignDE and the nine other methods (Fig. 4B; Methods 5.3.3). We projected grid-level calls onto cells in sample B and compared them with the implanted truth, requiring true-positive calls to have the correct direction. Naive inference performed better when the underlying alignment was more accurate. At a target FDR of 0.05, alignments produced by spAlignDE yielded a near-zero median FDP. When power was evaluated at FDP ≤ 0.05, the median power was approximately 0.7. SPOmiAlign and Spateo also yielded low FDP and high power, whereas methods with less accurate alignment had markedly inflated FDP and reduced power. This ordering broadly agreed with the independent alignment benchmark (Results 2.2), showing that residual correspondence errors can compromise local differential expression inference. We then compared naive inference with mismatch-aware adjustment for each alignment method. The mismatch-aware adjustment reduced FDP across all methods, with modest reductions for accurate alignments that left little residual mismatch and substantially larger reductions for less accurate alignments (Fig. 4B). For example, mismatch-aware adjustment helps STalign and CODA control the FDP under the 0.05 cutoff compared to their naive results. At the same FDP threshold, power was generally maintained and increased for some methods. However, several poorly aligned methods remained above the nominal FDP target after adjustment. Thus, mismatch-aware inference mitigated residual alignment errors but could not fully compensate for inaccurate spatial correspondence.

We then examined a representative simulation to determine where mismatch risk was elevated and how it related to falsepositive calls. The non-negative, robustly standardized mismatch-risk score 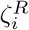 varied across the tissue. False-positive calls from the naive analysis were concentrated in high-risk regions, which overlapped the implanted distortion regions (Fig. 4C and Supplementary Figs. S27 and S28). For visualization, the maps display 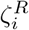 after median–MAD standardization and flooring at zero but before 95th-percentile capping and rescaling to 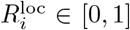. Unlike the raw cosine dissimilarity *M*_*i*_ ∈ [0, 2], 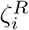 has no fixed upper bound (Supplementary Methods S3.3). Mismatch-aware testing removed many false-positive calls while retaining most implanted up- and down-regulated regions. FDP decreased from 0.108 to 0.007 for spAlignDE, from 0.210 to 0.015 for STalign and from 0.389 to 0.178 for the more severely misaligned CAST output. Consistent with the spatial maps, high mismatch-risk scores were enriched in the implanted distortion regions (Supplementary Fig. S29A). In an ablation analysis, the observed risk map reduced FDP relative to naive testing. Random permutation of the same risk values across grid locations produced an FDP close to that of the naive analysis (Supplementary Fig. S29B). This comparison indicates that the improvement depended on the observed spatial association between mismatch risk and excess variability in the initial local statistics, rather than on variance inflation alone. We also examined controlled coordinate perturbations. Under the perturbation scheme shown in Supplementary Fig. S30, shared-grid inference was more stable than cell-centered local testing (Supplementary Fig. S31), and mismatch-aware adjustment further reduced FDP relative to naive shared-grid testing (Supplementary Fig. S32). The adjustment was most effective when most of the tissue was well aligned but localized mismatch remained. We next assessed robustness to differential expression signal strength and deformation severity (Methods 5.3.2; Supplementary Methods S3.10). Across 20 matched simulation templates at each signal level, power increased with the strength of the implanted effects. Mismatch-aware testing maintained a low median FDP at the target FDR of 0.05 and power comparable to that of the naive analysis at FDP ≤ 0.05 (Fig. 4D). Increasing the severity of local deformation produced the same qualitative pattern (Supplementary Fig. S33). These results show that spatially resolved mismatch estimation improves post-alignment error control without systematic loss of power, particularly when the underlying alignment is sufficiently accurate.

### 2.5 Mismatch-aware inference identifies localized changes in aging brain and injured kidney

After evaluating statistical performance in simulations, we applied mismatch-aware post-alignment inference to two datasets: a cell-resolved MERFISH dataset of aging brain samples and a spot-based Visium dataset of injured kidney samples. The aging mouse brain dataset comprised 20 coronal sections spanning distinct ages and comparable anatomical levels [40]. We selected the 4.3-month section as the reference because it retained the most complete anatomy and aligned the remaining 19 sections to it (Methods 5.4.1; Supplementary Methods S3.11). The original study reported a broad age-associated decline in *Gamt* expression [40]. To determine the spatial distribution of this decline, spAlignDE estimated local *t*-statistics and *q*-values at each shared-grid location and grouped contiguous significant locations into regions (Fig. 5A; Methods 5.2.5; Supplementary Methods S3.2). *Gamt* down-regulation became more pronounced at later ages and was concentrated in an arch-shaped white-matter region along the corpus callosum.

**Fig 5:**
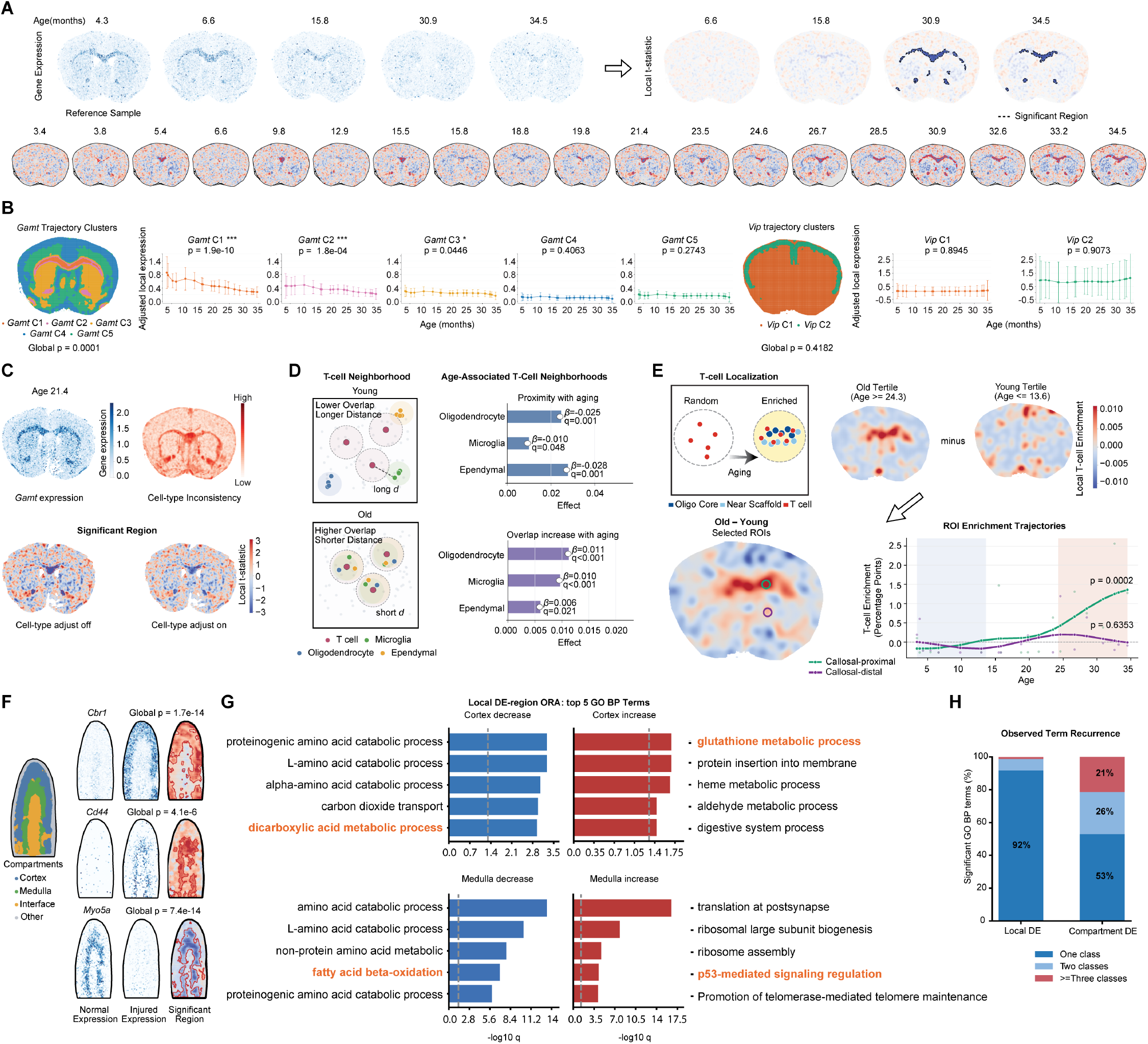
Mismatch-aware post-alignment inference in aging brain and injured kidney. A, Local differential expression across 20 aging mouse brain sections profiled by multiplexed error-robust fluorescence in situ hybridization (MERFISH), relative to the 4.3-month reference. Representative *Gamt* expression and local *t*-statistic maps are shown above; results for all 19 contrasts are shown below. **B**, Gene-level spatial Aggregated Cauchy Association Test (ACAT) results for age trends and spatial expression trajectories for *Gamt* (*P* = 1.1× 10^−4^) and *Vip* (*P* = 0.4182). The gene-level test fits a local linear age trend at each grid location and is performed without trajectory clustering. Points show cluster means, and bars indicate the mean ±1.96 spatial standard deviations; labels give raw two-sided Wald *P* values for cluster-level age trends. **C**, *Gamt* at 21.4 versus 4.3 months: expression, normalized cell-type-composition inconsistency and local statistics without and with in one, two or at least three classes. Contours mark connected regions with *q* ≤ 0.05 in **A** and **F** and with *q* ≤ 0.10 in **C**. cell-type adjustment. **D**, Age-associated changes in T-cell proximity to and spatial overlap with oligodendrocytes, microglia and ependymal cells. Positive *β* denotes decreasing distance or increasing spatial overlap with age. **E**, Local T-cell enrichment in young (≤13.6 months; *n* = 7) and old (≥24.3 months; *n* = 7) sections, the difference between the two age groups and enrichment trajectories in callosal-proximal and callosal-distal regions. **F**, Expression and mismatch-aware local differential expression in the normal NL3 and injured IL3 kidney sections. Labels indicate raw gene-level ACAT *P* values. **G**, The five highest-ranked Gene Ontology (GO) Biological Process terms for each displayed cortex or medulla compartment–direction class. The analysis included genes that passed gene-level Benjamini–Hochberg adjustment across 16,446 genes and were assigned to classes based on their local differential expression regions. Terms were ranked by increasing term *q*-value. Ties were resolved successively by increasing raw over-representation *P* value, decreasing overlap gene count and alphabetical GO term name. Bars show ™ log_10_(*q*), where *q* is the Benjamini–Hochberg-adjusted term *P* value; dashed lines mark *q* = 0.05, and orange labels denote kidney-injury-related terms annotated after ranking. **H**, Recurrence of significant terms across compartment–direction classes for local and compartment-level differential expression. Colors indicate whether terms occur in one, two or at least three classes. Contours mark connected regions with *q* ≤ 0.05 in **A** and **F** and with *q* ≤ 0.10 in **C**.

To summarize age-related evidence at the gene level, we fitted a linear age trend to the unsmoothed adjusted expression at each grid location using mismatch-aware precision weights. We then combined the resulting local trend *P* values across space using the Aggregated Cauchy Association Test to obtain a gene-level *P* value (ACAT; Methods 5.2.6; Supplementary Methods S3.9). *Gamt* showed evidence of an age-associated spatial trend (*P* = 1.1 ×10^−4^), but *Vip* did not (*P* = 0.4182; Fig. 5B). The *Vip* result was consistent with a previous mouse cortex study that reported little age-related change in *Vip* transcript or VIP protein abundance [51]. To characterize where and how expression changed with age across the tissue, we grouped locations with similar adjusted expression trajectories and tested the age trend within each spatial cluster using spatial-variability-weighted regression (Methods 5.2.6). The automatic procedure selected *K* = 5 for *Gamt* and *K* = 2 for *Vip*. For *Gamt*, clusters C1–C3 yielded two-sided Wald *P* values below 0.05 (*P* = 1.9 ×10^−10^, 1.8 ×10^−4^ and 0.0446, respectively). Clusters C4 and C5 showed no evidence of an age trend (*P* = 0.4063 and 0.2743). Neither *Vip* cluster showed evidence of an age trend (*P* = 0.8945 and 0.9073). Because the cluster number and spatial partition were selected from the observed trajectories, these raw Wald *P* values should be interpreted as conditional summaries. As a sensitivity analysis, we fixed *K* at 2, 4, 6 and 8; the resulting partitions mainly subdivided the same broad spatial domains (Supplementary Fig. S34).

Because sections may sample slightly different anatomical depths or contain incompletely preserved tissue, aligned neighborhoods can differ in cell-type composition; we therefore examined spAlignDE’s optional variance adjustment for this inconsistency. The adjustment reduces inferential precision at locations where the two neighborhoods differ strongly in composition without changing the estimated expression contrast (Methods 5.2.4; Supplementary Methods S3.5). In the aging-brain series, local composition differences were particularly evident around the lateral ventricles (Fig. 2H). For the comparison between the 21.4-month and 4.3-month sections, composition inconsistency was concentrated around central and ventricular regions.

Applying the adjustment primarily affected the local *Gamt* statistics in these regions (Fig. 5C).

Beyond differential expression analysis, we also used spAlignDE to examine age-associated changes in spatial cell-type organization. We quantified T-cell proximity by the median nearest-neighbor distance and spatial overlap by the mismatchrisk-weighted correlation between smoothed cell-density fields (Methods 5.4.1; Supplementary Methods S3.11). For the proximity analysis, a positive *β* indicates decreasing distance with age; for the overlap analysis, a positive *β* indicates increasing spatial overlap. T-cell proximity to oligodendrocytes, microglia and ependymal cells increased with age (*β* = 0.025, 0.010 and 0.028; *q* = 0.001, 0.048 and 0.001, respectively). Spatial overlap with the same cell classes also increased (*β* = 0.011, 0.010 and 0.006; *q <* 0.001, *q <* 0.001 and *q* = 0.021, respectively; Fig. 5D). These associations reflect increasing spatial colocalization, and the observed patterns were consistent with the source study [40]. To localize this redistribution, we mapped T-cell enrichment in the shared coordinate system (Methods 5.4.1). A comparison of young (≤ 13.6 months; *n* = 7) and old (≥24.3 months; *n* = 7) sections showed greater T-cell enrichment within and around an oligodendrocyte-associated region in older brains (Fig. 5E, top). T-cell enrichment increased with age in the callosal-proximal region (*P* = 2.0 ×10^−4^) but not in the callosal-distal region (*P* = 0.6353; Fig. 5E, bottom). This regional difference indicates that T-cell redistribution was concentrated near oligodendrocyte-associated neighborhoods rather than occurring uniformly across the tissue. More broadly, this analysis shows that spAlignDE extends post-alignment inference from local gene-expression changes to spatially resolved changes in cell-type organization across samples.

In the injured kidney application, we treated the normal NL3 Visium section as the reference and the injured IL3 section as the query, following the comparison in STcompare [28] (Methods 5.4.2; Supplementary Methods S3.12). Gene-level ACAT tests showed spatial expression differences for *Cbr1, Cd44* and *Myo5a* (*P* = 1.7 ×10^−14^, *P* = 4.1 ×10^−6^ and *P* = 7.4× 10^−14^, respectively). The local maps further showed where and in which direction these differences occurred: *Cbr1* and *Cd44* increased in spatially restricted regions of the injured section, whereas *Myo5a* decreased locally (Fig. 5F). Thus, local DE resolved the direction and spatial extent of expression differences within anatomical compartments that could be obscured by compartment-wide aggregation. Because this analysis compared one normal section with one injured section, the results characterize the observed NL3–IL3 section pair rather than a population-level injury effect.

We next asked whether the localized expression differences were associated with compartment-specific functional patterns. We first applied Benjamini–Hochberg adjustment to the gene-level ACAT *P* values across the predeclared family of 16,446 genes and retained genes with 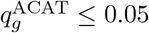 For each retained gene, we tested whether its significant local-grid locations were enriched within the annotated compartments and assigned each spatially concentrated gene to one dominant compartment– direction class. We then performed Gene Ontology (GO) Biological Process over-representation analysis for the genes assigned to each class (Methods 5.4.2; Supplementary Methods S3.12). The five highest-ranked cortex and medulla terms are shown in Fig. 5G, and the 15 highest-ranked terms for cortex, interface and medulla are shown in Supplementary Figs. S35–S36. Cortex-decrease terms included amino-acid catabolism and carbon dioxide transport, including dicarboxylic-acid metabolism. Cortex-increase terms included protein insertion into membranes and glutathione, heme, digestive-system and aldehyde metabolic processes. Interface-decrease terms included mitochondrial electron transport and ATP synthesis; interface-increase terms included extracellular-matrix organization and interleukin-4 responses. Medulla-decrease terms were dominated by amino-acid catabolism and fatty-acid beta-oxidation. Medulla-increase terms included postsynaptic translation, large-subunit biogenesis and ribosome assembly, p53-mediated signaling regulation and positive regulation of telomere maintenance via telomerase. For comparison, the corresponding ORA rankings from compartment-level DE are shown in Supplementary Figs. S37–S38. Among the significant GO terms identified by local DE, 92% occurred in one compartment–direction class, 7% in two classes and 1% in at least three classes. The corresponding proportions for compartment-level DE ORA were 53%, 26% and 21%, respectively (Fig. 5H). For this section pair, local DE therefore produced more compartment- and direction-specific functional enrichment patterns than compartment-level DE.

## 3 Discussion

We developed spAlignDE as an integrated framework for generalizable spatial alignment and mismatch-aware local differential expression analysis. By converting transcriptomic, epigenomic, histological and atlas-derived tissue organization into comparable continuous fields and identifying geometrically similar structures, spAlignDE enables S-LDDMM registration without requiring a shared molecular feature space. For post-alignment inference, spAlignDE incorporates residual mismatch into the variance of local expression contrasts rather than treating aligned coordinates as exact. Across the evaluated datasets, spAlignDE preserved global tissue geometry and local anatomy, scaled to a large multi-sample cohort and aligned spatial transcriptomics with histology, anatomical atlases and spatial ATAC-seq. Simulations showed that mismatch-aware variance inflation reduced false discoveries. The aging-brain analysis localized gene-expression and cell-neighborhood changes that section-wide summaries could obscure, and the injured-kidney analysis identified local expression patterns within anatomical compartments that compartment-level aggregation could obscure.

With one section per condition, spAlignDE’s local tests characterize spatial variation within the observed section pair rather than a population-level condition effect. At each location, spAlignDE tests whether the expression contrast differs from the section-wide baseline for that gene after optional adjustment for confounding variation, such as effects of local library size and detection rate. When the comparison-wide intercept *γ* _*g*_ is included, the test does not assess whether the raw between-section difference is zero. Consequently, spatially uniform shifts, whether technical or biological, may be absorbed by the section-wide baseline rather than identified as local deviations. When multiple sections are aligned, the current workflow fits pairwise or sequential contrasts separately according to the scientific comparison. Cells or spots improve estimation of the local expression surfaces but are not biological replicates. Results from a single section pair, including the injured-kidney analysis, should therefore be interpreted as sample-specific spatial contrasts. Extending spAlignDE to multiple biological replicates per condition will require a hierarchical model that captures between-section variability, potentially through section-level random effects. We anticipate that replicated spatial datasets will become more common and provide the data needed to develop and evaluate this extension for population-level inference.

Several limitations arise from the structure-guided alignment. Alignment accuracy depends on accurate identification and correspondence of spatial domains across samples or modalities. Clustering errors or differences in segmentation granularity can produce incompatible region partitions. For example, a single domain in one dataset may be divided into several subregions in another, and incorrect pairing of these regions can distort the estimated transformation. spAlignDE can automatically merge oversegmented regions and allows users to specify grouped one-to-many or many-to-many correspondences through the interactive interface. However, spAlignDE does not yet automatically infer general many-to-many mappings. Automatic resolution of one-to-many and many-to-many region correspondences would reduce sensitivity to clustering resolution and the need for manual specification.

The query-to-reference design introduces an additional dependence on the selected reference. The reference defines the common coordinate system and determines which regions have sufficient corresponding tissue support for downstream testing. When multiple suitable references are available, the sensitivity of both alignment and local inference to reference choice should be evaluated. S-LDDMM also assumes that corresponding tissues can be related by a smooth, one-to-one, topology-preserving transformation. This assumption may not hold when tissue is torn or missing, when a large lesion lacks a valid counterpart, or when sections collected at different depths contain different structures. Downweighting regions without reliable correspondence can limit their influence on registration, but it cannot establish valid location-level comparisons where no corresponding tissue exists. Accordingly, spAlignDE is intended primarily for two-dimensional sections with sufficiently shared anatomy and locally comparable neighborhoods. Three-dimensional reconstruction of serial sections will require substantial model extensions to accommodate structures that appear or disappear along the depth axis.

The mismatch-risk and subsampling-based measures also require careful interpretation. The mismatch-risk score measures local comparability and is not a direct estimate of alignment error. It assumes that accurately aligned neighborhoods have similar stable-gene expression and local observation density. Genuine biological differences may violate this assumption, increase the risk score and reduce sensitivity through conservative variance inflation. Mismatch-aware testing also cannot recover power lost through severe misalignment. The subsampling-based measure quantifies empirical transformation stability rather than calibrated alignment uncertainty. Future work could derive uncertainty directly from the alignment model and propagate it through a joint framework for alignment and statistical inference.

## 5 Methods

### 5.1 Structure-guided spatial alignment

spAlignDE aligns a query spatial dataset to a fixed reference coordinate system through five stages: spatial-structure construction, global pre-alignment, structure-pair identification, conversion to continuous multichannel fields and shooting-based large deformation diffeomorphic metric mapping (S-LDDMM) [32–34]. The query is the dataset whose coordinates are transformed, whereas the reference remains fixed. The same workflow accommodates cross-sample spatial transcriptomics and cross-modal alignment because modality-specific tissue organization is represented through a common spatial-structure abstraction.

#### 5.1.1 Construction of modality-specific spatial structures

Alignment begins by representing each spatial dataset as a two-dimensional image-like representation in which colors encode spatial structures. Depending on the modality, these structures may correspond to spatial domains, anatomical annotations or image-derived segmentation. They are typically identified using spatial clustering or segmentation procedures that produce spatially coherent domains, although spAlignDE can also accept user-defined structures. The resulting labels are subsequently converted into masks or continuous structural channels for searching cross-modality structure correspondence and S-LDDMM registration

##### Spatial transcriptomics

For cross-sample ST alignment, BANKSY combines gene-expression profiles with local spatial-neighborhood information within each section, after which the sample-specific representations are integrated with Harmony [30, 31]. Leiden clustering of the integrated representation assigns shared spatial-structure identities across sections; therefore, identically labeled structures have natural correspondence across samples. For cross-modality alignment, BANKSY clustering is performed independently within each ST sample, as correspondence is established later based on geometric compatibility. Detailed preprocessing and clustering settings are provided in Supplementary Methods S2.1.1.

##### Histological images

Histology-derived spatial structures are constructed from feature embeddings extracted using pretrained computational pathology models, including HIPT [52] and UNI [53]. Candidate representations based on HIPT embeddings alone or combined HIPT–UNI embeddings are clustered and spatially refined to obtain coherent tissue regions for alignment. Stain-specific preprocessing, feature-model configurations and final structure-map selection are described in Supplementary Methods S2.1.2.

##### Anatomical reference atlases

Atlas structures are obtained from predefined annotation labels and an anatomical hierarchy rather than inferred by clustering. In the Allen CCFv3 analysis, descendant labels are combined along hierarchical structure paths to generate candidate masks at multiple anatomical depths, with cortical layers additionally represented as layer-specific structures [9, 10]. Details are provided in Supplementary Methods S2.1.4.

##### Spatial ATAC-seq

Spatial ATAC-seq structures are identified by applying BANKSY to an ATAC-derived gene-activity score matrix and its spatial coordinates. The ST reference is clustered independently from its expression matrix, so ATAC and ST structures are connected only during geometric structure pairing. Gene-activity processing and clustering settings are provided in Supplementary Methods S2.1.3.

### 5.1.2 Global pre-alignment

Before deformable alignment, spAlignDE corrects major differences in tissue position, orientation and scale while keeping the reference fixed. Global pre-alignment is important for both structure-correspondence discovery and deformable registration. In cross-modal alignment, it places independently constructed structures into approximate spatial correspondence, making the overlap-, boundary-, size- and thickness-based components of the composite pairing score more meaningful (Section 5.1.3); inadequate initialization can therefore alter candidate-pair ranking and acceptance. It also provides a stable starting point for S-LDDMM, allowing the subsequent affine–diffeomorphic stage to refine residual global and local deformation rather than compensating for large initial differences in tissue position, orientation or scale (Section 5.1.5).

In the analyses reported here, shared-structure-centroid pre-alignment was used for the primary cross-sample ST analyses. Whole-tissue-mask-overlap pre-alignment was used for ST-to-atlas alignment and for the Xenium-to-H&E analysis. Interactive manual pre-alignment was used for MERFISH-to-Nissl and spatial ATAC-to-ST alignment and for the Xenium breast-cancer ST-to-ST partial-alignment example, for which automatic initialization was less reliable.

Under the centroid-based strategy, a weighted similarity transformation is estimated from shared-structure centroids. Let 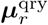 and 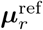 denote the query and reference centroids of shared structure *r*. The global transformation solves

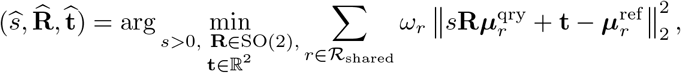

where *s* is an isotropic scaling factor, **R** ∈ SO(2) is a two-dimensional rotation matrix, thereby excluding reflection, and **t** ∈ ℝ^2^ is a translation vector. The set ℛ_shared_ contains the structure identities represented in both datasets. For structure *r*, let 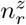 denote the number of observations assigned to that structure in dataset *z* ∈ {qry, ref}. The non-negative weights are normalized to sum to one and satisfy *ω* _*r*_ ∝ min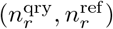.

When structure identities are unavailable but comparable whole-tissue masks can be constructed, spAlignDE uses the whole-tissue centers to determine translation and selects rotation and scale by maximizing whole-tissue-mask overlap. This strategy works well for tissues with fixed shapes, for example, coronal sections of the brain. Let **c**_qry_ and **c**_ref_ denote the fitted query and reference tissue centers, respectively. For candidate scale *s >* 0 and rotation angle *θ*, a query coordinate **x** is transformed as

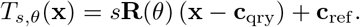

Here, **R**(*θ*) is the two-dimensional rotation matrix associated with *θ*. Centering the query coordinates at **c**_qry_ and adding **c**_ref_ determines the translation for each candidate *s* and *θ*, so translation is not optimized as an independent search parameter.

The transformed query coordinates are rasterized to obtain the binary mask 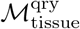 (*s, θ*) on the same grid as the fixed reference mask 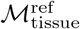 . Their overlap is

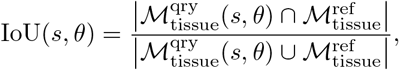

where | · | denotes the number of foreground pixels. The candidate rotation and scale producing the largest IoU are retained.

An interactive similarity pre-alignment is available when neither automatic strategy is sufficiently stable. Exact search ranges, tissue-center estimation, mask operations and interface behavior are described in Supplementary Methods S2.3.1.

### 5.1.3 Structure pairing

After global pre-alignment, spAlignDE determines which query and reference structures provide corresponding anatomical support. Cross-sample ST does not require structure pairing if structures are constructed by multisample joint clustering; cross-modal alignment compares independently constructed structures after representing them on a common raster domain. Candidate pairs without sufficient support are excluded rather than forced into alignment.

#### Structure-mask construction

Candidate cross-modal structures are represented as binary masks on a shared raster domain. Structures that are point-based (from cells, spots, or bins) are filtered only for mask construction, rasterized, adaptively smoothed and refined to connect small gaps, fill holes and remove isolated components; all original observations still receive the final transformation. Histology and atlas masks are obtained from their labeled image regions. Detailed procedures are provided in Supplementary Methods S2.3.2.

#### Cross-sample correspondence

For cross-sample ST alignment, structures assigned the same label through joint clustering are treated as corresponding across samples. Only structures present in both samples are used for correspondence-based prealignment and transformation estimation, but the cells, spots, or pixels in the remaining area still receive the transformation estimated from the shared structures.

#### Cross-modal correspondence

For query structure *i* and reference structure *j*, spAlignDE uses the composite score

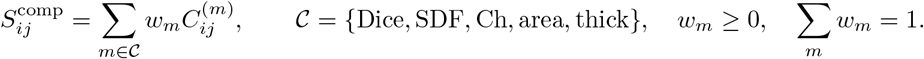

The five components summarize complementary aspects of geometric correspondence. *Dice similarity* measures direct foreground-mask overlap. *Signed-distance-field (SDF)* similarity compares signed distances within a boundary band and therefore captures similar boundary geometry despite modest displacement. *Chamfer similarity* converts the mean bidirectional nearest-boundary distance into a decreasing similarity score. *Area similarity* penalizes differences in total spatial extent through a symmetric relative-size comparison. *Thickness similarity* compares a distance-transform summary of characteristic width and is particularly informative for narrow or layer-like structures.

Pairing candidates are additionally screened using modality-specific quality-control criteria, including minimum overlap or maximum boundary displacement where appropriate. Accepted candidates are ranked and selected greedily, with each primitive query and reference structure used at most once unless structures have first been combined into an explicit correspondence group. Exact component definitions, weights, thresholds and soft penalties are provided in Supplementary Methods S2.4.

### 5.1.4 Continuous structural fields

S-LDDMM operates on image-like continuous multichannel fields rather than directly on discrete observations or categorical labels. For cross-sample ST, observations assigned to each shared structure are rasterized and spatially smoothed to define a local structure-composition channel. For single-cell datasets, an additional channel can represent the smoothed local density of cells assigned to shared structures when appropriate. No spot-density information contributed to transformation estimation for spot-based ST because the limited number of spots did not support a sufficiently continuous density field.

For cross-modal alignment, the discrete cells, spots or pixels assigned to each accepted structure are first rasterized as binary masks on a common spatial domain. Directly matching binary masks provides piecewise-constant fields whose spatial variation is concentrated at structure boundaries and can therefore provide limited guidance when paired structures remain separated after global pre-alignment. Each paired mask is instead converted into a signed-distance-transform (SDT) field, whose magnitude records distance to the nearest boundary and whose sign distinguishes the structure interior from the exterior. The SDT converts boundary displacement into a spatially graded matching signal, providing useful correspondence information even before the paired boundaries overlap and increasing the effective capture range of the deformable registration [54]. Smoothing, distance truncation and normalization reduce sensitivity to irregular mask boundaries and prevent distant interior or background locations from dominating the matching objective.

The query and reference fields are sampled on the same regular raster and arranged in the same channel order. For cross-sample alignment, the raster covers the joint spatial extent of the pre-aligned query and reference, and its dimensions are determined by the selected grid spacing rather than by the number of cells or spots. For cross-modal alignment, the raster is inherited from the fixed reference image, atlas section or common pre-alignment canvas, with optional resolution adjustment while preserving the same coordinate extent. Let *Ω* → ℝ^2^ denote this common raster domain and let *F* ^qry^, *F* ^ref^ : *Ω* ⊂ℝ^*C*^ denote the query and reference fields, where *C* is the total number of input channels. Detailed rasterization, normalization and grid-construction procedures are provided in Supplementary Methods S2.6.

### 5.1.5 Structure-guided shooting LDDMM

Let **x** ∈ *Ω* ⊂ ℝ^2^ denote a query coordinate after global pre-alignment. spAlignDE maps **x** into the fixed reference coordinate system by composing a smooth nonlinear deformation with a residual affine transformation:

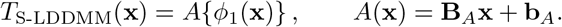

Here, *ϕ* _1_ corrects local nonlinear deformation, whereas *A* corrects remaining global linear differences. The matrix **B**_*A*_ ∈ ℝ^2*×*2^ and translation vector **b**_*A*_ ∈ ℝ^2^ define the affine transformation.

The nonlinear transformation *ϕ* _1_ is the endpoint of the diffeomorphic flow

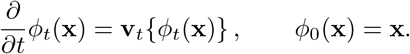

Here, *ϕ* _0_ is the identity transformation, *t* ∈ [0, 1] is an artificial deformation-time parameter, and **v**_*t*_ : ℝ^2^ → ℝ^2^ is the smooth velocity field that determines the instantaneous displacement of each coordinate. The transformation *ϕ* _1_ therefore represents the final nonlinear deformation at *t* = 1. Compared with a non-shooting, time-discretized LDDMM formulation that directly parameterizes the full velocity sequence, S-LDDMM generates the complete deformation trajectory from a single initial momentum field, thereby reducing the number of independently estimated deformation variables while retaining the smooth, invertible and topology-preserving properties of LDDMM [32, 33]. To accommodate unreliable local correspondence, spAlignDE adapts the probabilistic missing-data formulation of Tward et al. [34] to multichannel structural fields, using a three-component expectation–maximization mixture comprising one matched component and two unmatched appearance components. At each reference-grid location, *W*_*M*_ (**x**) ∈ [0, 1] denotes the posterior probability that the local multichannel fields belong to the matched component. With 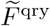 denoting the inverse-warped query field and *G* an affine mapping of query-channel values to the reference-channel scale, the principal matching term is

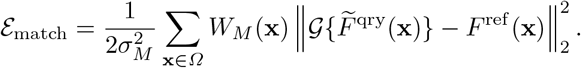

Here, *σ* _*M*_ *>* 0 controls the variability accommodated by the matched component. The complete objective ℰ = ℰ _match_ + ℰ_reg_ balances multichannel agreement with the LDDMM kinetic-energy penalty. Locations representing missing tissue, weak overlap or modality-specific information receive smaller *W*_*M*_(**x**) and therefore do not force unsupported deformation. The final query-to-reference mapping is

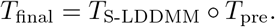

Here, *T*_pre_ denotes the selected global pre-alignment, and the composition applies *T*_pre_ before the S-LDDMM refinement. The complete shooting equations, matched/unmatched mixture, channel mapping, deformation regularization and optimization procedure are provided in Supplementary Methods S2.7.

### 5.1.6 Coarse-to-fine ST-guided atlas alignment and interactive pairing

For atlas alignment, spAlignDE uses the Allen CCFv3 hierarchy to construct a multiscale set of candidate anatomical regions, while representing the ST section by nested partitions ranging from broad transcriptomic domains to the finest BANKSY structures. Alignment proceeds from coarse to fine on the ST side: at each resolution of spatial clustering, every ST structure is compared with the same set of eligible atlas candidates, so the matched atlas region is not restricted to a particular hierarchy depth. Accepted ST–atlas pairs are converted into matched SDT channels and used to estimate an S-LDDMM transformation, which is applied to all ST coordinates and masks before pairing is reconsidered at the next finer resolution. After the finest ST partition is reached, pairing and alignment are repeated until a complete refinement cycle identifies no additional accepted pairs or the maximum number of cycles is reached.

When automatic geometric pairing remains ambiguous, spAlignDE Structure Pair allows users to inspect structures side by side, combine primitive structures into correspondence groups and save the resulting assumptions reproducibly. User-defined groups bypass automatic pair selection but enter the same SDT construction and S-LDDMM workflow. Interface details are provided in Supplementary Methods S2.5; stage-specific atlas settings are provided in Supplementary Methods S2.8.

## 5.2 Mismatch-aware post-alignment inference

The alignment component maps spatial transcriptomic samples into a shared coordinate system. The post-alignment model then defines contrasts according to the scientific comparison rather than the direction used to estimate the alignment. Throughout the post-alignment inference model, sample A denotes the case, treatment or time point of interest, and sample B denotes the control or baseline condition. These inferential labels are independent of which sample was treated as the moving query or fixed reference during alignment. Cells or spots provide the local observations, shared-grid locations are the local testing units, tissue sections are the biological sample units, and case–control or sequential condition comparisons define the fitted contrasts. To keep the notation readable, the location-level model below is written for a generic single contrast and suppresses the contrast index *c*; *c* is restored only where multiple contrasts are calibrated or analyzed jointly.

### 5.2.1 Definition and interpretation of the local DE test

For gene *g* at grid location *i*, let *µ*_A,*ig*_ and *µ*_B,*ig*_ denote the underlying local means in the case and control samples, respectively. The directional contrast is

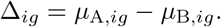

The primary local estimand is the departure of this contrast from a comparison-wide baseline and, when enabled, a grid-varying confounding-covariate term,

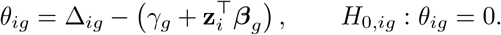

Here, *γ* _*g*_ represents the component of the between-section contrast that is approximately constant over the tested grid, while 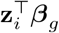 optionally captures confounding variation such as effects of local library size and detection rate. Including *γ* _*g*_ defines local DE as a spatial departure from the comparison-wide shift; the intercept can be omitted when an approximately uniform difference is itself part of the inferential target.

Cells or spots supply the local measurements used to estimate a section-level spatial contrast but do not constitute independent biological replicates. In an ordered series such as the aging-brain analysis, each non-baseline age section is treated as sample A and the common baseline-age section as sample B before evidence is aggregated across sections. The current formulation does not estimate within-age replication because only one section was available at each age.

### 5.2.2 Shared grid, kernel estimators and base local-contrast variance

Because aligned cells or spots are not themselves one-to-one matched observations, local contrasts are evaluated on a shared grid of common spatial locations. Let ***ξ***^*i*^, *i* = 1, …, *N*_grid_, denote candidate grid locations in the aligned coordinate system. The retained testing set consists of locations passing the common-support mask and sample-specific minimum-support filters; the exact support-mask construction and analysis-specific grid settings are reported in Supplementary Methods S3.1 and S3.2. Let 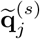 and 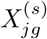 denote the aligned coordinate and expression of observation *j* in sample *s*. Around retained grid location *i*, spAlignDE defines the Gaussianweighted neighborhood

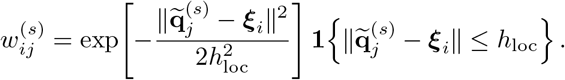

Here, *h*_loc_ is the local spatial bandwidth; it sets the Gaussian distance scale and excludes aligned observations farther than *h*_loc_ from grid location *i*. With normalized weights 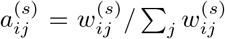, the local mean estimator is 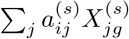. Under a working local approximation in which contributing observations are independent and have common variance *σ*^2^, its variance is

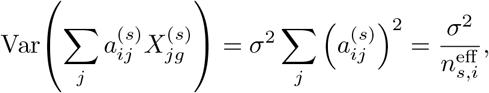

which gives the Kish effective local sample size [55],

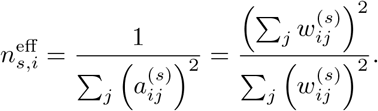

Thus, concentrated kernel weights provide less effective information than equal weights over the same number of observations. This calculation is a working local-sampling approximation: residual spatial correlation among nearby cells or spots is not modeled explicitly, and overlapping neighborhoods induce dependence among neighboring grid tests.

The kernel-weighted local mean, variance and contrast are

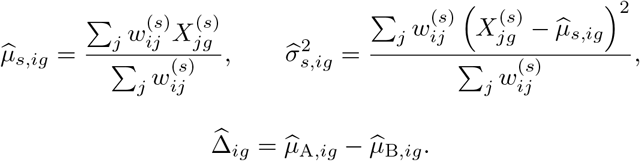

The effective-sample-size-adjusted Welch-type variance estimate of the local contrast is

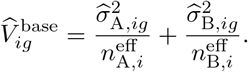

Grid locations with insufficient effective support in either sample are excluded. The effective spatial resolution is therefore determined by observation density relative to *h*_loc_, not by shared-grid density alone. In ordered multi-sample analyses, each case–baseline or sequential contrast is fitted independently; local means are not pooled across contrasts.

### 5.2.3 Putatively stable genes and local mismatch-risk estimation

The mismatch-risk map is estimated separately for each contrast from a panel of *putatively stable* genes. These genes are not assumed to be known negative controls; rather, they are selected as internal proxies for local comparability because they show informative spatial variation, positive cross-sample spatial correspondence and a small comparison-wide difference. For the generic single contrast considered here, define

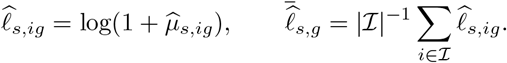

The putatively stable panel is

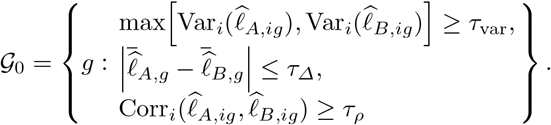

Exact thresholds, ranking rules and the maximum panel size are given in Supplementary Methods S3.3.

The risk map is constructed once for each pairwise or sequential contrast and is then shared across all target-gene fits for that contrast; it is not recomputed after removing the tested gene. Thus, if a target gene also belongs to *G*_0_, it remains in the panel used to construct the common risk map. Its three distributional channels form only one part of a pooled profile containing the corresponding channels from the full stable-gene panel together with the density channel, so its inclusion does not make the risk score gene specific. The use of empirically selected stably expressed genes as negative-control features for estimating shared unwanted variation follows related precedent in single-cell integration [35].

For a putatively stable gene *g*∈ *G*_0_, let 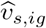 denote its kernel-weighted local variance. A moment estimator of the local negative-binomial size is

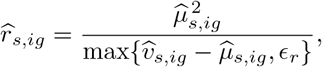

with corresponding fitted zero probability

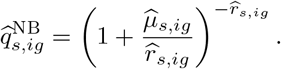

If 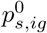 is the kernel-weighted observed zero fraction, the excess-zero probability is

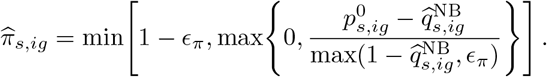

The three-channel local profile

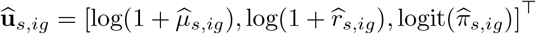

summarizes local expression magnitude, overdispersion and excess sparsity. Each channel is standardized across valid grid locations within its own sample; let Ƶ _*s*_ denote this sample-specific standardization. Local sampling density is summarized by

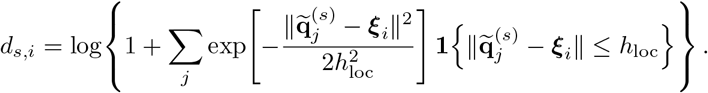

Let *d*_gene_ = 3|*G*_0_ | denote the number of standardized stable-gene channels. For a target density-energy share *s* ∈ (0, 1), the density multiplier is

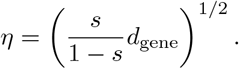

Thus, *s* controls the target contribution of the standardized density channel to the combined profile energy; it is not a direct weight on the final risk map or variance factor. The augmented profile is

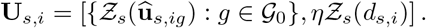

The raw mismatch score is the symmetric cosine dissimilarity

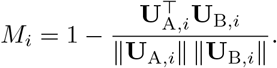

It is smoothed over the grid,

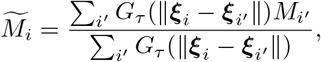

and robustly normalized within the contrast,

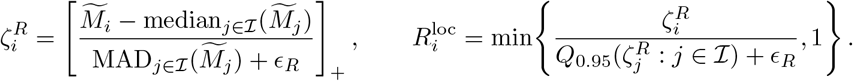

Thus, 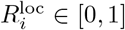. This score ranks relative local non-comparability within a contrast; it is neither a calibrated probability of misalignment nor a gene-specific biological effect.

### 5.2.4 Empirical mismatch calibration and cell-type-composition adjustment

For target gene *g*, the alignment-mismatch variance factor for the generic single contrast is

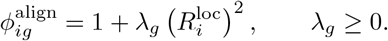

The location-specific risk map 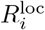 identifies where residual alignment mismatch may reduce precision within the current contrast, and the gene-specific coefficient *λ* _*g*_ determines how strongly this local risk inflates the variance for gene *g*. Calibration is based on the expectation that, if residual mismatch adds uncertainty, higher-risk strata will show greater excess dispersion in the initial local statistics relative to their corresponding Student-*t* null distributions.

The quadratic form is motivated by a local Taylor approximation of a smooth spatial expression field. A small alignment displacement ***δ*** _*i*_ gives

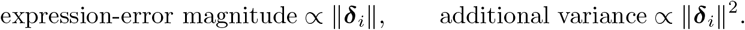

We therefore model alignment-induced variance inflation as quadratic in the local mismatch risk. This form also limits inflation at low-risk locations while concentrating the adjustment in regions with stronger mismatch.

To describe calibration in a form that covers both single- and multi-contrast analyses, we restore the contrast index *c*. The local model is first fitted with mismatch inflation disabled. Each contrast is calibrated separately using its initial local statistics 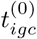 and risk map 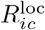. Locations are grouped into risk strata, the statistics are median-centered within each stratum and their robust scale is compared with the corresponding Student-*t* null scale. For bin *b*, with median risk *r*_*bc*_, size *n*_*bc*_ and residual degrees of freedom *v*_*gc*_,

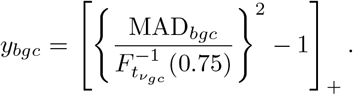

Positive excess dispersion is constrained to be nondecreasing with risk by weighted isotonic regression,

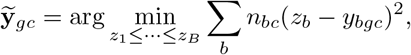

and a non-negative quadratic relation through the origin is fitted,

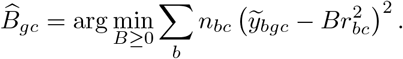

A bounded reference-bin adjustment converts 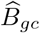 into a provisional contrast-specific coefficient 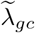. In a multi-contrast analysis, valid provisional coefficients are combined by an equal-weight Huber robust center to obtain one shared genespecific coefficient 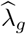. This avoids dependence on an arbitrarily selected anchor contrast and limits the influence of a contrast with unusually strong biological change or unstable calibration. If only one contrast is available, its valid provisional coefficient is used directly, leaving the single-contrast procedure unchanged. The resulting 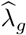 is held fixed when the final variances and local statistics are constructed, while every contrast retains its own risk map. Exact validity, aggregation and fallback rules are provided in Supplementary Methods S3.4.

This calibration assumes that risk-associated increases in the dispersion of the median-centered initial local statistics reflect residual mismatch-induced variability rather than systematic true local signal. The risk map, 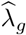 and the confoundingcovariate basis are treated as plug-in quantities in the final test; their estimation uncertainty is not propagated separately.

#### Cell-type-composition adjustment

Returning to the generic single-contrast notation, when cell-type annotations or spot-level deconvolution estimates are available, let 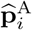 and 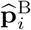 denote the kernel-smoothed local cell-type-composition vectors in the case and control samples. Their midpoint, normalized Jensen–Shannon distance and resulting variance factor are

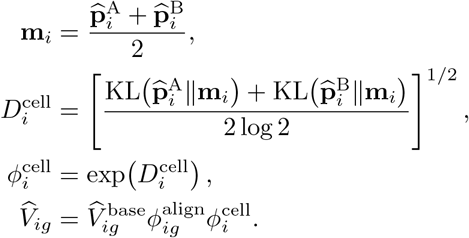

The Kullback–Leibler divergences use natural logarithms. The Jensen–Shannon divergence ranges from zero to log 2, so division by log 2 maps it to [0, 1]. Its square root gives the bounded composition distance 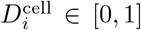. The exponential link then maps this distance to 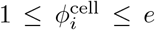. Identical local compositions give a variance factor of one, and increasing compositional inconsistency smoothly increases the local variance. The factor governs inferential precision; cell-type-specific expression effects require a separate model. Local composition estimation and fallback settings are described in Supplementary Methods S3.5.

### 5.2.5 Local regression with optional confounding-covariate adjustment

For sample *s* and grid location *i*, let *L*_*is*_ and *D*_*is*_ denote local library size and detection rate, and define

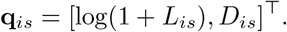

Sample B, or the designated set of control samples when several controls are supplied, defines the centering and scaling of these quality-control profiles. Singular value decomposition of the standardized control profiles supplies the retained directions, and the pooled sample-A batch profile is projected onto those directions to give **z**_*i*_. The resulting basis captures confounding variation such as effects of local library size and detection rate. Its construction is independent of which sample served as the query or reference during alignment and is detailed in Supplementary Methods S3.6.

For one contrast, the working observation model is

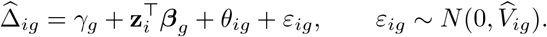

Let

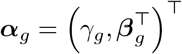

and let 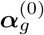 denote the gene-specific anchor estimate described in Supplementary Methods. The baseline coefficients are estimated by

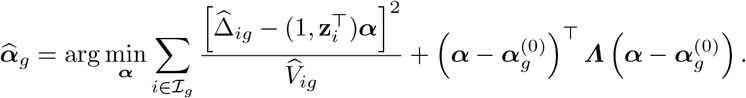

The confounding-covariate coefficients receive the default ridge penalty. When an intercept anchor is available, the intercept is also shrunk toward its anchor; otherwise its corresponding penalty entry is zero and the intercept is unpenalized. The adjusted residual and local statistic are

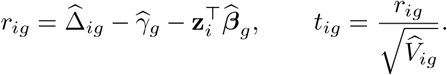

Let *n*_*g*_ be the number of valid grid locations for gene *g*, and let *d*_*g*_ be the number of fitted columns in the baseline design. The implementation uses

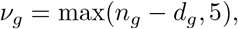

where *d*_*g*_ = 0 when neither the intercept nor a confounding-covariate basis is included and *d*_*g*_ = 1 for an intercept-only baseline. Two-sided plug-in *P* values are

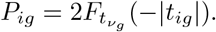

The denominator 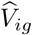 contains the estimated base local-contrast variance and the specified variance-inflation factors, but it does not include a separate weighted-regression leverage correction or an additional variance term for estimating 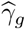 and 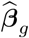 . Accordingly, *t*_*ig*_ is interpreted as a working plug-in statistic.

For each gene, local *q*-values are computed across valid grid locations,

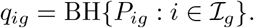

At target level *α*, connected components of {*i* : *q*_*ig*_ ≤*α* } define local DE regions. Each region is summarized by the median adjusted residual, its direction and its minimum local *q*-value. Connected regions inherit evidence from the locally adjusted grid calls and do not receive a separate region-level FDR guarantee. Numerical safeguards and null-reference details are described in Supplementary Methods S3.8 and S3.7.

### 5.2.6 Gene-level, multi-contrast and spatial-trajectory summaries

For a single contrast, spatially dependent local *P* values are combined by the Cauchy combination test (ACAT) [56]. For non-negative weights *a*_*i*_ normalized over valid locations,

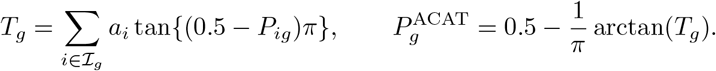

For an ordered sample series, such as aging, the gene-level test targets a spatially distributed trend rather than a difference in any single contrast. Let 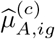 denote the kernel-smoothed local expression estimate for sample A in contrast *c*. The unsmoothed adjusted expression used in the age-trend analysis is

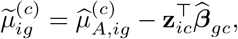

where 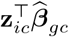 is the fitted grid-varying confounding component derived from local library size and detection rate. The comparison-wide intercept 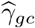 is not removed, so broad expression differences among the ordered sections remain available to the trend analysis. Here, “unsmoothed” means that no trajectory-level temporal smoothing has yet been applied; 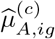is still a spatially kernel-smoothed local expression estimate.

In the aging analysis, *c* indexes the 19 non-reference sections. The 4.3-month reference is used to construct each ageversus-reference local fit but is not included as an additional observation in the age regression. At every valid grid location *i*, the adjusted expression estimates are fitted across the ordered sample variable *x*_*c*_,

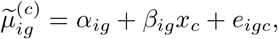

using the mismatch-aware inverse-variance weights from the corresponding local fits. The two-sided local trend *P* values 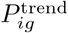 are then combined across valid spatial locations by equally weighted ACAT,

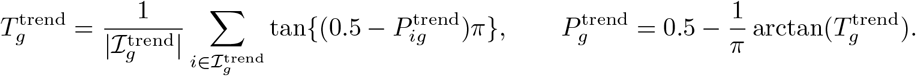

The global null is *β* _*ig*_ = 0 at every tested location. This gene-level trend test uses neither trajectory smoothing nor trajectory clusters and is therefore independent of the selected *K*. Both the single-contrast and ordered-series ACAT values are raw gene-level *P* values unless a separate across-gene adjustment is stated.

For visualization of ordered samples, adjusted local-expression trajectories are clustered into spatial domains. When *K* is not supplied, spAlignDE first evaluates whether cluster-specific time trends improve held-out prediction relative to a shared time trend. If no reliable improvement is detected, it returns the smallest candidate *K*; otherwise, it retains candidates within one standard error of the best held-out dynamic gain. Spatial resolution is then examined from the largest retained *K* toward smaller values. The procedure stops when the next coarser candidate would increase the fraction of grid locations in components smaller than one *R*_map_ footprint and retains the current, finer-side local minimum. If this fragmentation measure decreases throughout the scan and therefore supplies no elbow, the procedure makes one conservative coarsening step from the finest retained candidate rather than continuing to the coarsest candidate. This selection uses neither the gene-level trend test nor the subsequent cluster-level Wald *P* values.

For cluster C_*gκ*_ and contrast *c*, spAlignDE reports a reliability-weighted section-level mean 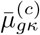 and weighted within-cluster spatial standard deviation 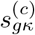. The plotted interval 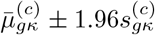 describes spatial dispersion among grid locations and is not a confidence interval for the section-level mean. A spatial-variability-weighted linear trend is fitted across section-level summaries using 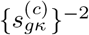 as a reliability weight, yielding a raw two-sided Wald *P* value for each selected cluster.

Because the trajectory partition and *K* are estimated from the same observed trajectories that are subsequently summarized, the cluster-level Wald values are conditional downstream localization summaries rather than tests independent of the clustering step. Selection-aware null calibration would require rerunning trajectory smoothing, clustering, *K* selection and trend fitting under the null and is not part of the current implementation. Complete clustering and trend-summary details are provided in Supplementary Methods S3.9.

The multiplicity scopes are distinct. For analyses with multiple contrasts, restoring the contrast index gives local *q*_*igc*_-values computed across grid locations within each gene and contrast; for a single contrast these reduce to *q*_*ig*_ -values. Cluster-level trajectory trends are reported as raw Wald *P* values without an additional within-gene BH adjustment. Gene-level ACAT values are also raw *P* values unless an across-gene adjustment is explicitly applied, and connected regions receive no additional region-level adjustment. In the injured-kidney analysis, valid local tests were equally weighted within the single IL3-versus-NL3 contrast. The resulting 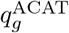-values were computed across the complete predeclared family of 16,446 genes before construction of the local-ORA gene sets.

## 5.3 Benchmark and simulation design

### 5.3.1 Alignment evaluation and robustness

#### Alignment benchmarking

Alignment performance was evaluated using information not supplied to transformation estimation whenever possible. Following the cross-sample spatial-alignment evaluation framework of Yan et al. [25], shared-gene expression was aggregated into corresponding spatial bins and evaluated using four metrics: Pearson correlation coefficient, cosine similarity, structural similarity index measure and mutual information. Compatible annotations were assessed by nearestneighbor label transfer, weighted Dice coefficient and weighted intersection over union. For cross-modal tasks, evaluation used task-specific evidence: molecular measurements paired with the reference histology but withheld from alignment; atlas-label coverage, label agreement between transferred atlas labels and RCTD-inferred class labels, and regional marker localization for ST-to-atlas alignment; and annotation agreement, gene-activity/expression concordance and native-neighborhood preservation for spatial ATAC-to-ST alignment. Wall-clock runtime and peak allocated GPU memory were recorded separately. Complete metric definitions are provided in Supplementary Methods S2.10.

#### Subsampling-based transformation stability

Ten replicate inputs were generated by independently retaining 80% of the cells in both sections and rerunning the complete cross-sample workflow. Each replicate-specific transformation was applied to the same fixed set of pre-aligned query points, and the mean Euclidean displacement from the replicate-mean aligned position was calculated. This quantity measures empirical sensitivity to input subsampling rather than a calibrated confidence interval for the unknown deformation. Full formulas are provided in Supplementary Methods S2.11.

#### Sensitivity to spatial-structure choices

Cross-sample sensitivity was assessed by varying the Leiden resolution used for joint structure identification, whereas cross-modal sensitivity was assessed by reducing the number of retained ST–atlas structure pairs while holding unrelated settings fixed. Complete designs are provided in Supplementary Methods S2.12.

### 5.3.2 Simulation data generation

The simulation benchmark used a real cell-resolved 3.8-month MERFISH mouse-brain section as the tissue and sampling template. Gene-wise negative-binomial spatial generalized additive models preserved the observed library-size, cell-type and spatial-expression structure when generating conditionally independent count profiles for a baseline reference (simulation sample A) and a signal-bearing sample (simulation sample B). Six non-overlapping circular DE regions, comprising three upregulated and three downregulated regions, were implanted in the signal-bearing sample. Three separate distortion regions were placed outside the DE regions and subjected to smoothly blended local affine deformation; global affine and radial-basis-function deformations and coordinate noise were then added to the signal-bearing sample. The simulation A/B labels refer only to data generation; in post-alignment inference, the signal-bearing sample was the condition of interest and the baseline reference was the control. This data-informed construction retained realistic tissue geometry and spatial sampling while preserving known expression and deformation ground truth [50]. Ten matched simulation groups were generated by varying the DE-and distortion-region placements while limiting pairwise overlap among their spatial masks. Further details are provided in Supplementary Methods S3.10.

### 5.3.3 Simulation evaluation

Post-alignment inference was performed at shared-grid locations. To compare the calls with the simulation truth, grid-level q-values, local statistics and significance calls were projected to sample-B cells, where the implanted DE regions and directions were defined. For one simulation group, let *D* be the number of projected cells called significant at the target FDR of 0.05, *F* the number of called cells outside the implanted DE regions and *T* the number of correctly signed called cells within those regions. Let *M*_1_ be the total number of cells in the implanted DE regions. The location-level false discovery proportion and power are

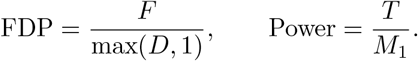

These quantities are calculated across sample-B spatial locations rather than across genes. Each simulation group therefore contributes one location-level FDP and one power value to the summaries in Fig. 4 and the Supplementary Figures.

For power at FDP ≤ 0.05, valid projected cells were ranked by increasing q-value, with ties ordered by decreasing absolute local statistic. FDP and power were evaluated along this ranking, and the largest power among prefixes satisfying the location-level FDP constraint was reported,

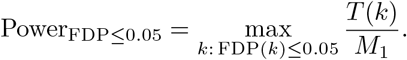

This truth-dependent thresholding is used only to compare sensitivity at a common realized error level in simulations; it is not an output of the real-data analysis.

## 5.4 Application-specific analyses

### 5.4.1 Aging-brain analysis

The processed coronal aging-brain MERFISH series comprised 20 sections from 3.4 to 34.5 months. The 4.3-month section was used as the reference because it retained the most complete overall anatomy, and each of the remaining 19 sections was aligned to it. Local expression testing was performed separately for every age-versus-reference contrast. Gene-level age evidence was obtained by fitting mismatch-aware local linear trends across the ordered sections and combining the resulting local trend *P* values across space by ACAT. Adjusted local-expression trajectories were clustered and summarized as described above.

Cell-type neighborhood analyses were performed on aligned coordinates. For each section, T-cell proximity to oligodendrocytes, microglia and ependymal cells was summarized by the median nearest-neighbor distance, transformed so that a positive age coefficient denotes decreasing distance. Spatial overlap was summarized by the mismatch-risk-weighted correlation between kernel-smoothed T-cell and target-cell density fields. BH adjustment was applied across target cell types separately for proximity and overlap. Local T-cell enrichment maps contrasted normalized T-cell mass with its smoothed expectation under local tissue area. Young and old summaries used the age groups shown in Fig. 5E (young, ≤ 13.6 months, *n* = 7; old, ≥24.3 months, *n* = 7). Detailed formulas are provided in Supplementary Methods S3.11.

Counts were normalized to a target total of 250. The shared grid contained 77,056 valid locations with spacing 27.05; the local neighborhood bandwidth was 121.1 and *R*_map_ = 40.57. The analyses in Fig. 5A–B included the comparison-wide intercept, adjusted for local library size and detection rate, used mismatch-aware variance inflation with density-energy share *s* = 0.25, and left cell-type adjustment off. For each gene, provisional mismatch-inflation coefficients were estimated separately in the 19 age-versus-reference contrasts and robustly combined into one shared gene-specific coefficient. Figure 5C used the 21.4-versus-4.3-month contrast from the same multi-contrast calibration, with cell-type adjustment switched off or on and *α* = 0.10. Putatively stable genes were selected using the default screen in Supplementary Methods S3.3. Gene-level aging summaries combined equally weighted local age-trend *P* values across valid grid locations.

### 5.4.2 Injured-kidney analysis

The injured IL3 Visium section was aligned as the query to the normal NL3 section as the reference. For post-alignment inference, injured IL3 was sample A (case) and normal NL3 was sample B (control), so positive local contrasts indicate higher expression in the injured section. The analysis estimated local expression contrasts on their shared aligned support and summarized gene-level evidence by equally weighted ACAT across valid grid locations. The resulting 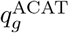-values were computed across the complete predeclared 16,446-gene testing family, and 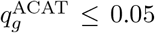 was required for entry into the local-ORA gene sets. For each gene and direction, significant local-grid locations were tested for enrichment in the annotated compartments after accounting for compartment area. A gene satisfying the spatial concentration requirements was assigned to one dominant compartment–direction class, and each class-specific gene set was tested for Gene Ontology Biological Process over-representation against the complete testing family, yielding term *q*-values within that class. For display, terms within each compartment–direction class were ranked by increasing term *q*-value. Ties were resolved successively by increasing raw over-representation *P* value, decreasing overlap gene count and alphabetical GO term name.

For comparison, spots were assigned to the cortex, interface or medulla and injured-versus-normal expression was tested within each compartment. Genes passing compartment-specific BH adjustment were divided by direction and subjected to the same GO over-representation framework. Term recurrence was summarized by the number of compartment–direction classes in which each significant term appeared. Detailed formulas are provided in Supplementary Methods S3.12.

Counts were library-size normalized to a target total of 10,000, and genes detected in at least 10 spots with at least 10 total counts were retained. The shared grid contained 6,187 valid locations with spacing 75.49; the local neighborhood bandwidth was 191.9 and *R*_map_ = 113.2. The formal full-gene and displayed-gene fits omitted the comparison-wide intercept. This choice treated broad injury-associated expression shifts as part of the injured-versus-normal contrast rather than as a nuisance baseline. The fits adjusted for local library size and detection rate, enabled mismatch-aware variance inflation with density-energy share *s* = 0.75, disabled cell-type adjustment, and used *α* = 0.05 with connected-region cleanup disabled. Gene-level Aggregated Cauchy Association Test values were adjusted across the complete 16,446-gene family before local DE-region over-representation analysis. The final Gene Ontology analysis used `clusterProfiler` 4.18.4 and `org.Mm.eg.db` 3.22.0.

## 5.5 Software and computational resources

### 5.5.1 Software and reproducibility

spAlignDE analyses were performed using spAlignDE 0.1.0 under Python 3.10.14. The software environment and workflow-specific random seeds are described in Supplementary Methods S1. Comparator versions, configurations and method-specific input limits are reported in Supplementary Methods S2.9. Runtime and GPU-memory measurements are defined in Supplementary Methods S2.10.1.

### 5.5.2 Computational resources

Computational analyses were performed on a workstation equipped with an AMD Ryzen Threadripper PRO 7985WX 64-Core Processor with 64 physical cores / 128 threads, 503 GB of system memory, and an NVIDIA RTX PRO 6000 Blackwell Max-Q Workstation Edition GPU with 96 GB of GPU memory. GPU-accelerated analyses used NVIDIA driver 570.195.03 with CUDA 12.8 support.

### 5.5.3 Use of generative AI tools

We used OpenAI’s ChatGPT and Codex for language polishing and limited programming assistance during the preparation of this work. For manuscript preparation, these tools were used selectively to improve readability, clarity, and flow while preserving the authors’ intended scientific meaning. Programming assistance was limited to code debugging, refactoring, and implementation suggestions. The study conception, methodological design, scientific analyses, interpretation of results, and conclusions were developed by the authors. All AI-assisted text was reviewed and revised by the authors, and all AI-assisted code was reviewed, tested, and modified as appropriate. The authors take full responsibility for the accuracy, integrity, and final content of the manuscript and associated code.

## 6 Data Availability

All datasets used in this study are publicly available. The 10x Genomics Visium mouse kidney sections NL3 (normal control) and IL3 (ischemia–reperfusion injury), together with their region annotations, were obtained from the STcompare Zenodo record (https://zenodo.org/records/20647680). Replicate 1 of the 10x Genomics Fresh Frozen Mouse Brain Replicates Xenium dataset was obtained from 10x Genomics (https://www.10xgenomics.com/datasets/fresh-frozen-mouse-brain-replicates-1-standard). The H&E image and paired Visium spatial gene-expression measurements for control Replicate 1 were obtained from the 10x Genomics Visium CytAssist Gene Expression Libraries of Post-Xenium Mouse Brain (FF) dataset (https://www.10xgenomics.com/datasets/visium-cytassist-gene-expression-libraries-of-post-xenium-mouse-brain-ff-using-the-mouse-whole-transcriptome-probe-set-2-standard). The Xenium In Situ Human Breast Cancer data, comprising In Situ Sample 1 Replicates 1 and 2 from serial sections of the same formalin-fixed, paraffin-embedded tissue block, were obtained from 10x Genomics and were originally described by Janesick et al. [42] (https://www.10xgenomics.com/products/xenium-in-situ/preview-dataset-human-breast). The MERFISH Mouse Brain Receptor Map datasets were obtained from Vizgen (https://info.vizgen.com/mouse-brain-map). The processed coronal mouse brain aging MERFISH dataset was obtained from the Zenodo record “Processed MERFISH Datasets for Brain Aging (Coronal, Sagittal) and Rejuvenation (Exercise, Partial Reprogramming)” (https://doi.org/10.5281/zenodo.13883177). The P22 mouse brain spatial ATAC–RNA-seq data originally reported by Zhang et al. [6] were obtained from the UCSC Cell Browser (https://brain-spatial-omics.cells.ucsc.edu) and the Gene Expression Omnibus under accession GSE205055. The processed data and manual anatomical annotations used in the COSMOS study [49] were obtained from Zenodo (https://doi.org/10.5281/zenodo.13932144). The Allen Mouse Brain Common Coordinate Framework version 3 (CCFv3) annotation volume was obtained from the Allen Institute (https://download.alleninstitute.org/informatics-archive/current-release/mouse_ccf/annotation/ccf_2022/). The Allen Institute Mouse Whole Cortex and Hippocampus 10x cell metadata and HDF5 gene-expression matrix used to construct the RCTD reference were obtained from the Allen Brain Map data portal [46, 47] (https://brain-map.org/our-research/cell-types-taxonomies/cell-types-database-rna-seq-data/mouse-whole-cortex-and-hippocampus-10x). The Nissl-stained reference section (Mouse, P56, Coronal; atlas image 100960252, section 269) was obtained from the Allen Mouse Brain Atlas (https://mouse.brain-map.org/static/atlas).

## 7 Code Availability

The spAlignDE Python package, including the source code, executable notebooks, interactive region-pairing interface, testing suite and environment specifications, is publicly available at https://github.com/dsong-lab/spAlignDE/. Comprehensive documentation and step-by-step tutorials are available at https://dsong-lab.github.io/spAlignDE/. The tutorials describe the input and output formats, model parameters and example workflows for the main functionalities of spAlignDE. The source notebooks for reproducing the analyses are available at https://github.com/dsong-lab/spAlignDE/tree/main/source_notebooks, and the archived source code and materials for reproducing the results are available in the Zenodo repository at https://zenodo.org/records/21983769. All datasets analyzed in this study are publicly available from the repositories listed in the Data Availability section. Large datasets, the Allen CCF annotation volume and pretrained HIPT checkpoints are not redistributed with the software; their sources and required input formats are provided in the corresponding tutorials.

## 9 Acknowledgements

The authors appreciate the comments and feedback from members of the DS Lab at UConn Health (https://dsong-lab.github.io/).

