## Supplementary Material for "spAlignDE unifies cross-sample and cross-modal spatial alignment with mismatch-aware differential expression"

#### S1 Software environments and reproducibility

##### S1.1 Software environments

All spAlignDE analyses were performed using spAlignDE 0.1.0 under Python 3.10.14. The specified reference environment included PyTorch 2.10.0 and torchvision 0.25.0 with CUDA 12.8, AnnData 0.10.9, NumPy 1.26.4, pandas 2.3.3, SciPy 1.10.1, Scanpy 1.10.3, scikit-image 0.24.0, scikit-learn 1.7.2, HarmonyPy 0.2.0 and pyBANKSY 1.3.4. RCTD annotations were generated using `spacexr` 1.2.0 under R 4.5.2. The complete environment specifications and details of the externally obtained model assets are distributed with the source code.

##### S1.2 Random seeds and repeated execution

All executable spAlignDE workflows used fixed workflow-level random seeds set before the first stochastic operation. Seed 1000 was used for cross-sample joint clustering and alignment, including the mouse-brain, kidney, aging-brain and breast-cancer analyses, and for the ten-replicate cross-sample transformation-stability workflow after replicate-specific subsampling. Seed 1234 was used for single-sample MERFISH and spatial ATAC-seq clustering and for automatic and user-guided atlas alignment and ATAC-to-ST alignment. Seed 0 was used for H&E feature processing, image-region clustering and ST-to-H&E alignment, as well as for Nissl image processing and MERFISH-to-Nissl alignment. Seed 1 was used for stochastic post-alignment inference and RCTD reference-cell subsampling. The H&E analysis used the same fixed, preclustered Xenium ST input as the public tutorial; seed 0 therefore controls the image-side processing, image clustering and downstream alignment conditional on this fixed ST input.

Each workflow-level seed initialized the Python, NumPy and PyTorch CPU and CUDA random-number generators before BANKSY feature construction and randomized principal component analysis (PCA). Explicit random states were additionally supplied to randomized PCA, Harmony, graph-partitioning, $k$-means and stochastic-sampling routines. General joint-clustering analyses used the `leidenalg` backend with `n_iterations=-1`, and the dedicated breast-cancer Xenium analysis used the Scanpy `igraph` backend with `n_iterations=2`. A change in the clustering backend was treated as a change in the analysis configuration.

Independent reruns used the same input data, observation order, workflow parameters and random seed. These runs retained the same cluster labels, hierarchy memberships and accepted structure pairs and reproduced the reported summary results. Small numerical differences in GPU-derived deformation coordinates did not change these results.

#### S2 Detailed alignment methods

##### S2.1 Spatial-data preprocessing and structure construction

The following subsections provide the dataset representations and parameters used to construct spatial structures for alignment.

**S2.1.1 Spatial transcriptomics** Spatial transcriptomic structures were constructed from a cell- or spot-by-gene count matrix and the corresponding two-dimensional coordinates. For cross-sample alignment, genes were matched by name across samples, and genes absent from an individual sample were assigned zero values in that sample. BANKSY features were then calculated separately within each sample so that spatial neighborhoods did not connect observations from different tissue sections [1]. Unless otherwise stated, 30 spatial neighbors and a BANKSY mixing parameter of 0.8 were used. The sample-specific BANKSY representations were concatenated, reduced by joint principal component analysis (PCA) and corrected with Harmony using the sample identifier as the batch variable [2]. A shared-nearest-neighbor graph was constructed from the Harmony-corrected BANKSY representation, followed by Leiden clustering to obtain a common set of spatial-domain labels across the query and reference sections. In the primary cross-sample benchmark, 20 principal components, 50 neighbors for the shared-nearest-neighbor graph, and a Leiden resolution of 1.4 were used. The cross-sample workflow used random seed 1000, initialized before BANKSY feature construction, joint PCA, Harmony integration and Leiden clustering. General joint-clustering analyses used the `leidenalg` backend with `n_iterations=-1`; the dedicated breast-cancer Xenium workflow used the Scanpy `igraph` backend with `n_iterations=2`. The effect of the Leiden resolution was evaluated separately in the robustness analysis.

Dataset-specific cross-sample clustering settings were as follows. The mouse-brain analysis used a BANKSY mixing parameter of 0.8, 20 principal components, 50 shared-nearest-neighbor graph neighbors, Harmony $\theta = 2$ with 30 iterations, and Leiden resolution 1.4. The kidney analysis used a BANKSY mixing parameter of 0.2, 30 principal components, 100 graph neighbors, Harmony $\theta = 2$ with 20 iterations, and Leiden resolution 0.2. The aging-brain analysis used a BANKSY mixing parameter of 0.8, 20 principal components, 50 graph neighbors, Harmony $\theta = 2$ with 30 iterations, and Leiden resolution 0.8. The breast-cancer analysis used a BANKSY mixing parameter of 0.2, 30 principal components, 50 graph neighbors, Harmony $\theta = 4$ with 30 iterations, and Leiden resolution 0.3; Leiden used the `igraph` backend with two iterations, and boundary refinement was disabled.

For cross-modality alignment, spatial transcriptomic structures were obtained by applying BANKSY to each ST sample independently, without Harmony integration. The selected BANKSY labels defined the finest ST structure partition. When a coarse-to-fine alignment was required, additional structure levels were constructed from the ST expression profiles by building a hierarchical tree. Raw gene expression counts were transformed using $\log(1+x)$, and variable genes were retained according to their cumulative variance, with at least 50 genes retained. Expression was averaged within each finest-level BANKSY cluster, standardized across clusters and hierarchically clustered using Ward linkage with Euclidean distance. Cutting this hierarchy into an increasing number of groups produced nested coarse-to-fine structure partitions, with the original BANKSY partition used at the final resolution. An optional boundary-aware refinement step reassigned locally inconsistent labels by distance-weighted voting among spatial neighbors, using smaller neighborhoods near estimated tissue boundaries to limit smoothing across anatomical interfaces. Mathematical details of the coarse-to-fine hierarchy and boundary-aware refinement are provided in Supplementary Methods S2.2. The single-sample MERFISH clustering used 20 principal components, 30 spatial neighbors, scaled-Gaussian neighborhood weighting, a BANKSY mixing parameter of 0.8, first-order neighborhood features ($m_{\max} = 1$), Leiden resolution 1.2 and boundary-aware label refinement. Random seed 1234 was initialized before BANKSY feature construction and randomized PCA. The H&E alignment instead used the same fixed, preclustered Xenium ST labels as the public tutorial.

**S2.1.2 Histology-image embedding and stain-specific spatial structure construction** Histology-derived spatial structures were constructed through four steps: image preparation, multiscale feature extraction, stain-specific tissue masking and clustering, and spatial refinement of the resulting image regions. The same image-preparation and feature-extraction framework was used for H&E and Nissl images, whereas the feature representation, tissue mask and clustering refinement were selected separately for each stain.

*Image preparation and multiscale feature extraction.* Histological images were represented as RGB images with intensity values scaled to $[0, 1]$. When the physical pixel size was available from image metadata or supplied explicitly, the image was resampled to $0.5\,\mu$m per pixel; otherwise, the native pixel grid was retained. The H&E and Nissl JPEG images used in this study did not contain embedded physical-pixel-size metadata and were therefore analyzed without rescaling. Each image was padded with white pixels so that both image dimensions were divisible by 224. Feature maps were represented on a common grid in which each grid location corresponded to a $16 \times 16$-pixel image region.

Pretrained HIPT encoders [3] were used to extract a 192-dimensional context-level embedding and a 384-dimensional subpatch-level embedding at each feature-grid location. To reduce sensitivity to the tiling origin, feature extraction was repeated using shifts of 0, 64, 128 and 192 pixels along each image axis, producing 16 shifted views. At locations represented in all shifted views, the corresponding embeddings were averaged across views. The context-level and subpatch-level feature maps were subsequently smoothed using square uniform filters with widths of 16 and 4 feature-grid units, respectively. Mean RGB intensities were also calculated for each $16 \times 16$-pixel region. The outer 256 image pixels were excluded from downstream clustering to reduce artifacts introduced by padding and shifted tiling.

*Candidate feature representations and selection.* Two candidate feature representations were retained for each histological image. The first contained only the HIPT-derived context, subpatch and RGB features and is denoted as VIT-only in the software outputs. The second combined dimension-reduced HIPT features with embeddings from the pretrained UNI pathology foundation model [4]. Within each image, the two candidates were processed using the same stain-specific tissue mask and clustering workflow. Candidate structure maps were compared according to spatial coherence, fragmentation and correspondence with visible tissue boundaries. The HIPT-only representation was retained for the reported H&E analysis, whereas the combined HIPT–UNI representation was retained for the reported Nissl analysis.

*H&E structure construction.* For the H&E image, a two-class K-means partition of channel-standardized RGB intensities was first used to separate tissue from background. Among classes occupying at least 1% of the valid feature grid, the darker class was selected as tissue, and only its largest connected component was retained.

Within the tissue mask, the smoothed HIPT context and subpatch embeddings were standardized and combined with mean RGB intensities and normalized spatial coordinates. RGB and coordinate contributions were weighted by 0.25 and 0.05, respectively. K-means clustering was then used to obtain 30 initial image regions. Clustering was run for 11 iterations with random seed 0, using at most 50,000 feature-grid locations for initialization and assignment batches of 50,000 locations.

Internal holes were assigned the label of the nearest classified location. Ward agglomerative clustering of the image-feature centroids reduced the initial partition from 30 to 26 meta-regions. The partition was then refined using the approximate bilateral symmetry of the coronal brain section. In the stored image orientation, the anatomical symmetry axis corresponded to an up–down array reflection. Regions containing at least 200 feature-grid locations were considered for reflection-based merging. Mutually selected region pairs were merged when their reflected intersection over union was at least 0.20 and their reflected Dice coefficient was at least 0.30, reducing the partition from 26 to 24 regions.

Starting from these 24 regions, up to three additional symmetry-supported merges were considered. Candidate regions were required to have a reflected Dice coefficient of at least 0.15 or a normalized reflected-centroid distance of at most 0.15, together with a feature-centroid cosine similarity of at least 0.30. A merge was accepted only when it increased the global bilateral label-overlap score by at least 0.02. Three merges were accepted, producing 21 regions. Finally, connected components smaller than 250 feature-grid locations were removed and reassigned using neighboring labels. The resulting 21 H&E-derived structures were used for structure pairing and ST-to-H&E alignment.

*Nissl structure construction.* For the Nissl image, a 1,024-dimensional embedding was extracted using UNI and combined with the HIPT representation. HIPT context, subpatch and RGB features were dimension-reduced as one block, whereas UNI embeddings, UNI-associated RGB values and normalized spatial coordinates were dimension-reduced as a second block. Principal component analysis was applied separately to the two blocks, retaining the smallest number of components required to explain at least 99% of the variance.

An adaptive brain mask was constructed from image color and brightness. Candidate tissue locations had HSV saturation above its 60th percentile and HSV value below its 90th percentile. The mask was refined by morphological closing with a disk of radius 10, opening with a disk of radius 6 and Gaussian smoothing with $\sigma = 1.95$. Holes smaller than 10,000 feature-grid locations and connected objects smaller than 7,056 locations were removed, and only the largest connected component was retained. Tissue support was further restricted to locations with mean RGB intensity below 0.965, followed by binary closing with a $7 \times 7$ kernel, hole filling and retention of the largest connected component. The outer 256 image pixels were excluded as in the H&E workflow.

Within the final tissue mask, the reduced HIPT and UNI feature blocks were standardized and combined with additional RGB and normalized-coordinate terms weighted by 0.25 and 0.05, respectively. K-means clustering with 25 clusters, 11 iterations and random seed 0 produced the initial Nissl partition. Connected components smaller than 300 feature-grid locations were removed and reassigned to the nearest retained region. One $3 \times 3$ neighborhood-majority update was then applied, requiring support from at least five neighboring locations. This refinement reduced the represented partition from 25 to 19 regions.

Ward agglomerative clustering of the 19 region centroids produced 18 meta-regions. The same feature- and symmetry-supported criteria used for the H&E image were then applied, with at most two additional merges. Both accepted merges satisfied the overlap, centroid-distance, feature-similarity and global-score-gain criteria, producing 16 regions. Finally, connected components smaller than 250 feature-grid locations were removed and reassigned using neighboring labels. The resulting 16 Nissl-derived structures were used for Nissl-to-ST structure pairing and alignment.

**S2.1.3 Spatial ATAC-seq** The spatial ATAC gene-activity scores used in this study were generated by Zhang et al. from spatial ATAC fragment files using the Gene Score model implemented in ArchR v1.0.1 [5, 6]. Spatial ATAC-seq structures were then constructed from this gene-activity matrix and the corresponding spatial coordinates. BANKSY was then applied as a single-sample clustering procedure using the gene-activity profiles and local spatial neighborhoods. For this analysis, we used 20 principal components, 30 spatial neighbors, scaled-Gaussian neighborhood weighting, a BANKSY mixing parameter of 0.6, first-order neighborhood features ($m_{\max} = 1$) and a Leiden resolution of 1.0. The resulting labels were optionally refined using the same boundary-aware spatial voting procedure described for single-sample ST data. Both the ATAC clustering and the independent clustering of the ST reference used random seed 1234, initialized before BANKSY feature construction and randomized PCA. The fixed-seed BANKSY partition contained 17 raw spatial structures. The `cluster_raw` labels were used for ATAC-to-ST structure pairing and alignment, whereas boundary-refined labels were retained only for quality control. Thus, ATAC and ST structures were defined without requiring a shared molecular feature space before structure pairing.

**S2.1.4 Anatomical reference atlases** For alignment to an anatomical atlas, spatial structures were obtained directly from the atlas annotation image and its accompanying hierarchical structure table rather than by data-driven clustering. In the Allen Common Coordinate Framework version 3 (CCFv3) analysis, a selected two-dimensional slice was extracted from the annotation volume, and each non-background annotation identifier was linked to its anatomical name, acronym and structure ID path in the Allen hierarchy metadata [7, 8]. Each prefix of a structure path represents an anatomical structure at a particular hierarchical depth. For every prefix represented in the selected slice, all descendant annotation labels present in that slice were combined to produce a candidate anatomical structure mask. Cortical-layer structures were additionally generated by combining labels annotated as layers 1, 2/3, 4, 5, 6a or 6b.

The atlas hierarchy defines candidate anatomical masks at multiple spatial scales. In the reported Allen CCFv3 analyses, all eligible hierarchy-prefix candidates from depths 2–10 were available at every alignment stage; the coarse-to-fine schedule was applied only to the ST structure partition. Although the Allen CCF was used as the default reference in this study, the same procedure can be applied to any other atlas provided that it supplies a rasterized annotation image or volume and a hierarchy table mapping each annotation identifier to its ancestral structure path.

#### S2.2 Coarse-to-fine ST structure hierarchy and boundary-aware refinement

*Coarse-to-fine structure hierarchy.* Consider $K$ ST samples indexed by $s \in \{1, \dots, K\}$. Sample $s$ contains $N_s$ cells or spots and $M$ genes, represented by the count matrix $X^{(s)} \in \mathbb{R}_{\geq 0}^{N_s \times M}$, with spatial coordinates $\mathbf{q}_i^{(s)} \in \mathbb{R}^2$. The following construction is performed independently for each sample; we therefore suppress the sample superscript for clarity. Let $X_{ig}$ denote the count of gene $g \in \{1, \dots, M\}$ in observation $i \in \{1, \dots, N_s\}$, and let $c_i \in \{1, \dots, R\}$ denote its finest-level BANKSY structure label, where $R$ is the number of finest-level structures. Counts are transformed as

$$Y_{ig} = \log(1 + X_{ig}).$$

For each gene, let $\sigma_g^2 = \mathrm{Var}_i(Y_{ig})$, and order genes such that $\sigma_{(1)}^2 \geq \cdots \geq \sigma_{(M)}^2$. Given a target cumulative variance fraction $\rho$, the number of retained genes is

$$M_{\text{ret}} = \min\left\{ M, \max\left[ M_{\min}, \min\left\{ p : \frac{\sum_{j=1}^{p} \sigma_{(j)}^2}{\sum_{g=1}^{M} \sigma_g^2} \geq \rho \right\} \right] \right\}.$$

The implementation uses $M_{\min} = 50$. For each finest-level structure $r$, a structure-level expression profile is calculated as

$$\bar{Y}_{rg} = \frac{1}{|C_r|} \sum_{i \in C_r} Y_{ig}, \qquad C_r = \{i : c_i = r\}.$$

Each retained gene is standardized across the $R$ structure profiles,

$$Z_{rg} = \frac{\bar{Y}_{rg} - \bar{Y}_{\cdot g}}{\sigma_g^{\text{structure}} + \epsilon},$$

where $\bar{Y}_{\cdot g}$ and $\sigma_g^{\text{structure}}$ are the mean and standard deviation across finest-level structures. Ward hierarchical clustering is then applied to the rows of $Z$ using Euclidean distance. For two groups of finest-level structures, $A$ and $B$, the Ward merge cost is

$$d_{\text{Ward}}(A, B) = \frac{|A||B|}{|A| + |B|} \left\| \bar{\mathbf{z}}_A - \bar{\mathbf{z}}_B \right\|_2^2,$$

where $\bar{\mathbf{z}}_A$ and $\bar{\mathbf{z}}_B$ are the mean standardized profiles of the two groups. Cutting the resulting tree at increasing structure numbers $R_1 < \cdots < R_L = R$ defines $L$ nested levels and mappings $h_\ell : \{1, \dots, R\} \to \{1, \dots, R_\ell\}$. The structure label of observation $i$ at level $\ell$ is

$$c_i^{(\ell)} = h_\ell(c_i).$$

By default, the intermediate $R_\ell$ values are approximately evenly spaced between 2 and $R$, constrained to be strictly increasing, and the original BANKSY labels are retained as the final-resolution partition.

*Boundary-aware label refinement.* To distinguish tissue-boundary observations from tissue-interior observations, spatial coordinates $\mathbf{q}_i$ are first min–max scaled and rasterized onto a $256 \times 256$ binary tissue grid. Morphological closing with a disk of radius 2 pixels and filling of holes smaller than 64 pixels are applied, followed by a Euclidean distance transform of the tissue mask. Observation $i$ is classified as a boundary observation when its rasterized position lies within $d_0$ pixels of the tissue boundary,

$$b_i = \mathbf{1}\{D_{\text{tissue}}(\mathbf{q}_i) \leq d_0\}.$$

The default boundary threshold is $d_0 = 3$ pixels. A smaller neighborhood is used at the boundary,

$$\nu_i = \begin{cases} \nu_{\text{boundary}}, & b_i = 1, \\ \nu_{\text{interior}}, & b_i = 0, \end{cases}$$

with default values $\nu_{\text{boundary}} = 8$ and $\nu_{\text{interior}} = 150$. Excluding observation $i$ itself, the inverse-distance vote for candidate label $c$ is

$$V_i(c) = \sum_{j \in \mathcal{N}_{\nu_i}(i)} \frac{\mathbf{1}\{c_j = c\}}{\|\mathbf{q}_i - \mathbf{q}_j\|_2 + \epsilon}, \qquad P_i(c) = \frac{V_i(c)}{\sum_{c'} V_i(c')}.$$

Let $c_i^* = \arg\max_c P_i(c)$ be the locally supported label. The original label $c_i$ is replaced by $c_i^*$ only when the winning label differs from the original label, support for the original label is weak, and support for the new label exceeds a location-dependent threshold:

$$c_i^{\text{ref}} = \begin{cases} c_i^*, & c_i^* \neq c_i, \quad P_i(c_i) < \tau_{\text{keep}}, \quad P_i(c_i^*) \geq \tau_i, \\ c_i, & \text{otherwise.} \end{cases}$$

The default values are $\tau_{\text{keep}} = 0.8$, $\tau_i = 0.2$ for interior observations and $\tau_i = 0.8$ for boundary observations. For user-specified protected structures, $\tau_i = 0.9$ is used. The smaller boundary neighborhood and higher reassignment threshold reduce label propagation across thin or sharply separated anatomical structures.

#### S2.3 Detailed global pre-alignment and structure-mask construction

**S2.3.1 Global pre-alignment** Before structure pairing and S-LDDMM alignment, spAlignDE places the query and reference datasets in approximately the same coordinate system. This global pre-alignment corrects major differences in tissue position, orientation and size by shifting, rotating and scaling the query coordinates while keeping the reference coordinates fixed. It provides only a coarse initialization and does not attempt to model local anatomical deformation, consistent with the standard distinction between global initialization and deformable alignment [9, 10].

Global pre-alignment is important because overlap- and geometry-based measures, such as Dice similarity and intersection over union (IoU), more meaningfully reflect structural correspondence once the query and reference tissues have been brought into approximate spatial agreement. It also allows S-LDDMM to focus on local deformation rather than first recovering large differences in global tissue position, orientation or scale.

spAlignDE provides three pre-alignment strategies for different data settings. In the primary cross-sample ST-to-ST analyses, rotation, scaling and shifting were estimated automatically from the centroids of jointly identified spatial structures. For ST-to-atlas and Xenium-to-H&E alignment, the global transformation was estimated automatically by maximizing the IoU between the query and reference whole-tissue masks. Interactive manual pre-alignment was used for MERFISH-to-Nissl and ATAC-to-ST alignment and for the Xenium breast-cancer ST-to-ST example in which automatic initialization was less reliable.

*Shared-structure-centroid pre-alignment.* For cross-sample spatial transcriptomics alignment, jointly inferred spatial-structure labels provide direct correspondence between the query and reference samples. Let $z \in \{\text{qry}, \text{ref}\}$ denote the dataset role. For observation $i$ in dataset $z$, let $\mathbf{q}_i^z \in \mathbb{R}^2$ denote its two-dimensional spatial coordinate and $c_i^z$ its spatial-structure label. Let $\mathcal{R}_{\text{shared}}$ denote the set of structure labels represented in both datasets.

For structure $r \in \mathcal{R}_{\text{shared}}$, its centroid in dataset $z$ is defined as

$$\boldsymbol{\mu}_r^z = \frac{1}{n_r^z} \sum_{i:c_i^z = r} \mathbf{q}_i^z,$$

where $n_r^z$ is the number of observations assigned to structure $r$ in dataset $z$, and $\boldsymbol{\mu}_r^z \in \mathbb{R}^2$ is their mean spatial position.

A weighted global alignment is estimated from the corresponding query and reference structure centroids by extending the standard least-squares similarity-alignment formulation:

$$(\widehat{s}, \widehat{\mathbf{R}}, \widehat{\mathbf{t}}) = \arg \min_{\substack{s>0,\ \mathbf{R}\in \text{SO}(2), \\ \mathbf{t}\in\mathbb{R}^2}} \sum_{r \in \mathcal{R}_{\text{shared}}} \omega_r \left\| s\mathbf{R}\boldsymbol{\mu}_r^{\text{qry}} + \mathbf{t} - \boldsymbol{\mu}_r^{\text{ref}} \right\|_2^2 .$$

Here, $s > 0$ is the global scaling factor, $R$ is a two-dimensional rotation matrix, and $\mathbf{t} \in \mathbb{R}^2$ shifts the query coordinates horizontally and vertically. Reflection is excluded. The notation $\|\cdot\|_2$ denotes Euclidean distance, and the hats indicate the estimated transformation parameters.

The weight assigned to structure $r$ is

$$\omega_r = \frac{\min\left(n_r^{\text{qry}}, n_r^{\text{ref}}\right)}{\sum_{r' \in \mathcal{R}_{\text{shared}}} \min\left(n_{r'}^{\text{qry}}, n_{r'}^{\text{ref}}\right)}.$$

Thus, structures supported by more observations contribute more strongly, while a structure that is large in only one dataset cannot dominate the transformation. The weights are normalized to sum to one. The estimated rotation, scaling and shifting are subsequently applied to all query observations, including those not used to calculate the shared-structure centroids.

*Whole-tissue-mask-overlap pre-alignment.* When direct correspondence between individual spatial structures is unavailable but reliable whole-tissue supports can be constructed, spAlignDE estimates the global pre-alignment by maximizing the overlap between the query and reference tissue masks. This strategy was used for automatic ST-to-atlas alignment, where the overall tissue boundaries are comparable even before individual transcriptomic and anatomical structures have been paired.

For the atlas reference, the whole-tissue mask consists of all non-background pixels in the selected atlas annotation section. For the ST query, spatial observations are rasterized onto the atlas pixel grid. Morphological closing connects nearby occupied pixels and removes small gaps caused by sparse spatial sampling, followed by dilation to produce a continuous query-tissue support. This yields a binary reference mask $\mathcal{M}^{\mathrm{ref}}_{\mathrm{tissue}}$ and a transformed query mask $\mathcal{M}^{\mathrm{qry}}_{\mathrm{tissue}}(s,\theta)$ on the same raster domain $\Omega$, where $s$ and $\theta$ denote the candidate scaling factor and rotation angle, respectively.

The overlap of the two masks is measured using intersection over union,

$$\mathrm{IoU}(s,\theta) = \frac{\left|\mathcal{M}^{\mathrm{qry}}_{\mathrm{tissue}}(s,\theta) \cap \mathcal{M}^{\mathrm{ref}}_{\mathrm{tissue}}\right|}{\left|\mathcal{M}^{\mathrm{qry}}_{\mathrm{tissue}}(s,\theta) \cup \mathcal{M}^{\mathrm{ref}}_{\mathrm{tissue}}\right|},$$

where $|\cdot|$ denotes the number of foreground pixels. IoU ranges from zero, indicating no tissue overlap, to one, indicating identical tissue supports.

To initialize the search, an ellipse is fitted to the largest outer contour of each tissue mask. The fitted tissue centers determine the horizontal and vertical shifting, whereas the ratio of the longer ellipse axes provides an initial scaling factor. Candidate rotations and scaling factors are then evaluated by transforming the query coordinates, reconstructing the query-tissue mask, and calculating its IoU with the atlas mask. In the reported analyses, rotations were evaluated over the full 360° range at 1° intervals, and five scaling factors were evaluated within ±5% of the initial estimate. Reflection was not considered.

The candidate producing the largest IoU was retained as the global pre-alignment. Its rotation, scaling and shifting were applied to all query observations, including observations not required for tissue-mask construction. This procedure establishes approximate whole-tissue correspondence without requiring prior matches between individual ST structures and atlas regions. Structure pairing and S-LDDMM subsequently refine the internal anatomical correspondence.

*Interactive manual pre-alignment.* When reliable shared-structure centroids or comparable whole-tissue masks are unavailable, spAlignDE provides an interactive tool for manual global pre-alignment. A live overlay of the query and reference datasets allows users to adjust scaling, rotation, and horizontal and vertical shifting until the overall tissue boundaries and recognizable anatomical landmarks are approximately aligned (Supplementary Fig. S1).

The selected parameters are saved and applied consistently to all query coordinates and query-derived spatial structures, allowing the same initialization to be reproduced in subsequent analyses. This step specifies only the approximate global position of the query tissue. It does not define structure correspondences or estimate local anatomical deformation. By reducing large global differences before structure pairing, it makes overlap-based quantities such as Dice similarity and IoU more informative and provides a stable initialization for S-LDDMM, which subsequently estimates the finer affine–diffeomorphic alignment [11–13]. In this study, interactive manual pre-alignment was used for MERFISH-to-Nissl and spatial ATAC-to-ST alignment and for the Xenium breast-cancer ST-to-ST example. The Xenium-to-H&E analysis instead used automatic whole-tissue-mask-overlap pre-alignment.

For the reported breast-cancer alignment, the Replicate-2 query initialization used scale 1.0, rotation 2°, and translation $(-250, 1750)$. For ATAC-to-ST alignment, the ST reference used scale 1.0 and rotation $-125°$, whereas the ATAC query used scale 1.6, rotation $-90°$, and translation $(-2600, -3300)$. The left half of the ST reference was retained at the median x-coordinate, and the common raster used scale 0.25 with ten-pixel padding. The MERFISH-to-Nissl initialization used scale 0.0651186, rotation $-12.7545°$, and translation $(-32.5316, 88.3956)$ in histology feature-grid coordinates.

**S2.3.2 Structure-mask construction** Following global pre-alignment, all query and reference structures were represented on a common two-dimensional raster grid. For each structure $r$ in dataset $z$, spAlignDE constructed a binary mask $M^z_r$, in which foreground pixels represent the spatial region occupied by that structure and background pixels represent the remaining field of view. Separate procedures were used for structures represented by discrete spatial observations and structures already represented as labeled image regions.

*Structure-wise filtering of point-based spatial data.* For spatial transcriptomics and spatial ATAC-seq, each structure initially consists of a set of observations (cells, spots, or bins) with the same structure label. Because isolated observations may generate artificial mask islands or substantially enlarge the inferred boundary, local spatial density was evaluated separately within each structure. For observation $i$, a local-spacing statistic was calculated as the mean distance to its $k$ nearest neighbors carrying the same structure label,

$$d_i = \frac{1}{k} \sum_{j \in \mathcal{N}_k(i)} \|\mathbf{q}_i - \mathbf{q}_j\|_2 \,,$$

where $\mathbf{q}_i$ is the pre-aligned coordinate of observation $i$ and $\mathcal{N}_k(i)$ denotes its within-structure nearest neighbors.

The distribution of $d_i$ was summarized using its median and median absolute deviation (MAD). Observations satisfying

$$d_i \leq \mathrm{median}(d) + \alpha\ \mathrm{MAD}(d)$$

were retained for mask construction, where $\alpha$ controls the degree of filtering. When the MAD was zero, an upper quantile of the local-spacing distribution was used instead. Structures containing too few observations for reliable neighborhood estimation were retained without filtering.

Filtering strength was adapted to structure size. A robust spatial extent was estimated from the central coordinate range of each structure, reducing the influence of isolated points. Structures below a modality-specific extent quantile were treated as spatially detailed structures and processed using settings that preserve narrow regions and fine boundaries. Optional grid-based thinning retained one locally representative observation within each occupied grid cell, thereby reducing oversampling in densely populated regions without changing the overall spatial coverage. Observations excluded during these steps were omitted only from mask construction; they remained in the original dataset and received the final estimated transformation.

For spatial ATAC-to-ST alignment, local-density statistics were additionally used to distinguish slender structures from broader structures. Structures with a comparatively high fraction of locally isolated observations were assigned a slender-mask profile with weaker smoothing and boundary refinement. This classification did not alter the structure labels or remove observations from the final aligned dataset.

*Rasterization and adaptive spatial smoothing.* Let $\Pi(\mathbf{q}_i)$ denote the pixel corresponding to coordinate $\mathbf{q}_i$ on the common grid $\Omega$. The initial occupancy image for structure $r$ was defined as

$$H_r^z(\mathbf{u}) = \sum_{i:c_i^z=r} \mathbf{1}\{\Pi(\mathbf{q}_i^z) = \mathbf{u}\}, \qquad \mathbf{u} \in \Omega,$$

where $c_i^z$ is the structure label of observation $i$. Because the observations provide discrete samples of an underlying tissue region, the occupancy image was convolved with a Gaussian kernel. The smoothing bandwidth was adapted to the median within-structure point spacing,

$$\sigma_r^z = \mathrm{clip}\left(\gamma \ \mathrm{median}_i \, d_{i,(4)}, \sigma_{\min}, \sigma_{\max}\right),$$

where $d_{i,(4)}$ is the distance from observation $i$ to its fourth nearest within-structure neighbor, $\gamma$ is a scale factor, and $\sigma_{\min}$ and $\sigma_{\max}$ prevent excessive under- or over-smoothing. The smoothed occupancy image was normalized by its maximum and thresholded to obtain an initial binary mask. Detailed or slender structures used a smaller smoothing range and a more restrictive threshold than broader structures to reduce unintended merging of nearby anatomical regions.

*Morphological and connected-component refinement.* Initial masks were refined using standard binary morphological operations. Closing connected small gaps and discontinuities, whereas opening removed isolated boundary noise. Internal holes were optionally filled, and Gaussian boundary smoothing reduced pixel-scale irregularities. Components smaller than a modality-specific area threshold were removed. When a mask remained highly fragmented, connected components were ranked by area and only the largest components accounting for a predefined fraction of the total mask area were retained. Different refinement profiles were used for broad and spatially detailed structures so that narrow anatomical features were not eliminated by aggressive smoothing or component removal.

*Masks derived from histology and anatomical labels.* Histology and atlas structures were already defined on image grids and therefore did not require point rasterization. Given a label image $\Lambda^z$ and a set of labels $\mathcal{A}_r^z$ defining structure $r$, the corresponding mask was constructed as

$$M_r^z(\mathbf{u}) = \mathbf{1}\left\{\Lambda^z(\mathbf{u}) \in \mathcal{A}_r^z\right\}.$$

For histological images, small labeled regions were removed before mask construction, after which closing, opening, hole filling and boundary smoothing were applied. Small disconnected islands were removed relative to the area of the largest connected component. For anatomical atlases, one mask could contain either a single annotation label or the union of multiple descendant labels selected from the anatomical hierarchy. Atlas boundaries were otherwise retained directly from the annotation image.

The resulting masks provide a common representation of structure location, extent and boundary geometry across all modalities. They were used only for structure-pair scoring and for constructing the matched channels supplied to S-LDDMM; the final estimated transformation was applied to every observation in the original query dataset.

##### S2.4 Automatic structure-pairing metrics and modality-specific settings

For cross-modality structure pairing, let $\mathcal{M}_i^{\mathrm{qry}}$ and $\mathcal{M}_j^{\mathrm{ref}}$ denote the processed binary masks of query structure $i$ and reference structure $j$, respectively. spAlignDE evaluates each candidate using five possible similarity components, $\mathcal{C} = \{\mathrm{Dice}, \mathrm{SDF}, \mathrm{Ch}, \mathrm{area}, \mathrm{thick}\}$. Each component $C_{ij}^{(m)}$ lies in $[0, 1]$, with larger values indicating stronger correspondence. The modality-specific composite score is

$$S_{ij}^{\mathrm{comp}} = \sum_{m\in\mathcal{C}} w_m C_{ij}^{(m)}, \qquad w_m \geq 0, \qquad \sum_{m\in\mathcal{C}} w_m = 1.$$

Components that are not used for a modality pair have weight zero. Throughout this subsection, $d(\mathbf{u}, \mathcal{A})$ denotes the shortest Euclidean distance from raster location $\mathbf{u}$ to set $\mathcal{A}$, and $\epsilon > 0$ is a small constant that prevents numerical instability when an area or thickness estimate approaches zero.

*Dice overlap.* The Dice coefficient measures the direct spatial overlap between two binary structure masks:

$$C_{ij}^{\mathrm{Dice}} = \frac{2|\mathcal{M}_i^{\mathrm{qry}} \cap \mathcal{M}_j^{\mathrm{ref}}|}{|\mathcal{M}_i^{\mathrm{qry}}| + |\mathcal{M}_j^{\mathrm{ref}}|}.$$

It ranges from zero to one, with one indicating identical masks and zero indicating no spatial overlap. Dice similarity provides an intuitive measure of whether two structures occupy the same spatial region. Because it depends directly on mask intersection, it is most informative after the datasets have undergone a reasonable global pre-alignment.

*Signed-distance-field agreement.* Binary-mask overlap alone does not indicate how far non-overlapping boundaries are separated or whether two structures have similar surrounding geometry. For structure-pair evaluation, spAlignDE defines the signed distance field

$$\phi_{\mathrm{pair}}(\mathcal{M})(\mathbf{u}) = d(\mathbf{u}, \mathcal{M}^c) - d(\mathbf{u}, \mathcal{M}),$$

where $\mathcal{M}^c$ is the mask complement. Under this convention, $\phi_{\mathrm{pair}}$ is positive for locations inside the mask and negative for locations outside the mask, with its magnitude reflecting distance from the mask interface.

The union of the two boundary bands with half-width $h_{\mathrm{SDF}} > 0$ is

$$\mathcal{B}_{ij}(h_{\mathrm{SDF}}) = \left\{\mathbf{u} : |\phi_{\mathrm{pair}}(\mathcal{M}_i^{\mathrm{qry}})(\mathbf{u})| \le h_{\mathrm{SDF}} \text{ or } |\phi_{\mathrm{pair}}(\mathcal{M}_j^{\mathrm{ref}})(\mathbf{u})| \le h_{\mathrm{SDF}}\right\}.$$

Signed-distance-field agreement is defined as

$$C_{ij}^{\mathrm{SDF}} = \frac{1 + \mathrm{corr}_{\mathbf{u}\in\mathcal{B}_{ij}(h_{\mathrm{SDF}})}\left[\phi_{\mathrm{pair}}\left(\mathcal{M}_i^{\mathrm{qry}}\right)(\mathbf{u}), \phi_{\mathrm{pair}}\left(\mathcal{M}_j^{\mathrm{ref}}\right)(\mathbf{u})\right]}{2}.$$

The affine transformation $(1+\rho)/2$ maps the boundary-band correlation $\rho \in [-1, 1]$ to $[0, 1]$. If the band contains fewer than 10 raster locations or either signed-distance vector has zero variance, the implementation assigns zero similarity. Restricting the calculation to the boundary bands prevents distant interior and background locations from dominating the correlation. A smaller $h_{\mathrm{SDF}}$ emphasizes local boundary geometry, whereas a larger value incorporates broader interior and exterior shape information.

*Chamfer boundary similarity.* The bidirectional Chamfer distance measures the average nearest-neighbor separation between the two structure boundaries. Let $\partial\mathcal{M}$ denote the set of boundary pixels of mask $\mathcal{M}$. The distance is

$$d_{ij}^{\mathrm{Ch}} = \frac{1}{2}\left[\frac{1}{|\partial\mathcal{M}_i^{\mathrm{qry}}|}\sum_{\mathbf{x}\in\partial\mathcal{M}_i^{\mathrm{qry}}} d(\mathbf{x}, \partial\mathcal{M}_j^{\mathrm{ref}}) + \frac{1}{|\partial\mathcal{M}_j^{\mathrm{ref}}|}\sum_{\mathbf{y}\in\partial\mathcal{M}_j^{\mathrm{ref}}} d(\mathbf{y}, \partial\mathcal{M}_i^{\mathrm{qry}})\right].$$

Using both directions prevents the comparison from depending on which mask is designated as the query or reference and penalizes cases in which one boundary covers only a subset of the other. The distance is converted into a similarity by

$$C_{ij}^{\mathrm{Ch}} = \exp\left(-d_{ij}^{\mathrm{Ch}}/\tau_{\mathrm{Ch}}\right),$$

where $\tau_{\mathrm{Ch}} > 0$ is a decay scale in pairing-raster units. Increasing $\tau_{\mathrm{Ch}}$ makes the similarity more tolerant of boundary displacement, whereas decreasing it makes the boundary comparison stricter. This transformation maps non-negative distances to $(0, 1]$.

*Area similarity.* Area similarity evaluates whether the two masks have comparable spatial extents:

$$C_{ij}^{\mathrm{area}} = \exp\left[-\left|\log\frac{|\mathcal{M}_i^{\mathrm{qry}}| + \epsilon}{|\mathcal{M}_j^{\mathrm{ref}}| + \epsilon}\right|\right].$$

This symmetric measure equals one when the masks have equal area and decreases as their relative sizes diverge. Unlike Dice similarity, area similarity does not require direct spatial overlap. It therefore prevents a small structure from being paired with a substantially larger structure solely because their boundaries or centroids are partially aligned.

*Thickness consistency.* For thin or layer-like anatomical structures, similar area and boundary placement do not necessarily imply similar width. spAlignDE summarizes characteristic thickness using the within-mask Euclidean distance transform:

$$T_q(\mathcal{M}) = Q_q\left\{d(\mathbf{u}, \mathcal{M}^c) : \mathbf{u} \in \mathcal{M}\right\},$$

where $d(\mathbf{u}, \mathcal{M}^c)$ is the Euclidean distance from an interior raster location to the nearest location outside the mask. Thickness consistency is

$$C_{ij}^{\mathrm{thick}} = \exp\left[-\left|\log\frac{T_q(\mathcal{M}_i^{\mathrm{qry}}) + \epsilon}{T_q(\mathcal{M}_j^{\mathrm{ref}}) + \epsilon}\right|\right].$$

This similarity equals one for equal characteristic thickness and decreases when one structure is substantially wider or narrower than the other. The atlas–ST workflow uses $q = 0.75$, which is less sensitive than the maximum to isolated thick locations while representing the wider support of a structure more strongly than the median.

*Average symmetric surface distance for quality control.* Average symmetric surface distance (ASD) is retained as a raw boundary-distance quality-control measure. Let

$$n_i = |\partial \mathcal{M}_i^{\mathrm{qry}}|, \qquad n_j = |\partial \mathcal{M}_j^{\mathrm{ref}}|.$$

The ASD is

$$d_{ij}^{\mathrm{ASD}} = \frac{\sum_{\mathbf{x} \in \partial \mathcal{M}_i^{\mathrm{qry}}} d(\mathbf{x}, \partial \mathcal{M}_j^{\mathrm{ref}}) + \sum_{\mathbf{y} \in \partial \mathcal{M}_j^{\mathrm{ref}}} d(\mathbf{y}, \partial \mathcal{M}_i^{\mathrm{qry}})}{n_i + n_j}.$$

The bidirectional Chamfer distance assigns equal weight to the two directed mean distances, whereas ASD pools all directed boundary distances and therefore weights the two directions according to their numbers of boundary pixels. They are identical when the masks have equal numbers of boundary pixels and are generally strongly related when their boundary lengths are similar. To avoid including two highly overlapping boundary-distance similarities in the composite score, spAlignDE uses only $C_{ij}^{\mathrm{Ch}}$ as a weighted component. The raw $d_{ij}^{\mathrm{ASD}}$, which remains in pairing-raster units and is not normalized to $[0, 1]$, is used only as an absolute QC threshold for histology–ST and atlas–ST pairing.

*Modality-specific score composition and acceptance criteria.* All reported pairing scores used non-negative weights that sum to one. The SDF band half-width was $h_{\mathrm{SDF}} = 20$ pairing-raster units in all three cross-modality workflows. Distance-dependent parameters and ASD thresholds are expressed in the corresponding pairing-raster units and should be rescaled if the raster resolution is changed.

*Histology–ST.* The histology–ST composite score was

$$S_{ij}^{\mathrm{hist}} = 0.20C_{ij}^{\mathrm{SDF}} + 0.40C_{ij}^{\mathrm{Ch}} + 0.15C_{ij}^{\mathrm{Dice}} + 0.25C_{ij}^{\mathrm{area}},$$

with $\tau_{\mathrm{Ch}} = 30$. The relatively high Chamfer weight emphasizes boundary agreement between the histology- and ST-derived structures, whereas the raw ASD provides an independent absolute-distance check. Candidate masks were required to intersect by at least 20 raster locations before full scoring. A candidate was accepted when $S_{ij}^{\mathrm{hist}} \geq 0.40$ and $d_{ij}^{\mathrm{ASD}} \leq 30$. No additional soft penalty was applied, so the final pairing score equaled the composite score.

*ATAC–ST.* The ATAC–ST composite score was

$$S_{ij}^{\mathrm{ATAC}} = 0.35C_{ij}^{\mathrm{SDF}} + 0.25C_{ij}^{\mathrm{Ch}} + 0.10C_{ij}^{\mathrm{Dice}} + 0.30C_{ij}^{\mathrm{area}},$$

with $\tau_{\mathrm{Ch}} = 30$. A soft boundary-distance gate was defined as

$$G_{ij}^{\mathrm{Ch}} = \left[1 + \exp\left(\frac{d_{ij}^{\mathrm{Ch}} - 16}{6}\right)\right]^{-1}, \qquad S_{ij}^{\mathrm{final}} = S_{ij}^{\mathrm{ATAC}}\left(0.5 + 0.5G_{ij}^{\mathrm{Ch}}\right).$$

Here, 16 is the center of the logistic boundary-distance penalty and 6 controls the smoothness of its transition, both in pairing-raster units. Thus, candidates with larger boundary separation were downweighted smoothly without being reduced below one half of their composite score by this gate alone. Candidates were accepted when $S_{ij}^{\mathrm{final}} \geq 0.21$ and $C_{ij}^{\mathrm{Dice}} \geq 0.01$.

*Atlas–ST.* The atlas–ST composite score was

$$S_{ij}^{\mathrm{atlas}} = 0.05C_{ij}^{\mathrm{SDF}} + 0.05C_{ij}^{\mathrm{Ch}} + 0.20C_{ij}^{\mathrm{Dice}} + 0.50C_{ij}^{\mathrm{area}} + 0.20C_{ij}^{\mathrm{thick}},$$

with $\tau_{\mathrm{Ch}} = 25$ and thickness quantile $q = 0.75$. To reduce the scores of candidates with weak overlap, SDF agreement or thickness consistency, define

$$g(x; t) = \begin{cases} 1, & x \geq t, \\ \max(x/t, 0.45), & x < t. \end{cases}$$

Here, $x$ is the corresponding component similarity, $t$ is its soft threshold, and 0.45 is the minimum component-level penalty factor.

The atlas soft-gate factor and final score were

$$G_{ij}^{\mathrm{atlas}} = \max\left\{g(C_{ij}^{\mathrm{Dice}}; 0.25)g(C_{ij}^{\mathrm{SDF}}; 0.55)\left[g(C_{ij}^{\mathrm{thick}}; 0.65)\right]^{1.2}, 0.45\right\}, \qquad S_{ij}^{\mathrm{final}} = S_{ij}^{\mathrm{atlas}}G_{ij}^{\mathrm{atlas}}.$$

The thresholds 0.25, 0.55 and 0.65 apply to Dice, SDF and thickness similarity, respectively. The exponent 1.2 places additional emphasis on insufficient thickness agreement, and the outer maximum limits the overall soft-gate factor to a minimum of 0.45. Candidates were accepted when $S_{ij}^{\mathrm{final}} \geq 0.50$ and $d_{ij}^{\mathrm{ASD}} \leq 50$. After thresholding, candidates were ranked by their final pairing scores and selected greedily subject to one-to-one, non-overlapping structure use.

#### S2.5 Interactive cross-modality structure pairing

spAlignDE Structure Pair was implemented as a Streamlit interface with side-by-side query and reference panels (Supplementary Fig. S2). The interface accepts either a spatial point table in comma-separated format or a two-dimensional NumPy label image. For a point table, users specify the two coordinate columns and a cluster or structure-label column. In a label image, numeric values identify structures, whereas `NaN` and $-1$ represent background. The interface provides native support for the Allen CCF atlas. Users supply the Allen CCF 2022 annotation volume as an external input, after which the atlas slice and display orientation can be selected directly within the interface.

Point tables and label images can be rotated within the interface. For an anatomical atlas, users select the two-dimensional reference slice and can independently reverse its horizontal and vertical orientations. These display settings are stored in the exported correspondence table so that the selected structures can subsequently be reconstructed in the same coordinate configuration.

Each panel provides separate navigation and structure-selection modes. Selecting a point-table cluster, image label or atlas label activates the complete structure rather than only the clicked location. Multiple primitive structures can be combined into a named custom structure. In flexible-reuse mode, a primitive structure can contribute to more than one custom structure. In exclusive-assignment mode, a primitive structure that has already been assigned cannot be included in another custom structure. These options support both overlapping anatomical hypotheses during exploratory pairing and disjoint structure definitions when exclusive correspondences are required.

Users organize the selected query and reference structures into correspondence groups. Within each group, all selected primitive query structures are combined by pixelwise union, and all selected primitive reference structures are combined in the same way. The two resulting grouped masks define one query–reference correspondence. A group can therefore represent one-to-one, many-to-one, one-to-many or many-to-many relationships between primitive structures.

When multiple structures are selected on both sides, the exported table contains their pairwise row combinations under a shared `group_id`. Downstream processing treats all rows with the same `group_id` as one grouped correspondence rather than as independent pairwise channels. Thus, the correspondence group, rather than each individual exported row, is the unit supplied to alignment.

For each saved group, the exported comma-separated file records the query and reference dataset identities and input types, primitive and custom structure identifiers, the primitive labels included in each custom structure, the custom-structure reuse mode, point, pixel or voxel counts when applicable, and the display-orientation settings. In the reported ST-to-atlas workflow, primitive ST cluster identifiers and atlas annotation identifiers sharing the same `group_id` were de-duplicated and combined to construct the grouped query and reference masks. Groups lacking a non-empty mask on either side were discarded.

The retained grouped masks were converted into corresponding signed-distance-transform channels. Automatic structure-pair discovery was skipped, but all subsequent multichannel construction and S-LDDMM optimization steps were identical to those used for automatically identified pairs. The resulting channels were used for the final high-resolution S-LDDMM alignment.

#### S2.6 Construction of multichannel structural inputs for S-LDDMM

Following global pre-alignment, spAlignDE converts the query and reference structures into image-like inputs defined on the same regular two-dimensional pixel grid. The two inputs have the same spatial dimensions and channel order, so that corresponding pixels and channels represent comparable spatial information. Three types of structural channels are constructed. Shared-structure-composition channels, optionally augmented by a cell-density channel for single-cell datasets, are used for cross-sample spatial transcriptomics alignment, whereas paired signed-distance-transform channels are used for cross-modality alignment. No density channel contributed to transformation estimation for the spot-based ST analysis.

*Common raster domain and input-grid resolution.* For cross-sample alignment, the raster domain was constructed after global pre-alignment from the pooled query and reference coordinates. Let $x_{\min}, x_{\max}, y_{\min}$ and $y_{\max}$ denote the coordinate extrema across both datasets, let $(c_x, c_y)$ denote the center of this joint bounding box and let

$$L = \max\left(x_{\max} - x_{\min}, y_{\max} - y_{\min}\right).$$

A square domain centered at $(c_x, c_y)$, with side length $\eta L$, was used, where $\eta > 1$ is a margin-expansion factor. Given input-grid spacing $\Delta$, the grid coordinates were generated at intervals of $\Delta$ along both axes. The resulting number of grid locations along each axis was therefore approximately

$$N_x = N_y = \left\lceil \frac{\eta L}{\Delta} \right\rceil + 1,$$

up to endpoint rounding. Thus, the raster dimensions were determined by the spatial extent and grid spacing, rather than by a prescribed number of pixels or a fixed ratio to the number of cells or spots. Smaller values of $\Delta$ preserve finer spatial detail but increase memory and computation, whereas larger values produce a coarser and less expensive representation. The reported cross-sample workflows used $\eta = 1.05$, $\Delta = 30$ in the coordinate units supplied after any workflow-specific coordinate scaling and Gaussian smoothing with a standard deviation of one raster pixel.

*Shared-structure-composition channels.* For cross-sample spatial transcriptomics alignment, each jointly identified spatial structure that is present in both datasets defines one corresponding channel. The cells or spots assigned to a structure are first counted according to their positions on the pixel grid. The resulting count map is spatially smoothed so that nearby observations form a continuous field rather than isolated occupied pixels. This reduces sensitivity to irregular sampling and to small differences in cell or spot positions between sections.

At each pixel, the smoothed count for a structure is divided by the total smoothed count across all shared structures. Each resulting channel therefore represents the local proportion of observations assigned to one spatial structure. Corresponding structures occupy the same channel in the query and reference inputs. This local normalization reduces the influence of differences in sampling density while preserving the relative spatial organization of the shared structures. Structures absent from either dataset are excluded.

*Cell-density channel.* The structure-composition channels primarily describe the relative mixture of spatial structures and intentionally remove information about the total number of nearby cells. Consequently, regions with similar structure composition can have similar channel values even when one is densely populated and the other contains only weak cellular support. spAlignDE can therefore include an additional cell-density channel for single-cell cross-sample alignment when local cell density provides informative spatial support. No spot-density information contributed to transformation estimation in the reported spot-based Visium analysis because the limited number of spots did not support a sufficiently continuous density field.

The cell-density channel is calculated by summing the smoothed cell-count maps across all shared structures. The resulting density is log-transformed to limit the influence of unusually dense regions and normalized using the pooled query and reference cell-density distribution. Specifically, the pooled 99th percentile is used as the common scaling value, after which values are restricted to the range from zero to one. When enabled, the cell-density channel is appended to the structure-composition channels. Fixed weights control the relative contributions of spatial-structure composition and cell density during alignment.

*Paired signed-distance-transform channels.* For cross-modality alignment, each accepted query–reference structure pair defines one corresponding channel. The binary mask on each side of the pair is converted into a signed-distance-transform (SDT) field, which records the distance of every pixel from the structure boundary [14]. Values are positive inside the structure, zero at its boundary and negative outside. Compared with a binary mask, the SDT provides graded spatial information on both sides of the boundary and remains informative when the paired structures do not overlap perfectly after global pre-alignment.

Each SDT field is spatially smoothed to reduce small boundary irregularities. Distances beyond a specified range are truncated and the remaining values are normalized to the range from $-1$ to 1, preventing distant interior or background regions from dominating the alignment. Query and reference channels are arranged according to the same accepted-pair order. Optional pair-specific weights balance the contributions of structures with different spatial support. A whole-tissue support channel can additionally be appended when the overall tissue outline provides useful alignment information.

*Assembly and use in S-LDDMM.* For cross-sample alignment, the shared-structure-composition channels and, when enabled, the cell-density channel are concatenated to form the query and reference multichannel inputs. For cross-modality alignment, the corresponding SDT channels, together with any optional whole-tissue support channel, form the multichannel inputs. These fields are used only to estimate the S-LDDMM transformation and do not replace or aggregate the original observations. After optimization, the estimated transformation is applied directly to the original query cells, spots, image positions or atlas coordinates.

##### S2.7 Mathematical formulation and optimization of S-LDDMM

After global pre-alignment and construction of the multichannel structural inputs, spAlignDE estimates an affine–diffeomorphic transformation within the pre-aligned coordinate system. Let $\Omega \subset \mathbb{R}^2$ denote the common pixel domain and $C$ the number of structural channels. The fixed query and reference fields are functions

$$F^{\text{qry}}, F^{\text{ref}} : \Omega \to \mathbb{R}^C.$$

They have the same spatial dimensions and channel order, with each channel representing corresponding structural information. A spatial location is denoted by $\mathbf{x} \in \Omega$.

*Affine–diffeomorphic transformation.* The transformation estimated by S-LDDMM is

$$T_{\text{S-LDDMM}}(\mathbf{x}) = A(\phi_1(\mathbf{x})),$$

where $\phi_1$ is the endpoint of a diffeomorphic flow and $A$ is an affine map,

$$A(\mathbf{x}) = \mathbf{B}_A\mathbf{x} + \mathbf{b}_A.$$

Here, $\mathbf{B}_A \in \mathbb{R}^{2\times 2}$ is an invertible linear-transformation matrix and $\mathbf{b}_A \in \mathbb{R}^2$ is a shifting vector. The matrix $\mathbf{B}_A$ can represent rotation, scaling and shear, whereas $\mathbf{b}_A$ represents horizontal and vertical shifting.

The affine component is part of the S-LDDMM model and is distinct from the global pre-alignment $T_{\text{pre}}$. Because major differences in tissue position, orientation and size have already been addressed by global pre-alignment, the affine component is initialized as the identity transformation:

$$\mathbf{B}_A = I_2, \qquad \mathbf{b}_A = \mathbf{0},$$

where $I_2$ is the $2\times 2$ identity matrix. The affine component can subsequently correct broad spatial discrepancies that remain within the pre-aligned coordinate system.

*Geodesic shooting.* The diffeomorphic component is generated by a smooth time-dependent velocity field

$$\mathbf{v}_t : \mathbb{R}^2 \to \mathbb{R}^2, \qquad t \in [0,1],$$

through the flow equation [11, 15]

$$\frac{\partial}{\partial t}\phi_t(\mathbf{x}) = \mathbf{v}_t(\phi_t(\mathbf{x}))\,, \qquad \phi_0(\mathbf{x}) = \mathbf{x}.$$

Here, $\phi_t : \mathbb{R}^2 \to \mathbb{R}^2$ is the transformation at time $t$, $\phi_0$ is the identity transformation and $\phi_1$ is the final diffeomorphic deformation.

Under the shooting formulation, the complete trajectory is determined by an initial momentum field

$$\mathbf{m}_0 : \Omega_v \to \mathbb{R}^2,$$

where $\Omega_v \subset \mathbb{R}^2$ is the velocity-field domain. At time $t$, the momentum and velocity fields satisfy

$$\mathbf{m}_t = \mathcal{L}\mathbf{v}_t, \qquad \mathbf{v}_t = \mathcal{K}\mathbf{m}_t, \qquad \mathcal{K} = \mathcal{L}^{-1}.$$

The operator $\mathcal{L}$ defines the smoothness metric on the velocity field, whereas $\mathcal{K}$ converts momentum into a smooth velocity field.

In the implementation, $\mathcal{K}$ is applied in the Fourier domain using the discrete counterpart of

$$\mathcal{L} = \left(\mathrm{Id} - a^2\nabla^2\right)^{2p},$$

where Id is the identity operator, $\nabla^2$ is the spatial Laplacian, $a > 0$ is the smoothing scale and $p > 0$ is the operator power. Larger values of $a$ couple motion over a broader spatial range, whereas larger values of $p$ impose higher-order spatial smoothing.

The momentum evolves according to the geodesic equation

$$\frac{\partial \mathbf{m}_t}{\partial t} + (\nabla \mathbf{m}_t)\mathbf{v}_t + (\nabla \mathbf{v}_t)^{\mathsf{T}}\mathbf{m}_t + \mathbf{m}_t\,\nabla\cdot\mathbf{v}_t = \mathbf{0},$$

where $\nabla\mathbf{m}_t$ and $\nabla\mathbf{v}_t$ are the spatial Jacobian matrices and $\nabla\cdot\mathbf{v}_t$ is the divergence of the velocity field. Starting from $\mathbf{m}_0$, this equation is integrated forward together with the flow equation to obtain $\mathbf{m}_t$, $\mathbf{v}_t$ and $\phi_t$.

The velocity domain is constructed from the spatial extent of the query field. For spatial axis $d \in \{1,2\}$, let $l_d$ and $u_d$ denote the lower and upper coordinate limits, respectively. Its center and half-width are

$$c_d = \frac{l_d + u_d}{2}, \qquad r_d = \frac{u_d - l_d}{2}.$$

The velocity domain is

$$\Omega_v = \prod_{d=1}^{2} \left[c_d - \rho_v r_d,\, c_d + \rho_v r_d\right],$$

where $\rho_v \geq 1$ is the velocity-domain expansion factor. The expanded domain reduces boundary effects when the smoothing operator is applied using Fourier transforms.

The velocity field is represented on a regular grid with spacing $\Delta_{v,d} > 0$ along axis $d$. The trajectory is integrated using $n_t \in \mathbb{N}$ equal time steps of length

$$\Delta t = \frac{1}{n_t}.$$

Smaller velocity-grid spacing provides a more spatially detailed deformation field, whereas larger $n_t$ improves numerical integration accuracy at additional computational cost.

*Inverse warping of the query field.* Because $T_{\text{S-LDDMM}}$ maps query coordinates into the reference coordinate system, image matching is evaluated by inverse warping. For each $\mathbf{x} \in \Omega$, the warped query field is

$$\widetilde{F}^{\text{qry}}(\mathbf{x}) = F^{\text{qry}}\left[\phi_1^{-1}\left(A^{-1}(\mathbf{x})\right)\right].$$

Thus, each reference-grid location is mapped back through the inverse affine and diffeomorphic transformations to the corresponding position in the query field. The query-channel values at that position are obtained by spatial interpolation.

*Probabilistic multichannel matching with missing-data components.* Not every location in $\Omega$ provides reliable correspondence information. Non-overlapping tissue, missing regions, modality-specific structures and local acquisition artifacts may produce signals that should not strongly influence the estimated transformation. To accommodate these locations, spAlignDE adapts the probabilistic missing-data registration framework of Tward et al. [16] to multichannel structural fields, using an expectation–maximization mixture with one matched component and two unmatched appearance components.

Differences in channel amplitude and baseline intensity are accommodated by a linear channel mapping with an intercept,

$$\mathcal{G}(\mathbf{f}) = \mathbf{b}_G + \mathbf{B}_G\mathbf{f},$$

where $\mathcal{G} : \mathbb{R}^C \to \mathbb{R}^C$, $\mathbf{B}_G \in \mathbb{R}^{C\times C}$ is the channel-mapping matrix and $\mathbf{b}_G \in \mathbb{R}^C$ is the channel-specific intercept vector. This mapping changes only the channel values and does not alter spatial coordinates.

At each reference-grid location, the reference field is modeled using one matched component, denoted by $M$, and two unmatched appearance components, denoted by $U_1$ and $U_2$:

$$\begin{aligned} F^{\text{ref}}(\mathbf{x}) \sim\ & \pi_M \mathcal{N}\left[\mathcal{G}\left(\widetilde{F}^{\text{qry}}(\mathbf{x})\right), \sigma_M^2 I_C\right] \\ & + \pi_{U_1}\mathcal{N}\left(\boldsymbol{\mu}_{U_1}, \sigma_{U_1}^2 I_C\right) + \pi_{U_2}\mathcal{N}\left(\boldsymbol{\mu}_{U_2}, \sigma_{U_2}^2 I_C\right). \end{aligned}$$

Here, $\mathcal{N}(\boldsymbol{\mu}, \Sigma)$ denotes a $C$-dimensional Gaussian distribution with mean $\boldsymbol{\mu}$ and covariance matrix $\Sigma$, and $I_C$ is the $C \times C$ identity matrix. The vectors

$$\boldsymbol{\mu}_{U_1}, \boldsymbol{\mu}_{U_2} \in \mathbb{R}^C$$

represent the mean channel profiles of the unmatched components. The positive parameters

$$\sigma_M, \sigma_{U_1}, \sigma_{U_2} > 0$$

control the variability accommodated by the matched and unmatched components.

The mixture proportions satisfy

$$\pi_M, \pi_{U_1}, \pi_{U_2} \geq 0, \qquad \pi_M + \pi_{U_1} + \pi_{U_2} = 1.$$

The posterior probabilities of the three components define the spatial weights

$$W_M(\mathbf{x}),\ W_{U_1}(\mathbf{x}),\ W_{U_2}(\mathbf{x}) \in [0, 1],$$

which satisfy

$$W_M(\mathbf{x}) + W_{U_1}(\mathbf{x}) + W_{U_2}(\mathbf{x}) = 1$$

at every $\mathbf{x} \in \Omega$. Locations with consistent query–reference structural information receive a high matched-component weight $W_M(\mathbf{x})$, whereas locations representing missing tissue, weak overlap or modality-specific information can be assigned to an unmatched component.

Given the current matched-component weights, the channel mapping is estimated by weighted ridge regression:

$$\begin{aligned} (\widehat{\mathbf{b}}_G, \widehat{\mathbf{B}}_G) = \underset{\mathbf{b}_G, \mathbf{B}_G}{\operatorname{argmin}} & \sum_{\mathbf{x}\in\Omega} W_M(\mathbf{x}) \left\| \mathbf{b}_G + \mathbf{B}_G \widetilde{F}^{\text{qry}}(\mathbf{x}) - F^{\text{ref}}(\mathbf{x}) \right\|_2^2 \\ & + \lambda_G \left( \|\mathbf{B}_G\|_F^2 + \|\mathbf{b}_G\|_2^2 \right), \end{aligned}$$

where $\lambda_G \geq 0$ is the ridge-stabilization parameter, $\|\cdot\|_2$ is the Euclidean norm and $\|\cdot\|_F$ is the Frobenius norm.

The resulting multichannel matching term is

$$\mathcal{E}_{\text{match}} = \frac{1}{2\sigma_M^2} \sum_{\mathbf{x}\in\Omega} W_M(\mathbf{x}) \left\| \mathcal{G}\left(\widetilde{F}^{\text{qry}}(\mathbf{x})\right) - F^{\text{ref}}(\mathbf{x}) \right\|_2^2.$$

Locations supported by both datasets therefore contribute strongly to transformation estimation, whereas poorly matched locations are downweighted rather than forcing unsupported local deformation. Smaller values of $\sigma_M$ impose a stronger penalty for disagreement within the matched component.

*Deformation regularization.* The complete S-LDDMM objective balances multichannel structural agreement and deformation regularity:

$$\mathcal{E} = \mathcal{E}_{\text{match}} + \mathcal{E}_{\text{reg}}.$$

The deformation-regularization term is the LDDMM kinetic energy:

$$\mathcal{E}_{\text{reg}} = \frac{1}{2\sigma_R^2} \int_0^1 \langle \mathbf{m}_t, \mathbf{v}_t \rangle_{\Omega_v} \, dt,$$

where

$$\langle \mathbf{m}_t, \mathbf{v}_t \rangle_{\Omega_v} = \int_{\Omega_v} \mathbf{m}_t(\mathbf{x})^{\mathsf{T}} \mathbf{v}_t(\mathbf{x}) \, d\mathbf{x}.$$

The parameter $\sigma_R > 0$ controls the balance between multichannel matching and deformation regularity. Smaller values impose stronger regularization, whereas larger values permit greater deformation when supported by the matched structural fields.

*Alternating optimization.* The affine parameters $\mathbf{B}_A$ and $\mathbf{b}_A$, together with the initial momentum field $\mathbf{m}_0$, are estimated by gradient-based minimization of $\mathcal{E}$. The affine map is initialized using $\mathbf{B}_A = I_2$ and $\mathbf{b}_A = \mathbf{0}$, and the initial momentum is initialized as the zero vector field unless an estimate from a preceding alignment stage is supplied.

When enabled, an initial affine-only optimization stage corrects broad differences that remain after global pre-alignment. Optimization of $\mathbf{m}_0$ then begins, and the affine learning rate can be reduced so that subsequent local refinement is driven primarily by the diffeomorphic component. Momentum-gradient clipping and learning-rate decay can be used to stabilize optimization.

The channel mapping and spatial correspondence weights are updated alternately with the transformation parameters. Given the current transformation and $W_M$, $\mathcal{G}$ is refitted by weighted ridge regression. After an initial warm-up period, the mixture proportions, unmatched-component means and posterior weights are periodically updated by expectation–maximization. The current transformation therefore determines which locations provide consistent correspondence, and the resulting correspondence weights determine how strongly those locations influence subsequent transformation updates. The channel-mapping ridge constant was $\lambda_G = 0.1$. The matched and two unmatched spatial mixture-weight fields were initialized to 0.5, 0.4 and 0.1, respectively. Expectation–maximization updates were performed every five iterations beginning at iteration 50.

*Composition with global pre-alignment.* For an original query coordinate $\mathbf{q} \in \mathbb{R}^2$, the globally pre-aligned coordinate is

$$\mathbf{q}^{\text{pre}} = T_{\text{pre}}(\mathbf{q}).$$

The final aligned coordinate is

$$\mathbf{q}^{\text{final}} = T_{\text{S-LDDMM}}\left(\mathbf{q}^{\text{pre}}\right) = \left(A \circ \phi_1 \circ T_{\text{pre}}\right)(\mathbf{q}),$$

where $\circ$ denotes function composition. Therefore, the complete query-to-reference transformation is

$$T_{\text{final}} = T_{\text{S-LDDMM}} \circ T_{\text{pre}}.$$

The multichannel structural fields remain fixed during optimization and are used only to estimate this transformation. The complete transformation is finally applied to every original query observation, including cells, spots, image positions or atlas coordinates that were not directly used to construct the structural channels.

*Dataset-specific S-LDDMM settings for the reported alignments.* For cross-sample alignment, the mouse-brain S2R3-to-S2R2 fit used $n_t = 3$, $a = 300$, $p = 2$, domain-expansion factor 2, velocity-grid spacing 100, 500 optimization iterations and momentum learning rate $2 \times 10^3$. All 19 aging-brain source-to-4.3-month fits used the same settings except that optimization was run for 800 iterations. For the kidney IL3-to-NL3 alignment, coordinates were multiplied by 50 internally and S-LDDMM used $n_t = 5$, $a = 500$, $p = 2$, expansion factor 2, velocity-grid spacing 250, 5,000 iterations and momentum learning rate 50. The breast-cancer Replicate-2-to-Replicate-1 fit used $n_t = 3$, $a = 300$, $p = 2$, expansion factor 2, velocity-grid spacing 100, 500 iterations and momentum learning rate $4 \times 10^3$. All reported cross-sample structural fields used structure-composition weight 1. The cell-density channel weight was 1 for the mouse-brain and aging-brain cell-level analyses and 0 for the kidney and breast-cancer fits. All cross-sample S-LDDMM fits used `float32` precision. The breast-cancer fit additionally used momentum-gradient clipping at 1,000.

The reported Xenium-to-H&E alignment retained two matched structure pairs and used $n_t = 5$, $a = 60$, $p = 2$, expansion factor 2, velocity-grid spacing 6 and 300 optimization iterations. The affine-linear, affine-translation and momentum learning rates were $5 \times 10^{-9}$, $5 \times 10^{-2}$ and $2 \times 10^3$, respectively; the whole-tissue channel received a weight factor of 1.6, and optimization used `float64` precision. The reported MERFISH-to-Nissl alignment used three user-defined correspondence groups plus a whole-tissue channel. Equalized signed-distance inputs were clipped at 60 pixels, smoothed with $\sigma_{\text{SDT}} = 0.9$, normalized within a four-pixel boundary band and transformed using $\tanh(d/2)$. Area-based channel weights used exponent 0.8 and range 0.5–2.5, the whole-tissue channel received a factor of 1.2 and the structural fields were evaluated at 0.6 times the feature-grid resolution. S-LDDMM used $n_t = 5$, $a = 30$, $p = 2$, expansion factor 2, velocity-grid spacing 6 and 1,000 iterations. The affine-linear, affine-translation and momentum learning rates were $5 \times 10^{-9}$, $5 \times 10^{-2}$ and $2 \times 10^3$, respectively, and optimization used `float32` precision.

The reported ATAC-to-ST alignment retained eight matched structure pairs and used $n_t = 8$, $a = 100$, $p = 2$, expansion factor 2, velocity-grid spacing 50 and 500 optimization iterations, with diffeomorphic optimization beginning at iteration 20. The affine-linear, affine-translation and momentum learning rates were $2 \times 10^{-11}$, $2 \times 10^{-5}$ and $10^3$, respectively. Momentum-gradient clipping was 1.0, deformation regularization was $10^6$, the matching scale was 0.5 and optimization used `float64` precision.

#### S2.8 Stage-specific settings for coarse-to-fine ST and multiscale atlas alignment

*ST and atlas hierarchy levels.* The automatic ST-to-atlas analyses used three nested ST structure resolutions. The hierarchy was constructed by Ward clustering of the finest-level structure-average expression profiles after retaining genes that explained 80% of the cumulative expression variance, with at least 50 genes retained. For the primary S2R1 analysis, the three ST partitions contained 7, 16 and 25 structures, respectively. The corresponding partitions contained 7, 17 and 27 structures for S1R1 and 7, 14 and 21 structures for S3R1. The final partition in each analysis was the original refined BANKSY partition.

Only the ST resolution changed across the three hierarchical stages. At every stage, the atlas search space included all hierarchy prefixes of depth 2 or greater that were represented in the selected Allen CCFv3 slice; no stage-specific atlas-depth restriction was applied. The prefixes represented hierarchy depths 2–10 in all three slices. This produced 227 hierarchy-prefix candidates for the S2R1 reference slice ($z = 675$), 204 for the S1R1 slice ($z = 890$) and 145 for the S3R1 slice ($z = 485$). Six additional cortical-layer candidates, corresponding to layers 1, 2/3, 4, 5, 6a and 6b, were available in each slice.

*Candidate-mask construction.* At each stage, ST masks were reconstructed on the atlas raster from the currently transformed coordinates and the ST labels at the corresponding hierarchy level. Point densities were smoothed using an adaptive Gaussian bandwidth based on the fourth-nearest-neighbor distance. For normal structures, the bandwidth was scaled by 1.1 and restricted to 1.2–6.0 pixels; the normalized density was thresholded at 0.03, closed with a radius-10 disk, refined using closing and opening radii of 3 and 1, filled internally and filtered with a minimum connected-component size of 180 pixels. For structures classified as spatially detailed or thin, the bandwidth was scaled by 0.8 and restricted to 0.6–2.2 pixels; the density threshold was 0.11, the initial and refinement closing radii were both 1, no opening or hole filling was applied, and the minimum component size was 200 pixels.

For an atlas hierarchy prefix $p$, let $\mathcal{L}_p$ denote the set of descendant annotation identifiers present in the selected slice. Its candidate mask was

$$M_p(\mathbf{x}) = \mathbb{1}\{S(\mathbf{x}) \in \mathcal{L}_p\},$$

where $S(\mathbf{x})$ is the Allen annotation identifier at pixel $\mathbf{x}$. A cortical-layer mask was constructed analogously by taking the union of all slice labels whose anatomical names denoted the corresponding cortical layer. Atlas masks were generated directly from annotation-label unions and were not morphologically modified.

The same candidate-screening rules were used at all stages. A candidate was required to overlap the ST mask by at least 20 raster pixels. Hierarchy-prefix candidates were prescreened at Dice similarity 0.05, with the 30 highest-Dice candidates evaluated using the complete geometric score and at most 10 retained per ST structure. Cortical-layer candidates were prescreened at Dice similarity 0.03, with at most six retained per ST structure. Accepted pairs required a final gated score of at least 0.50 and an average symmetric surface distance no greater than 50 pixels. Layer correspondences were selected first; hierarchy-prefix pairs were then added greedily while preventing reuse of an ST structure or overlap between selected atlas-label sets.

*Stage-specific S-LDDMM settings.* The first two ST hierarchy levels used standard signed-distance-transform inputs. ST and atlas masks were preprocessed with Gaussian smoothing parameters 1.4 and 0.4, respectively; the corresponding binary thresholds were 0.5, closing radii were 2 and 1, opening radii were 1 and 1, and minimum mask areas were 50 and 80 pixels. Signed distances were clipped at 60 pixels and smoothed with $\sigma_{\mathrm{SDT}} = 1.2$. Channel weights used area exponent 0.9 and were restricted to 0.25–4.5. The structural fields were optimized at 0.3 times the atlas-raster resolution.

For these two stages, S-LDDMM used five time steps, kernel parameters $a = 500$ and $p = 2$, and a domain-expansion factor of 2. The first and second hierarchy stages were optimized for 100 and 500 iterations, respectively, with diffeomorphic optimization beginning at iteration 100. The learning rates were $2 \times 10^{-8}$, $2 \times 10^{-1}$ and $2 \times 10^3$ for the affine-linear, affine-translation and momentum parameters, respectively. The affine slowdown factor was 10, and the momentum learning rate was multiplied by 0.9995 with a lower bound of 200.

The final hierarchical stage and all subsequent final-resolution refinement cycles used equalized signed-distance inputs. Distances were clipped at 4 pixels, smoothed with $\sigma_{\mathrm{SDT}} = 0.9$, normalized within a four-pixel boundary band and transformed using $\tanh(d/2)$. Area-based channel weights used exponent 0.8 and were restricted to 0.5–2.5; the whole-tissue channel received an additional factor of 1.6. These fields were optimized at 0.6 times the atlas-raster resolution. S-LDDMM used five time steps, $a = 200$, $p = 2$, expansion factor 2 and grid step 50. The third hierarchy stage was optimized for 100 iterations, whereas each subsequent final-resolution continuation transformation was optimized for 200 iterations. Diffeomorphic optimization was active from the first iteration for the third stage and continuation transformations. All other learning-rate parameters were unchanged. At every stage, the matched and unmatched component weights were updated every five iterations beginning at iteration 50, using mixture scales

$$\sigma_M = 1, \qquad \sigma_{U_1} = 5, \qquad \sigma_{U_2} = 2.$$

The deformation-regularization scale was $\sigma_R = 5 \times 10^5$.

*UI-guided atlas alignment settings.* The reported UI-guided S2R1 atlas analysis used nine manually specified correspondence groups and automatic whole-tissue-mask pre-alignment. It used the same equalized signed-distance preprocessing, area-based channel weighting, whole-tissue channel factor and raster zoom as the automatic final-resolution stage. S-LDDMM used $n_t = 5$, $a = 200$, $p = 2$, expansion factor 2, velocity-grid spacing 50 and 500 optimization iterations, with diffeomorphic optimization active from the first iteration. The affine-linear, affine-translation and momentum learning rates were $2 \times 10^{-8}$, $2 \times 10^{-1}$ and $2 \times 10^3$, respectively. The affine slowdown factor was 10, and the momentum learning rate was multiplied by 0.9995 with a lower bound of 200. Matched and unmatched component weights were updated every five iterations beginning at iteration 50, using $\sigma_M = 1$, $\sigma_{U_1} = 5$ and $\sigma_{U_2} = 2$. The deformation-regularization scale was $\sigma_R = 5 \times 10^5$. Optimization used `float64` precision.

*Transformation composition and stopping rule.* Within stage $r$, the forward S-LDDMM mapping was

$$T_r = A_r \circ \phi_r,$$

so that the diffeomorphic flow was applied before the residual affine transformation. Each stage transformation was applied immediately to all ST coordinates, and the transformed coordinates were used to reconstruct masks and identify pairs at the next stage. If $C$ final-resolution refinement cycles were performed, the complete transformation was

$$T_{\text{total}} = T_C^{\text{cont}} \circ \cdots \circ T_1^{\text{cont}} \circ T_3 \circ T_2 \circ T_1 \circ T_{\text{pre}},$$

where the rightmost transformation was applied first.

A maximum of ten additional final-resolution refinement cycles was allowed. In each cycle, pairs were identified before S-LDDMM, one complete 200-iteration transformation was estimated and applied, and pairs were identified again. If $n_{\text{before}}$ and $n_{\text{after}}$ denote the accepted-pair counts before and after that transformation, iteration stopped when

$$n_{\text{after}} - n_{\text{before}} < 1.$$

Thus, the implementation tested the accepted-pair count rather than equality of the pair identities. It also stopped without estimating a transformation if no pairs were available at the beginning of a cycle.

For S2R1, the three hierarchy stages accepted 3, 8 and 16 pairs. Re-evaluation after the third-stage transformation yielded 17 pairs. The first continuation cycle increased the accepted-pair count from 17 to 18, whereas the second retained 18 pairs and triggered stopping, yielding 18 final pairs. For S1R1, the three hierarchy stages accepted 3, 7 and 11 pairs. Re-evaluation after the third-stage transformation yielded 12 pairs, and the subsequent continuation retained 12 pairs and triggered stopping, yielding 12 final pairs. For S3R1, the three hierarchy stages accepted 4, 5 and 10 pairs. The first continuation increased the accepted-pair count from 10 to 11, whereas the second retained 11 pairs and triggered stopping, yielding 11 final pairs.

##### S2.9 Comparator implementations

*General implementation protocol.* Cross-sample comparators were implemented using their published workflows and, where available, the parameterization used by the SABench framework [17]. In every pairwise comparison, the query was designated as the moving sample and the reference as the alignment target. Original spatial coordinates were used unless a method-specific initialization is stated below. GPU-enabled methods were run as separate processes on one NVIDIA RTX PRO 6000 Blackwell Max-Q GPU; CPU preprocessing was performed on the workstation described in Methods 5.5.2. CODA used the GPU for SuperPoint/LightGlue feature matching and one CPU thread for its NumPy/SciPy global and local coordinate optimization. Fixed iteration schedules or the native stopping criteria of each implementation were used. A run was retained only when it terminated normally and produced the expected number of finite aligned coordinates. Compatibility patches addressed software-interface, device-selection or numerical-precision issues without altering the corresponding alignment objective.

*PASTE.* PASTE 1.4.0 [18] was run using all shared input genes without external highly variable gene selection. Pairwise alignment used the default Kullback–Leibler expression dissimilarity, $\alpha = 0.1$, `norm=True` and the POT Torch backend on CUDA, followed by `stack_slices_pairwise` to place the query in the reference coordinate system. For the mouse-brain comparison, the supplied query pre-alignment coordinates were used as initialization. Because PASTE constructs dense pairwise-distance and transport matrices, optimization was restricted to a reproducible random subset of at most 30,000 observations per section using seed 0.

*PASTE2.* The PASTE2 partial-alignment objective [19] was evaluated through the maintained PASTE3 0.0.0 GPU backend. All shared genes were retained, and the final configuration used Kullback–Leibler dissimilarity, overlap fraction $s = 0.8$, $\alpha = 0.1$, normalized spatial costs, float32 transport matrices, 100 partial-EMD dummy points, an EMD iteration limit of $10^7$, and relative and absolute fused-Gromov–Wasserstein tolerances of $10^{-6}$. The optimization used at most 30,000 observations per section. Partial stacking produced the aligned fitting coordinates. For the aging-brain analysis, the affine representation of the resulting stacking transformation was recovered from the fitting subset and applied to all query cells, while the reference coordinates were retained unchanged.

*STAligner.* STAligner 1.0.0 [20] was run on the complete sections. For each section, a radius-based spatial graph was constructed with cutoff 150, up to 5,000 highly variable genes were selected using the `seurat_v3` procedure, and expression was normalized to 10,000 counts per observation and log transformed. The concatenated sections were trained with `knn_neigh=100`, 600 epochs and pairwise order `iter_comb=[(1,0)]` on CUDA. Louvain domains were identified from the STAligner embedding at resolution 0.2 using Scanpy random state 666. All detected domains were supplied to the ICP step, and the resulting ICP transformation was applied to the complete query section.

*GPSA.* GPSA 0.6 [21] was run using at most 10,000 observations per section. Counts were normalized by library size, log transformed and reduced to at most 3,000 highly variable genes. Spatial coordinates were independently rescaled to $[0, 10]^2$, and expression features were standardized within each view. The reference was assigned to view 0 and fixed using `fixed_view_idx=0`. Variational GPSA used an identity-fixed mean function, radial-basis-function warp and data kernels, data-based initialization, five Monte Carlo samples, 50 inducing locations for both the warp and data Gaussian processes, and Adam optimization with learning rate $10^{-2}$ for 3,000 epochs. The posterior common-coordinate means were retained as the aligned coordinates. All GPSA alignments included in the reported analyses completed successfully under this fixed configuration.

*SLAT.* scSLAT 0.3.0 [22] was run on the complete sections. The two sections were joined using their shared genes, normalized by total counts, log transformed and scaled separately. The package's DPCA representation was constructed in 50 dimensions; because the analyzed datasets contained fewer genes than the internal 12,000-gene cap, all available shared genes were retained. A 10-nearest-neighbor spatial graph was constructed for each section. SLAT was trained for six epochs with one LGCN layer, after which feature-based spatial matching was performed with `reorder=False`. A two-dimensional rigid rotation and translation were estimated by singular-value decomposition between query coordinates and their matched reference coordinates and applied to every query observation.

*STalign.* STalign 1.0 [23] used spatial point-density images and therefore required neither expression normalization nor gene selection. Query and reference coordinates were rasterized using the package defaults, followed by `LDDMM` with `niter=10000` and `epV=50` on CUDA. For the mouse-brain comparison, the same supplied query pre-alignment coordinates used by the benchmark wrapper were provided before rasterization. The estimated source-to-target point transformation was subsequently evaluated at every query coordinate.

*CAST.* CAST 0.4 [24] was run on the complete sections. Expression was normalized to 10,000 counts per observation without additional log transformation or highly variable gene selection. CAST-MARK used Delaunay spatial graphs and 400 training epochs. CAST-STACK used 150 affine iterations, `dist_penalty1=0`, `bleeding=500`, $d = [3, 2, 1, 0.5, 1/3]$, and affine basis weights $[10^{-3}, 10^{-3}, 1/50, 5, 5]$. The spot-level kidney analysis included one B-spline refinement iteration, whereas the larger cell-level datasets used no additional B-spline iteration. CAST-MARK and CAST-STACK were executed on CUDA, and the returned coordinates were retained for all observations.

*STAIR.* STAIR-tools 1.3.1 [25] was run on the complete sections. Its internal preprocessing normalized total expression and applied $\log(1 + x)$, without highly variable gene selection or feature scaling. The negative-binomial autoencoder used 128 hidden units, a 32-dimensional latent representation, dropout 0.2, learning rate $10^{-3}$, batch size 128 and 100 training epochs. Homogeneous spatial graphs used ten neighbors, and the heterogeneous graph-attention model used `c_neigh_het=0.9`, $\gamma = 0.8$ and 150 epochs. The production runs used the prespecified 13-cluster $k$-means fallback when the requested R-based mclust step was unavailable. Location alignment used one mutual-nearest-neighbor match, one spatial domain, $\alpha = 500$, at most 20 fine-alignment iterations and tolerance $10^{-10}$. The resulting fine-alignment coordinates were retained for all observations.

*Spateo.* Spateo-release 1.1.1 [26] was run on the complete sections. Counts were normalized by total expression, log transformed and annotated with up to 2,000 highly variable genes. Non-rigid `morpho_align` was performed in `SN-N` mode with stochastic variational inference enabled, precomputed distances and a maximum of 200 iterations on CUDA. The query coordinates stored in `align_spatial_nonrigid` were used as the final aligned coordinates.

*SANTO.* SANTO 0.0.2 [27] was run as a global rigid-alignment method. Expression was normalized to 10,000 counts per observation and log transformed. For the MERFISH mouse-brain data, 166 `Blank-*` control probes were excluded and the remaining 483 biological genes were used; all 300 genes were used for the aging-brain sections, and the 300 genes with the largest normalized variance were used for the kidney sections. Large datasets were represented during fitting by up to 4,000 spatially balanced cells per section. SANTO was run in fine-only mode with `mode=None`, 40 epochs, $k = 20$, $\alpha = 0.1$, learning rate $10^{-3}$, two spatial dimensions and `diff_omics=False`. The fitting subset was used only to estimate the rigid rotation and translation. The resulting transformation matrix was then applied to every query cell, while every reference cell was retained unchanged; thus, the reported SANTO outputs contain the complete sections rather than only the fitting cells.

*SPOmiAlign.* SPOmiAlign 0.1.0 [28] was run on the complete sections using the released compatibility wrapper. No highly variable gene selection was applied. Each H5AD was rendered as a grayscale scatter image using total expression across its genes, a long-side resolution of 2,200 pixels, square points with radius 5, target rotation 0°, query rotation −90°, reference blur $\sigma = 6.0$ and SSIM smoothing $\sigma = 1.5$. The `affine+bspline` workflow used RoMa matching on CUDA, estimated the affine and B-spline transformations in image space and back-projected the combined transformation to every original query coordinate.

*CODA.* CODA 2.0.0 [29] was run on the complete sections using the Tutorial 1 full-alignment workflow. Counts were normalized to 10,000 per observation, log transformed and restricted to at most 3,000 highly variable genes. The sections were integrated using BBKNN with 30 principal components, followed by a three-dimensional UMAP representation and Louvain clustering at resolution 2. The resulting shared Louvain labels were used for G1 rigid alignment with 10% trimming. The mouse-brain comparison used the same supplied query pre-alignment coordinates as the STalign benchmark, whereas the kidney and aging-brain comparisons used their original coordinates. Embedding images were generated on automatically selected canvases, and SuperPoint/LightGlue common-domain matching used at most 2,048 keypoints on CUDA. Local LDDMM used $T = 32$, $K = 20$, $\sigma = 0.2$, $\alpha = 1$, $\gamma = 1$, $\epsilon = 0.001$ and 20 iterations. The local transformation was back-projected to all query coordinates and retained only when nearest-neighbor shared-label consistency improved by at least 0.02; otherwise, the G1 rigid coordinates were used. This quality-control fallback was treated as part of the fixed CODA workflow rather than as a failed run.

*xIV-LDDMM.* The atlas comparison used xIV-LDDMM-Particle 1.0.0 [30, 31]. xIV-LDDMM received the same pre-aligned ST coordinates used to initialize the corresponding spAlignDE analysis; no spAlignDE structures, matched channels or aligned coordinates were supplied. Blank probes were excluded, and the remaining 483 gene-count features were transformed using $\log(1 + x)$. A seed-0 random subset of 5,000 ST cells was used as source control particles. The Allen atlas slice was downsampled by a factor of eight to 9,359 target particles represented by one-hot indicators for 153 atlas regions. The RKHS kernel scales were $[0.20, 0.10, 0.05]$, the varifold scales were $[0.20, 0.10, 0.05, 0.02]$, and $\gamma = 0.1$, $c_A = 1$, $c_T = 1$ and $c_S = 10$. Optimization was allowed up to 300 steps on CUDA and terminated after 47 steps according to the implementation's native stopping behavior. The estimated shooting transformation was subsequently evaluated for all 83,546 ST cells before atlas labels were assigned using the common atlas-coordinate lookup.

*3d-OT.* The spatial ATAC-to-ST comparison used 3d-OT 0.1.1 [32]. Both methods were initialized in the same manually pre-aligned ATAC–ST coordinate frame. The analysis retained 476 shared biological genes after excluding blank probes and used ST expression and ATAC-derived gene-activity counts. The two modalities were jointly normalized by total counts, log transformed, scaled within modality and represented by a 50-dimensional DPCA embedding; the internal 12,000-gene highly variable gene cap therefore retained all 476 shared genes. Spatial graphs used six nearest neighbors. Modality-specific graph encoders were trained for 800 epochs using Adam with learning rate $10^{-3}$. The unified alignment model used `simk=5`, `otk=300`, `reconk=2`, smooth- and divergence-flow regularization, and one alignment epoch with learning rate $10^{-4}$. All 35,422 ST reference cells and all 9,215 ATAC query cells were used, and the reconstructed ATAC coordinates were mapped back to the shared ST coordinate system.

*Comparator selection for the post-alignment simulation benchmark.* The final simulation benchmark evaluated ten methods: spAlignDE, CAST, STAligner, SLAT, STalign, STAIR, Spateo, SANTO, SPOmiAlign and CODA. Each synthetic sample contained 62,296 observations, and local false-discovery proportion and power were evaluated on the same complete set of sample-B locations for every method. Full-data PASTE and PASTE2 runs were computationally infeasible because their implementations construct dense pairwise-distance and transport matrices, whereas full-data GPSA exceeded feasible memory because of its Gaussian-process covariance and variational computations. Feasible runs of these three methods required reducing each synthetic sample to 10,000 observations, and their benchmark wrappers did not return a validated out-of-sample transformation for the remaining observations. Including their subsampled outputs would therefore have changed the testing units and truth denominator, whereas method-specific interpolation would have introduced an additional unvalidated postprocessing procedure. PASTE, PASTE2 and GPSA were consequently excluded from the ten-method post-alignment inference benchmark, although they were retained in the separate biological alignment benchmarks.

*Randomization and run validation.* Cross-sample comparator wrappers initialized NumPy and PyTorch with random seed 0, except SANTO, which used seed 42; STAligner additionally used a Scanpy random state of 666. xIV-LDDMM used seeds 0 and 1 for spatially stratified subsampling of the ST and atlas data, respectively, whereas 3d-OT used seed 7. Runs that raised exceptions, exhausted available memory or returned missing or non-finite coordinates were classified as failures. No automatic retry procedure was used; after diagnosis, failed runs were restarted from the beginning using the same locked configuration.

#### S2.10 Detailed evaluation metrics for spatial alignment

Alignment performance was evaluated using molecular, annotation-based, anatomical and geometric criteria selected for each alignment task. Whenever possible, evaluation used biological information that was not supplied to the alignment procedure, thereby avoiding assessment based on the same spatial structures used to estimate the transformation. The following sections describe the construction of the evaluation inputs, the mathematical definitions of metrics requiring explicit specification, and the procedures used to summarize performance across genes, anatomical classes and dataset pairs.

##### S2.10.1 Metrics for cross-sample alignment

*Shared-grid gene-expression concordance.* For cross-sample spatial transcriptomics alignment, molecular concordance was evaluated by comparing the post-alignment spatial distributions of shared genes. The overlapping rectangular extent of the aligned query and reference sections was divided into corresponding spatial bins. An $m \times n$ grid divides the overlapping coordinate range into $m$ equal-width intervals along one spatial axis and $n$ equal-width intervals along the other, producing $mn$ rectangular bins before filtering.

For each grid resolution, the observation-count threshold was calculated as one-half of the smaller of the query and reference mean numbers of observations across all bins. A bin was retained only when both datasets met this threshold. This filtering reduced instability caused by bins containing very few cells or spots.

For each retained bin $b = 1, \ldots, B$ and shared gene $g = 1, \ldots, G$, mean expression was calculated separately in the query and reference datasets, producing the spatial profiles

$$\mathbf{e}_g^{\text{qry}} = (e_{g1}^{\text{qry}}, \ldots, e_{gB}^{\text{qry}}), \qquad \mathbf{e}_g^{\text{ref}} = (e_{g1}^{\text{ref}}, \ldots, e_{gB}^{\text{ref}}).$$

Here, $B$ is the number of bins retained in both datasets, $G$ is the number of evaluated genes, $e_{gb}^z$ is the mean expression of gene $g$ in bin $b$, and $z \in \{\text{qry}, \text{ref}\}$ denotes the aligned query or fixed reference dataset.

Following the spatial-alignment benchmarking framework of Yan et al. [17], agreement between the two profiles was quantified independently for each gene using the Pearson correlation coefficient (PCC), cosine similarity, structural similarity index measure (SSIM) and mutual information (MI).

PCC measures linear agreement between the centered profiles:

$$\text{PCC}_g = \frac{\sum_{b=1}^{B}(e_{gb}^{\text{qry}} - \bar{e}_g^{\text{qry}})(e_{gb}^{\text{ref}} - \bar{e}_g^{\text{ref}})}{\sqrt{\sum_{b=1}^{B}(e_{gb}^{\text{qry}} - \bar{e}_g^{\text{qry}})^2}\sqrt{\sum_{b=1}^{B}(e_{gb}^{\text{ref}} - \bar{e}_g^{\text{ref}})^2}},$$

where

$$\bar{e}_g^z = \frac{1}{B}\sum_{b=1}^{B} e_{gb}^z$$

is the mean expression of gene $g$ across retained bins in dataset $z$. PCC was set to zero when either profile had zero variance.

Cosine similarity measures agreement in the direction of the two non-centered profiles:

$$\text{Cosine}_g = \frac{\mathbf{e}_g^{\text{qry}} \cdot \mathbf{e}_g^{\text{ref}}}{\|\mathbf{e}_g^{\text{qry}}\|_2 \|\mathbf{e}_g^{\text{ref}}\|_2},$$

where $\|\cdot\|_2$ denotes the Euclidean norm. Cosine similarity was set to zero when either profile had zero norm.

For SSIM, each spatial profile was divided by its maximum value:

$$\widetilde{e}_{gb}^z = \frac{e_{gb}^z}{\max_{b'} e_{gb'}^z}.$$

A profile with a maximum value of zero was represented by a vector of zeros. Let

$$\mu_z = \frac{1}{B}\sum_{b=1}^{B} \widetilde{e}_{gb}^z$$

and

$$\sigma_z = \sqrt{\frac{1}{B}\sum_{b=1}^{B}(\widetilde{e}_{gb}^{z} - \mu_z)^2}$$

denote the population mean and standard deviation of the normalized profile. The population covariance between the query and reference profiles was

$$\sigma_{\text{qry,ref}} = \frac{1}{B}\sum_{b=1}^{B}(\widetilde{e}_{gb}^{\text{qry}} - \mu_{\text{qry}})(\widetilde{e}_{gb}^{\text{ref}} - \mu_{\text{ref}}).$$

SSIM was then calculated as

$$\text{SSIM}_g = \frac{2\mu_{\text{qry}}\mu_{\text{ref}} + C_1}{\mu_{\text{qry}}^2 + \mu_{\text{ref}}^2 + C_1}\,\frac{2\sigma_{\text{qry}}\sigma_{\text{ref}} + C_2}{\sigma_{\text{qry}}^2 + \sigma_{\text{ref}}^2 + C_2}\,\frac{\sigma_{\text{qry,ref}} + C_3}{\sigma_{\text{qry}}\sigma_{\text{ref}} + C_3},$$

with $C_1 = 0.01^2$, $C_2 = 0.03^2$ and $C_3 = C_2/2$. These constants stabilize the calculation when the profile means or variances are small. The three factors measure agreement in average intensity, contrast and spatial variation, respectively.

MI measures statistical dependence between the query and reference bin values and can capture relationships not restricted to linear association. MI was estimated using the nearest-neighbor regression estimator implemented in `sklearn.feature_selection.r`[illegible] with the query profile as the predictor, the reference profile as the response, $k = 3$ nearest neighbors and random seed 0. MI was set to zero when either profile was constant or fewer than two retained bins were available.

Higher PCC, cosine similarity, SSIM and MI values indicate stronger post-alignment spatial concordance. The primary paired-section and aging-cohort mouse-brain evaluations used $m = n = 10$; both analyses were additionally repeated using $m = n = 30$ and $m = n = 50$ to evaluate finer spatial patterns and assess sensitivity to grid resolution. For the aging-brain cohort, each metric was first calculated separately for every query-to-reference alignment and was then averaged across the available alignments for each gene.

*Annotation consistency after spatial label transfer.* When compatible biological annotations were available, we evaluated whether aligned query observations were located near reference observations with consistent labels. Let $(\mathbf{x}_i^{\text{qry}}, y_i^{\text{qry}})$ denote the aligned coordinate and original annotation of query observation $i$, and let $(\mathbf{x}_j^{\text{ref}}, y_j^{\text{ref}})$ denote the coordinate and annotation of reference observation $j$. Euclidean distance was used throughout the annotation-transfer analysis.

The nearest reference observation was

$$j^*(i) = \arg\min_j \|\mathbf{x}_i^{\text{qry}} - \mathbf{x}_j^{\text{ref}}\|_2,$$

and its label was transferred to the query observation:

$$\widehat{y}_i^{\text{qry}} = y_{j^*(i)}^{\text{ref}}.$$

For each annotation class $c$, Dice similarity and intersection over union (IoU) were calculated as

$$\text{Dice}_c = \frac{2TP_c}{2TP_c + FP_c + FN_c}, \qquad \text{IoU}_c = \frac{TP_c}{TP_c + FP_c + FN_c}.$$

Here, $TP_c$ is the number of query observations for which both the original and transferred labels equal $c$, $FP_c$ is the number transferred to $c$ whose original label differs from $c$, and $FN_c$ is the number originally annotated as $c$ but transferred to another class.

Let $n_c$ be the number of query observations originally assigned to class $c$. Class-specific scores were summarized as

$$\text{Weighted Dice} = \frac{\sum_c n_c \text{Dice}_c}{\sum_c n_c}, \qquad \text{Weighted IoU} = \frac{\sum_c n_c \text{IoU}_c}{\sum_c n_c}.$$

Local annotation consistency was additionally evaluated over a reference neighborhood rather than using only the nearest observation:

$$\text{kNN same-label} = \frac{1}{N_{\text{qry}}k}\sum_{i=1}^{N_{\text{qry}}}\sum_{j\in\mathcal{N}_k^{\text{ref}}(i)} \mathbf{1}(y_j^{\text{ref}} = y_i^{\text{qry}}).$$

Here, $N_{\text{qry}}$ is the number of query observations, $\mathcal{N}_k^{\text{ref}}(i)$ is the set of the $k$ nearest reference observations to aligned query observation $i$, and $\mathbf{1}(\cdot)$ is the indicator function. We used $k = 5$ for the kidney benchmark and $k = 20$ for the aging-brain benchmark. Aging-brain scores were calculated separately for each query section and then averaged across alignments.

*Computational performance.* Computational performance was measured for the core alignment procedure, excluding method-specific preprocessing. Running time was recorded as the wall-clock duration from the start to the completion of the core alignment and was reported in minutes.

For CUDA-based methods, the GPU allocator was reset before alignment and the largest amount of memory occupied by allocated GPU tensors during the procedure was recorded in gigabytes. This measurement did not include unused memory retained in the reserved CUDA cache. Methods for which no CUDA allocation was recorded were omitted from the GPU-memory comparison. Dataset subsampling and method-specific input-size restrictions are reported with the corresponding benchmarks.

Shared input loading, preprocessing or subsampling performed outside the published alignment routine, output serialization and plotting were excluded. Operations intrinsic to a method, including spatial-graph construction, rasterization and learned-feature estimation, were included. End-to-end runtime was recorded separately and was not used in the comparative analysis.

**S2.10.2 Metrics for cross-modality alignment** Because different modalities do not generally contain directly comparable measurements, cross-modality alignment was evaluated using task-specific molecular, anatomical and spatial readouts that were independent of the structural fields used to estimate the transformation.

*Common-grid molecular concordance.* For histology-guided and spatial ATAC-to-ST alignment, molecular measurements from the aligned datasets were aggregated into corresponding square spatial bins. Let $c^z_{gb}$ denote the summed raw expression count or gene-activity value for gene $g$, bin $b$ and modality $z$. Within each bin, the molecular profile was library-size normalized and log-transformed:

$$x^z_{gb} = \log\left(1 + 10^4 \frac{c^z_{gb}}{\sum_{h=1}^{G} c^z_{hb}}\right),$$

where $G$ is the number of evaluated genes and the denominator is the total molecular signal across these genes in bin $b$. A bin with zero total signal was assigned zero for all genes. Only bins satisfying the specified observation-count threshold in both modalities were retained.

PCC and cosine similarity were calculated from the vectors of retained bin values. A gene was excluded from PCC when either profile had a standard deviation no greater than $10^{-12}$, and from cosine similarity when the product of the two Euclidean norms was no greater than $10^{-12}$.

For two-dimensional SSIM, the retained bin values were placed at their corresponding positions in a rectangular two-dimensional array. Positions not retained in both modalities were set to zero. The SSIM data range was defined as the difference between the largest and smallest values across the two arrays. Genes with a data range no greater than $10^{-12}$ were excluded from the SSIM summary.

For MI, the retained values in each modality were independently divided using 20 equally spaced quantiles between 0 and 1. Repeated quantile boundaries were removed before discretization. MI was calculated from the joint frequency table of the resulting intervals and was set to zero when either profile was constant or could not be divided into at least two intervals. This quantile-based estimator differs from the nearest-neighbor estimator used for cross-sample alignment.

*Reliability-weighted gene-level summaries.* Cross-modality similarity estimates may be unstable for genes with weak detection or little spatial variation. Each gene was therefore assigned a reliability weight based on its detection support, spatial variability and detection balance between modalities.

Let $f^z_g$ be the fraction of retained bins with a positive raw molecular value for gene $g$ in modality $z$, and let $v^z_g$ be the population variance of its log-normalized profile across retained bins. For modalities $z_1$ and $z_2$, we defined

$$d_g = \sqrt{f^{z_1}_g f^{z_2}_g}, \qquad s_g = \sqrt{v^{z_1}_g v^{z_2}_g},$$

and

$$b_g = \frac{\min(f^{z_1}_g, f^{z_2}_g) + \epsilon}{\max(f^{z_1}_g, f^{z_2}_g) + \epsilon}, \qquad \epsilon = 10^{-9}.$$

Here, $d_g$ measures shared detection support, $s_g$ measures shared spatial variability, and $b_g$ penalizes strongly imbalanced detection between modalities.

For any gene-level quantity $a_g$, $R(a_g)$ denotes its percentile rank among the $G$ evaluated genes, calculated as its average rank in the presence of ties divided by $G$. The unnormalized and normalized reliability weights were

$$u_g = R(d_g)R(s_g)b_g, \qquad w_g = \frac{u_g}{\sum_{h=1}^{G} u_h}.$$

If all $u_g$ values were zero or non-finite, equal weights $w_g = 1/G$ were used.

For a gene-level metric $m_g$, the weighted summary was calculated over genes with finite metric values:

$$\overline{m}_{\text{weighted}} = \frac{\sum_{g \in \mathcal{G}_{\text{valid}}} w_g m_g}{\sum_{g \in \mathcal{G}_{\text{valid}}} w_g},$$

where $\mathcal{G}_{\text{valid}}$ is the set of genes with finite values for the evaluated metric. Reliability weights were calculated separately for each alignment result.

*Gene-expression concordance after ST-to-histology alignment.* spAlignDE estimated the transformation using spatial structures derived from Xenium gene expression and the reference histological image. The Visium gene-expression measurements paired with this image were withheld from transformation estimation and used only for evaluation. After alignment, we assessed molecular concordance by comparing spatial gene-expression patterns between the Xenium and Visium datasets.

Both datasets were represented in the shared histology-image coordinate system. Square bins had a side length of approximately 11.0 pixels in this coordinate system. A bin was retained when it contained at least three Xenium cells and at least one Visium spot. PCC, cosine similarity, two-dimensional SSIM and quantile-discretized MI were calculated for the 246 shared genes. Overall performance was summarized using both unweighted and reliability-weighted gene-level means.

*Atlas-label coverage.* For ST-to-atlas alignment, the transformed coordinate of each ST observation was sampled from the reference atlas annotation image to obtain a transferred anatomical label. Atlas label 0 represented background or unlabeled atlas space. Atlas-label coverage was defined as the percentage of all evaluated ST observations assigned a nonzero atlas label. Conversely, the background-assignment rate was the percentage assigned label 0. Background assignments were also summarized separately by RCTD-inferred class label to identify populations preferentially mapped outside the annotated atlas domain.

*Label agreement between RCTD-inferred class labels and transferred atlas labels.* ST observations were independently annotated using RCTD [33] with the Allen Institute Mouse Whole Cortex and Hippocampus 10x single-cell RNA-sequencing dataset as the reference [34, 35]. A reproducible subset of 1,000 reference cells was selected using random seed 1. The ST and reference count matrices were restricted to their shared genes, RCTD was run in doublet mode, and the primary `first_type` assignment was retained as the expression-derived cell-type label.

After alignment, Allen CCF labels transferred to the ST observations were collapsed into coarse anatomical categories. A predefined high-confidence mapping specified the atlas categories considered compatible with each evaluated RCTD label. The mappings were: `L2_3_IT_CTX` to `CTX_L2_3`; `L4_5_IT_CTX` to `CTX_L4` or `CTX_L5`; `L5_IT_CTX` and `L5_PT_CTX` to `CTX_L5`; `L6_IT_CTX` and `L6_CT_CTX` to `CTX_L6a`; `L5_6_NP_CTX` to `CTX_L5` or `CTX_L6a`; `L6b_CTX` to `CTX_L6b`; `CA1_ProS` to `CA1`; `CA3` to `CA3`; and `DG` to `DG`. For presentation, `L5_IT_CTX` and `L5_PT_CTX` were combined as `L5_CTX`, whereas `L6_IT_CTX` and `L6_CT_CTX` were combined as `L6_CTX`.

An ST observation was considered anatomically consistent when its transferred atlas category belonged to the expected category set for its RCTD-inferred class label. Broadly distributed glial, vascular and inhibitory subclasses without a unique anatomical expectation were excluded from this strict analysis.

We reported the overall label-agreement rate across all eligible observations, the class-specific label-agreement rate and a balanced label-agreement rate obtained by averaging category-specific rates with equal weight. The balanced rate prevents abundant RCTD categories from dominating the comparison. Atlas-background assignments were summarized for all RCTD labels, including labels excluded from strict anatomical matching.

*Atlas marker-gene localization.* For each regional marker gene $g$, an expected atlas-category set $E_g$ was specified before evaluation. The primary expected sets were cortex and hippocampus for *Slc17a7*, cortical layers 6a and 6b for *Ntsr1*, striatum for *Drd1* and *Adora2a*, thalamus and hypothalamus for *Slc17a6*, habenula for *Gpr151*, and reticular thalamic nucleus for *Cckar*.

Let $x_{ig}$ denote the raw expression count of marker gene $g$ in ST observation $i$. Marker-high observations, denoted by $\mathcal{Q}_g$, were defined as observations with positive expression at or above the 75th percentile of the positive-count distribution for that gene. The same gene-specific threshold was used for the primary automatic alignment, the UI-guided alignment and the structure-pair sensitivity analysis.

Let $C_i$ denote the transferred atlas category of observation $i$, and let $n_{C_i}$ be the number of observations assigned to that category. Each observation received the inverse-frequency weight

$$q_i = \frac{1}{n_{C_i}}.$$

Category sizes and weights were calculated separately for each alignment method. This weighting prevents large transferred atlas regions from dominating the evaluation.

Region-balanced marker enrichment was calculated as

$$\text{Enrichment}_g = \frac{\dfrac{\sum_{i\in\mathcal{Q}_g} q_i \mathbf{1}(C_i \in E_g)}{\sum_{i\in\mathcal{Q}_g} q_i}}{\dfrac{\sum_i q_i \mathbf{1}(C_i \in E_g)}{\sum_i q_i}}.$$

The numerator is the weighted fraction of marker-high observations assigned to expected regions, whereas the denominator is the weighted fraction of all observations assigned to those regions. Values greater than one indicate preferential localization of marker-high observations within the expected anatomical regions.

We additionally evaluated marker localization using a region-balanced AUROC and the contrast in weighted mean expression between expected and non-expected regions. Let

$$z_{ig} = \log(1 + x_{ig})$$

be the log-transformed expression of gene $g$ in observation $i$. For any atlas-category set $S$, the weighted mean expression was

$$\bar{z}_{g,S} = \frac{\sum_i q_i z_{ig} \mathbf{1}(C_i \in S)}{\sum_i q_i \mathbf{1}(C_i \in S)}.$$

The expected-to-non-expected expression contrast was

$$R_g = \log_2\left(\frac{\bar{z}_{g,E_g} + 0.01}{\bar{z}_{g,E_g^c} + 0.01}\right),$$

where $E_g^c$ includes all non-expected assignments, including atlas background, and 0.01 is a pseudocount preventing division by zero.

For the region-balanced AUROC, membership in $E_g$ was treated as the positive class, $z_{ig}$ was used as the prediction score and $q_i$ was used as the observation weight. Marker-enrichment variability was evaluated using 250 bootstrap samples of the marker-high observations, sampled with replacement using random seed 7 while holding the region-frequency weights fixed.

*ATAC–ST anatomical annotation consistency.* Spatial ATAC-seq and ST annotations were first harmonized into a shared set of coarse anatomical categories. Euclidean distance in the shared evaluation coordinate system was used to identify the ten nearest ST observations to each aligned ATAC pixel.

An ST neighbor at distance $d$ received weight $1/(d+10^{-8})$. Weights were summed separately for each anatomical category and divided by the total weight across all ten neighbors. The category with the largest normalized weight was transferred to the ATAC pixel.

The exact-match rate was the fraction of ATAC pixels for which the transferred ST category equaled the original ATAC annotation. Annotation support was the normalized voting weight assigned to the original ATAC category, averaged across ATAC pixels. The median Euclidean distance from each aligned ATAC pixel to its nearest ST observation was retained as a spatial-proximity diagnostic.

*Preservation of the native ATAC neighborhood structure.* To determine whether alignment distorted the internal organization of the ATAC dataset, local ATAC neighborhoods were compared before and after alignment. The same set of ATAC pixels was used in both coordinate systems, and each pixel itself was excluded from its neighborhood.

Let $\mathcal{N}_k^{\mathrm{orig}}(i)$ and $\mathcal{N}_k^{\mathrm{aln}}(i)$ denote the sets of the $k$ nearest ATAC neighbors of pixel $i$ in the original and aligned coordinate systems, respectively. Neighborhood preservation for pixel $i$ was

$$P_i(k) = \frac{\left|\mathcal{N}_k^{\mathrm{orig}}(i) \cap \mathcal{N}_k^{\mathrm{aln}}(i)\right|}{k}.$$

A value of one indicates that all original neighbors were retained after alignment, whereas a value of zero indicates that none were retained. Mean and median values of $P_i(k)$ were reported across ATAC pixels for $k \in \{10, 20, 30, 50\}$. This metric measures deformation within the query dataset and does not by itself quantify ATAC-to-ST correspondence.

*ATAC–ST gene-level spatial concordance.* ATAC-derived gene-activity scores and ST gene expression were compared for their shared genes using the common-bin molecular-concordance procedure. Square bins had a side length of 25 units in the shared ATAC–ST evaluation coordinate system. A bin was retained when it contained at least three ATAC spatial pixels and at least three ST observations.

Within each retained bin, ATAC gene-activity and ST expression profiles were independently normalized to a total of $10^4$ and transformed using $\log(1 + x)$. PCC, cosine similarity, two-dimensional SSIM and quantile-discretized MI were then calculated independently for each shared gene. Reliability weights were estimated separately for each alignment method using the corresponding ATAC and ST detection fractions and spatial variances. All methods were evaluated using the same ATAC pixels, ST observations, shared-gene set and spatial-bin configuration.

#### S2.11 Subsampling-based transformation stability

To quantify the local stability of the cross-sample deformation estimated by spAlignDE, we performed a repeated subsampling analysis using the MERFISH Mouse Brain Receptor Map data obtained from Vizgen. We used the same query-to-reference direction as in the primary cross-sample benchmark, with section S2R3 as the query and section S2R2 as the reference. This analysis measured the mean Euclidean displacement of each fixed query location from its replicate-mean aligned position when the complete alignment pipeline was re-estimated using independently subsampled but biologically comparable inputs.

We generated $R = 10$ replicate datasets by independently sampling, without replacement, 80% of the cells from each section using fixed replicate- and sample-specific random seeds. For every replicate, we recomputed the complete cross-sample workflow. This comprised per-sample BANKSY representation learning, principal-component and Harmony integration, shared-nearest-neighbor Leiden clustering at resolution 1.4, boundary-aware structure refinement, weighted shared-structure-centroid pre-alignment that estimates rotation, translation and scaling without reflection, rasterization into multichannel structure-composition and cell-density fields, and S-LDDMM alignment. All non-sampling parameters were held fixed. Each replicate therefore yielded an independently estimated final S-LDDMM transformation $T_r$, $r = 1, \dots, R$, reflecting variation propagated through structure construction, pre-alignment and non-rigid alignment.

For replicate $r \in \{1, \dots, 10\}$, subsampling used seed $20260401 + 1000r + \delta_s$, where $\delta_{\mathrm{S2R2}} = 17$ and $\delta_{\mathrm{S2R3}} = 31$. Each replicate used clustering seed 1000. The replicate-specific S-LDDMM fits used $n_t = 3$, $a = 300$, $p = 2$, velocity-grid spacing 100 and 500 iterations. The affine-linear, affine-translation and momentum learning rates were $2 \times 10^{-8}$, 0.2 and $2 \times 10^3$, respectively; the affine slowdown factor was 10, momentum-gradient clipping was disabled and the momentum learning rate was held at $2 \times 10^3$.

Directly comparing the output coordinates within each replicate would confound transformation variability with differences in subsample membership. We therefore fixed the evaluation support to the pre-aligned query points from replicate 1. Let $p_i \in \mathbb{R}^2$ denote the pre-aligned coordinate of fixed query point $i$, $i = 1, \dots, n$. Each replicate-specific transformation $T_r$ was applied to the same point $p_i$, yielding the aligned coordinate

$$z_{r,i} = T_r(p_i) = (x_{r,i}, y_{r,i}) \in \mathbb{R}^2.$$

The repeat-mean aligned coordinate for point $i$ was defined as

$$\bar{z}_i = \frac{1}{R} \sum_{r=1}^{R} z_{r,i}.$$

We then computed the replicate-wise displacement from the mean aligned position,

$$d_{r,i} = \|z_{r,i} - \bar{z}_i\|_2 .$$

Coordinate-wise variance was calculated as

$$s_{x,i}^2 = \frac{1}{R-1} \sum_{r=1}^{R} (x_{r,i} - \bar{x}_i)^2, \qquad s_{y,i}^2 = \frac{1}{R-1} \sum_{r=1}^{R} (y_{r,i} - \bar{y}_i)^2,$$

where $\bar{z}_i = (\bar{x}_i, \bar{y}_i)$. We summarized the total positional spread by

$$s_i = \sqrt{s_{x,i}^2 + s_{y,i}^2}.$$

The primary pointwise transformation-stability measure was the mean Euclidean displacement from the repeat-mean aligned position,

$$\bar{d}_i = \frac{1}{R} \sum_{r=1}^{R} d_{r,i}.$$

For secondary characterization of replicate-to-replicate dispersion in displacement magnitude, we also calculated the sample variance and standard deviation,

$$v_{d,i} = \frac{1}{R-1} \sum_{r=1}^{R} \left(d_{r,i} - \bar{d}_i\right)^2, \qquad s_{d,i} = \sqrt{v_{d,i}}.$$

Here, $\bar{d}_i$ measures the average magnitude of the displacement of point $i$ from its repeat-mean aligned position. The coordinate-spread measure $s_i$ and the displacement-variance measure $v_{d,i}$ provide secondary summaries of transformation variability. The spatial stability map in Fig. 2E displays the replicate-1 aligned query coordinates colored by $\bar{d}_i$, corresponding to the `dist_mean` output. Locations above the 95th percentile were outlined.

This procedure produced a pointwise empirical stability measure without requiring a ground-truth deformation field. Low values of $\bar{d}_i$ indicate that independently subsampled alignments placed the same query location close to its repeat-mean aligned position, whereas high values indicate that the estimated local transformation was sensitive to the sampled tissue support. Most locations showed small mean Euclidean displacement, whereas the largest values were concentrated near the partially missing or weakly overlapping boundary. The measure should therefore be interpreted as subsampling-based transformation stability rather than as a calibrated probability or confidence interval for the unknown true deformation.

An additional aging-brain stability analysis aligned the 26.7-month section to the 4.3-month reference. Ten independent 80% subsamples used seeds $267043+r$, $r=1,\ldots,10$, and each replicate used clustering seed 1000. BANKSY representations, joint PCA, Harmony integration, Leiden clustering at resolution 1.4 and boundary-aware refinement were recomputed for every replicate. The same fixed manual similarity initialization was applied to all replicates, after which S-LDDMM used $n_t=3$, $a=300$, $p=2$, velocity-grid spacing 100, 500 iterations and momentum learning rate $2\times10^3$.

##### S2.12 Robustness to spatial-structure choices

The robustness analyses examined two structural choices that may vary across datasets or parameter settings: the resolution of jointly inferred structures in cross-sample alignment and the number of accepted structure pairs in cross-modality alignment. In each analysis, one structural factor was varied while the input observations, evaluation features and all unrelated alignment parameters were held fixed.

*Cross-sample sensitivity to spatial-structure resolution.* Sensitivity to the granularity of jointly inferred spatial structures was evaluated using the MERFISH mouse-brain benchmark, with section S2R3 as the query and section S2R2 as the reference. The Leiden resolution was varied over 0.6, 0.8, 1.0, 1.2 and 1.4, producing 15, 17, 21, 24 and 27 boundary-refined shared structures, respectively.

The BANKSY features, principal-component representation, Harmony-adjusted embedding and shared-nearest-neighbor graph were held fixed across all conditions. For each resolution, the Leiden labels and all subsequent structure-dependent steps were recomputed. These steps included boundary-aware structure refinement, shared-structure-centroid pre-alignment, construction of the structure-composition and cell-density channels, and S-LDDMM alignment. Rasterization settings, channel weights and S-LDDMM parameters were held fixed. The completed alignment at the primary resolution of 1.4 was retained as the baseline, whereas the complete structure-dependent workflow was rerun for resolutions 0.6–1.2.

All resolution settings were evaluated using the same 483 shared genes and the same 65 retained bins from the common $10\times10$ spatial grid. PCC, cosine similarity, SSIM and MI were calculated using the cross-sample gene-expression concordance procedure described in Supplementary Methods S2.10.1. Because genes were the evaluated features rather than independent biological replicates, metric distributions were treated as descriptive summaries, and no formal hypothesis test was performed between resolutions.

Query–reference tissue overlays were additionally compared using the same spatial display limits. Aligned query and reference structure maps were visualized separately for each resolution. Structure colors were matched between the query and reference within a resolution but not across resolutions because the inferred spatial partitions differed. Gene-pattern similarity and whole-tissue overlays are shown in Supplementary Fig. S5, and the corresponding spatial-structure maps are shown in Supplementary Fig. S6.

*Cross-modality sensitivity to the number of retained structure pairs.* Sensitivity to the amount of cross-modality structural guidance was evaluated using the primary automatic ST-to-Allen-CCF alignment. The completed fixed-seed S2R1 pipeline retained 18 accepted ST–atlas structure pairs. This completed multistage alignment remained the primary Atlas result and was used only to define the fixed pair set for the sensitivity analysis. The 18 pairs were ranked once by the final gated alignment score, with the ungated alignment score used to resolve remaining ties. The controlled sensitivity conditions retained the highest-ranking 18, 16, 14, 12, 10 or 8 pairs.

The ST dataset, Allen CCFv3 reference section, global pre-alignment and all S-LDDMM parameters were held fixed. For every pair-count condition, including the 18-pair condition, signed-distance-transform channels were reconstructed from the corresponding subset of the locked 18-pair set, and one final-resolution S-LDDMM alignment was performed for 200 optimization iterations from the same pre-aligned coordinates. Structure-pair discovery was not repeated. Thus, the one-step 18-pair rerun served as the controlled full-pair baseline, and the fitting protocol differed across conditions only in the number of retained structure pairs. After each alignment, Allen CCFv3 labels were sampled at the transformed ST coordinates using the same atlas section with $10\,\mu$m pixel spacing. Label agreement was evaluated using the same RCTD-inferred class labels, predefined RCTD-to-atlas compatibility mapping and eligible ST observations used in the primary atlas evaluation. Overall, balanced and class-specific label-agreement rates were calculated as described in Supplementary Methods S2.10.2. These comparisons are shown in Supplementary Fig. S22.

Marker-gene localization was additionally evaluated for *Slc17a7*, *Ntsr1*, *Drd1*, *Slc17a6*, *Adora2a*, *Gpr151* and *Cckar*. For each gene, marker-high ST observations were defined as observations with positive raw expression at or above the 75th percentile of the positive-count distribution. Their transferred atlas labels were classified according to whether they belonged to the predefined expected anatomical regions described in Supplementary Methods S2.10.2. The same gene-specific expression threshold, expected-region definition and spatial display extent were used across all pair-count conditions. Marker-localization comparisons are shown in Supplementary Fig. S23.

### S3 Detailed post-alignment inference

#### S3.1 Shared-grid construction, bandwidth selection and support thresholds

For aligned sample $s$, let $\widetilde{\mathbf{q}}_j^{(s)}$ denote the aligned coordinate of the $j$th observed spot or cell, and let $X_{jg}^{(s)}$ be its expression of gene $g$. Separately, let $\boldsymbol{\xi}_i$, $i = 1, \ldots, N_{\text{grid}}$, denote the constructed shared-grid locations within the common aligned tissue support. Each retained $\boldsymbol{\xi}_i$ is a testing unit, whereas the nearby observations at $\widetilde{\mathbf{q}}_j^{(s)}$ provide the local data used at that testing unit. Regions outside the aligned support, holes, or locations with insufficient nearby observations are excluded from testing. Around each grid location, the local test defines a Gaussian-weighted neighborhood with a hard distance cutoff,

$$w_{ij}^{(s)} = \exp\left\{-\frac{\|\widetilde{\mathbf{q}}_j^{(s)} - \boldsymbol{\xi}_i\|^2}{2h_{\text{loc}}^2}\right\} \mathbf{1}\left\{\|\widetilde{\mathbf{q}}_j^{(s)} - \boldsymbol{\xi}_i\| \le h_{\text{loc}}\right\},$$

where $h_{\text{loc}}$ sets the Gaussian distance scale and also truncates observations outside the local neighborhood. These weights define the local estimation neighborhoods across aligned samples; they support the grid-level tests but are not themselves testing units. The same grid is used for all genes in a comparison, so differences across genes arise from expression rather than from changes in the spatial testing domain.

To express the amount of independent information represented by unequal kernel weights, define the normalized weights

$$a_{ij}^{(s)} = \frac{w_{ij}^{(s)}}{\sum_j w_{ij}^{(s)}}.$$

Under a local working approximation in which the observations are independent with common variance $\sigma^2$,

$$\operatorname{Var}\left(\sum_j a_{ij}^{(s)} X_j^{(s)}\right) = \sigma^2 \sum_j \left(a_{ij}^{(s)}\right)^2 = \frac{\sigma^2}{n_{s,i}^{\text{eff}}}.$$

Equating these two expressions gives the Kish effective local sample size [36],

$$n_{s,i}^{\text{eff}} = \frac{1}{\sum_j \left(a_{ij}^{(s)}\right)^2} = \frac{\left(\sum_j w_{ij}^{(s)}\right)^2}{\sum_j (w_{ij}^{(s)})^2}.$$

When the weights are equal, $n_{s,i}^{\text{eff}}$ equals the number of contributing observations; when the weights are concentrated on a few observations, it is smaller. Grid locations with insufficient effective support in any compared sample are removed from the gene-level local testing set $\mathcal{I}_g$. This filtering prevents unstable local tests in regions where the kernel neighborhood is too sparse to reliably estimate a local mean and variance.

The effective spatial resolution of the local test is therefore determined by observation density relative to the neighborhood bandwidth, rather than by shared-grid density alone. Denser spot or cell sampling can provide sufficient effective support within a smaller physical neighborhood and thereby resolve finer spatial changes. Under sparse sampling, stable local estimation necessarily borrows information over a broader area, which can smooth narrow or sharply bounded expression changes. Increasing the number of shared-grid testing locations alone does not replace missing observations or increase the underlying local information.

For sample $s$, define its coordinate scale as

$$L_s = \max\left\{\text{range}(x_s), \text{range}(y_s)\right\}.$$

Let $d_{sj}^{(5)}$ denote the fifth-nearest-neighbor distance for observation $j$. The local observation density and target neighborhood radius are estimated as

$$\widehat{\rho}_s = \text{median}_j \frac{5}{\pi\left(d_{sj}^{(5)}\right)^2}, \qquad R_s = \left(\frac{18}{\pi\widehat{\rho}_s}\right)^{1/2}.$$

The relative bandwidth is

$$R_{\text{rel}} = \text{clip}\left[Q_{0.65}\left(\frac{R_s}{L_s}\right), 0.010, 0.020\right].$$

With $L = \text{median}_s L_s$, the local neighborhood bandwidth is $h_{\text{loc}} = R_{\text{rel}} L$.

For each analysis, a tissue mask was constructed from the aligned sample geometry, and shared support retained locations inside the common occupied tissue after excluding holes and unsupported boundary locations. Let $N_{\text{typ}}$ denote the median number of observations per sample. The implementation first evaluates the R-driven candidate $n_{\text{raw}} = \text{round}(4.5/R_{\text{rel}})$, constructs its masked shared grid and counts the valid locations $m_{\text{grid}}$. The R-driven resolution is retained when

$$N_{\text{typ}} \le m_{\text{grid}} \le 2N_{\text{typ}}.$$

Otherwise, an integer search selects the grid resolution whose masked valid-location count is closest to the violated boundary. A user-specified grid resolution overrides this automatic rule. The local bandwidth is $h_{\text{loc}} = R_{\text{rel}}L$, where $L$ is the coordinate scale used by the geometry rule. In the real-data analyses, the minimum effective support was the empirical 10th percentile of positive local Kish effective sample sizes, rounded down and clipped to $[2, 25]$. The risk-map smoothing radius was $R_{\text{map}} = 1.5h_{\text{grid}}$, where $h_{\text{grid}}$ is the final grid spacing.

#### S3.2 Analysis-specific settings for post-alignment inference

The settings used in the two real-data analyses are summarized below; simulation settings are specified in Supplementary Methods S3.10 and S3.10.1.

| Setting | Aging-brain analysis | Injured-kidney analysis |
|---|---|---|
| Valid shared-grid locations | 77,056 | 6,187 |
| Grid spacing | 27.05 | 75.49 |
| $h_{\text{loc}}$ | 121.1 | 191.9 |
| $R_{\text{map}}$ | 40.57 | 113.2 |
| Minimum effective support | Empirical rule, clipped to $[2, 25]$ | Empirical rule, clipped to $[2, 25]$ |
| Comparison-wide intercept | Included | Omitted |
| Confounding covariates | Library size; detection rate | Library size; detection rate |
| Mismatch adjustment | On ($s = 0.25$) | On ($s = 0.75$) |
| Cell-type adjustment | Off; on only in Fig. 5C | Off |
| Local FDR threshold | 0.05; 0.10 in Fig. 5C | 0.05 |

Putatively stable-gene selection used the default screen in Supplementary Methods S3.3. Gene-level Aggregated Cauchy Association Test weights are given in Supplementary Methods S3.9.

#### S3.3 Putatively stable-gene selection and mismatch-risk construction

*Putatively stable-gene selection.* The alignment mismatch risk score reflects the putative level of local mismatch; a high score means that the compared neighborhoods remain unreliable after alignment. It is estimated from a data-selected panel of putatively stable genes so that the score primarily reflects local alignment or sampling reliability rather than the candidate DE signal. The putatively stable-gene set and local risk map are constructed separately for each pairwise or sequential contrast. To keep the single-contrast construction readable, the contrast index $c$ is suppressed below; in a multi-sample analysis the resulting local score is denoted by $R^{\text{loc}}_{ic}$. Let $\mathcal{G}_0$ denote a set of putatively stable genes selected to have sufficient spatial variation, positive cross-sample local correspondence, and small global expression difference between the compared samples. These genes are used as internal proxies for local comparability after alignment and are not assumed to be known negative controls, although the use of empirically identified stably expressed genes as negative-control features has precedent in the estimation of shared unwanted variation [37].

For putatively stable-gene screening, define

$$\widehat{\ell}_{s,ig} = \log(1 + \widehat{\mu}_{s,ig}), \qquad \bar{\widehat{\ell}}_{s,g} = |\mathcal{I}|^{-1} \sum_{i \in \mathcal{I}} \widehat{\ell}_{s,ig}.$$

Putatively stable genes are selected according to

$$\mathcal{G}_0 = \left\{ g : \begin{array}{l} \max\Big[\text{Var}_i(\widehat{\ell}_{\text{A},ig}), \text{Var}_i(\widehat{\ell}_{\text{B},ig})\Big] \ge \tau_{\text{var}}, \\ |\bar{\widehat{\ell}}_{\text{A},g} - \bar{\widehat{\ell}}_{\text{B},g}| \le \tau_{\Delta}, \\ \text{Corr}_i\left(\widehat{\ell}_{\text{A},ig}, \widehat{\ell}_{\text{B},ig}\right) \ge \tau_{\rho} \end{array} \right\}.$$

The current implementation sets $\tau_{\text{var}}$ to the 50th percentile of the maximum cross-grid log-mean variance across the two samples and sets $\tau_{\rho} = 0.20$. Among genes passing these variation and correlation criteria, $\tau_{\Delta}$ is the larger of 0.05 and the 10th percentile of the absolute comparison-wide log-mean differences. At most 300 putatively stable genes are retained; if more genes pass, they are ranked by the product of their non-negative cross-sample spatial correlation and spatial variance. These criteria retain genes with informative spatial variation and concordant spatial structure but without a large comparison-wide expression difference. The resulting map is a contrast-level diagnostic constructed once from the pooled stable-gene profiles and density channel and reused for every target-gene fit in that contrast. The implementation does not perform leave-one-gene-out risk construction: if a tested gene belongs to $\mathcal{G}_0$, its three profile channels remain among the channels used to form the common risk map. Because the map pools the full retained panel and the density channel, this contribution does not make the risk map gene specific.

*Alignment-mismatch risk construction.* For each putatively stable gene $g \in \mathcal{G}_0$, sample $s$, and grid location $i$, let $\widehat{\mu}_{s,ig}$ and $\widehat{v}_{s,ig}$ denote the kernel-weighted mean and variance estimates within the neighborhood used to construct the risk profile. We summarize the fitted local expression distribution using its mean, negative-binomial dispersion, and excess-zero probability. Under the negative-binomial mean–variance relation, the local negative-binomial size is estimated by moment matching as

$$\widehat{r}_{s,ig} = \frac{\widehat{\mu}_{s,ig}^2}{\max\{\widehat{v}_{s,ig} - \widehat{\mu}_{s,ig}, \epsilon_r\}}.$$

Smaller values of $\widehat{r}_{s,ig}$ indicate greater fitted local overdispersion, whereas large values approach the Poisson limit.

The expected zero probability under the fitted negative-binomial distribution is

$$\widehat{q}_{s,ig}^{\mathrm{NB}} = \left(1 + \frac{\widehat{\mu}_{s,ig}}{\widehat{r}_{s,ig}}\right)^{-\widehat{r}_{s,ig}}.$$

Let $p_{s,ig}^0$ denote the kernel-weighted observed zero fraction. The excess-zero probability is estimated as

$$\widehat{\pi}_{s,ig} = \min\left[1 - \epsilon_\pi, \max\left\{0, \frac{p_{s,ig}^0 - \widehat{q}_{s,ig}^{\mathrm{NB}}}{\max\left(1 - \widehat{q}_{s,ig}^{\mathrm{NB}}, \epsilon_\pi\right)}\right\}\right].$$

The three-channel local profile is

$$\widehat{\mathbf{x}}_{s,ig} = \left(\log(1 + \widehat{\mu}_{s,ig}), \log(1 + \widehat{r}_{s,ig}), \operatorname{logit} \widehat{\pi}_{s,ig}\right).$$

The three channels describe local expression magnitude, overdispersion and excess sparsity, respectively. The vectors are stacked over putatively stable genes to form $\widehat{\mathbf{x}}_{s,i}$.

Before the local profiles are compared, each channel is standardized across valid grid locations within its own sample. For a generic channel $u_{s,i\ell}$, define

$$\mathcal{Z}_s(u_{s,i\ell}) = \frac{u_{s,i\ell} - \overline{u}_{s,\ell}}{s_{s,\ell} + \epsilon_Z},$$

where $\overline{u}_{s,\ell}$ and $s_{s,\ell}$ are the within-sample mean and standard deviation of channel $\ell$. This prevents high-scale channels from dominating the cosine risk score.

In addition to stable-gene concordance, corresponding local neighborhoods are expected to have similar observation density. Let $d_{s,i}$ denote the local density of sample $s$,

$$d_{s,i} = \log\left\{1 + \sum_j \exp\left[-\frac{\|\widetilde{\mathbf{q}}_j^{(s)} - \boldsymbol{\xi}_i\|^2}{2h_{\mathrm{loc}}^2}\right] \mathbf{1}\left\{\|\widetilde{\mathbf{q}}_j^{(s)} - \boldsymbol{\xi}_i\| \le h_{\mathrm{loc}}\right\}\right\}.$$

With $d_{\mathrm{gene}} = 3|\mathcal{G}_0|$ standardized stable-gene channels and target density-energy share $s \in (0, 1)$, set

$$\eta = \left(\frac{s}{1-s} d_{\mathrm{gene}}\right)^{1/2}.$$

The augmented local profile is

$$\widetilde{\mathbf{x}}_{s,i} = \left[\mathcal{Z}_s(\widehat{\mathbf{x}}_{s,i}), \eta \mathcal{Z}_s(d_{s,i})\right].$$

The raw local alignment-mismatch score is

$$M_i = 1 - \frac{\widetilde{\mathbf{x}}_{\mathrm{A},i}^\top \widetilde{\mathbf{x}}_{\mathrm{B},i}}{\|\widetilde{\mathbf{x}}_{\mathrm{A},i}\| \, \|\widetilde{\mathbf{x}}_{\mathrm{B},i}\|}.$$

The raw risk map is smoothed over neighboring grid locations,

$$\widetilde{M}_i = \frac{\sum_{i'} G_\tau(\|\boldsymbol{\xi}_i - \boldsymbol{\xi}_{i'}\|) M_{i'}}{\sum_{i'} G_\tau(\|\boldsymbol{\xi}_i - \boldsymbol{\xi}_{i'}\|)},$$

where $G_\tau$ is a spatial smoothing kernel on the grid. The local risk map is median–MAD standardized, floored at zero, capped at the 95th percentile among valid grid locations, and rescaled to $[0, 1]$,

$$\zeta_i^{\mathrm{R}} = \max\left[0, \frac{\widetilde{M}_i - \mathrm{median}_{j\in\mathcal{I}}(\widetilde{M}_j)}{\mathrm{MAD}_{j\in\mathcal{I}}(\widetilde{M}_j) + \epsilon_R}\right],$$

$$R_i^{\mathrm{loc}} = \min\left\{\frac{\zeta_i^{\mathrm{R}}}{Q_{0.95}(\zeta_j^{\mathrm{R}} : j \in \mathcal{I}) + \epsilon_R}, 1\right\},$$

where $Q_{0.95}$ is computed over valid grid locations. High risk indicates that putatively stable-gene structure and/or local observation density remains inconsistent after alignment. The score distinguishes relative reliability across grid locations within a contrast and is not interpreted as a gene-specific biological effect or as a direct estimate of the true alignment error.

For visualization, Fig. 4C and Supplementary Figs. S27 and S28 show the non-negative robust-standardized score $\zeta_i^{\mathrm{R}}$ after median–MAD standardization and flooring at zero, but before 95th-percentile capping and rescaling. Consequently, the displayed values have no fixed upper bound. Only the rescaled score $R_i^{\mathrm{loc}} \in [0, 1]$ enters the variance-inflation model.

##### S3.4 Empirical calibration of mismatch inflation

For each gene $g$, the local model is first fitted with mismatch inflation disabled. Each contrast $c$ supplies initial local statistics $t_{igc}^{(0)}$ and its corresponding risk map $R_{ic}^{\mathrm{loc}}$. An exact-zero-risk bin is retained when it contains sufficient locations, and positive-risk locations are divided into quantile bins. The default calibration requests ten bins and retains a bin only when it contains at least 200 locations. Let $B_{bc}$ denote a retained bin, with median risk $r_{bc}$ and size $n_{bc}$.

Within risk bin $b$, define

$$m_{bgc} = \mathrm{median}_{i\in B_{bc}}\, t_{igc}^{(0)}$$

and

$$\mathrm{MAD}_{bgc} = \mathrm{median}_{i\in B_{bc}} \left| t_{igc}^{(0)} - m_{bgc} \right|.$$

For residual degrees of freedom $\nu_{gc}$, the MAD of the corresponding standard Student-$t$ null distribution is

$$q_{0.75,gc} = F_{t_{\nu_{gc}}}^{-1}(0.75).$$

The robust scale relative to this null and its non-negative excess variance are

$$\widehat{s}_{bgc} = \frac{\mathrm{MAD}_{bgc}}{q_{0.75,gc}}, \qquad y_{bgc} = \left[\widehat{s}_{bgc}^{2} - 1\right]_{+}.$$

For a retained zero-risk bin, the target excess is fixed at zero, ensuring that a location with normalized local risk zero receives no mismatch inflation.

The raw values $y_{bgc}$ are constrained to not decrease with risk by weighted isotonic regression,

$$\widetilde{\mathbf{y}}_{gc} = \arg \min_{z_1 \le \cdots \le z_B} \sum_b n_{bc}(z_b - y_{bgc})^2.$$

With $d_{\mathrm{risk}} = 2$, a non-negative quadratic relation through the origin is fitted,

$$\widehat{B}_{gc} = \arg\min_{B\ge 0} \sum_b n_{bc} \left(\widetilde{y}_{bgc} - B r_{bc}^2\right)^2.$$

The square reflects the variance scale of a first-order spatial perturbation: if mismatch risk tracks the magnitude of local displacement, the induced expression error is first order in that displacement and its additional variance is proportional to its squared magnitude.

Let $b_{\star c}$ be the retained bin whose median risk is nearest to $Q_{0.80}(r_{bc})$. The bounded reference-bin factor is

$$\widehat{\tau}_{gc} = \left[\frac{\widetilde{y}_{b_{\star c}gc}}{\widehat{B}_{gc} r_{b_{\star c}c}^2}\right]_{\mathrm{clip}(0,1)}.$$

If the denominator is zero or non-finite, $\widehat{\tau}_{gc}$ is set to zero. The provisional contrast-specific coefficient is

$$\widetilde{\lambda}_{gc} = \min\left\{\widehat{\tau}_{gc}\widehat{B}_{gc}, \lambda_{\max}\right\},$$

with $\lambda_{\max} = 5 \times 10^4$.

A contrast-specific calibration is considered valid only when calibration completes successfully, at least 800 usable grid locations and at least four retained risk bins are available, the retained bins provide at least four distinct finite risk values including positive risk, and the fitted quantities are finite and within their stated bounds. A successful estimate equal to zero remains valid evidence; a zero produced only as a failed-calibration fallback is excluded.

Let $\mathcal{C}_g^{\text{valid}}$ denote the valid contrasts. For multiple valid contrasts, define

$$m_g = \text{median}_{c\in\mathcal{C}_g^{\text{valid}}} \widetilde{\lambda}_{gc},$$

$$s_g = \max\left\{1.4826 \ \text{median}_{c\in\mathcal{C}_g^{\text{valid}}} \left|\widetilde{\lambda}_{gc} - m_g\right|, 10^{-8}(1+|m_g|)\right\}.$$

The shared gene-specific coefficient is the equal-weight Huber location

$$\widehat{\lambda}_g = \arg\min_{\lambda\geq 0} \sum_{c\in\mathcal{C}_g^{\text{valid}}} \rho_{1.345}\left(\frac{\widetilde{\lambda}_{gc} - \lambda}{s_g}\right),$$

where

$$\rho_\kappa(u) = \begin{cases} u^2/2, & |u| \leq \kappa, \\ \kappa|u| - \kappa^2/2, & |u| > \kappa. \end{cases}$$

The result is truncated to $[0, \lambda_{\max}]$. With one valid contrast, $\widehat{\lambda}_g$ equals its provisional coefficient exactly; if no contrast is valid, $\widehat{\lambda}_g = 0$. The final contrast-specific variance factor is

$$\phi_{igc}^{\text{align}} = 1 + \widehat{\lambda}_g (R_{ic}^{\text{loc}})^2.$$

The calibration treats increases in the robust dispersion of median-centered initial local statistics across risk strata as mismatch-associated excess variability. This interpretation assumes that the risk–dispersion relationship is not driven predominantly by true spatially heterogeneous biological signal. The estimated risk maps and $\widehat{\lambda}_g$ are treated as fixed plug-in quantities; their estimation uncertainty is not propagated separately into the final local reference distribution.

##### S3.5 Cell-type-composition inconsistency map

For sample $s$, contrast $c$, grid location $i$ and cell type $\ell$, kernel smoothing with symmetric Dirichlet regularization estimates the local composition,

$$\widehat{p}_{ic\ell}^{(s)} = \frac{\sum_j w_{ij}^{(s)} a_{j\ell}^{(s)} + \alpha_0}{\sum_j w_{ij}^{(s)} + \alpha_0 L},$$

where $a_{j\ell}^{(s)}$ is a one-hot cell-type indicator for annotated single-cell data or an estimated cell-type proportion for deconvolved spot-level data. Here, $L$ is the number of retained cell types and the default regularization parameter is $\alpha_0 = 1$. The regularized vector sums to one and remains well defined when individual cell types are locally absent.

The target and reference composition vectors enter Eq. (5.2.4). The displayed inconsistency map is $D_{ic}^{\text{cell}}$. Its normalization by $\log 2$ places the map on a common zero-to-one scale across contrasts. The exponential variance link gives a factor between one and $e$, providing a bounded and smoothly increasing adjustment. Missing cell-type information or a disabled cell-type adjustment sets $\phi_{ic}^{\text{cell}} = 1$.

##### S3.6 Confounding-covariate basis

For sample $s$, let $L_{is}$ and $D_{is}$ denote the grid-level local library-size and detection-rate summaries, and define

$$\mathbf{q}_{is} = [\log(1 + L_{is}), D_{is}]^\top.$$

A control set $\mathcal{S}_{\text{ctrl}}$ consists of the designated sample-B controls. The pooled entries of $\mathbf{q}_{is}$ over $i$ and $s \in \mathcal{S}_{\text{ctrl}}$ define the feature-wise control mean $\boldsymbol{\mu}_{\text{ctrl}}$ and standard deviation $\boldsymbol{\sigma}_{\text{ctrl}}$. After centering and scaling the control profiles, singular value decomposition gives

$$H_{\text{ctrl}} = U\Sigma V^\top.$$

Let $V_{(2)}$ denote the first two right singular vectors. Samples assigned to the same case batch are pooled at each grid location before projection. If $b(c)$ is the batch containing sample A in contrast $c$, and $\overline{\mathbf{q}}_{i,b(c)}$ is its pooled local quality-control profile, then

$$Z_c = \left[\frac{\overline{\mathbf{q}}_{i,b(c)} - \boldsymbol{\mu}_{\text{ctrl}}}{\boldsymbol{\sigma}_{\text{ctrl}}}\right]_{i\in\mathcal{I}} V_{(2)}.$$

For gene $g$, the basis is fixed and only its coefficients are gene-specific. Let

$$\widehat{\boldsymbol{\Delta}}_{gc} = \left(\widehat{\Delta}_{igc} : i \in \mathcal{I}_{gc}\right)^\top, \qquad Z_{+,c} = [\mathbf{1}, Z_c], \qquad W_{gc} = \operatorname{diag}\left(\widehat{V}_{igc}^{-1} : i \in \mathcal{I}_{gc}\right),$$

where the intercept column is omitted when `include_intercept=False`. Before fitting, the columns of $Z_c$ are centered under the local inverse-variance weights.

A gene-specific coefficient anchor $\boldsymbol{\alpha}_g^{(0)}$ is estimated from grid locations whose absolute initial local statistic lies in the lowest 30% in at least 60% of the fitted contrasts. The final contrast-specific coefficients are obtained from

$$\widehat{\boldsymbol{\alpha}}_{gc} = \left(Z_{+,c}^\top W_{gc} Z_{+,c} + \Lambda\right)^{-1} \left(Z_{+,c}^\top W_{gc} \widehat{\boldsymbol{\Delta}}_{gc} + \Lambda \boldsymbol{\alpha}_g^{(0)}\right),$$

where

$$\widehat{\boldsymbol{\alpha}}_{gc} = \left(\widehat{\gamma}_{gc}, \widehat{\boldsymbol{\beta}}_{gc}^\top\right)^\top.$$

The diagonal matrix $\Lambda$ applies ridge shrinkage to the confounding-covariate coefficients, with default penalty $\lambda_\beta = 10$, and to the intercept when an intercept anchor is fitted. Columns and penalty entries corresponding to disabled model components are omitted.

The adjusted local effect passed to the test statistic is

$$r_{igc} = \widehat{\Delta}_{igc} - \widehat{\gamma}_{gc} - \mathbf{z}_{ic}^\top \widehat{\boldsymbol{\beta}}_{gc},$$

with omitted terms set to zero. The residual reference distribution and its degrees of freedom are defined in the following subsection.

#### S3.7 Null reference distribution and residual degrees of freedom

For local statistic $t_{igc} = r_{igc}/\sqrt{\widehat{V}_{igc}}$, let $n_{gc}$ denote the number of grid locations with a valid residual and variance for gene $g$ and contrast $c$, and let $d_{gc}$ denote the number of columns fitted in the corresponding baseline design. The implemented residual degrees of freedom are

$$\nu_{gc} = \max(n_{gc} - d_{gc}, 5).$$

Thus, $d_{gc} = 0$ when neither an intercept nor a confounding-covariate basis is fitted, $d_{gc} = 1$ for an intercept-only baseline, and otherwise equals the number of columns in the fitted intercept-plus-covariate design. The lower bound of five is a numerical safeguard for the finite-sample reference. The implementation evaluates two-sided probabilities as

$$P_{igc} = 2F_{t_{\nu_{gc}}}(-|t_{igc}|).$$

The variance $\widehat{V}_{igc}$ combines the estimated base local-contrast variance with the alignment-mismatch and optional cell-type-composition variance factors. The fitted baseline coefficients, variance components, risk map and calibration coefficient enter the local reference distribution as plug-in estimates. The implementation adds no separate leverage or coefficient-estimation variance term. A small numerical variance floor stabilizes nearly constant neighborhoods.

#### S3.8 Implementation settings and fallback behavior

The implementation stores the shared grid, kernel-neighborhood structures, effective local support, risk maps, the optional $Z$ matrix for confounding-covariate adjustment, local weights, test statistics, q-values, and significant masks as reusable objects. Local tests require finite local variance and sufficient effective support in both samples. A small variance floor stabilizes local weights in nearly constant neighborhoods. The alignment-risk map is standardized, floored at zero, capped at the 95th percentile, and rescaled before entering $\phi_{igc}^{\text{align}}$. Unavailable cell-type information or a disabled cell-type adjustment sets $\phi_{ic}^{\text{cell}} = 1$. Disabling alignment-risk adjustment sets $\phi_{igc}^{\text{align}} = 1$, yielding the naive local weighted DE model. The final statistic uses the risk map, mismatch-calibration coefficient, cell-type-composition factor, confounding-covariate basis and fitted baseline as plug-in quantities.

##### S3.9 Gene-level and trajectory summaries

For a single contrast $c$, valid local $P$ values are combined by ACAT,

$$T_{gc} = \sum_{i \in \mathcal{I}_{gc}} a_{ic} \tan\{(0.5 - P_{igc})\pi\}, \qquad P_{gc}^{\mathrm{ACAT}} = 0.5 - \frac{1}{\pi} \arctan(T_{gc}),$$

where $a_{ic} \geq 0$ and $\sum_i a_{ic} = 1$.

For an ordered series, let $\widehat{\mu}_{A,ig}^{(c)}$ denote the kernel-smoothed local expression estimate for sample A in contrast $c$. The unsmoothed adjusted expression is

$$\widetilde{\mu}_{ig}^{(c)} = \widehat{\mu}_{A,ig}^{(c)} - \mathbf{z}_{ic}^{\top} \widehat{\boldsymbol{\beta}}_{gc},$$

where $\mathbf{z}_{ic}$ is the confounding-covariate basis constructed from local library size and detection rate, and $\widehat{\boldsymbol{\beta}}_{gc}$ is its fitted gene- and contrast-specific coefficient vector. Only this grid-varying component is removed; the fitted comparison-wide intercept $\widehat{\gamma}_{gc}$ is retained. No B-spline or other trajectory-level temporal smoothing is applied at this stage.

Let $x_c$ denote the ordered sample value, such as age. In the aging analysis, $c$ runs over the 19 non-reference sections; the 4.3-month reference is used in the corresponding local fits but does not provide a separate response value in the age regression. For every grid location with at least three valid sample values, weighted least squares fits

$$\widetilde{\mu}_{ig}^{(c)} = \alpha_{ig} + \beta_{ig} x_c + e_{igc},$$

using the mismatch-aware local precision $w_{igc} = \widehat{V}_{igc}^{-1}$. The two-sided statistic

$$t_{ig}^{\mathrm{trend}} = \frac{\widehat{\beta}_{ig}}{\mathrm{se}(\widehat{\beta}_{ig})}$$

is evaluated against a Student-$t$ distribution with $n_{ig} - 2$ degrees of freedom, yielding $P_{ig}^{\mathrm{trend}}$. These local trend $P$ values are combined with equal weights,

$$T_g^{\mathrm{trend}} = \frac{1}{|\mathcal{I}_g^{\mathrm{trend}}|} \sum_{i \in \mathcal{I}_g^{\mathrm{trend}}} \tan\{(0.5 - P_{ig}^{\mathrm{trend}})\pi\}, \qquad P_g^{\mathrm{trend}} = 0.5 - \frac{1}{\pi} \arctan(T_g^{\mathrm{trend}}).$$

The global null is zero linear age slope at every tested grid location. This test uses the unsmoothed adjusted-expression values and mismatch-aware local precisions and does not use trajectory smoothing, trajectory clusters or the selected $K$. Per-contrast spatial ACAT values may be retained as diagnostics but are not combined to form the reported ordered-series $P$ value. These are raw gene-level omnibus $P$ values and do not themselves provide across-gene FDR adjustment.

For ordered samples represented by contrasts $c = 1, \ldots, N_{\mathrm{ctr}}$, the trajectory-clustering input at grid location $i$ is

$$\widetilde{\boldsymbol{\mu}}_{ig} = \left( \widetilde{\mu}_{ig}^{(1)}, \ldots, \widetilde{\mu}_{ig}^{(N_{\mathrm{ctr}})} \right).$$

For the aging analysis, $N_{\mathrm{ctr}} = 19$. A missing trajectory entry, when present, is imputed by the average of its location-specific mean across samples and its sample-specific mean across locations.

Each adjusted-expression trajectory is smoothed across the ordered sample values using a cubic B-spline basis with eight basis functions and ridge parameter $10^{-2}$. The clustering feature vector combines the standardized smoothed trajectory, its standardized second differences and robustly scaled spatial coordinates,

$$\mathbf{f}_{ig} = \begin{bmatrix} \mathcal{Z}(\widetilde{\boldsymbol{\mu}}_{ig}^{\mathrm{sm}}) \\ 0.08\, \mathcal{Z}(\Delta^2 \widetilde{\boldsymbol{\mu}}_{ig}^{\mathrm{sm}}) \\ 0.20\, \mathcal{R}(\mathbf{s}_i) \end{bmatrix},$$

where $\mathcal{Z}$ denotes column-wise standardization across grid locations, $\Delta^2$ denotes the second difference across ordered samples, and $\mathcal{R}$ robustly centers and scales the spatial coordinates $\mathbf{s}_i = (x_i, y_i)^{\top}$ using their 2nd and 98th percentiles. The adjusted-expression trajectory therefore remains the primary clustering signal, while curvature and spatial position serve as weaker regularizing features. The values 0.08 and 0.20 are fixed implementation defaults and are not selected by the automatic-$K$ procedure.

A symmetric ten-nearest-neighbor graph is constructed on the shared grid using Gaussian edge weights with bandwidth equal to the median neighbor distance. Let $P$ be its row-normalized weight matrix. Before clustering, the feature matrix is smoothed for five iterations according to

$$F^{(m+1)} = 0.45 F^{(m)} + 0.55 P F^{(m)}.$$

This step promotes local spatial continuity without replacing the expression-trajectory features.

When local-statistic maps are available, reliability weights reduce the influence of weakly supported grid locations. Let $S_i \in [0,1]$ be the robustly rescaled, graph-smoothed mean absolute local statistic across contrasts, and let $U_i \in [0,1]$ be the corresponding rescaled, graph-smoothed fraction of contrasts in which location $i$ is significant. The raw reliability score is

$$r_i = (0.75 + 0.25S_i)(0.85 + 0.15U_i).$$

The second factor is omitted when significant masks are unavailable. The clustering weight is

$$\omega_i = \text{clip}\left[\frac{r_i}{\text{median}_{j:r_j>0}(r_j)}, 0.60, 1.80\right].$$

Thus, a typical location has weight near one, and no location receives less than 0.60 or more than 1.80 times the median influence. When local-statistic maps are unavailable or reliability weighting is disabled, $\omega_i = 1$. These fixed bounds prevent weak locations from being discarded and strong locations from dominating the partition.

For each candidate $K$, reliability-weighted k-means minimizes

$$\sum_i \omega_i \left\|\mathbf{f}_{ig} - \mathbf{m}_{z_i}\right\|^2,$$

using weighted k-means++ initialization and a fixed random seed. Candidate partitions use three starts and at most 80 iterations. A spatial refinement step then penalizes assignments that disagree with neighboring grid locations, using penalty weight 2.8 and at most eight refinement iterations. After $K$ has been selected, the final partition is refitted on the complete grid using six starts, at most 100 weighted k-means iterations and at most ten spatial-refinement iterations.

When $K$ is not supplied, the implementation evaluates $K = 2, \ldots, 9$ by default. The resulting candidate partitions are compared using the automatic selection procedure below. The feature weights, reliability weights and spatial-smoothing parameters above are fixed before candidate-$K$ evaluation. Cluster-level trend $P$ values are calculated only after the final partition has been selected and are not used to choose $K$.

Automatic selection first asks whether the candidate partition supports distinct time trends. For each $K$, the complete observed adjusted-expression trajectory is averaged within each cluster at every time point. On interleaved held-out time folds, a shared-trend model, which has cluster-specific intercepts but a common polynomial time trend, is compared with a cluster-specific-trend model, which additionally contains cluster-by-time polynomial terms. If $L_{Kf}^{\text{shared}}$ and $L_{Kf}^{\text{cluster}}$ are their cluster-mass-weighted squared prediction losses in held-out fold $f$, the dynamic gain is

$$G_{Kf} = \frac{L_{Kf}^{\text{shared}} - L_{Kf}^{\text{cluster}}}{\max(L_{Kf}^{\text{shared}}, \epsilon)}.$$

The implementation records the mean gain $\overline{G}_K$ and its standard error across folds. With at most three distinct time points, automatic selection returns the smallest candidate $K$. With four or five time points, it uses a linear trend and leave-one-time-point-out validation. With at least six time points, the maximum number of interleaved folds is $\min\{5, \lfloor T/2 \rfloor\}$; the highest feasible polynomial degree in $\{3, 2, 1\}$ and the largest feasible fold count are used subject to retaining at least $2(d+1)$ training time points for degree $d$.

Let $K^\star = \arg\max_K \overline{G}_K$ and $G_{\text{low}} = \overline{G}_{K^\star} - \text{SE}(\overline{G}_{K^\star})$. If $G_{\text{low}} \le 0$, there is no reliable held-out evidence for cluster-specific time trends and the smallest candidate $K$ is selected. Otherwise, the dynamic-evidence set is

$$\mathcal{K}_{\text{dyn}} = \left\{K : \overline{G}_K \ge G_{\text{low}}\right\}.$$

The second stage selects spatial resolution within this set. One $R_{\text{map}}$ footprint is defined as the median number of valid grid locations within radius $R_{\text{map}}$ of a grid location. For each $K \in \mathcal{K}_{\text{dyn}}$, connected components smaller than this footprint are treated as sub-resolution fragments, and $F_K$ denotes the fraction of grid locations belonging to such components. The dynamic candidates are examined from the largest $K$ to the smallest. Beginning with the finest candidate, the procedure identifies the first coarser candidate that fails to reduce $F_K$ and retains the immediately preceding, finer candidate, corresponding to the first fine-side local minimum. If $F_K$ decreases across the complete fine-to-coarse scan, the fragmentation sequence supplies no elbow; in that case, the procedure retains the candidate one step coarser than the finest dynamically supported candidate rather than continuing to the coarsest candidate. This rule uses fragmentation to locate a spatial-resolution elbow and does not treat global minimization of $F_K$ as the selection objective. Candidate screening may use a reliability-weighted subsample of at most 25,000 grid locations and keeps only the small center matrices needed for initialization; after selection, the final partition is fitted once on the full grid.

For trajectory cluster $\mathcal{C}_{g\kappa}$, the sample-level cluster mean for contrast $c$ is

$$\bar{\mu}_{g\kappa}^{(c)} = \frac{\sum_{i\in\mathcal{C}_{g\kappa}} \omega_i \widetilde{\mu}_{ig}^{(c)}}{\sum_{i\in\mathcal{C}_{g\kappa}} \omega_i},$$

using the trajectory-clustering reliability weight $\omega_i$ defined above. The corresponding weighted within-cluster spatial variance is

$$\left(s_{g\kappa}^{(c)}\right)^2 = \frac{\sum_{i\in\mathcal{C}_{g\kappa}} \omega_i \left(\widetilde{\mu}_{ig}^{(c)} - \bar{\mu}_{g\kappa}^{(c)}\right)^2}{\sum_{i\in\mathcal{C}_{g\kappa}} \omega_i}.$$

Each contrast contributes one value $\bar{\mu}_{g\kappa}^{(c)}$ to the cluster trajectory; in the aging analysis, each value corresponds to one non-reference section. The plotted error bars are

$$\bar{\mu}_{g\kappa}^{(c)} \pm 1.96 s_{g\kappa}^{(c)}$$

and depict the spatial dispersion among grid locations within that cluster and section.

The cluster-level trend test fits

$$\bar{\mu}_{g\kappa}^{(c)} = \beta_{0,g\kappa} + \beta_{1,g\kappa} x_c + e_{g\kappa}^{(c)}$$

over the ordered sample variable $x_c$, such as age. The weighted least-squares weight for contrast $c$ is the inverse squared spatial standard deviation,

$$v_{g\kappa}^{(c)} = \left\{\max\left(s_{g\kappa}^{(c)}, 10^{-8}\right)\right\}^{-2}.$$

If a spatial standard deviation is unavailable or non-positive, it is replaced by the median positive value across the available contrasts before the weight is calculated. Weighted least squares gives $\widehat{\beta}_{1,g\kappa}$ and $\mathrm{se}(\widehat{\beta}_{1,g\kappa})$. The Wald statistic is

$$W_{g\kappa} = \left\{\frac{\widehat{\beta}_{1,g\kappa}}{\mathrm{se}(\widehat{\beta}_{1,g\kappa})}\right\}^2,$$

with the corresponding two-sided Wald $P$ value

$$P_{g\kappa}^{\mathrm{trend}} = \Pr\left(\chi_1^2 \geq W_{g\kappa}\right).$$

This cluster-level trend calculation describes changes in a selected spatial trajectory domain and is distinct from the gene-level ACAT omnibus test. The weights $\{s_{g\kappa}^{(c)}\}^{-2}$ use within-cluster spatial variability as a reliability measure; they are not derived as inverse estimated variances of the section-level cluster means. Because the trajectory features, partition and $K$ are estimated from the same observed age series, the resulting Wald values are conditional downstream summaries rather than selection-independent confirmatory tests. A selection-aware analysis would need to repeat trajectory smoothing, clustering, $K$ selection and trend fitting under the null.

Cluster-level trend results are reported as raw two-sided Wald $P$ values; no additional within-gene BH adjustment was applied across the selected clusters.

For the aging-brain gene-level test, the local age-trend $P$ values received equal weights across valid grid locations. The values in Fig. 5B are raw gene-level age-trend ACAT $P$ values. For the single-contrast kidney analysis, valid local $P$ values were combined with equal weights within the IL3-versus-NL3 contrast. The resulting kidney gene-level values were adjusted by the Benjamini–Hochberg procedure across all 16,446 fitted genes; $q_g^{\mathrm{ACAT}} \leq 0.05$ defined eligibility before compartment–direction classification and Gene Ontology analysis.

##### S3.10 Simulation data generation and controlled coordinate perturbation

The simulations used the observed coordinates, cell-type labels and library sizes from the 3.8-month S2R2 mouse-brain section. For *Gamt* and the 400 genes with the highest mean expression, baseline count models were fitted in R using `mgcv::bam` with a negative-binomial response, an offset for log library size, a cell-type effect and a two-dimensional thin-plate spatial smooth. The fits used fast restricted maximum likelihood and smooth-basis dimension $k = 80$. Their fitted means and dispersion parameters defined gene-wise count distributions from which samples A and B were generated conditionally independently on the same tissue template.

Six non-overlapping circular signal regions were placed within well-sampled tissue, with three assigned positive and three assigned negative effects. Each region contained at least 200 cells, and its radius was sampled between $0.025L$ and $0.06L$, where $L$ is the larger side length of the tissue bounding box. In sample B, the fitted mean of *Gamt* was multiplied by 4 in positive regions and by 0.20 in negative regions before negative-binomial count generation. The other simulated genes retained the fitted baseline or received region-specific modulation according to the simulation design.

Three distortion regions were generated separately from the signal regions. Within each distortion region, coordinates were transformed by a local affine perturbation whose contribution was smoothly tapered at the region boundary, thereby producing localized correspondence error without a discontinuity in the coordinate field. Sample B additionally received a smooth radial-basis-function deformation, a global affine transformation and coordinate noise; sample A retained the reference coordinate system.

Ten matched simulation groups were accepted. DE-region placement and distortion-region placement varied across groups, whereas the non-region random streams were held fixed to make differences among groups attributable primarily to their spatial configurations. A candidate group was rejected if its signal, distortion, combined, upregulated or downregulated mask had Jaccard overlap greater than 0.25 with the corresponding mask in any previously accepted group.

**S3.10.1 Controlled coordinate-perturbation analyses** We evaluated the sensitivity of local inference to residual coordinate error by perturbing the oracle sample-B coordinates after the simulation truth had been defined. At perturbation level $u$, independent Gaussian noise was added to both coordinate axes with per-axis standard deviation $uh_{\mathrm{grid}}$, where $h_{\mathrm{grid}}$ denotes the shared-grid spacing. Alignment was not rerun after introducing this noise.

We first compared the default shared-grid workflow with a cell-centered local-kernel baseline. The shared-grid workflow estimates local contrasts at fixed grid anchors, while the baseline performs local inference directly around cell-centered neighborhoods. We then held the shared-grid construction fixed and compared naive testing with mismatch-aware variance inflation. The first comparison evaluates the stability provided by fixed grid anchors and smoothing; the second isolates the additional contribution of mismatch-aware adjustment.

Robustness was summarized by correlation with the unperturbed local statistic map, location-level FDP for calls made at a target FDR of 0.05, and power at FDP $\leq 0.05$, as defined above.

##### S3.11 Aging-brain application settings

The aging-brain analysis used 20 age-specific coronal MERFISH sections and the 4.3-month section as the fixed reference. Each non-reference section supplied one age-versus-reference contrast. Young and old summaries used sections aged $\leq 13.6$ months ($n = 7$) and $\geq 24.3$ months ($n = 7$), respectively.

*Spatial cell-type neighborhood analysis.* Cell-type neighborhood analyses were performed on the aligned cell coordinates. Let $\mathcal{T}_s$ denote the T cells in section $s$, and let $\mathcal{C}_{\ell s}$ denote cells of target type $\ell$. The section-level nearest-neighbor distance was

$$d_{\ell s} = \mathrm{median}_{j\in\mathcal{T}_s} \min_{k\in\mathcal{C}_{\ell s}} \left\| \widetilde{\mathbf{q}}_j^{(s)} - \widetilde{\mathbf{q}}_k^{(s)} \right\|.$$

We defined the proximity score as $P_{\ell s} = -\log(d_{\ell s} + \epsilon)$ and regressed it on age. A positive fitted age coefficient therefore indicates decreasing T-cell-to-target distance with age.

To measure spatial overlap, we constructed kernel-smoothed density fields for T cells and each target cell type on the shared grid. For target type $\ell$,

$$D_{\ell s}(\boldsymbol{\xi}_i) = \sum_{j\in\mathcal{C}_{\ell s}} K_h\left( \left\| \widetilde{\mathbf{q}}_j^{(s)} - \boldsymbol{\xi}_i \right\| \right),$$

with an analogous field $D_{Ts}(\boldsymbol{\xi}_i)$ for T cells. Spatial overlap was defined as the weighted correlation between $D_{Ts}$ and $D_{\ell s}$ across valid grid locations, using weights $w_{is} = \{1 + \widetilde{R}_{is}^{\mathrm{loc}}\}^{-1}$. The overlap summaries were regressed on age, and positive coefficients indicate increasing spatial co-localization. Separate $q$-values were computed across the tested target cell types for the proximity and overlap analyses.

Continuous local T-cell enrichment maps were constructed by subtracting the kernel-smoothed tissue-area expectation from the kernel-smoothed local T-cell mass. Young and old maps were obtained by averaging within the corresponding age strata. For each prespecified region of interest, the association between mean local enrichment and age was tested, followed by BH adjustment across the displayed regions. The displayed ROIs were defined by their distance from the corpus callosum and are referred to as callosal-proximal and callosal-distal.

Figure 5C compares the 21.4-month section with the 4.3-month reference for *Gamt*. Both panels use the same local model settings, with the cell-type-composition adjustment switched off or on.

The young stratum comprised sections aged 3.4, 3.8, 4.3, 5.4, 6.6, 9.8 and 12.9 months; the old stratum comprised sections aged 24.6, 26.7, 28.5, 30.9, 32.6, 33.2 and 34.5 months. Counts were normalized to a target total of 250. The shared grid contained 77,056 valid locations with spacing 27.05, $h_{\mathrm{loc}} = 121.1$ and $R_{\mathrm{map}} = 40.57$. Each age-versus-reference fit included the comparison-wide intercept and local library-size and detection-rate covariates. For each gene, provisional mismatch-inflation coefficients were calibrated separately within the 19 age-versus-reference contrasts and combined by an equal-weight Huber robust center to obtain the shared $\widehat{\lambda}_g$; each contrast retained its own local risk map. Cell-type adjustment was disabled for Fig. 5A–B and switched off or on for Fig. 5C; the latter used $\alpha = 0.10$ for the local $q$-value maps.

Cell-type-composition vectors used the common set of annotated cell types after local kernel smoothing. T-cell neighborhood analyses included oligodendrocytes, microglia and ependymal cells as target classes. The local tissue-area expectation was estimated from the kernel-smoothed all-cell density on the same shared grid. The callosal-proximal and callosal-distal circular regions had radius 3.2% of the maximum aligned coordinate span and were specified in the aligned reference coordinate system from the old-minus-young local T-cell enrichment map. Their trajectories used Gaussian age smoothing with bandwidth 4.5 months.

##### S3.12 Injured-kidney analysis and over-representation analysis

The injured IL3 section was aligned as the query to the normal NL3 section as the reference. For post-alignment inference, injured IL3 was sample A and normal NL3 was sample B, so positive contrasts indicate higher expression in injury.

*Direction-specific local-grid classification and gene-level eligibility.* Let $\mathcal{I}_g$ be the valid shared-grid locations for gene $g$, let $\mathcal{A}_h$ denote grid locations assigned to kidney compartment $h$, and let $d \in \{-1, +1\}$ denote decrease and increase, respectively. The direction-specific significant set is

$$\mathcal{S}_{gd} = \left\{ i \in \mathcal{I}_g : q_{ig} \leq 0.05,\ \operatorname{sign}(\widehat{\Delta}_{ig}) = d \right\}.$$

For each gene, direction and compartment, we calculated the fraction of significant locations falling in the compartment and the corresponding fraction of valid tissue area,

$$\rho_{gdh} = \frac{|\mathcal{S}_{gd} \cap \mathcal{A}_h|}{|\mathcal{S}_{gd}|}, \qquad \pi_{gh} = \frac{|\mathcal{I}_g \cap \mathcal{A}_h|}{|\mathcal{I}_g|}.$$

A one-sided Fisher exact test compared membership in $\mathcal{S}_{gd}$ with membership in $\mathcal{A}_h$ over the valid grid. A compartment–direction pair was eligible for spatial classification when $|\mathcal{S}_{gd}| \geq 5$, $|\mathcal{S}_{gd} \cap \mathcal{A}_h| \geq 3$, $\rho_{gdh} - \pi_{gh} \geq 0.03$, and the one-sided Fisher $P$ value was at most 0.05. If more than one pair was eligible, the gene was assigned to a single dominant class by the smallest Fisher $P$ value, followed by the largest $\rho_{gdh} - \pi_{gh}$ and then the largest number of significant locations in the compartment. Genes without an eligible pair were not assigned to an anatomical local-ORA class. This Fisher screen was used for spatial classification and was not interpreted as a separate region-level FDR guarantee.

Raw local $P$ values were combined with equal weights by ACAT within the single IL3-versus-NL3 contrast. Genes with no valid local tests were assigned an ACAT $P$ value of one. The resulting gene-level $q_g^{\mathrm{ACAT}}$-values were computed across the complete predeclared family of 16,446 fitted genes. For compartment $h$ and direction $d$, the formal local-ORA gene set was

$$\mathcal{G}_{h,d}^{\mathrm{local}} = \left\{ g : C_g = (h, d), q_g^{\mathrm{ACAT}} \leq 0.05 \right\},$$

where $C_g$ is the dominant class selected above. Thus, local spatial concentration and $q_g^{\mathrm{ACAT}} \leq 0.05$ were both required, while the ORA universe remained the complete 16,446-gene testing family.

*GO Biological Process over-representation and term recurrence.* Each local compartment–direction gene set was analyzed with `clusterProfiler::enrichGO`, using `org.Mm.eg.db`, gene symbols as keys and the Biological Process ontology. GO categories represented by fewer than 10 or more than 100 genes in the analysis universe were excluded by `minGSSize=10` and `maxGSSize=100`. Term $q$-values were computed separately within each compartment–direction class, and $q \leq 0.05$ defined significant terms. Redundant GO terms were simplified at semantic-similarity cutoff 0.7, retaining the term with the smallest $q$-value. Displayed terms were ranked within each compartment–direction class by increasing term $q$-value, with increasing raw over-representation $P$ value, decreasing `Count` and alphabetical GO term name used successively to resolve ties. Here, `Count` is the number of genes from the class-specific input set that overlap the GO term. These term-level $q$-values are distinct from the gene-level $q_g^{\mathrm{ACAT}}$-values used for eligibility.

For a significant GO term $u$, recurrence is defined as

$$\kappa(u) = \sum_{h,d} \mathbf{1}\{u \in \mathcal{T}_{h,d}\},$$

where $\mathcal{T}_{h,d}$ is the set of significant terms for a cortex-, interface- or medulla-direction class. The recurrence plot reports the fractions of significant terms with $\kappa(u) = 1$, $\kappa(u) = 2$, or $\kappa(u) \geq 3$.

*Descriptive annotation of kidney-injury-associated terms.* Orange labels were assigned after statistical ranking using a fixed keyword-based annotation rule covering renal function; injury, stress and repair; p53 and apoptosis; immune and inflammatory processes; mitochondrial energy and central-carbon metabolism; fatty-acid, coenzyme A and triglyceride metabolism; and antioxidant defense. The nonspecific cardiac-muscle apoptosis term was explicitly excluded. This annotation affected neither term selection nor ordering.

*Compartment-level differential expression baseline.* For comparison, compartment DE ORA began with predefined anatomical groups. Each spot was assigned to the cortex, interface or medulla. Within each compartment containing at least 20 valid spots from each section, expression was compared between the injured IL3 and normal NL3 sections for every gene shared by the two sections using a two-sided Wilcoxon rank-sum test. The direction was defined by the sign of

$$\log\bigl(1+\overline{X}_{\mathrm{IL3},cg}\bigr)-\log\bigl(1+\overline{X}_{\mathrm{NL3},cg}\bigr)\,,$$

where $\overline{X}_{s,cg}$ is the mean expression of gene $g$ among spots assigned to compartment $c$ in section $s$. Gene-level $q$-values were computed separately within each compartment, and genes with $q \leq 0.05$ were divided into increase and decrease sets. Each compartment–direction gene set was then tested for GO Biological Process over-representation against the common set of detectable genes, yielding term $q$-values within that class. We summarized term recurrence by the number of compartment–direction classes in which each significant term appeared. This baseline differs from local DE ORA because the anatomical groups are fixed before differential testing and all spots in a compartment contribute to one compartment-level result.

The final Gene Ontology analyses used `clusterProfiler` version 4.18.4 and `org.Mm.eg.db` version 3.22.0.

### S4 Supplementary Figures

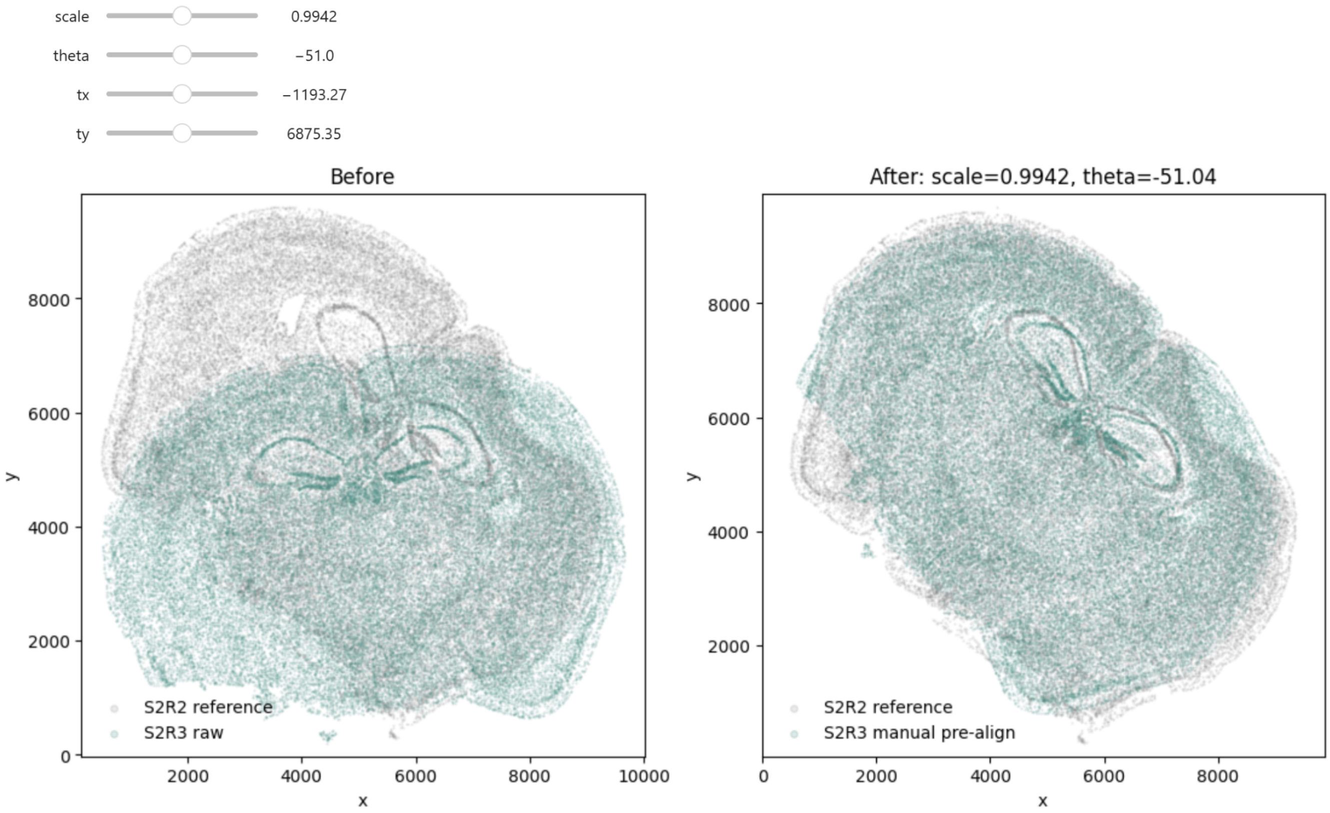


Fig. S1: **Interactive global pre-alignment interface.** The interface displays the query and reference datasets in an overlaid view and allows users to interactively adjust global scaling, rotation, and horizontal and vertical translation. The selected parameters define an approximate similarity transformation that places the two tissues into a common coordinate frame before structure pairing and S-LDDMM refinement. The parameters and transformation matrix are saved for reproducible application to all query coordinates and query-derived spatial structures. This step estimates only the global initialization and does not define structure correspondences or local anatomical deformation.

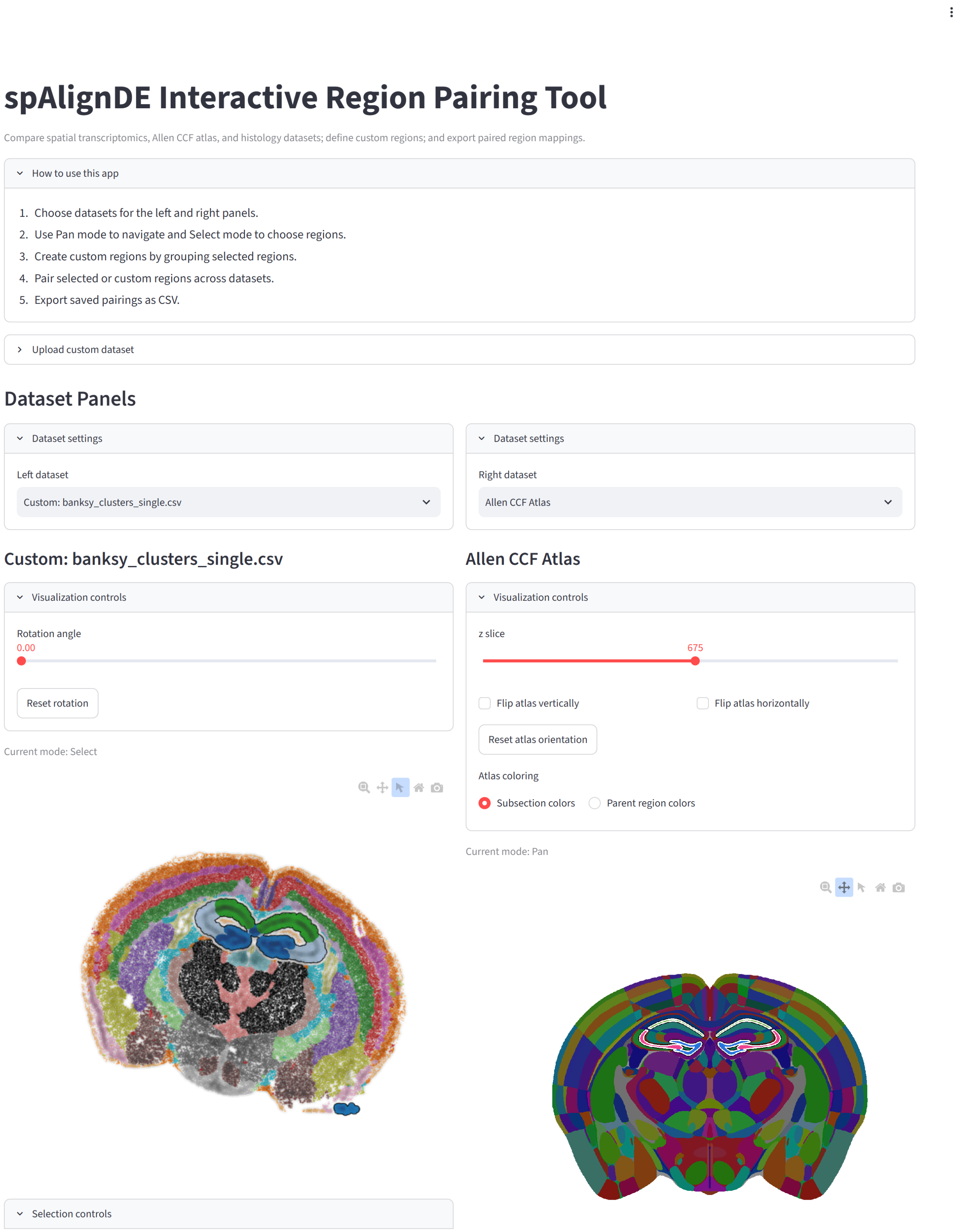


Fig. S2: **Interactive cross-modality structure pairing with spAlignDE Structure Pair.** The interface displays query and reference spatial structures side by side and allows users to inspect, select and group corresponding structures. Multiple primitive structures can be combined into custom structures, supporting one-to-one, one-to-many, many-to-one and many-to-many correspondences. The selected correspondence groups and display-orientation settings are exported reproducibly and converted into matched signed-distance-transform channels for downstream S-LDDMM alignment.

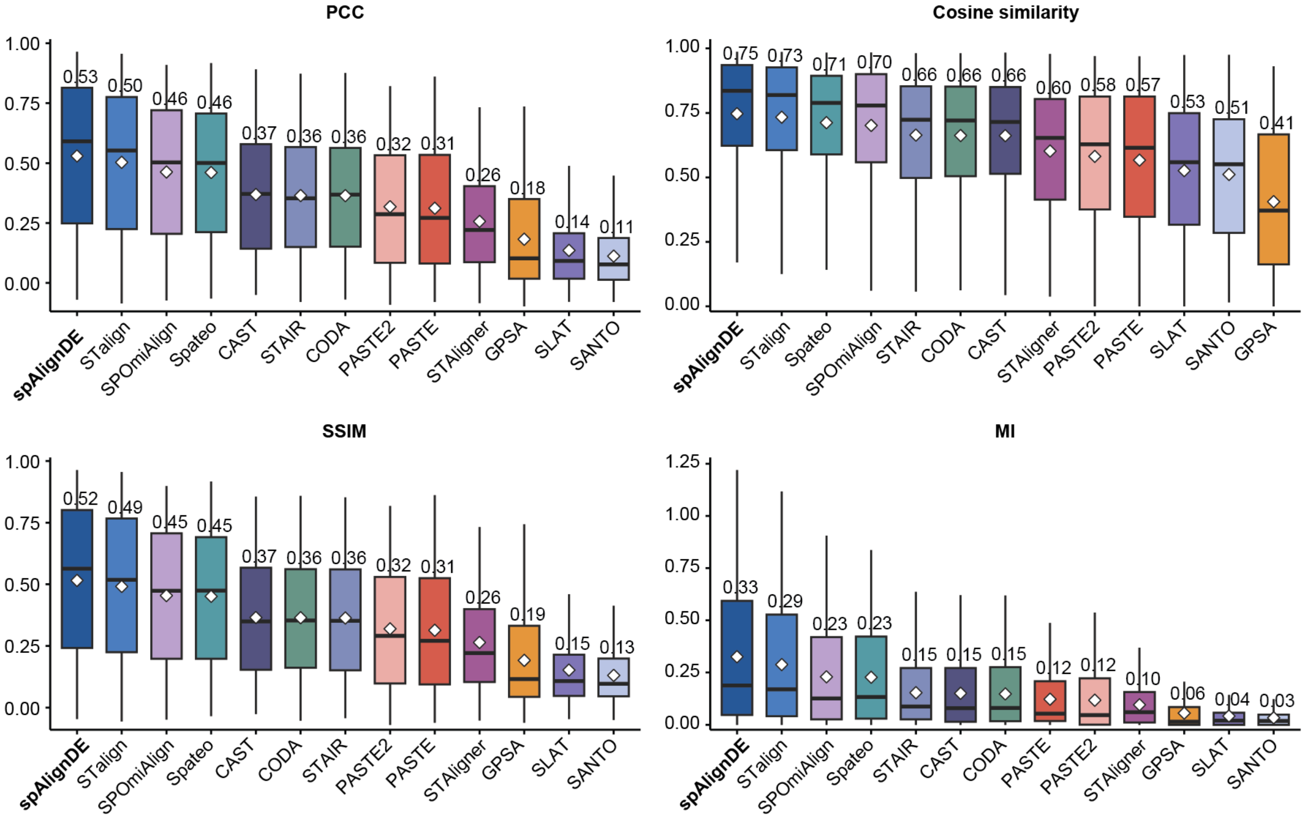


Fig. S3: **Gene-expression concordance after cross-sample alignment using a $30 \times 30$ evaluation grid.** The aligned MERFISH mouse-brain sections S2R3 and S2R2 were evaluated across 483 shared genes and 13 alignment methods. From top left to bottom right, the boxplots show the distributions of Pearson correlation coefficient (PCC), cosine similarity, structural similarity index measure (SSIM) and mutual information (MI) across genes. The aligned coordinates were unchanged from the primary benchmark; only the spatial grid used to aggregate expression and evaluate concordance was refined from $10 \times 10$ to $30 \times 30$. Boxes indicate the interquartile range, horizontal lines indicate medians, whiskers extend to the most extreme values within $1.5\times$ the interquartile range, and white diamonds indicate method-level means. Numeric labels above the boxes report the corresponding means, and outliers are not shown. Methods are ordered independently by their mean value for each metric. Higher values indicate stronger spatial expression concordance.

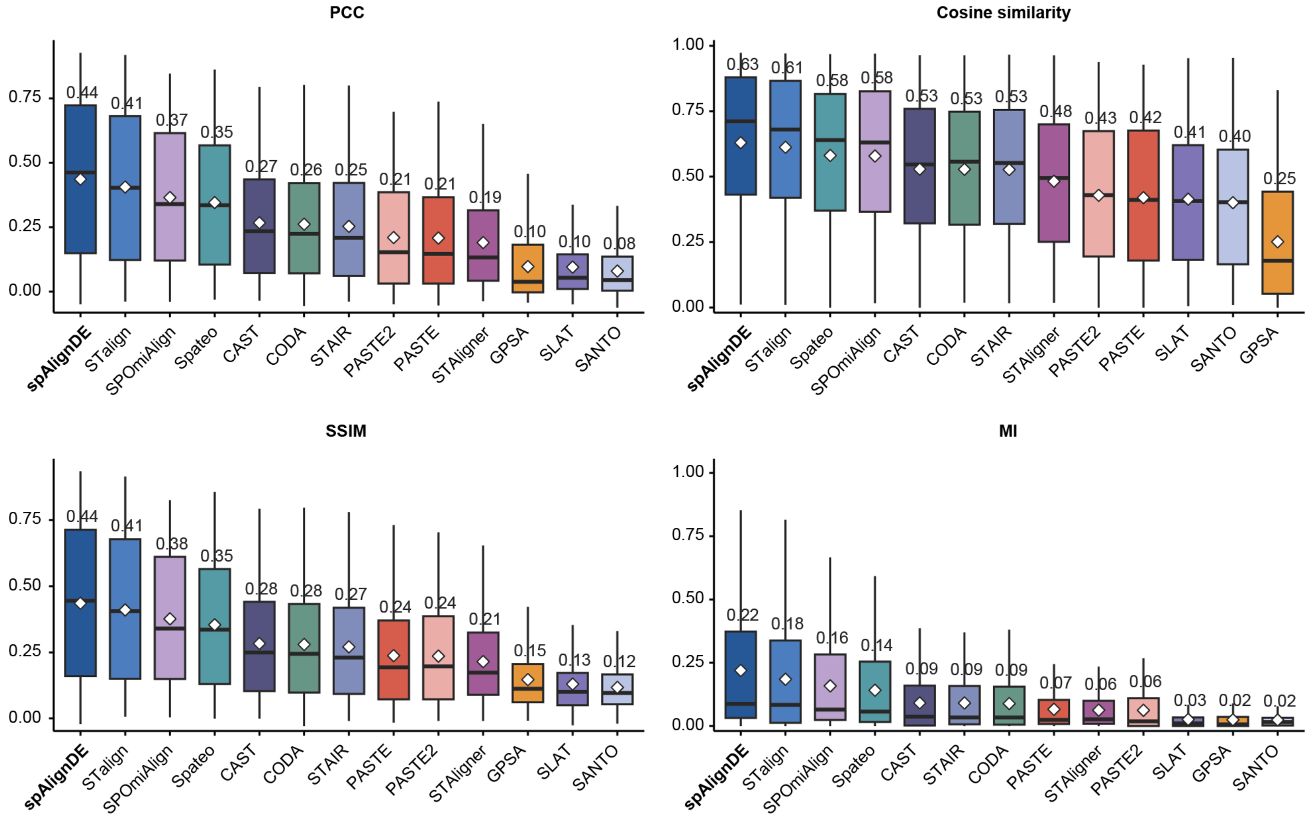


Fig. S4: **Gene-expression concordance after cross-sample alignment using a $50 \times 50$ evaluation grid.** The aligned MERFISH mouse-brain sections S2R3 and S2R2 were evaluated across 483 shared genes and 13 alignment methods. From top left to bottom right, the boxplots show the distributions of Pearson correlation coefficient (PCC), cosine similarity, structural similarity index measure (SSIM) and mutual information (MI) across genes. The aligned coordinates were unchanged from the primary benchmark; only the spatial grid used to aggregate expression and evaluate concordance was refined from $10 \times 10$ to $50 \times 50$. Boxes indicate the interquartile range, horizontal lines indicate medians, whiskers extend to the most extreme values within $1.5\times$ the interquartile range, and white diamonds indicate method-level means. Numeric labels above the boxes report the corresponding means, and outliers are not shown. Methods are ordered independently by their mean value for each metric. Higher values indicate stronger spatial expression concordance.

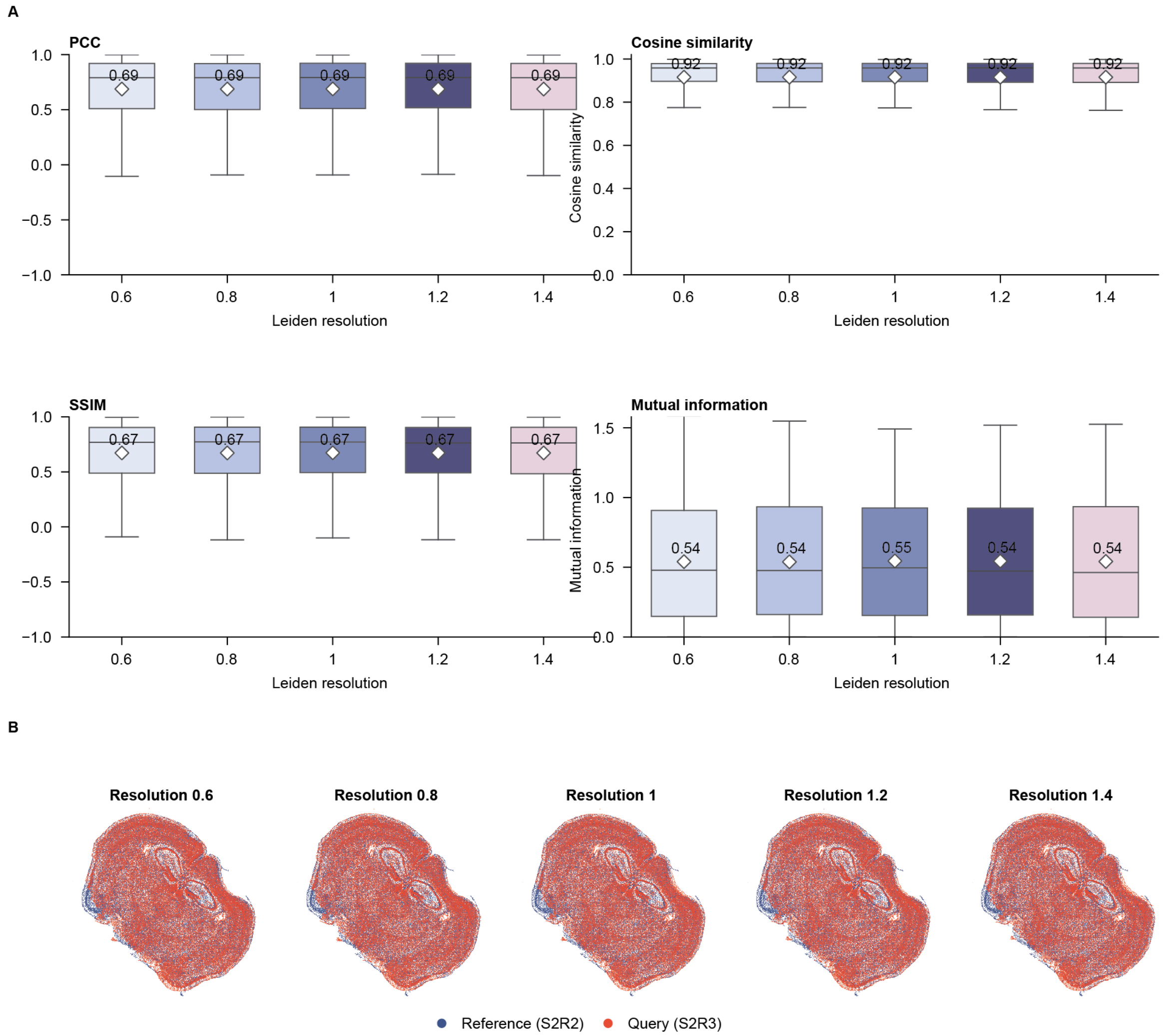


Fig. S5: **Cross-sample alignment is robust to joint-clustering resolution.** Section S2R3 was aligned to the S2R2 reference using Leiden resolutions of 0.6, 0.8, 1.0, 1.2 and 1.4; only the joint-clustering resolution was varied between runs, and 1.4 was the default setting used in the primary benchmark. **A,** Gene-pattern preservation after alignment, quantified across 483 shared genes using Pearson correlation coefficient (PCC), cosine similarity, structural similarity index measure (SSIM) and mutual information (MI). Boxes show the interquartile range, center lines indicate medians, whiskers extend to 1.5× the interquartile range and white diamonds indicate means; outliers are not shown. Values above the diamonds report the corresponding means. **B,** Whole-section overlays after alignment at the five respective resolutions. Reference cells are shown in blue and aligned query cells in red.

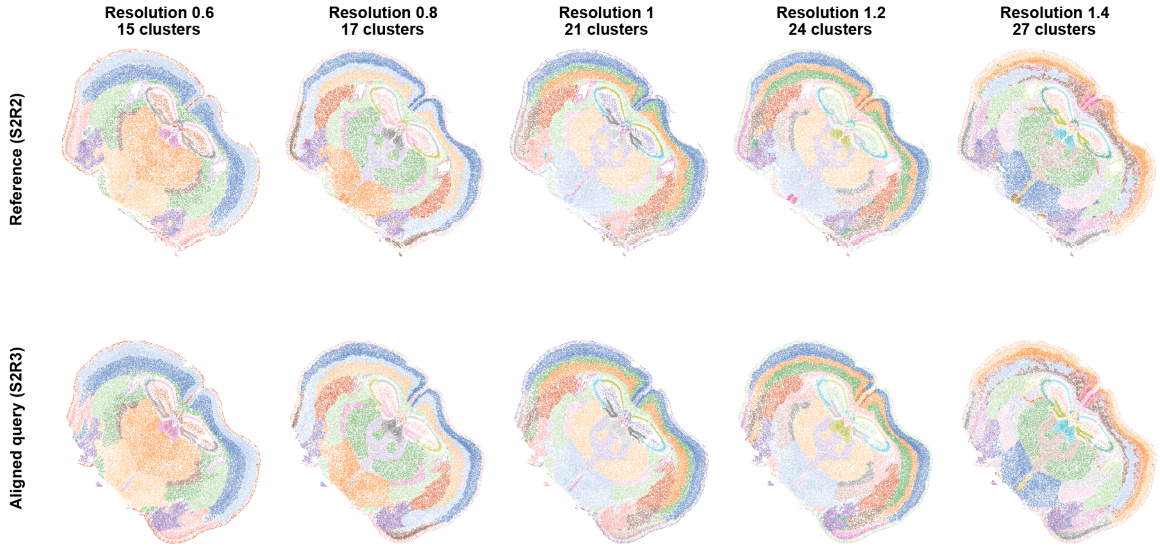


Fig. S6: **Spatial domains after alignment across joint-clustering resolutions.** The top row shows joint spatial clusters in the S2R2 reference at Leiden resolutions of 0.6, 0.8, 1.0, 1.2 and 1.4, yielding 15, 17, 21, 24 and 27 clusters, respectively. The bottom row shows the corresponding joint-cluster assignments in the aligned S2R3 query after alignment at each resolution. Cluster colors are shared between the reference and aligned query within each resolution but are resolution-specific and therefore do not denote cluster correspondence across columns. All maps use identical spatial limits.

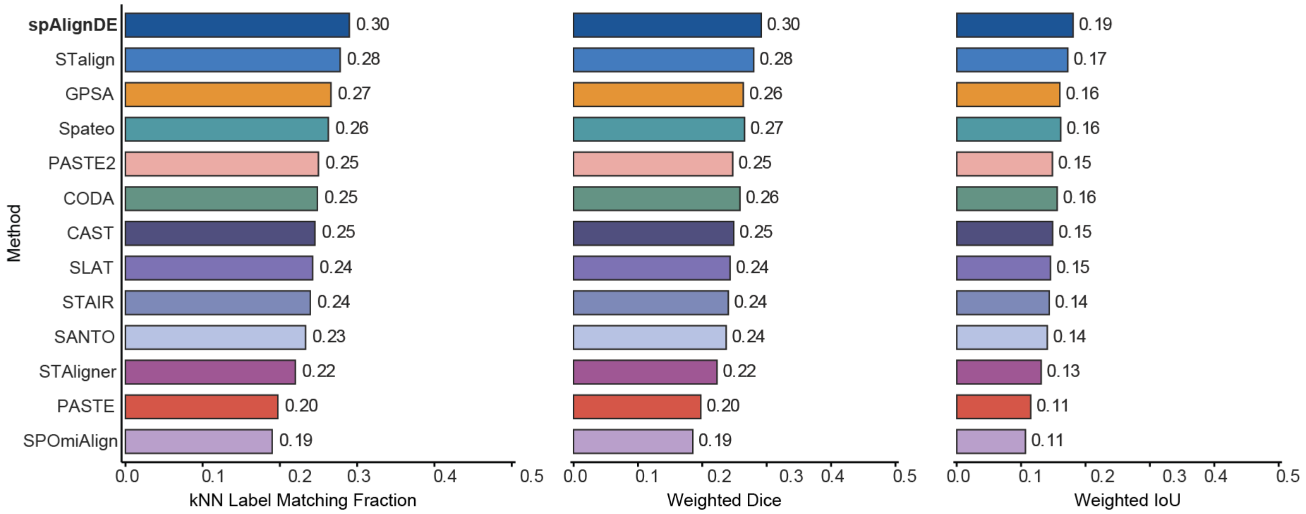


Fig. S7: **Cell-type annotation concordance after cross-sample alignment in the aging mouse brain MERFISH cohort.** Each of the 19 non-reference sections was aligned to the 4.3-month reference section. Alignment consistency was evaluated by comparing the original cell-type annotations of aligned query cells with annotations transferred from spatially neighboring reference cells, using the 20-nearest-neighbor same-label fraction, weighted Dice coefficient and weighted intersection over union (IoU). Bars show means across the 19 source-to-reference alignments, and higher values indicate greater annotation concordance. Methods are ordered by their mean 20-nearest-neighbor same-label fraction. GPSA used 10,000 cells per section, PASTE and PASTE2 used 30,000 cells per section because of computational constraints, and the remaining methods used all available cells.

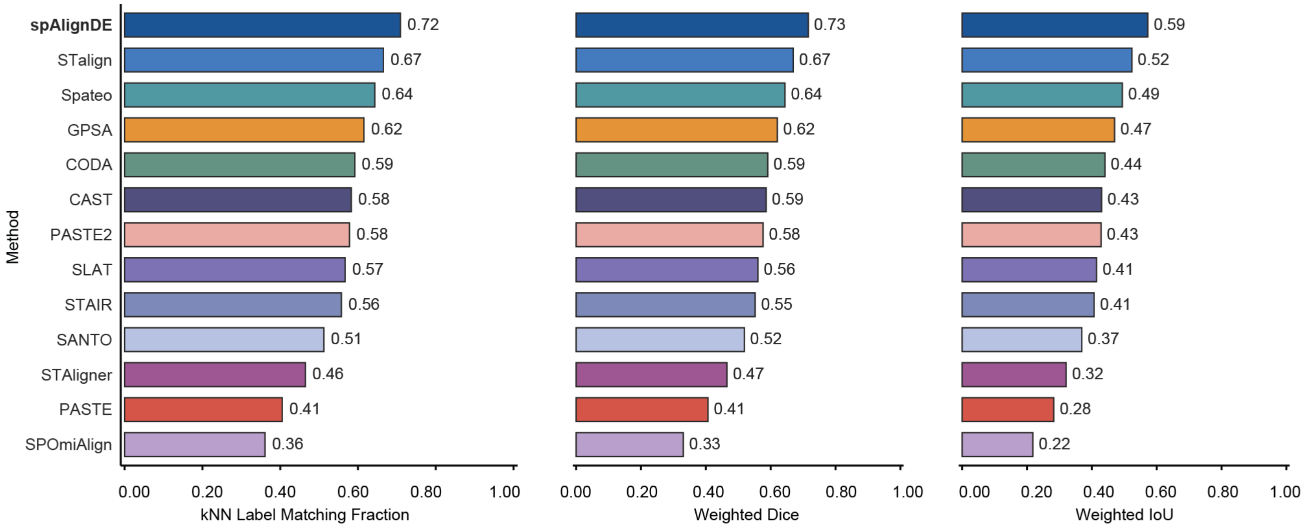


Fig. S8: **Subregion annotation concordance after cross-sample alignment in the aging mouse brain MERFISH cohort.** Each of the 19 non-reference sections was aligned to the 4.3-month reference section. Alignment consistency was evaluated by comparing the original subregion annotations of aligned query cells with annotations transferred from spatially neighboring reference cells, using the 20-nearest-neighbor same-label fraction, weighted Dice coefficient and weighted intersection over union (IoU). Bars show means across the 19 source-to-reference alignments, and higher values indicate greater annotation concordance. Methods are ordered by their mean 20-nearest-neighbor same-label fraction. GPSA used 10,000 cells per section, PASTE and PASTE2 used 30,000 cells per section because of computational constraints, and the remaining methods used all available cells.

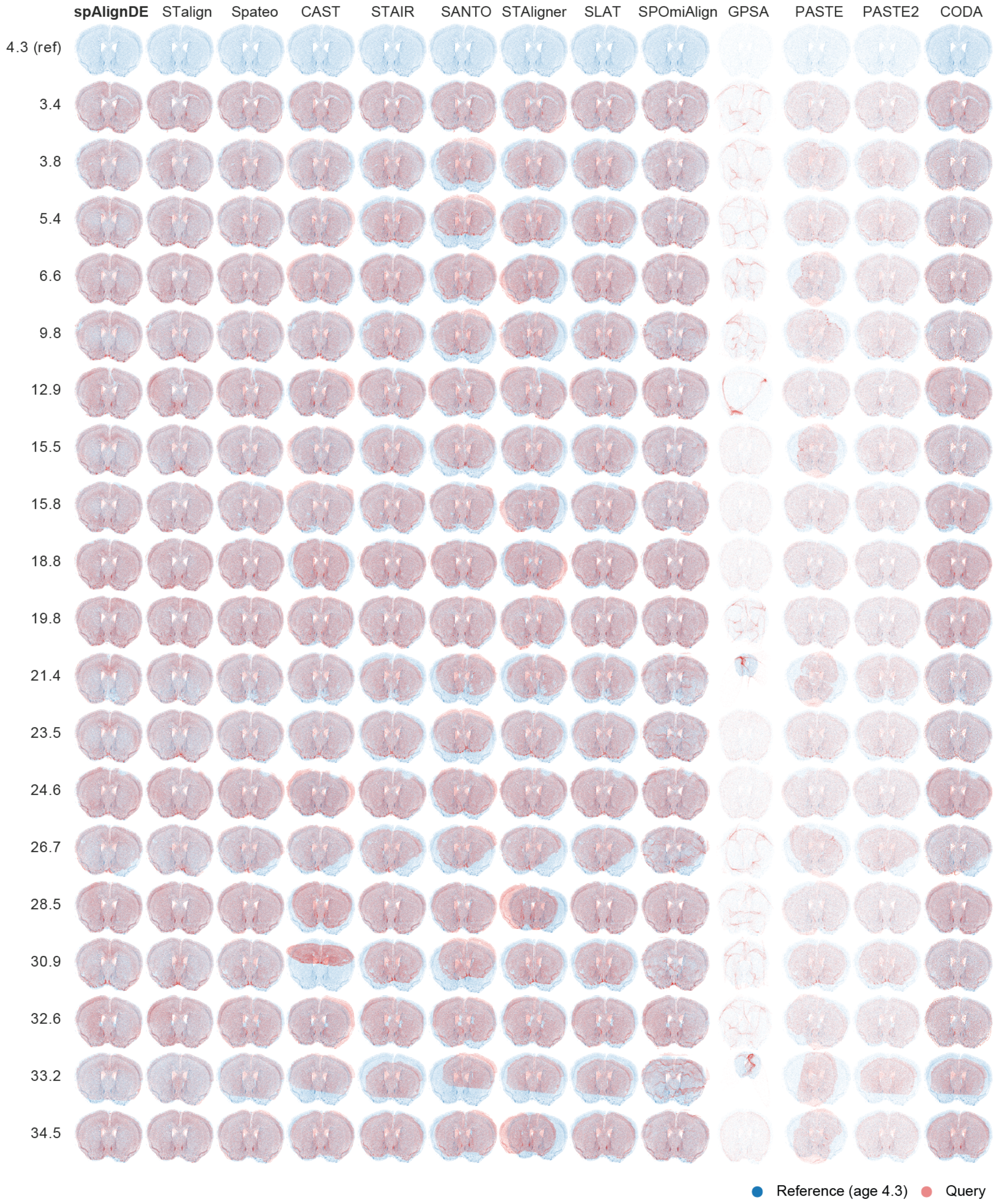


Fig. S9: **Post-alignment spatial overlap across the aging mouse brain cohort.** Rows correspond to the 13 evaluated alignment methods, and columns correspond to the 4.3-month reference section followed by the 19 non-reference age-specific sections. The first column shows the 4.3-month reference coordinates in blue. In each remaining panel, the query cells from the indicated age are shown in red after alignment and overlaid on the fixed 4.3-month reference cells shown in blue. Red–blue spatial overlap indicates global correspondence between the aligned sections, whereas regions dominated by one color indicate incomplete shared tissue coverage or residual local mismatch. GPSA used 10,000 cells per section, PASTE and PASTE2 used 30,000 cells per section because of computational constraints, and the remaining methods used all available cells.

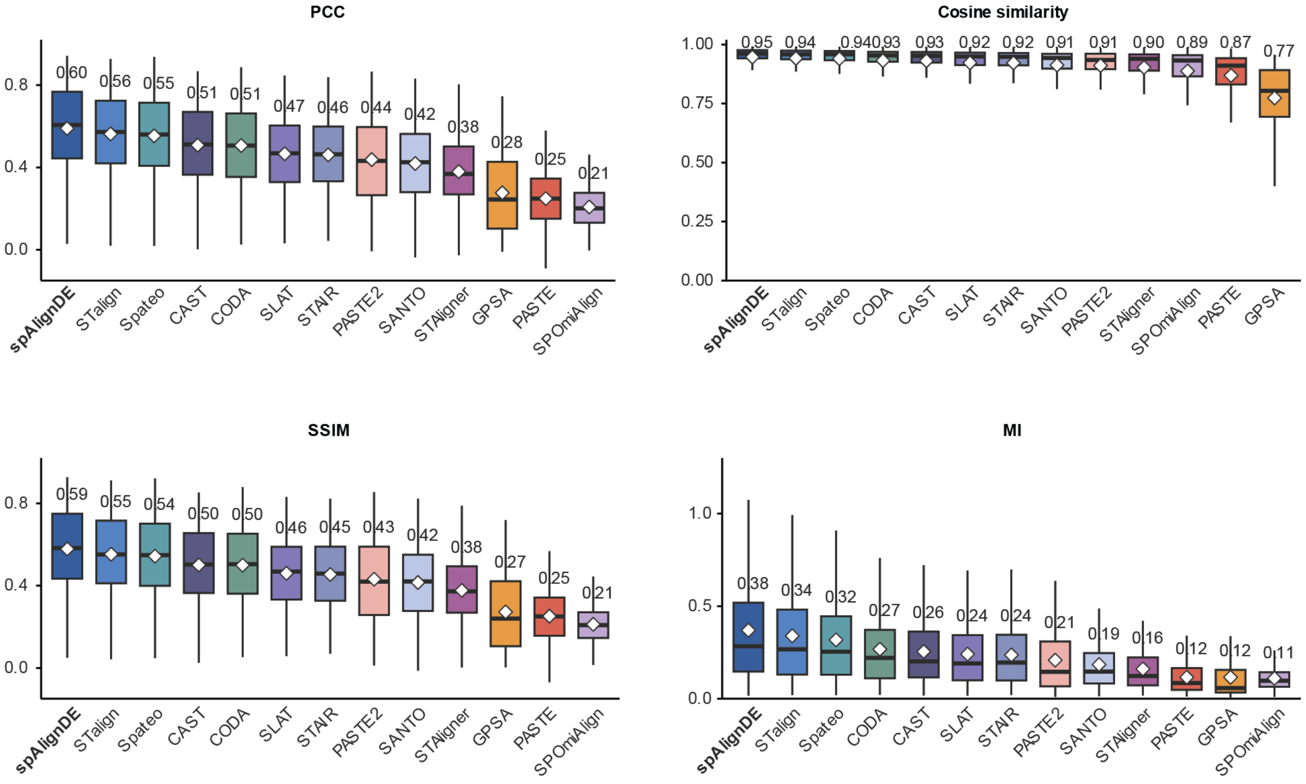


Fig. S10: **Quantitative benchmarking of cross-sample alignment in the aging mouse brain MERFISH cohort using a** $10{\times}10$ **evaluation grid.** All 19 non-reference sections were aligned to the 4.3-month reference section. Post-alignment gene-pattern concordance was evaluated across 300 genes using Pearson correlation coefficient (PCC), cosine similarity, structural similarity index measure (SSIM) and mutual information (MI). For each gene and query section, expression patterns were aggregated on a $10 \times 10$ spatial grid and compared with the 4.3-month reference; each metric was then averaged across the available query-to-reference alignments for that gene. Boxes show interquartile ranges across genes, center lines indicate medians, whiskers extend to 1.5× the interquartile range and white diamonds indicate method-level means; outliers are not shown. Numeric labels above the boxes report the corresponding means. All 13 methods were evaluated across all 19 query-to-reference alignments and were ordered independently by decreasing mean within each metric. Higher values indicate stronger post-alignment spatial expression concordance. spAlignDE achieved the highest mean value for all four metrics. GPSA used 10,000 cells per section, PASTE and PASTE2 used 30,000 cells per section because of computational constraints, and the remaining methods used all available cells.

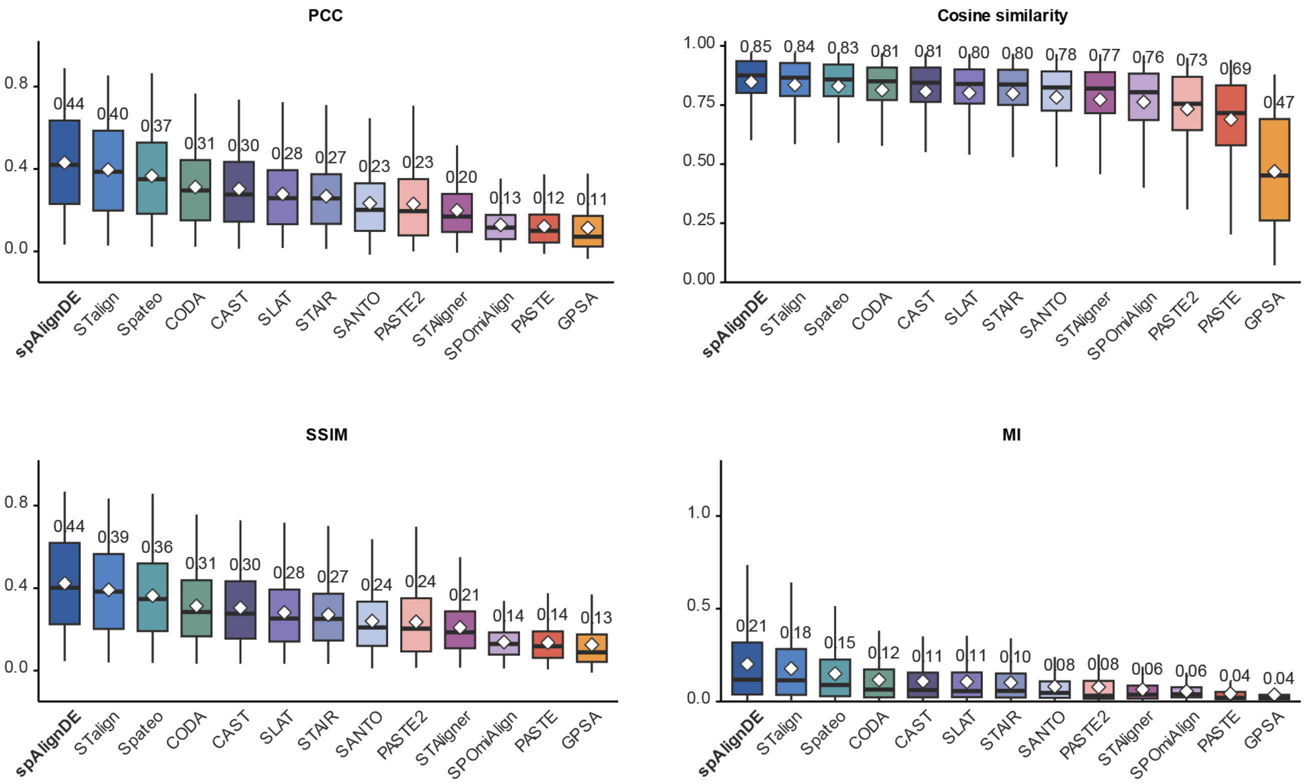


Fig. S11: **Quantitative benchmarking of cross-sample alignment in the aging mouse brain MERFISH cohort using a 30×30 evaluation grid.** All 19 non-reference sections were aligned to the 4.3-month reference section. Post-alignment gene-pattern concordance was evaluated across 300 genes using Pearson correlation coefficient (PCC), cosine similarity, structural similarity index measure (SSIM) and mutual information (MI). For each gene and query section, expression patterns were aggregated on a $30 \times 30$ spatial grid and compared with the 4.3-month reference; each metric was then averaged across the available query-to-reference alignments for that gene. The aligned coordinates were unchanged from the primary $10 \times 10$ evaluation; only the spatial grid used to aggregate expression and calculate concordance was refined. Boxes show interquartile ranges across genes, center lines indicate medians, whiskers extend to 1.5× the interquartile range and white diamonds indicate method-level means; outliers are not shown. Numeric labels above the boxes report the corresponding means. All 13 methods were evaluated across all 19 query-to-reference alignments and were ordered independently by decreasing mean within each metric. Higher values indicate stronger post-alignment spatial expression concordance. spAlignDE achieved the highest mean value for all four metrics. GPSA used 10,000 cells per section, PASTE and PASTE2 used 30,000 cells per section because of computational constraints, and the remaining methods used all available cells.

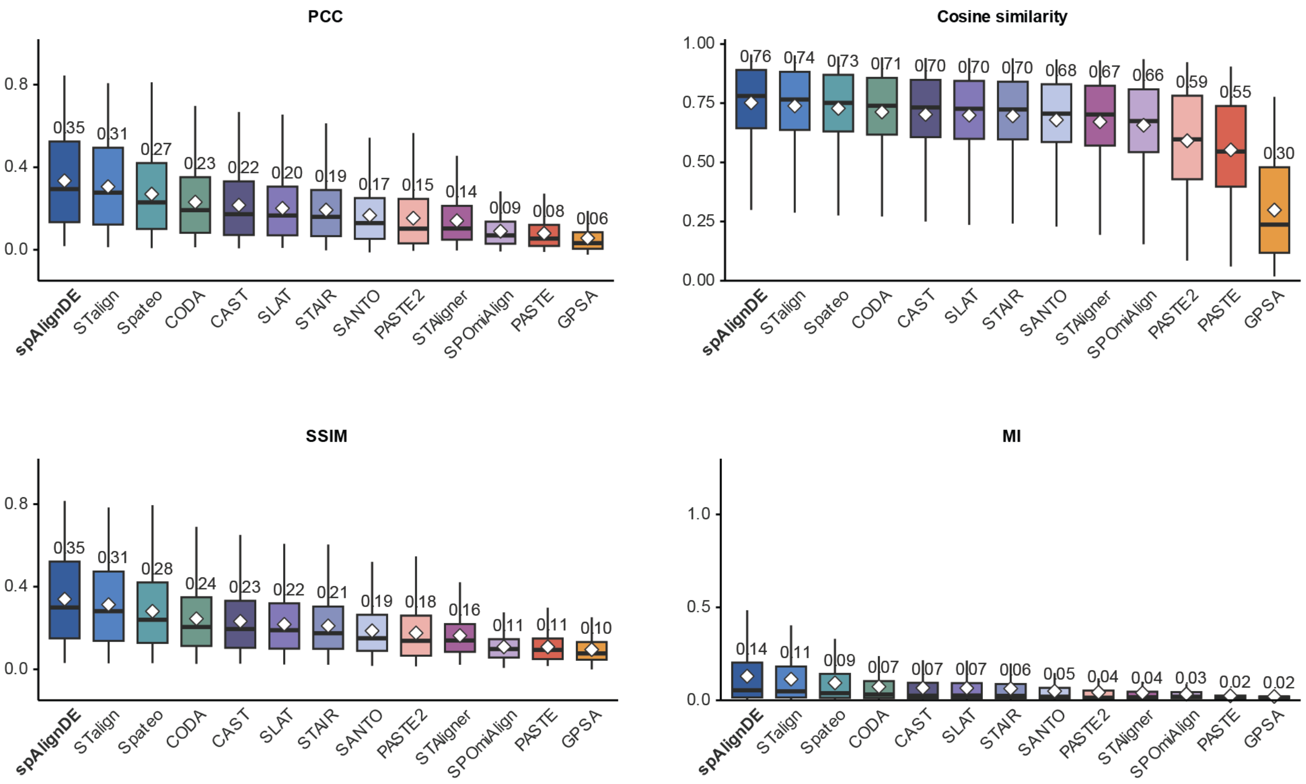


Fig. S12: **Quantitative benchmarking of cross-sample alignment in the aging mouse brain MERFISH cohort using a 50×50 evaluation grid.** All 19 non-reference sections were aligned to the 4.3-month reference section. Post-alignment gene-pattern concordance was evaluated across 300 genes using Pearson correlation coefficient (PCC), cosine similarity, structural similarity index measure (SSIM) and mutual information (MI). For each gene and query section, expression patterns were aggregated on a $50 \times 50$ spatial grid and compared with the 4.3-month reference; each metric was then averaged across the available query-to-reference alignments for that gene. The aligned coordinates were unchanged from the primary $10 \times 10$ evaluation; only the spatial grid used to aggregate expression and calculate concordance was refined. Boxes show interquartile ranges across genes, center lines indicate medians, whiskers extend to 1.5× the interquartile range and white diamonds indicate method-level means; outliers are not shown. Numeric labels above the boxes report the corresponding means. All 13 methods were evaluated across all 19 query-to-reference alignments and were ordered independently by decreasing mean within each metric. Higher values indicate stronger post-alignment spatial expression concordance. spAlignDE achieved the highest mean value for all four metrics. GPSA used 10,000 cells per section, PASTE and PASTE2 used 30,000 cells per section because of computational constraints, and the remaining methods used all available cells.

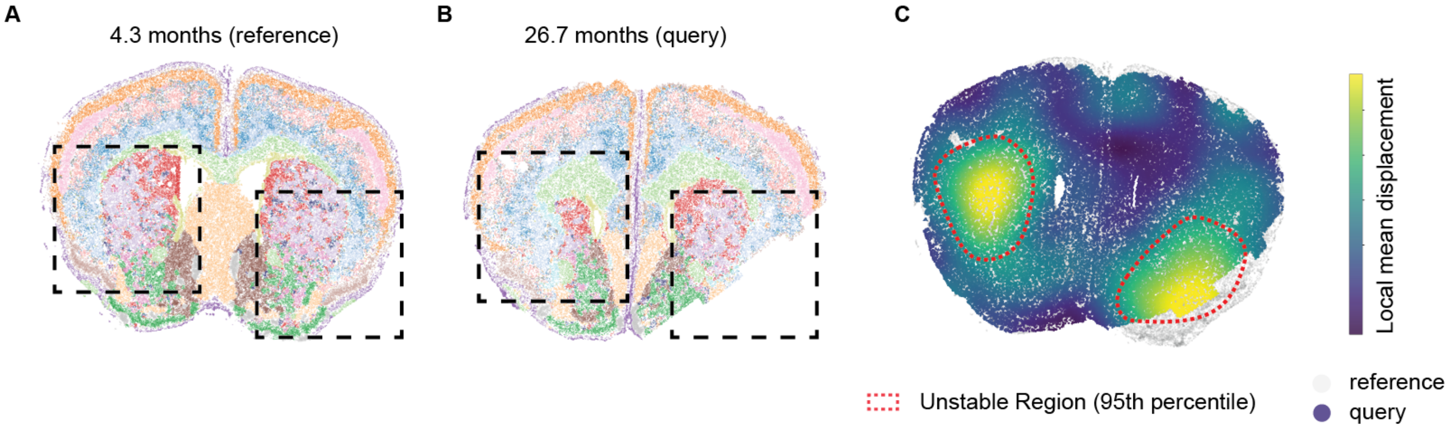


Fig. S13: **Subsampling-based local transformation stability in the aging mouse brain. A,B,** Joint spatial-structure assignments for the 4.3-month reference **A** and 26.7-month query **B**; black dashed boxes highlight local structural discordance. **C,** Mean transformation displacement after aligning the 26.7-month query to the 4.3-month reference. Red dashed contours mark locations above the 95th percentile. Elevated mean displacement coincides with the spatial-structure discordance in **A,B** and the missing-tissue boundary in the query.

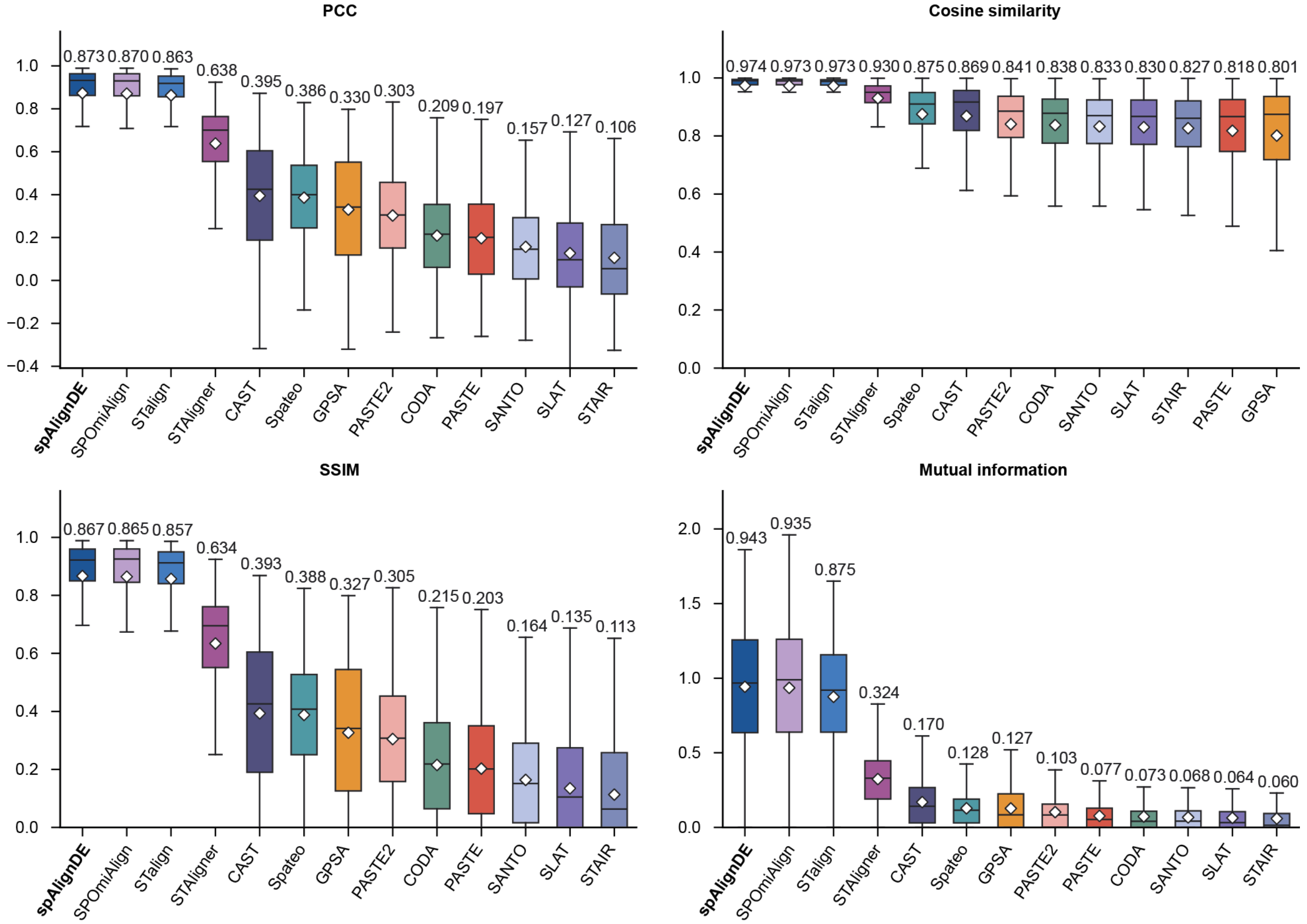


Fig. S14: **Gene-expression concordance after alignment of Xenium breast-cancer replicates using a** $10 \times 10$ **evaluation grid.** Xenium In Situ Sample 1, Replicate 2 (117,630 QC-passing cells) was aligned as the query to Replicate 1 (161,995 QC-passing cells) as the fixed reference. All 313 Gene Expression genes were retained without highly variable gene selection and were normalized using log-transformed counts per 10,000 before evaluation. From upper left to lower right, the four boxplots show the gene-level distributions of Pearson correlation coefficient (PCC), cosine similarity, structural similarity index measure (SSIM) and mutual information (MI). Boxes indicate the interquartile range, center lines indicate medians, whiskers extend to the most extreme values within 1.5× the interquartile range, and white diamonds indicate means; outliers are not shown. Numeric labels above the boxes report method-level means. spAlignDE is fixed at the left of each panel, and the remaining methods are ordered independently by decreasing mean performance. Higher values indicate greater post-alignment spatial expression concordance. PASTE and PASTE2 used 30,000 cells per replicate, GPSA used 10,000 cells per replicate, and all other methods used all QC-passing cells.

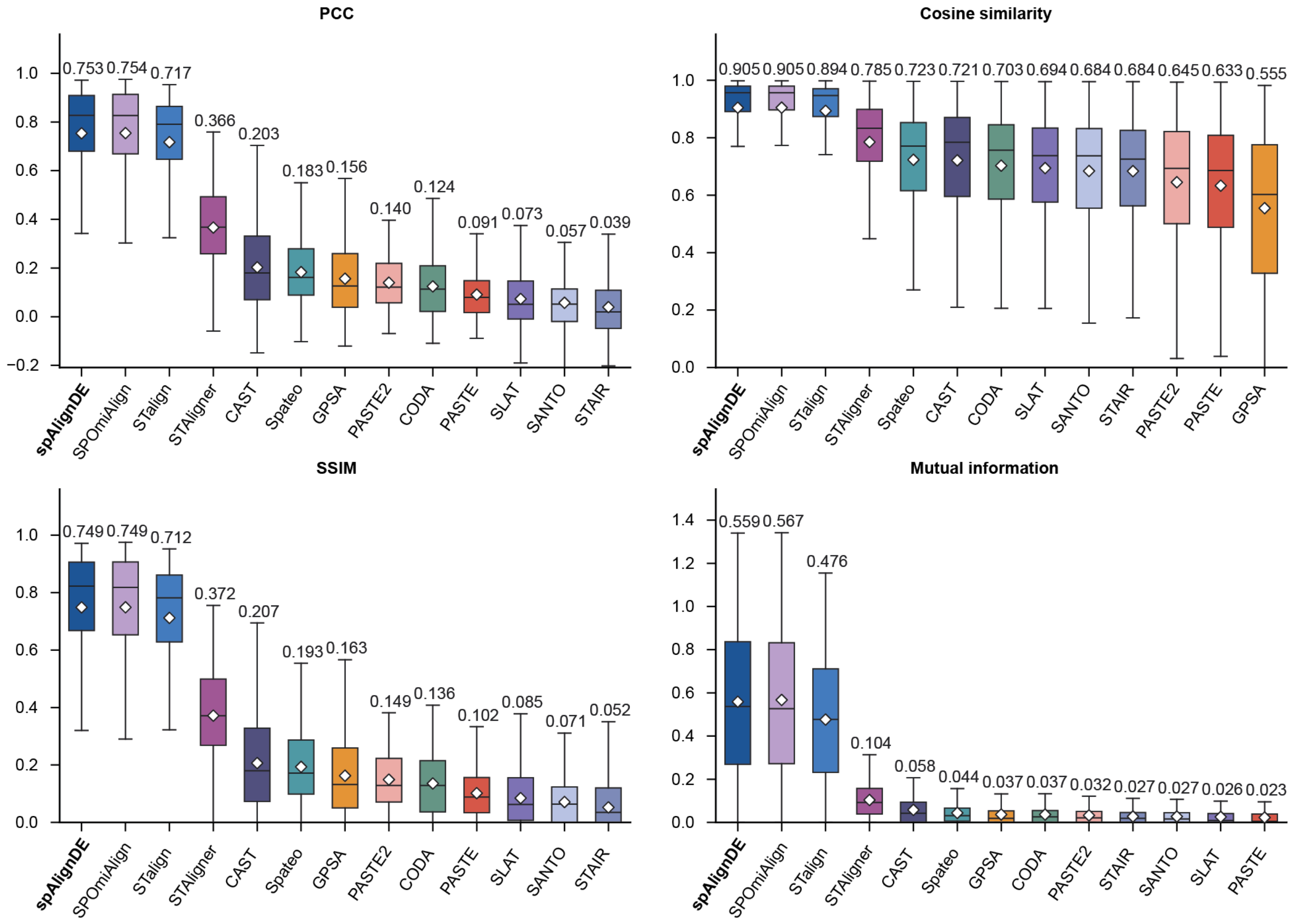


Fig. S15: **Gene-expression concordance after alignment of Xenium breast-cancer replicates using a** $30 \times 30$ **evaluation grid.** Xenium In Situ Sample 1, Replicate 2 (117,630 QC-passing cells) was aligned as the query to Replicate 1 (161,995 QC-passing cells) as the fixed reference. All 313 Gene Expression genes were retained without highly variable gene selection and were normalized using log-transformed counts per 10,000 before evaluation. From upper left to lower right, the four boxplots show the gene-level distributions of Pearson correlation coefficient (PCC), cosine similarity, structural similarity index measure (SSIM) and mutual information (MI). Boxes indicate the interquartile range, center lines indicate medians, whiskers extend to the most extreme values within 1.5× the interquartile range, and white diamonds indicate means; outliers are not shown. Numeric labels above the boxes report method-level means. spAlignDE is fixed at the left of each panel, and the remaining methods are ordered independently by decreasing mean performance. Higher values indicate greater post-alignment spatial expression concordance. PASTE and PASTE2 used 30,000 cells per replicate, GPSA used 10,000 cells per replicate, and all other methods used all QC-passing cells.

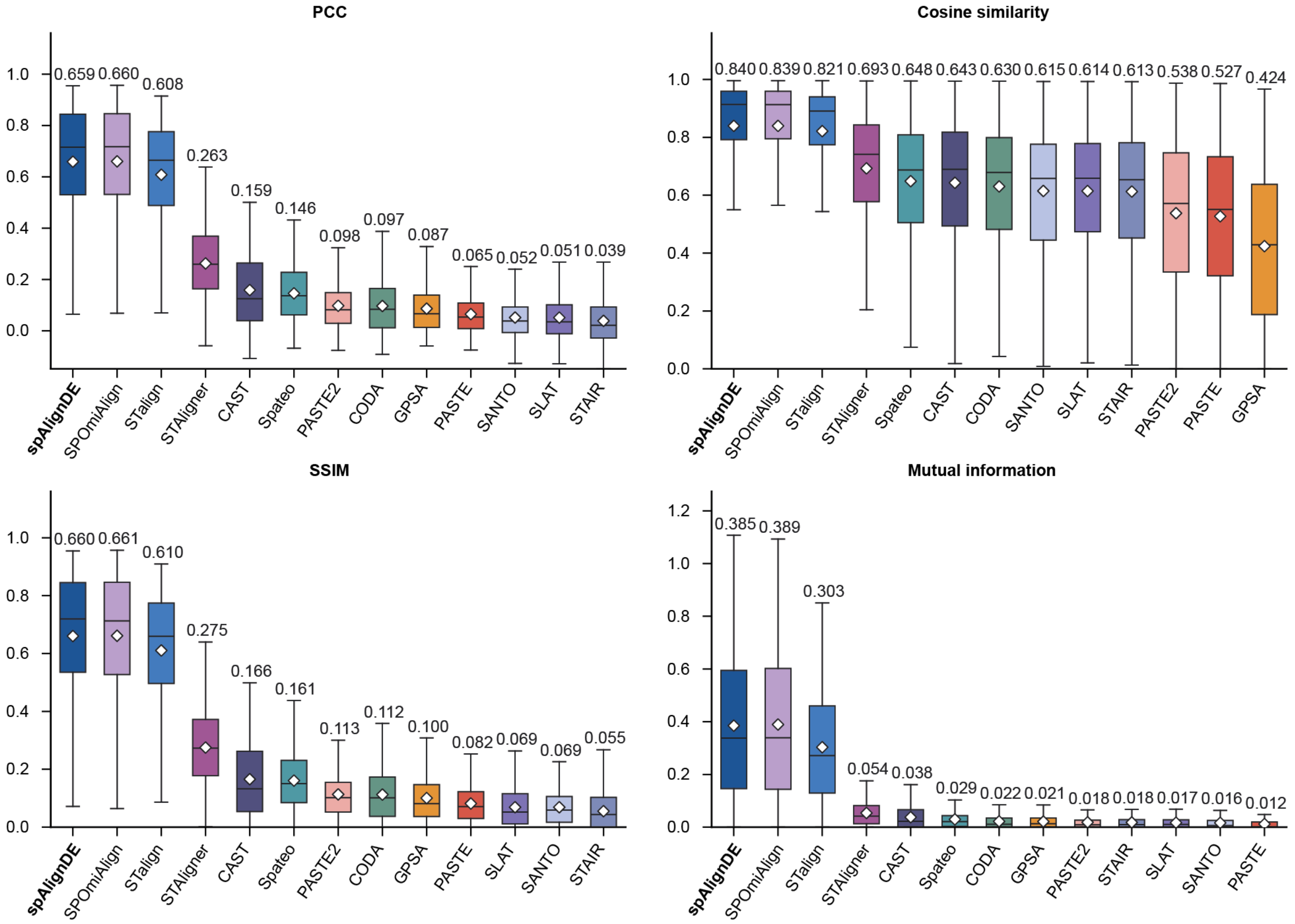


Fig. S16: **Gene-expression concordance after alignment of Xenium breast-cancer replicates using a** $50 \times 50$ **evaluation grid.** Xenium In Situ Sample 1, Replicate 2 (117,630 QC-passing cells) was aligned as the query to Replicate 1 (161,995 QC-passing cells) as the fixed reference. All 313 Gene Expression genes were retained without highly variable gene selection and were normalized using log-transformed counts per 10,000 before evaluation. From upper left to lower right, the four boxplots show the gene-level distributions of Pearson correlation coefficient (PCC), cosine similarity, structural similarity index measure (SSIM) and mutual information (MI). Boxes indicate the interquartile range, center lines indicate medians, whiskers extend to the most extreme values within 1.5× the interquartile range, and white diamonds indicate means; outliers are not shown. Numeric labels above the boxes report method-level means. spAlignDE is fixed at the left of each panel, and the remaining methods are ordered independently by decreasing mean performance. Higher values indicate greater post-alignment spatial expression concordance. PASTE and PASTE2 used 30,000 cells per replicate, GPSA used 10,000 cells per replicate, and all other methods used all QC-passing cells.

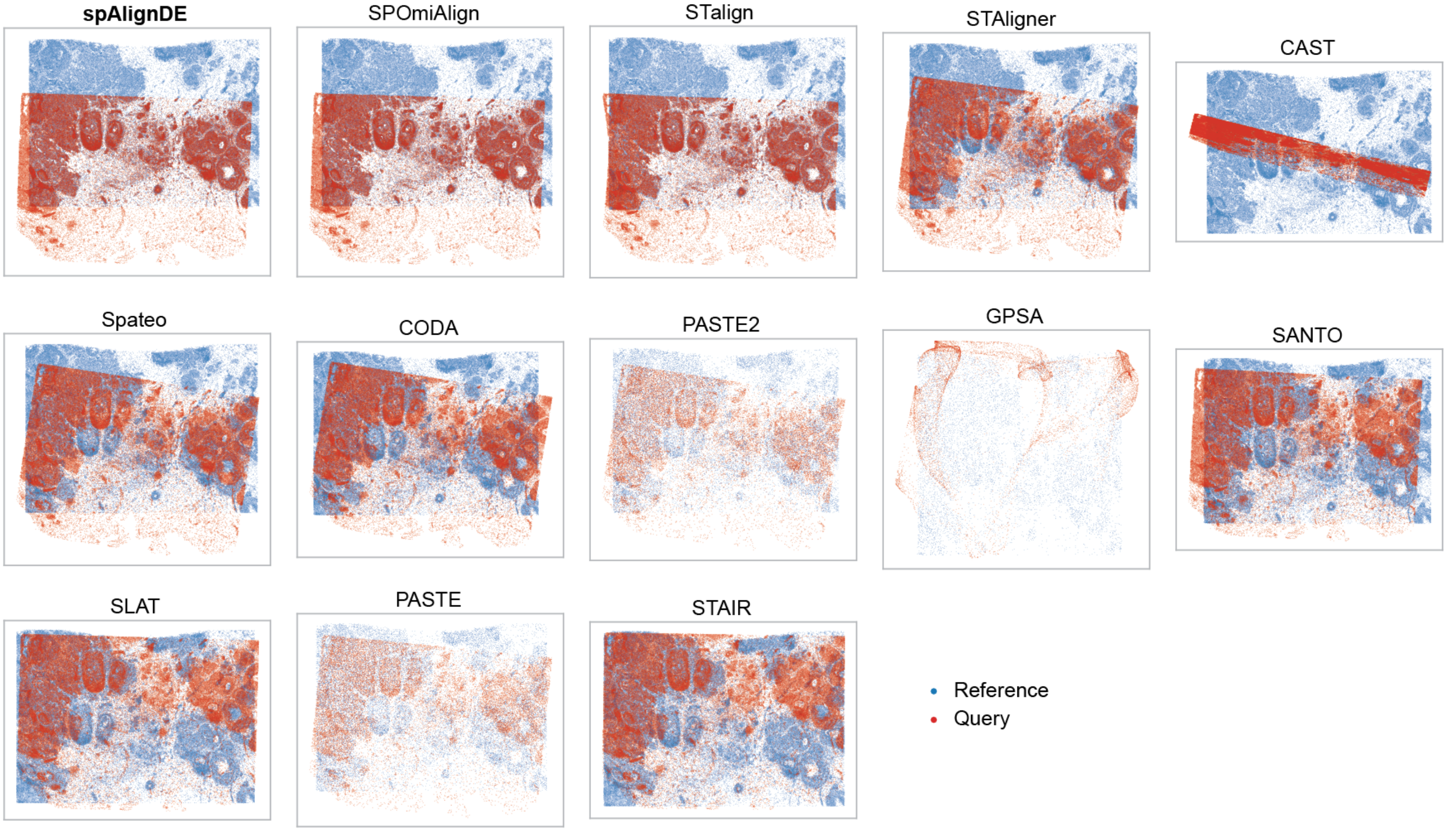


Fig. S17: **After-alignment overlap of Xenium breast-cancer replicates across 13 methods.** Xenium In Situ Sample 1, Replicate 2 was aligned as the query to Replicate 1 as the fixed reference. Reference cells are shown in blue and aligned query cells in red. Red–blue co-localization indicates global spatial correspondence, whereas regions dominated by one color indicate incomplete shared tissue coverage or residual alignment mismatch. spAlignDE is fixed at the left, and the remaining methods are ordered by their overall mean rank across PCC, cosine similarity, SSIM and MI evaluated using the $10 \times 10$, $30 \times 30$ and $50 \times 50$ grids. PASTE and PASTE2 used 30,000 cells per replicate, GPSA used 10,000 cells per replicate, and all other methods used all QC-passing cells.

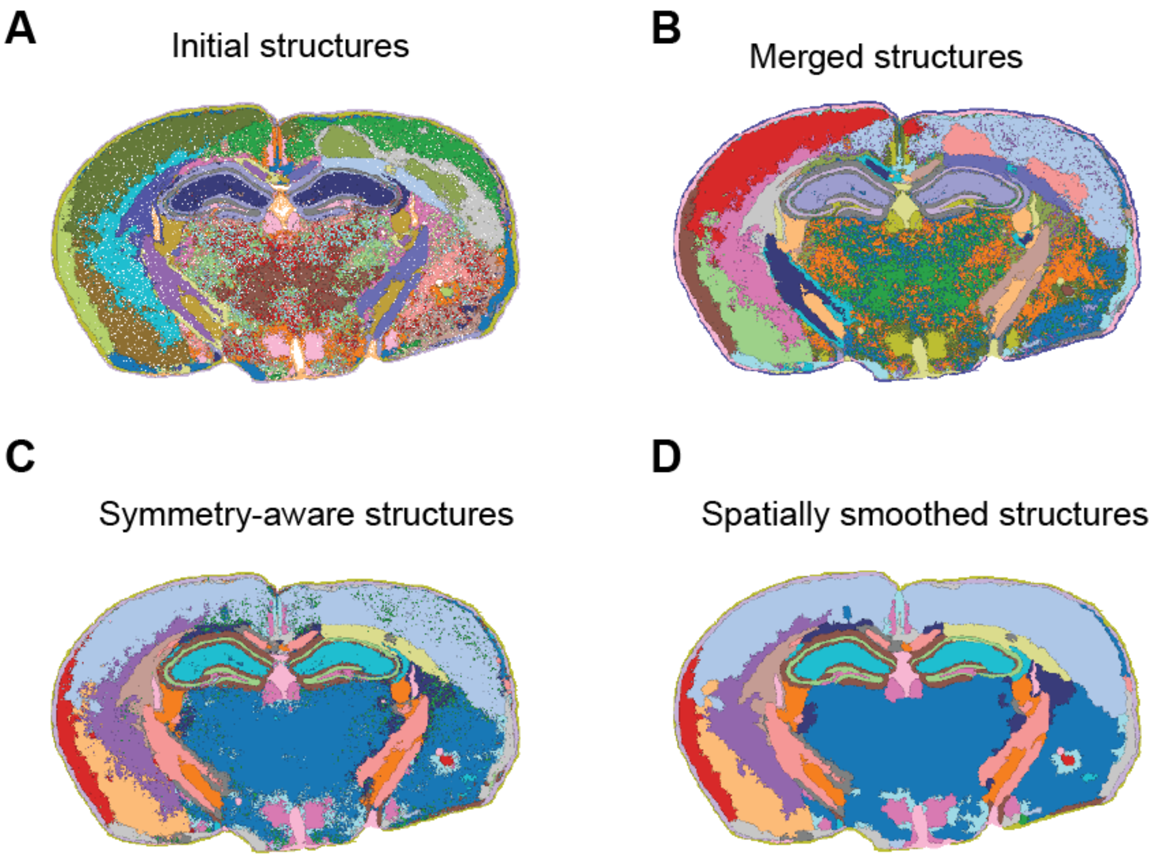


Fig. S18: **Construction of histology-derived spatial structures from the H&E image. A,** Initial H&E image-feature assignments generated on the downsampled embedding grid after tissue–background separation. Histology-derived embedding channels were combined with lightly weighted RGB intensities and spatial coordinates and partitioned into 30 initial regions ($k_{\text{slide}} = 30$, $w_{\text{RGB}} = 0.25$ and $w_{\text{xy}} = 0.05$). **B,** Ward agglomeration consolidated the initial assignments to 26 meta-regions, after which reflected-mask pairing across the predefined bilateral anatomical symmetry axis merged two mutually selected region pairs and yielded 24 regions. **C,** Three additional symmetry-supported merges passed the minimum global bilateral-overlap gain of 0.02, with score gains of 0.0357, 0.0492 and 0.0300, reducing the partition from 24 to 21 regions. **D,** Spatial cleaning removed small disconnected islands, reassigned locally inconsistent regions and filled remaining unlabeled gaps while retaining 21 represented regions. The resulting 21-region H&E structure map was used for cross-modality structure pairing and S-LDDMM alignment. Background pixels were excluded, and colors denote histology-derived spatial structures.

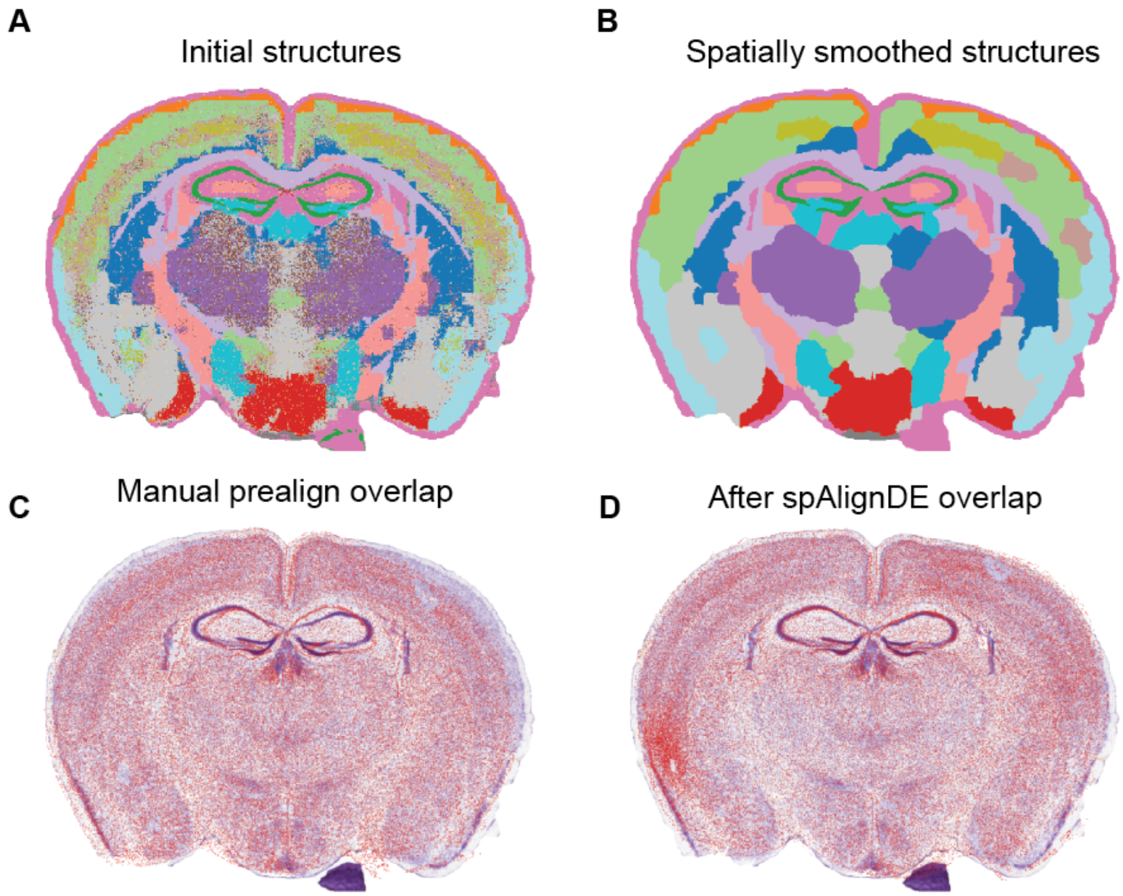


Fig. S19: **Histology-derived spatial structures and alignment of MERFISH data to Nissl histology. A,** Initial fine-grained histology-feature assignments derived from a Nissl-stained coronal section of the Allen Mouse Brain Atlas. **B,** Spatially smoothed assignments defining the coherent histology-derived structures used for alignment. **C,** Overlay of cells from the MERFISH Mouse Brain Receptor Map S2R1 section and the Nissl image after manual global pre-alignment. **D,** Overlay after S-LDDMM refinement, showing improved visual correspondence with the tissue boundary and internal cytoarchitectural structures of the Nissl section. MERFISH cells are shown in red.

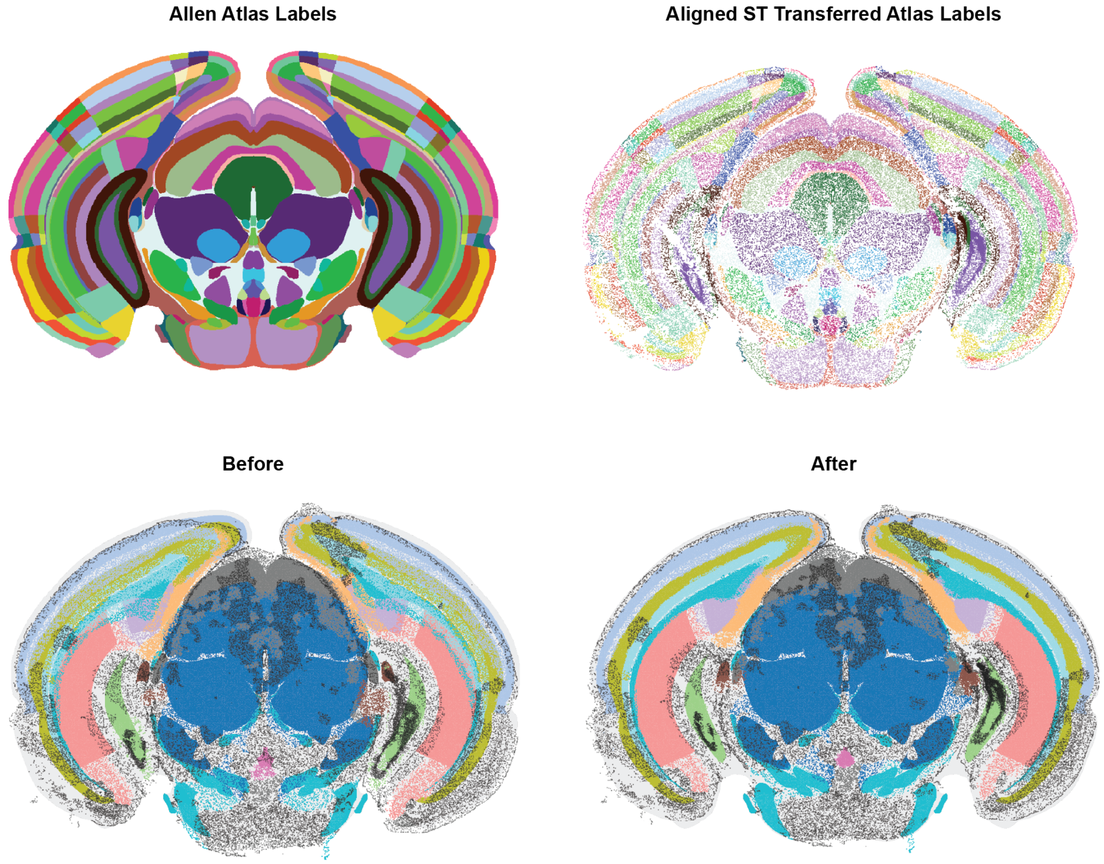


Fig. S20: **Alignment of the MERFISH S1R1 section to the Allen CCFv3 reference.** The fixed-seed automatic pipeline retained 12 final matched ST–atlas structure pairs. Top, anatomical labels from the anatomically corresponding Allen CCFv3 coronal section and their transfer to the MERFISH cells after alignment with spAlignDE. Bottom, automatically identified spatial-structure correspondences before and after alignment. Corresponding MERFISH spatial structures and atlas regions are shown using matched colors.

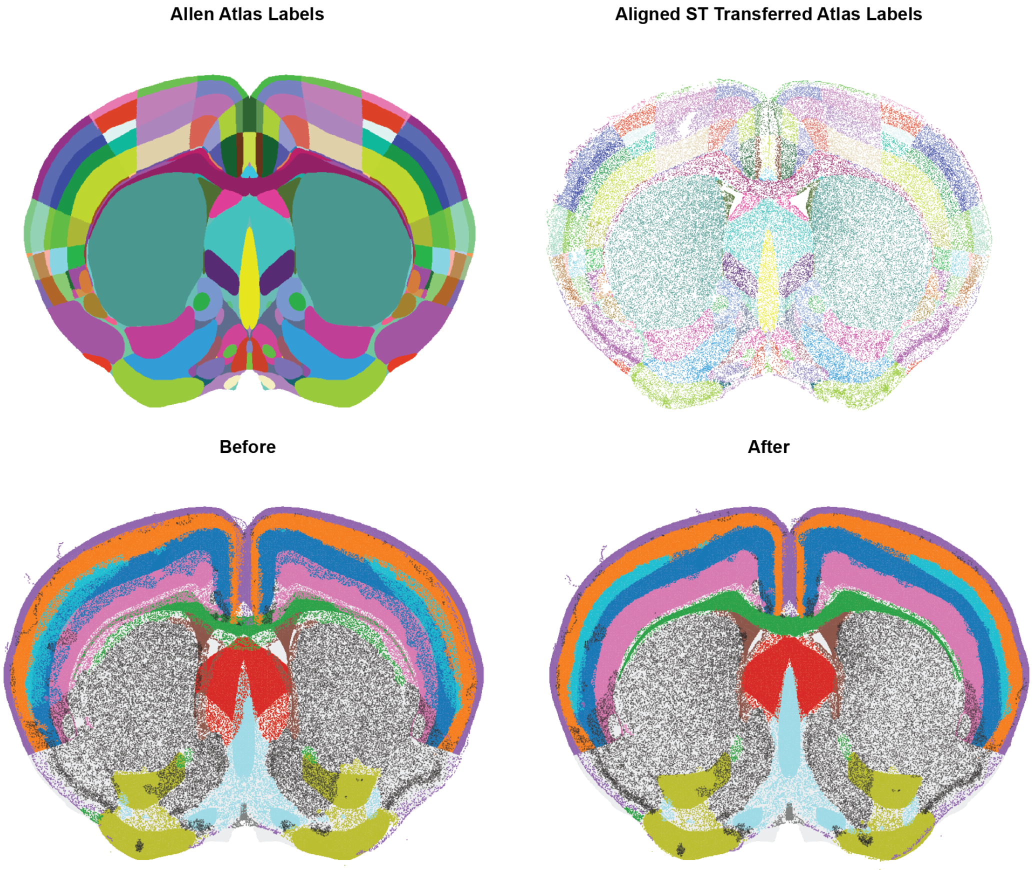


Fig. S21: **Alignment of the MERFISH S3R1 section to the Allen CCFv3 reference.** The fixed-seed automatic pipeline retained 11 final matched ST–atlas structure pairs. Top, anatomical labels from the anatomically corresponding Allen CCFv3 coronal section and their transfer to the MERFISH cells after alignment with spAlignDE. Bottom, automatically identified spatial-structure correspondences before and after alignment. Corresponding MERFISH spatial structures and atlas regions are shown using matched colors.

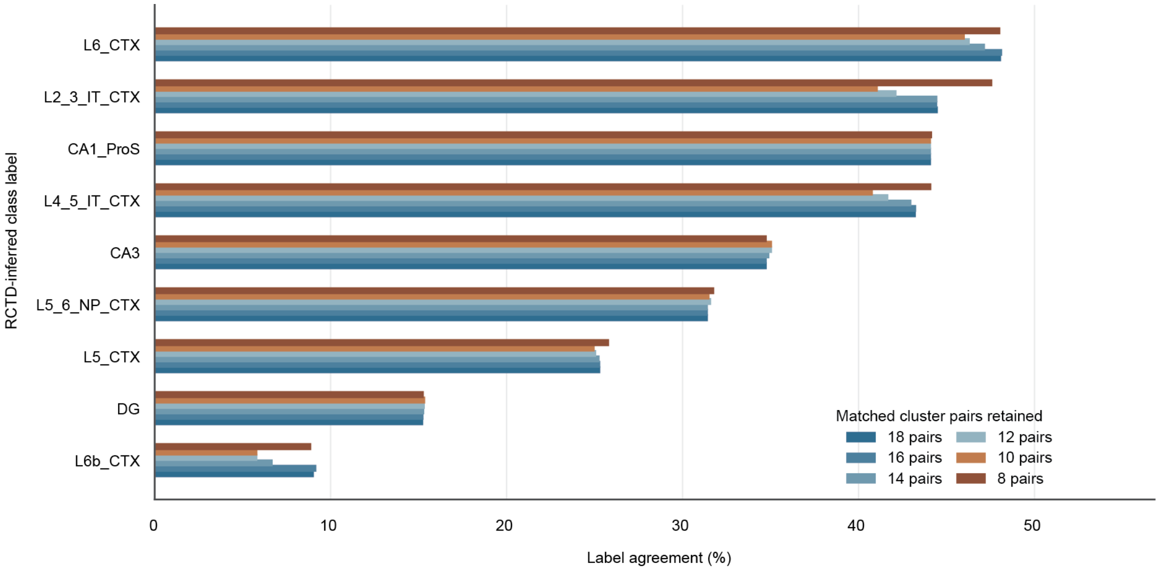


Fig. S22: **Label agreement across RCTD-inferred class labels is robust to reductions in the number of retained matched spatial-structure pairs.** The completed fixed-seed automatic ST-to-atlas pipeline retained 18 matched spatial-structure pairs. For the controlled sensitivity analysis, these pairs were ranked once and the highest-ranking 18, 16, 14, 12, 10 or 8 pairs were retained. For every condition, the corresponding signed-distance-transform channels were reconstructed and one final-resolution S-LDDMM alignment was performed for 200 optimization iterations from the same fixed global pre-alignment. Structure-pair discovery was not repeated, and the ST dataset, Allen CCFv3 reference section and all other S-LDDMM parameters were held fixed. The one-step 18-pair rerun served as the full-pair sensitivity baseline; it is distinct from the completed multistage 18-pair primary alignment. Bars show label agreement (%) for each RCTD-inferred class label, defined as the percentage of eligible ST cells whose transferred Allen CCFv3 label belonged to one of the predefined anatomical categories compatible with that class label. Colors indicate the number of matched spatial-structure pairs retained.

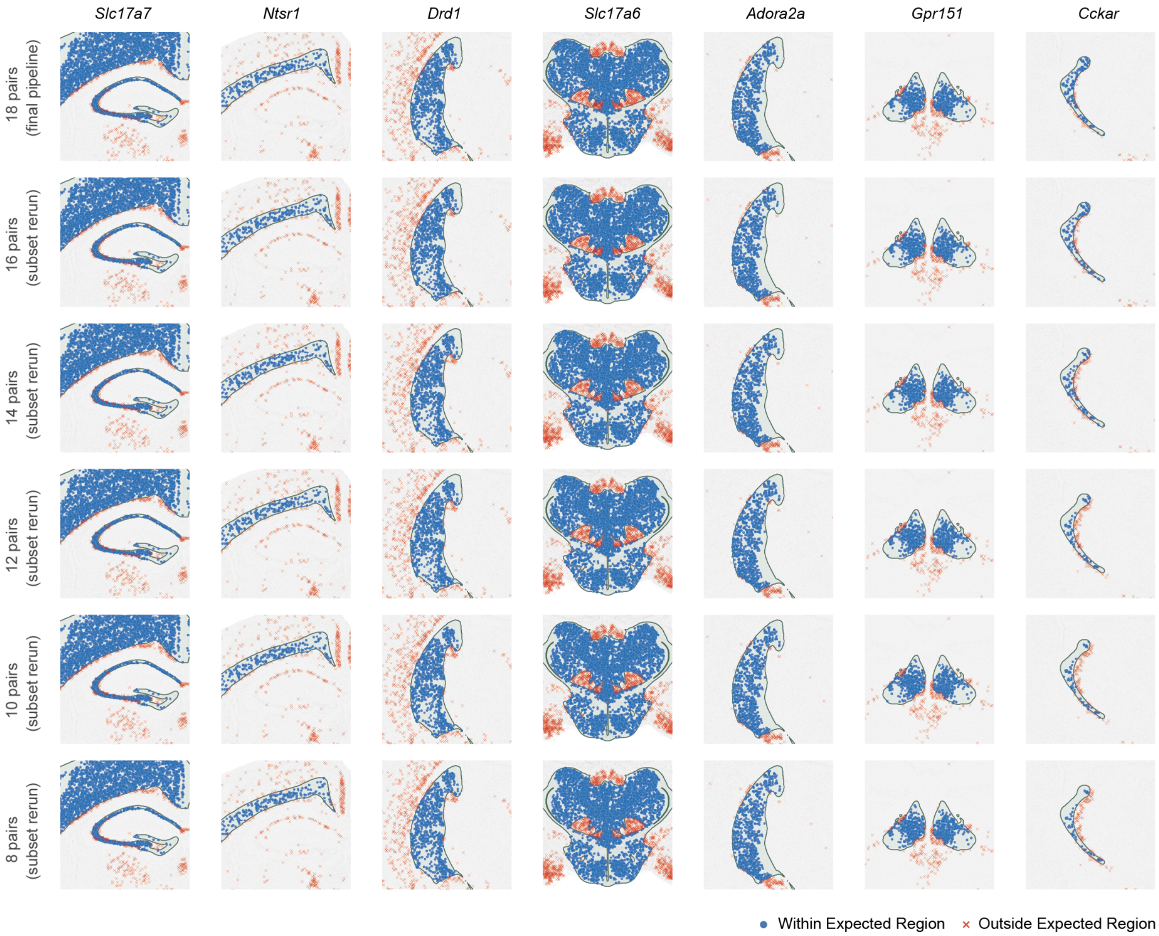


Fig. S23: **Regionally enriched marker-gene localization is robust to reductions in the number of retained matched spatial-structure pairs.** Starting from the 18 matched spatial-structure pairs retained by the fixed-seed automatic pairing procedure, the pairs were ranked once by the final gated alignment score, with the ungated alignment score used to resolve remaining ties. The highest-ranking 18, 16, 14, 12, 10 or 8 pairs were retained. For each condition, including the 18-pair condition, the corresponding signed-distance-transform channels were reconstructed and an independent one-step final-resolution S-LDDMM alignment was run for 200 optimization iterations from the same fixed global pre-alignment. Structure-pair discovery was not repeated, and the ST dataset, Allen CCFv3 reference section and all other S-LDDMM parameters were held fixed. The one-step 18-pair rerun served as the full-pair sensitivity baseline and is distinct from the completed multistage 18-pair alignment used as the primary atlas result. Columns show seven regionally enriched marker genes, and rows indicate the number of matched spatial-structure pairs retained. For each gene, high-expression cells were defined as cells with positive raw counts at or above the 75th percentile among cells with positive expression. Blue points indicate high-expression cells whose transferred Allen CCFv3 labels fell within the predefined expected anatomical region, red crosses indicate high-expression cells transferred outside that region and black outlines delineate the expected anatomical regions.

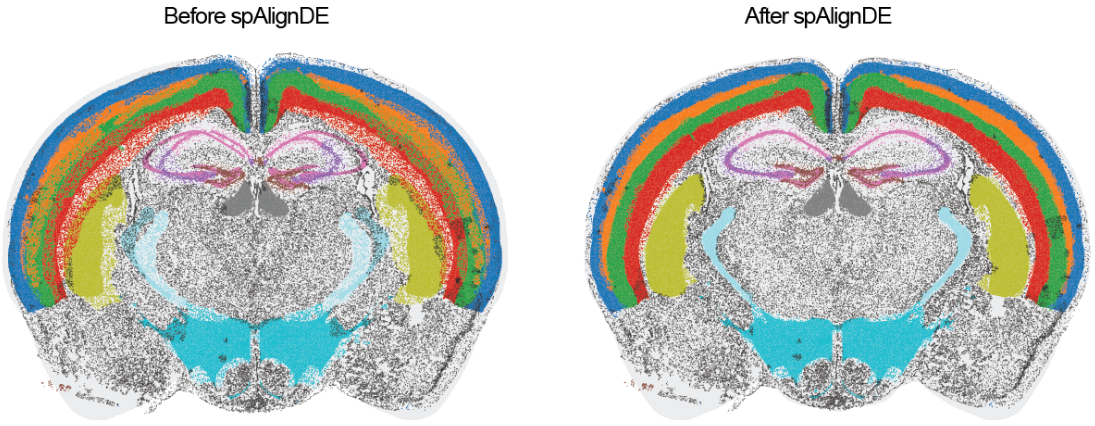


Fig. S24: **Atlas alignment guided by UI-defined spatial-structure pairs.** Nine correspondence groups between spatial transcriptomic structures and Allen CCFv3 atlas regions were UI-defined using the spAlignDE Structure Pair user interface. Spatial transcriptomics coordinates are shown before and after high-resolution S-LDDMM alignment using these UI-defined structure pairs. Corresponding ST structures and atlas regions within each paired group share a color, whereas structures not included in the selected correspondence groups are shown in gray.

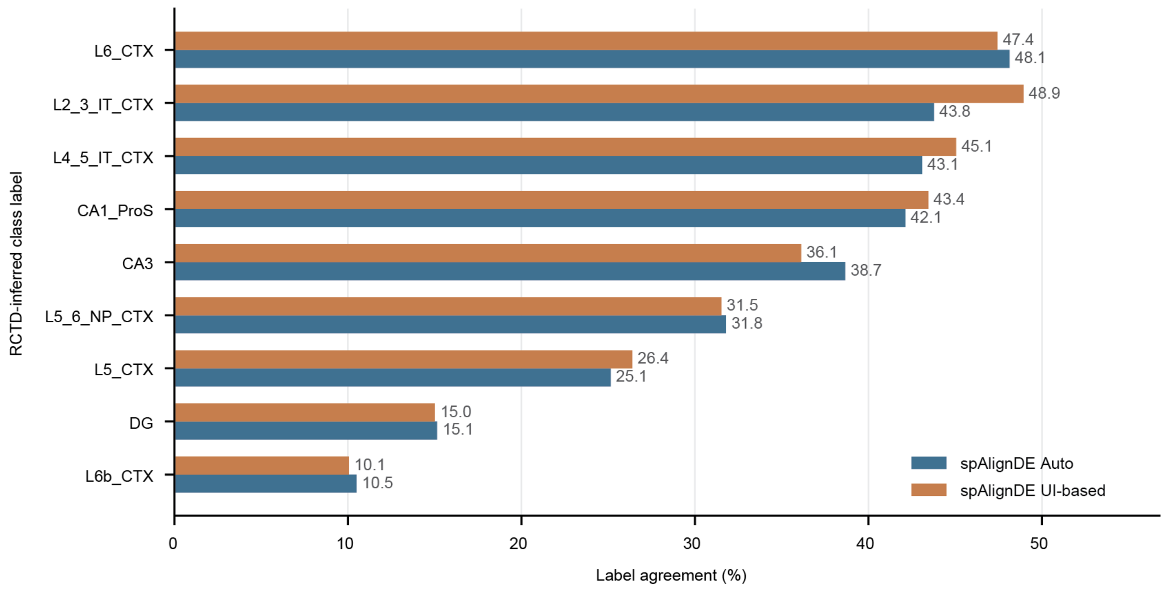


Fig. S25: **Label agreement across RCTD-inferred class labels after automatic or UI-defined spatial-structure pairing.** The same spatial transcriptomics section was aligned to the Allen CCFv3 reference using either 18 matched spatial-structure pairs retained by the fully automatic spAlignDE pipeline or nine UI-defined correspondence groups constructed with the spAlignDE Structure Pair user interface. Bars show label agreement (%) for each RCTD-inferred class label after automatic-pairing alignment (blue) or UI-defined-pairing alignment (orange). For each class, label agreement was calculated post hoc as the percentage of eligible ST cells whose transferred Allen CCFv3 label belonged to one of the predefined anatomical categories compatible with that class label. RCTD-inferred labels were used only for evaluation and were not used to construct the structure correspondences or estimate either alignment.

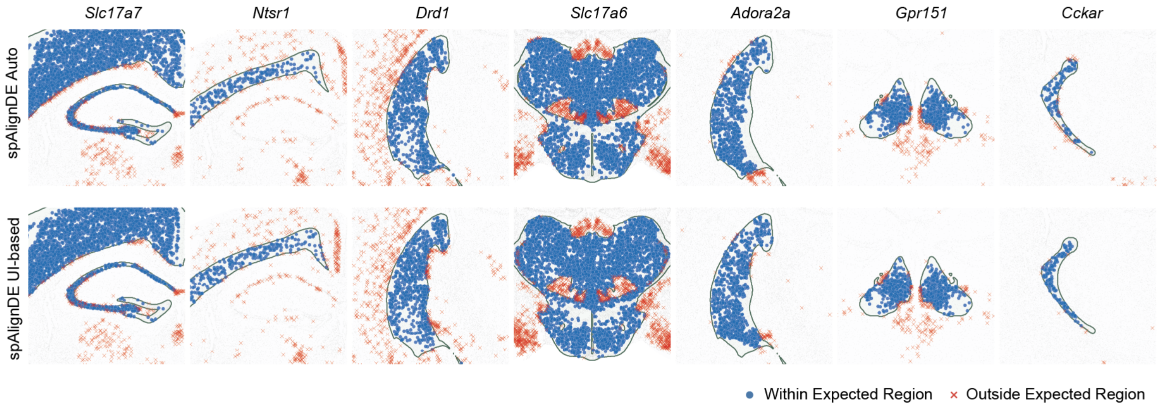


Fig. S26: **Regionally enriched marker-gene localization after automatic or UI-defined spatial-structure pairing.** Spatial localization of seven regionally enriched marker genes is shown after alignment using 18 matched spatial-structure pairs retained by the fully automatic spAlignDE pipeline (top) or nine UI-defined correspondence groups constructed with the spAlignDE Structure Pair user interface (bottom). For each gene, high-expression ST cells were defined as cells with positive raw counts at or above the 75th percentile among cells with positive expression. Blue circles indicate high-expression cells whose transferred Allen CCFv3 labels fell within the predefined expected anatomical region, red crosses indicate high-expression cells transferred outside that region and green outlines delineate the expected atlas regions. Light-gray points show the remaining ST cells.

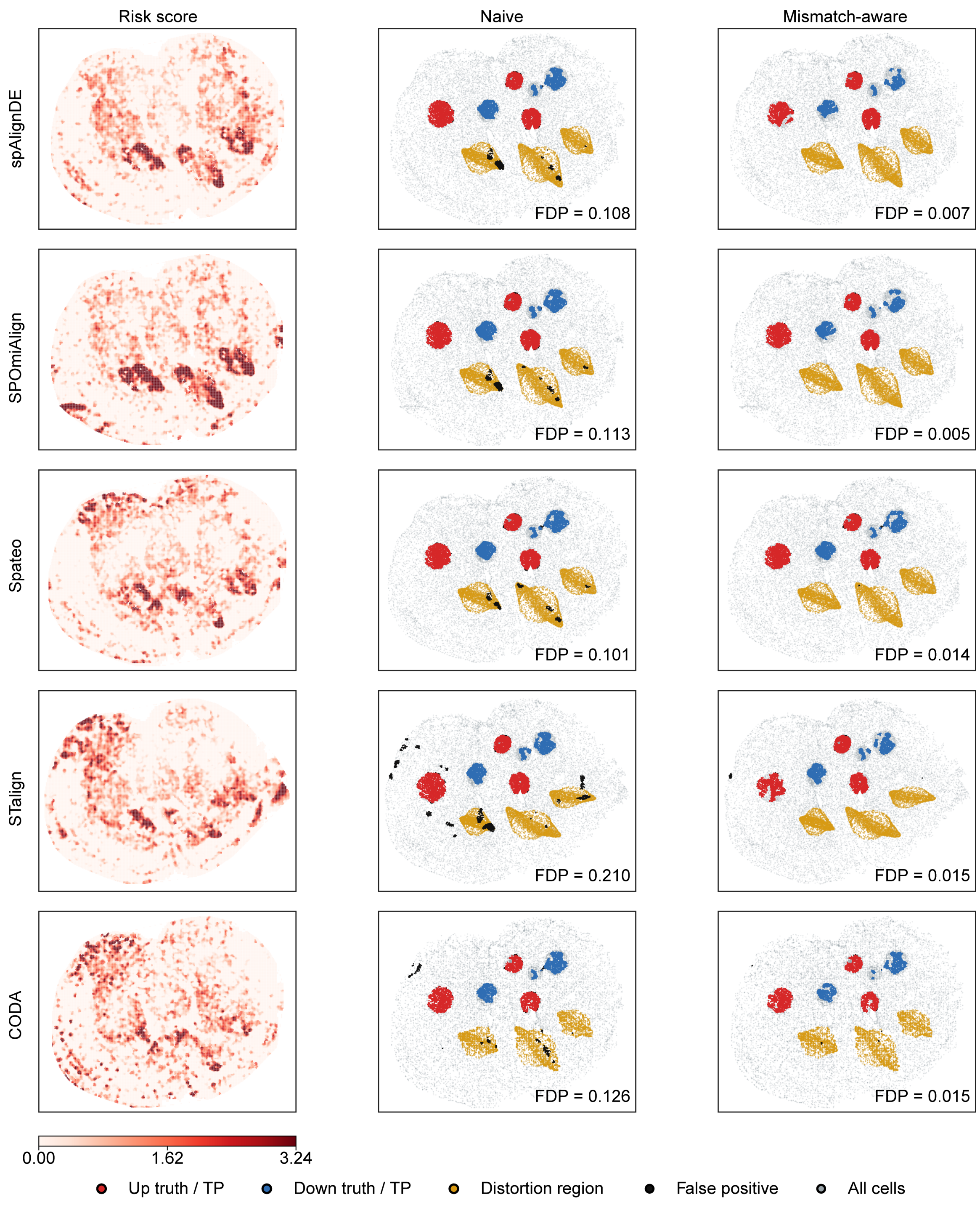


Fig. S27: **Complete spatial testing outputs across alignment methods, part 1.** Non-negative robust-standardized mismatch-risk, naive and mismatch-aware maps are shown for spAlignDE, SPOmiAlign, Spateo, STalign and CODA using the representative simulation in Fig. 4C. The displayed risk scores are evaluated on the shared grid after median–MAD standardization and flooring at zero, but before 95th-percentile capping and rescaling; they therefore have no fixed upper bound. A common color scale is shared with Supplementary Fig. S28. Significant grid calls are projected to sample-B cells. Red and blue points denote correctly signed true-positive calls in up- and down-regulated regions, orange points denote implanted distortion regions, black points denote false-positive calls and gray points denote all cells. Displayed values are location-level FDP for calls made at a target FDR of 0.05.

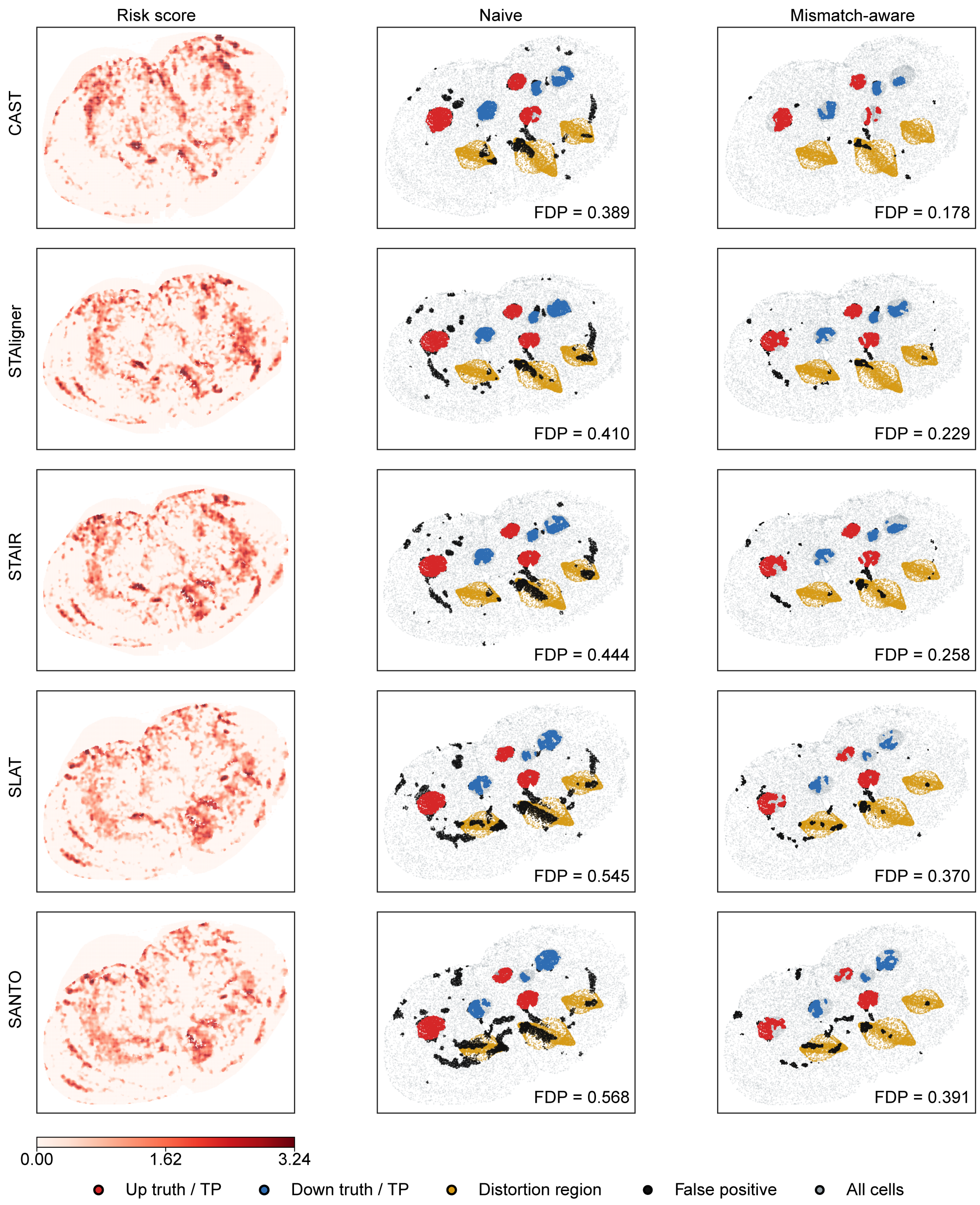


Fig. S28: **Complete spatial testing outputs across alignment methods, part 2.** Non-negative robust-standardized mismatch-risk, naive and mismatch-aware maps are shown for CAST, STAligner, STAIR, SLAT and SANTO using the representative simulation in Fig. 4C. The displayed risk scores are evaluated on the shared grid after median–MAD standardization and flooring at zero, but before 95th-percentile capping and rescaling; they therefore have no fixed upper bound. A common color scale is shared with Supplementary Fig. S27. Significant grid calls are projected to sample-B cells. Red and blue points denote correctly signed true-positive calls in up- and down-regulated regions, orange points denote implanted distortion regions, black points denote false-positive calls and gray points denote all cells. Displayed values are location-level FDP for calls made at a target FDR of 0.05.

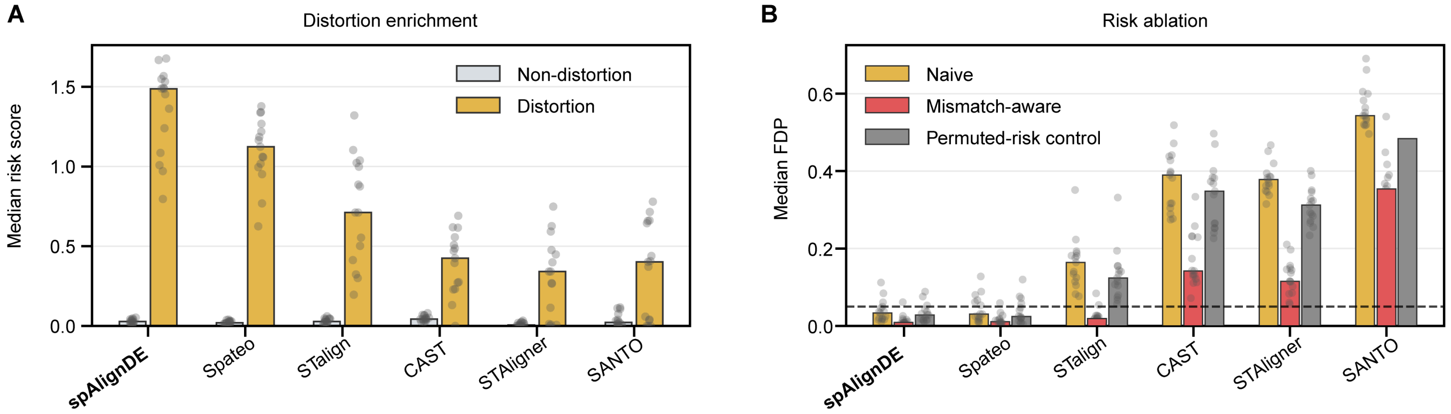


Fig. S29: **Supplementary validation of the alignment mismatch risk score. A,** Distortion enrichment analysis. Implanted synthetic distortion regions were tested for enrichment of high non-negative robust-standardized mismatch-risk scores $\zeta_i^{\mathrm{R}}$. These scores are shown before 95th-percentile capping and rescaling and therefore have no fixed upper bound. The analysis evaluates whether the risk map is spatially informative for the simulated mismatch regions, rather than treating the risk score as a direct estimator of true alignment error. **B,** Risk ablation analysis. Location-level FDP for calls made at a target FDR of 0.05 is compared among naive testing, mismatch-aware testing using the observed spatial risk map, and a permuted-risk control. The permuted-risk control retains the observed risk values but randomly reassigns them across grid locations. Mismatch-aware testing using the observed risk map improves FDP control. The permuted-risk control remains close to the naive analysis. Bars show medians across simulation replicates, and points show individual replicates.

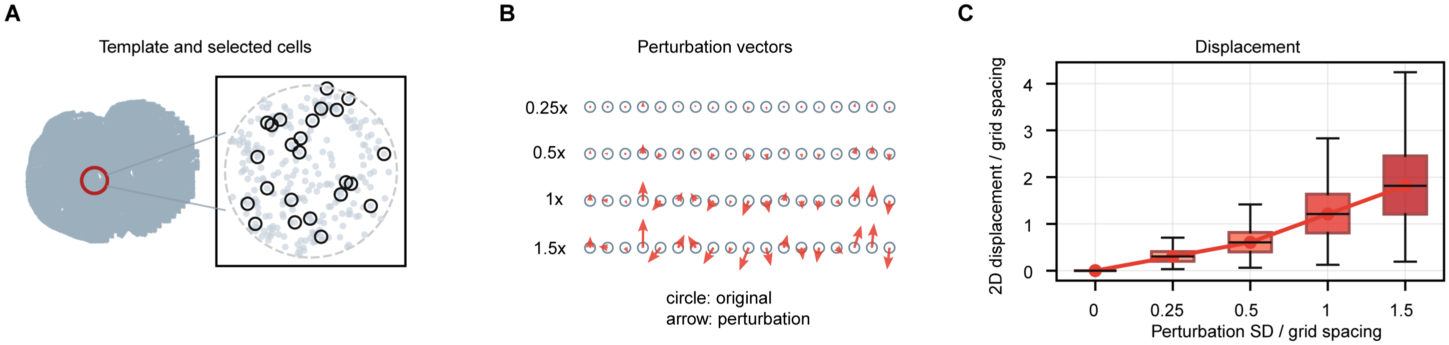


Fig. S30: **Controlled cell-level coordinate perturbation analysis. A,** A local set of cells selected from the spatial simulation template. **B,** Gaussian perturbation vectors at increasing noise levels. At perturbation level $u$, the standard deviation along each coordinate axis is $uh_{\text{grid}}$. **C,** Two-dimensional displacement divided by shared-grid spacing. Noise was added to the oracle sample-B cell coordinates after simulation truth was defined, without rerunning alignment.

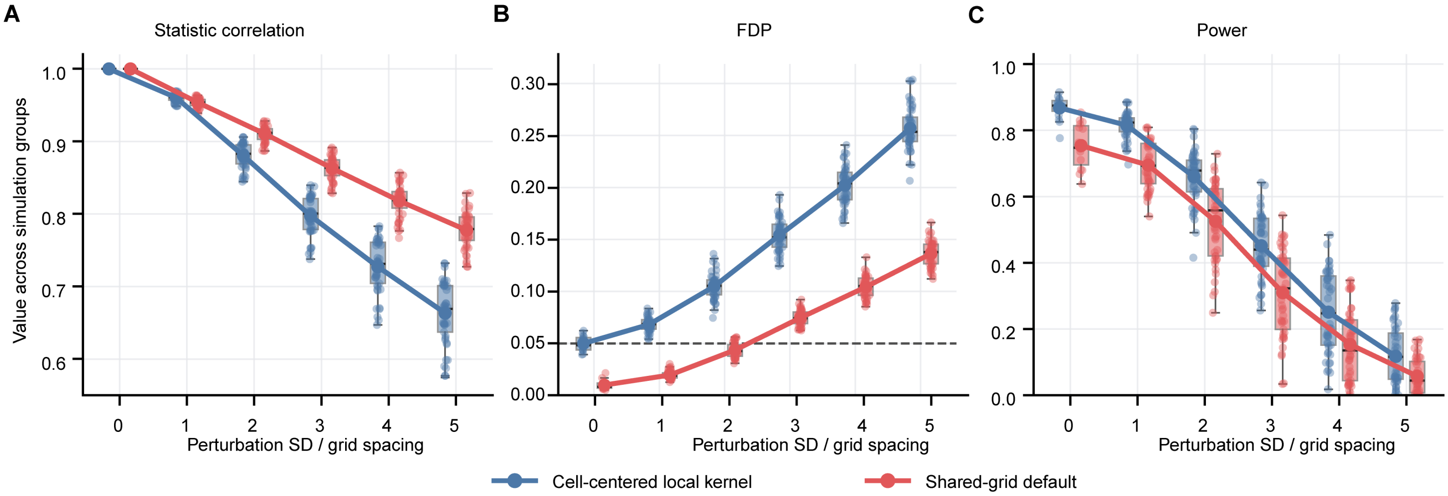


Fig. S31: **Shared-grid local inference under controlled coordinate perturbation.** The shared-grid workflow is compared with a cell-centered local-kernel baseline. **A,** Correlation between perturbed and unperturbed local statistic maps. **B,** Location-level FDP for calls made at a target FDR of 0.05. **C,** Power at FDP $\leq 0.05$. Grid-level results were projected to sample-B cells before comparison with the implanted truth, as defined in Methods. Fixed shared-grid anchors preserve local statistics more stably and slow FDP inflation as coordinate perturbation increases.

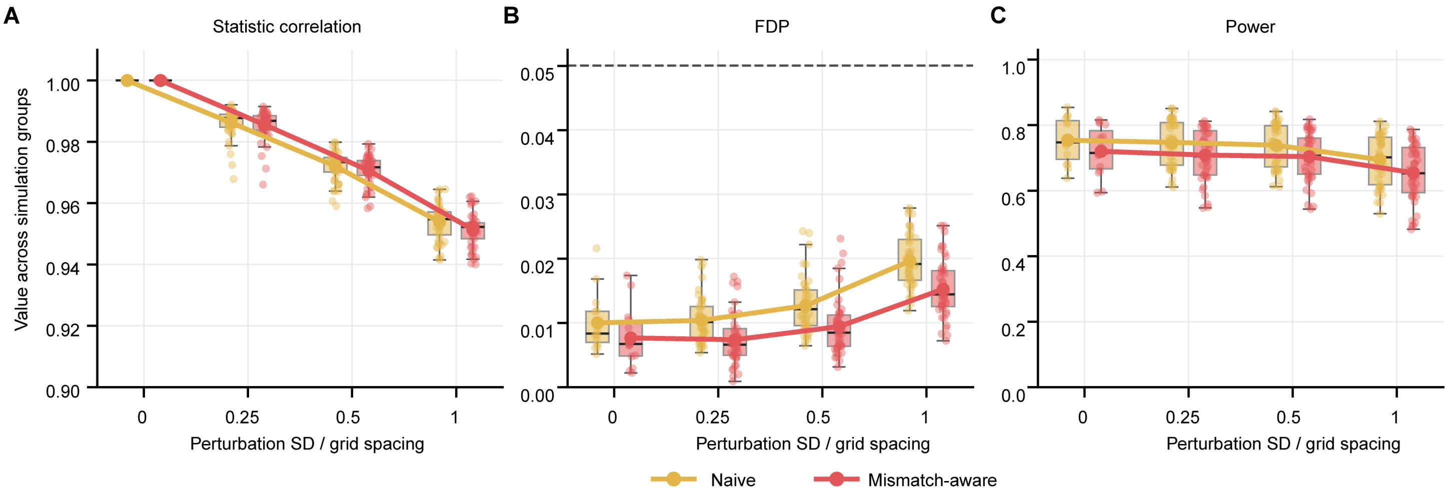


Fig. S32: **Naive and mismatch-aware testing under controlled coordinate perturbation.** Both analyses use the same shared-grid workflow and differ only in whether mismatch-aware variance inflation is applied. **A,** Local statistic map correlation. **B,** Location-level FDP for calls made at a target FDR of 0.05. **C,** Power at FDP $\leq 0.05$. Grid-level results were projected to sample-B cells before comparison with the implanted truth, as defined in Methods. Mismatch-aware testing further reduces FDP under coordinate perturbation, with some loss of power at larger perturbation levels.

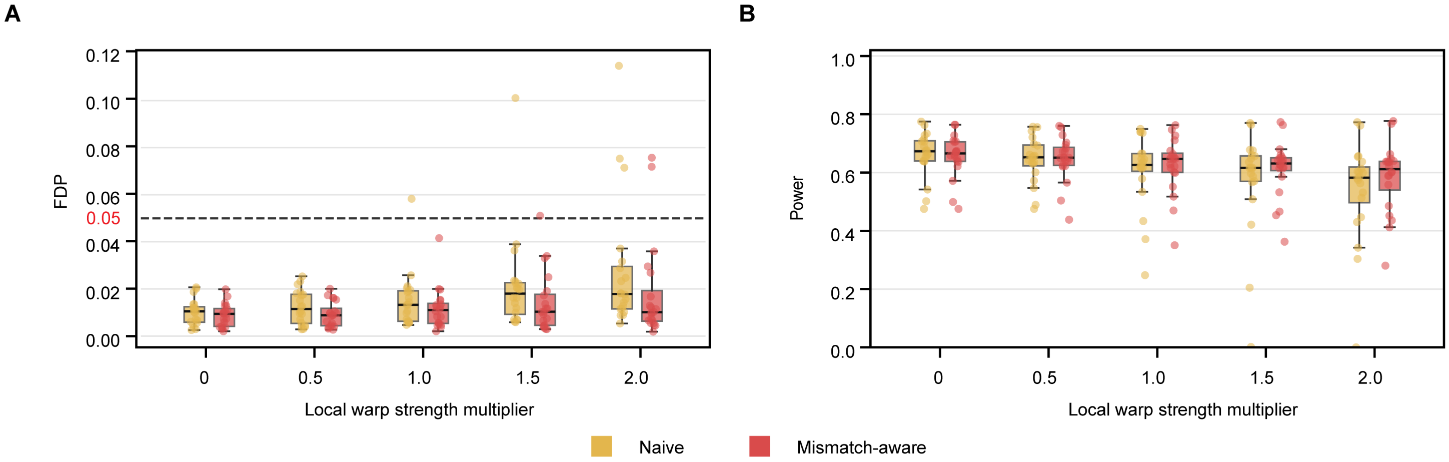


Fig. S33: **Sensitivity to synthetic local warp strength.** Twenty matched simulation templates were evaluated while varying the local warp strength. Naive and mismatch-aware testing were compared using location-level FDP for calls made at a target FDR of 0.05 and power at FDP $\leq 0.05$, as defined in Methods. Mismatch-aware testing shows lower FDP as local deformation increases while retaining similar power under the common FDP constraint.

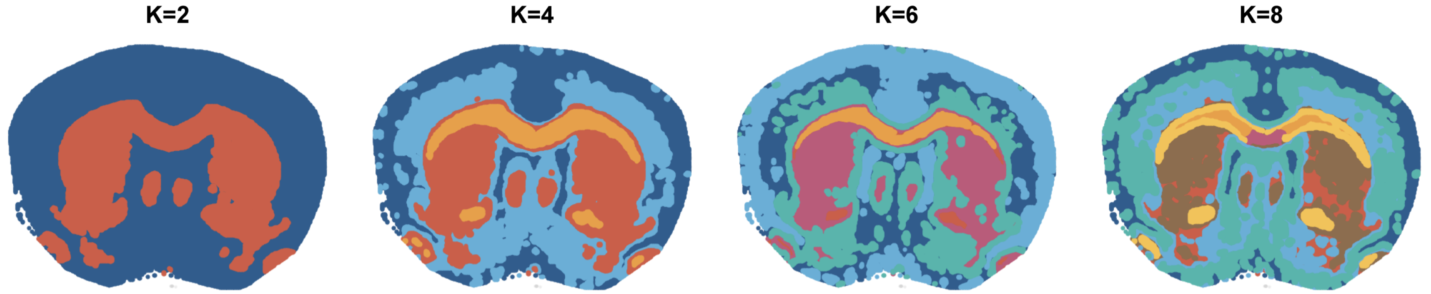


Fig. S34: ***Gamt* spatial trajectory clusters across different cluster numbers.** Spatial trajectory clustering for *Gamt* is shown after manually fixing $K$ at 2, 4, 6 and 8 in the aging brain application. These values were used to examine the effect of cluster resolution and were not outputs of the automatic $K$-selection procedure. Grid locations were clustered by adjusted local expression trajectories across age. Larger $K$ values subdivide broad spatial domains while preserving the main regional organization, supporting the interpretation of the *Gamt* trajectory clusters shown in Fig. 5B.

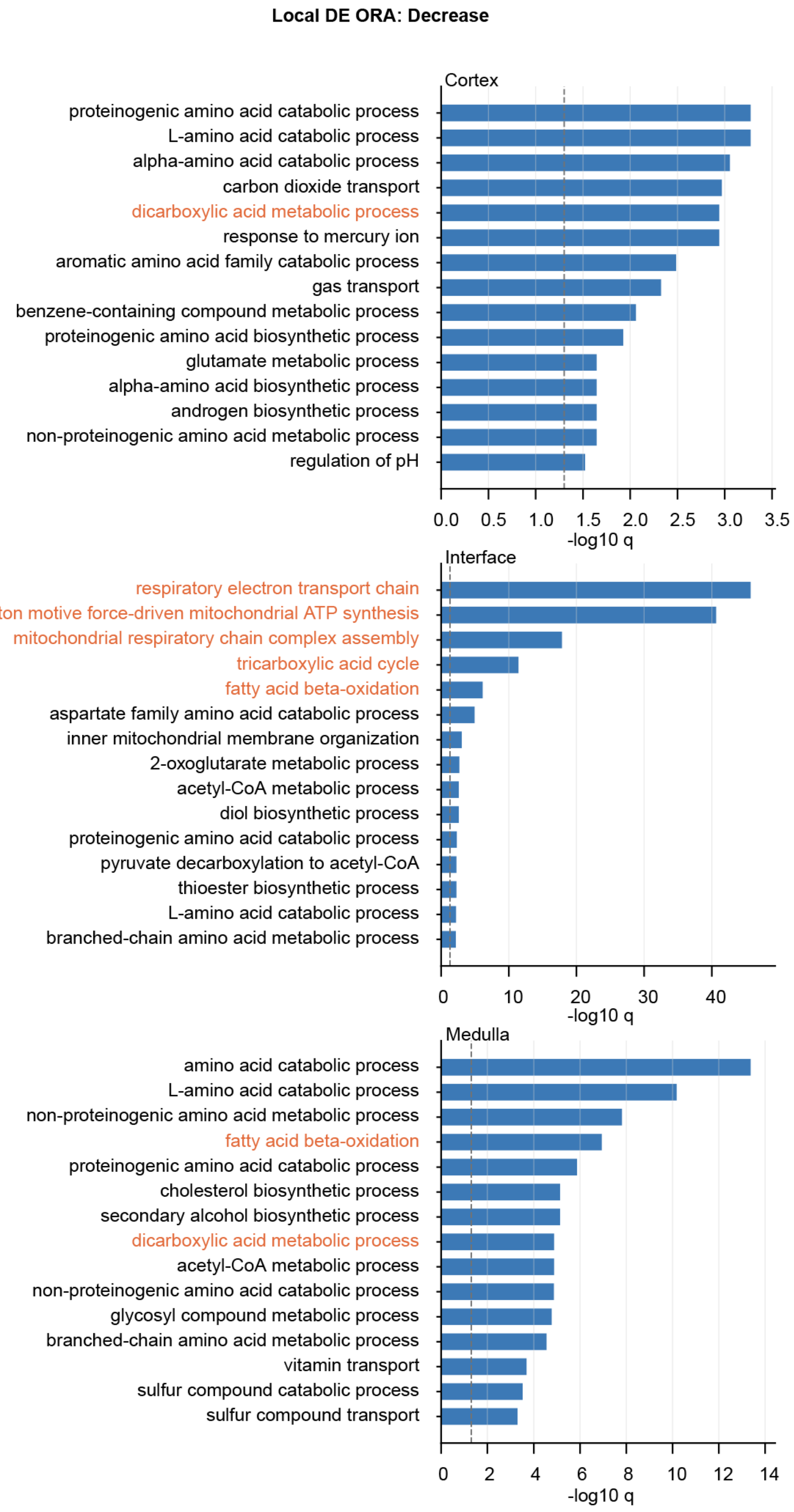


Fig. S35: **Local DE-region over-representation analysis for decreased expression in kidney injury.** Panels show the 15 highest-ranked Gene Ontology Biological Process terms for cortex, interface and medulla classes defined from direction-specific significant local-grid patterns after the gene-level $q_g^{\mathrm{ACAT}}$ eligibility screen. Bars indicate $-\log_{10} q$, dashed lines mark $q = 0.05$, and orange labels identify kidney-injury-related terms using the fixed descriptive rule applied after ranking. Highlighting affected neither term selection nor order.

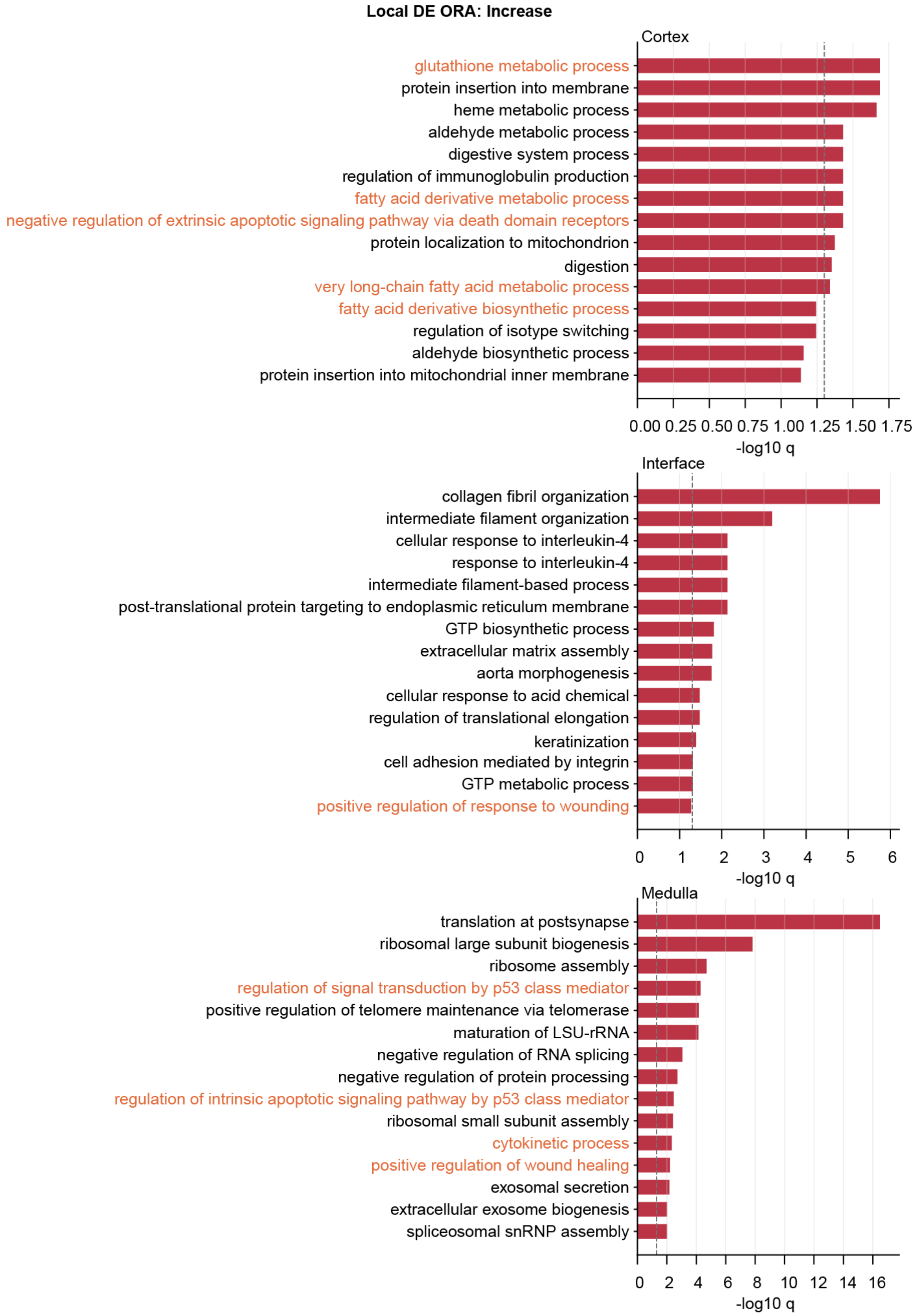


Fig. S36: **Local DE-region over-representation analysis for increased expression in kidney injury.** Panels show the 15 highest-ranked Gene Ontology Biological Process terms for cortex, interface, and medulla classes defined from direction-specific significant local-grid patterns after the gene-level $q_g^{\mathrm{ACAT}}$ eligibility screen; when fewer than 15 terms were returned, all available terms are shown. Bars indicate $-\log_{10} q$, dashed lines mark $q = 0.05$, and orange labels identify kidney-injury-related terms using the fixed descriptive rule applied after ranking. Highlighting affected neither term selection nor order.

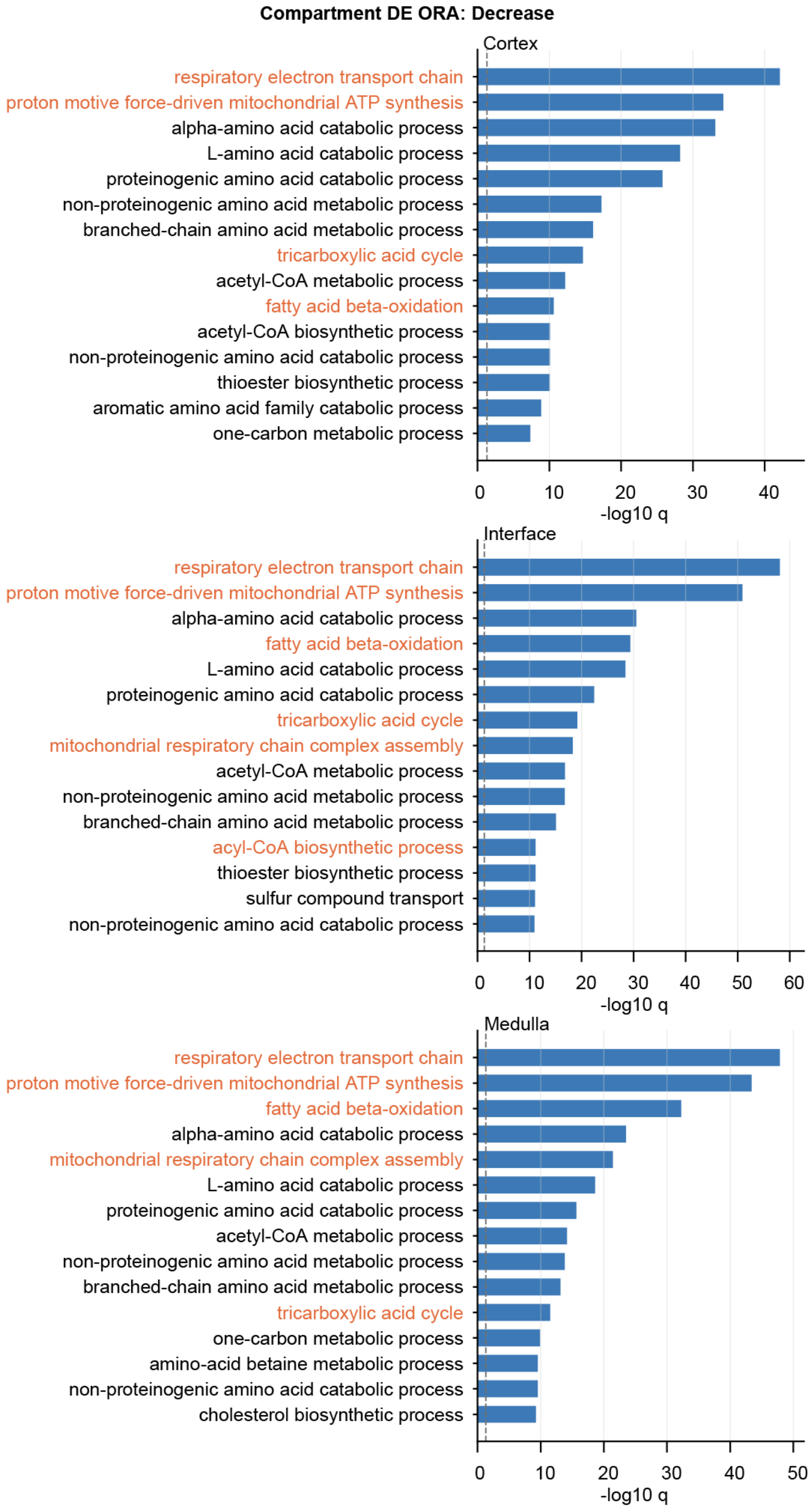


Fig. S37: **Compartment-level over-representation analysis for decreased expression in kidney injury.** Panels show the 15 highest-ranked Gene Ontology Biological Process terms obtained after direct injured-versus-normal differential expression within cortex, interface and medulla. Bars indicate $-\log_{10} q$, dashed lines mark $q = 0.05$, and orange labels identify kidney-injury-related terms using the fixed descriptive rule applied after ranking. Highlighting affected neither term selection nor order.

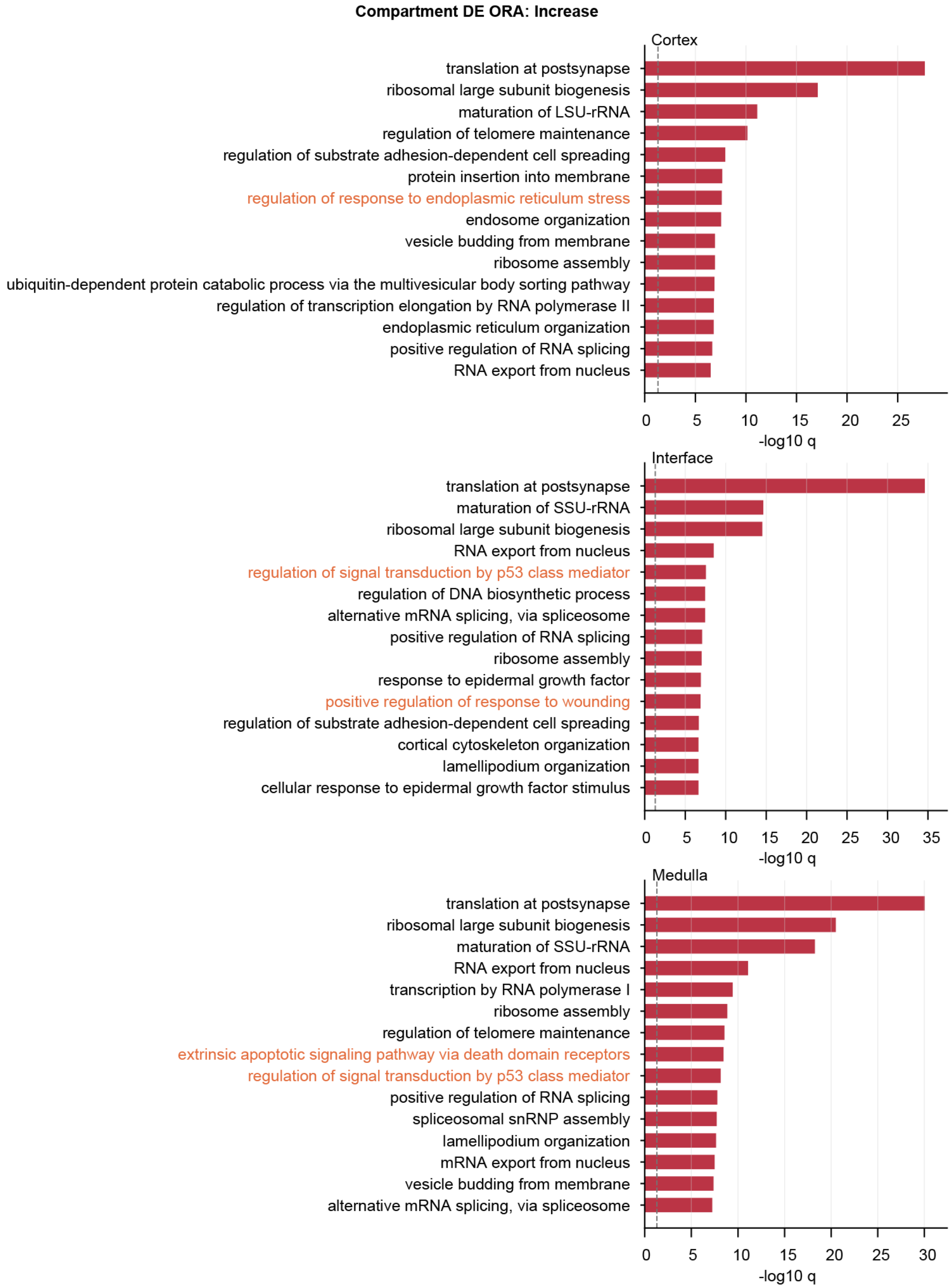


Fig. S38: **Compartment-level over-representation analysis for increased expression in kidney injury.** Panels show the 15 highest-ranked Gene Ontology Biological Process terms obtained after direct injured-versus-normal differential expression within cortex, interface and medulla. Bars indicate $-\log_{10} q$, dashed lines mark $q = 0.05$, and orange labels identify kidney-injury-related terms using the fixed descriptive rule applied after ranking. Highlighting affected neither term selection nor order.
